# Multiplex PCR reveals unique trends in pathogen and parasitoid infestations of alfalfa leafcutting brood cells

**DOI:** 10.1101/2022.01.09.475550

**Authors:** Justin Clements, Maggie Haylett, Brenda Nelson, Silas Shumate, Nicole Young, Benjamin Bradford, Doug Walsh, Kurt Lamour

## Abstract

The alfalfa leafcutting bee *Megachile rotundata* Fabricius (HYMENOPTERA: Megachilidae) is an important pollinator for multiple agricultural seed commodities in the United States. *Megachile rotundata* is a solitary bee that forms brood cocoons where its larvae can develop. During the developmental stages of growth, broods can be preyed upon by multiple different fungal and bacterial pathogens and insect predators and parasitoids, resulting in the loss of the developing larvae. Larval loss is a major concern for alfalfa (*Medicago sativa* L.) seed producers because they rely on pollinator services provided by *Megachile rotundata* and reduced pollination rates result in lower yields and increased production costs. In the present study, we examined the taxonomic composition of organisms found within *M. rotundata* brood cells using a multiplex PCR assay which was developed for the detection of the most common bacterial, fungal, and invertebrate pests and pathogens of *M. rotundata* larvae. Known pests of *M. rotundata* were detected, including members of the fungal genus *Ascosphaera*, the causative agent of chalkbrood. Co-infection of single brood cells by multiple *Ascosphaera* species was confirmed, with potential implications for chalkbrood disease management. The multiplex assay also identified DNA from more than 2,400 total species including multiple new predators and pathogenetic species not previously documented in associated with *M. rotundata* brood cells.

## Introduction

The alfalfa leafcutting bee *Megachile rotundata* Fabricius (HYMENOPTERA: Megachilidae) is an important pollinator for multiple agricultural commodities in the United States including alfalfa seed, canola, melons and carrots. One of the primary agricultural commodities that uses *M. rotundata* for pollination is alfalfa (*Medicago sativa* L.) seed production, which provides the germplasm for alfalfa used for hay [1–5]. While alfalfa seed production is predominately limited to the Pacific Northwest region of the United States, alfalfa hay production encompasses more than 17 million acres with a production value of more than 8.8 billion dollars annually in the United States [5–6] and is only possible with the production of high-quality seed. An insect pollinator is necessary to generate a seed crop in alfalfa, as alfalfa is self-incompatible and unable to self-pollinate, and alfalfa seed producers rely on *M. rotundata* and the ground dwelling alkali bee, *Nomia melanderi*, to fill this role [2,4,7–9]. The reliance on these bees has resulted in *M. rotundata* being one of the most heavily managed pollinator species in the world [9].

*Megachile rotundata* is a solitary bee that constructs a nest (cocoon) for oviposition and larval development. A single female *M. rotundata* will construct a nest that consists of individual brood cells that are lined with cut leaves (cocoon) [9,10–11] and filled with pollen to provide nutrition for the developing larva. After construction, a female leafcutting bee will lay a single egg within the cell and seal the egg within the cocoon to undergo larval stage development and pupation into the adult stage [9]. This cocoon construction requires multiple trips to flowering plants and makes alfalfa leafcutting bees a highly effective pollinator species [1,9]. As with most pollinating species, including honey bees (*Apis sp.*), mason bees (*Osmia bicornis*), and blue orchard bees (*Osmia lignaria*), *N. melanderi* and *M. rotundata* are preyed upon by predatory insects and play host to numerous fungal pathogens [12–16]. The presence of invertebrate pests and fungal pathogens within *M. rotundata* brood cells is a major concern for alfalfa seed producers, as they can cause reduced pollination efficiency by resulting in brood loss. Multiple pathogens and parasitoids can infect a developing larva, often resulting in the loss of brood. Common fungal pathogens include multiple *Ascosphaera* species, which cause chalkbrood. Alfalfa seed growers are predominately concerned with the presence of *Ascosphaera aggregata* within brood cells, as it is currently thought to be the major *Ascosphaera* species which results in *M. rotundata* brood loss [17]. However, multiple different *Ascosphaera* species have been detected in *M. rotundata* nest cells. For example, *A. subglobosa, A. acerosa, A. asterophora, A. larvis, A. pollenicola*, and *A. proliperda* [18-21] have all been found in *M. rotundata* brood cells and can cause the chalkbrood phenotype. Broods may also be attacked by other insects, including parasitic wasp species (such as *Monodontomerus obscurus, Leucospis affinis, Pteromalus venustus*, and *Sapyga pumila* [22]), nest destroying beetles (including *Tribolium audax, Tribolium brevicornis*, and *Trichodes ornatus* [22]), and cuckoo bees (*Epeoloides pilosula*) [22], all of which feed on developing larvae and further decimate producers’ bee stocks. The presence of these pathogens and parasitoids can significantly impact the pollination of the alfalfa seed crop and subsequently decrease the crop yield by reducing pollinator populations and efficiency. Additionally, these pathogens and predators can reproduce within grower stocks, bee boards, and housing, and if not controlled, can result in a high abundance of dead bee larvae. With these bees being heavily managed, sold, and transported throughout growing regions, this can cause significant concern for growers.

In order to reduce brood loss, growers use a combination of disinfectants and lures to protect brood from different pathogens and parasitoids [23–26]. Growers monitor the presence of these pathogens using X-ray (Faxitron) imaging as a diagnostic technique [27]. With current detection practices limited to X-ray diagnostics and few published polymerase chain reaction (PCR) amplifications of *Ascosphaera* species [28], we set out to examine the taxonomic makeup of *M. rotundata* brood cells through a multiplex deep sequencing PCR reaction. The multiplex was designed to amplify short sequence reads (150-200bp) to known and unknown pathogen, predators, and *Ascosphaera* species. If primers are designed correctly, multiplex technology can provide millions of short sequences which can be mapped back to reference sequences revealing the identity and composition of all species within a sample. As such, this technique can provide a glimpse into the complex biological niche of the *M. rotundata* brood cell.

In the current investigation we examined the taxonomic makeup of *M. rotundata* brood cells to determine if we could generate a multiplex PCR reaction that could confirm the presence of known pathogens and potentially identify new pathogens present in *M. rotundata* brood cells. Currently, diagnostic techniques rely on the visual assessment of pathogen and predator infection using Faxitron (X-ray) imaging. Pathogens and predators can be challenging to identify and classify using imaging alone. In order to develop a new, more definitive, and more sensitive method to detect infection, we explored the use of an Illumina-based DNA multiplex PCR assay. To develop this assay, we classified the presence of different pathogens and predators within 4 populations of *M. rotundata* cells encompassing 4 grower populations from Idaho and Washington using Faxitron imaging. We extracted total DNA from a set of 200 *M. rotundata* brood cells encompassing healthy brood cells, pathogen/predator infested brood cells, and brood cells with unknown infection status. Using an Illumina-based multiplex PCR assay we examined taxonomic makeup of the brood cells, including known and novel pathogens and predators. The analysis resulted in the identification of multiple, new potential pathogens of *M. rotundata* bee cells. Further, the multiplex analysis provided insight into which species of *Ascosphaera* can infect *M. rotundata* brood cells, including the detection of multiple *Ascosphaera* species co-infections within individual brood cells.

## Materials and methods

### Data availability

All relevant data are contained within the paper and its supporting information files.

### Ethical statement

This article does not contain studies with any human participants and no specific permits were required for collection or experimental treatment of *Megachile rotundata* for the study described.

### *Megachile rotundata* sample collection

*Megachile rotundata* samples were acquired from growers as part of the Parma Cocoon Diagnostic Laboratory service (Parma, Idaho). This service examines bee health for alfalfa seed growers through Faxitron diagnostics. The Parma Cocoon Diagnostic Laboratory (PCDL) is an extension-oriented service that classifies the proportion of pathogen and parasitoid infected *M. rotundata* brood cells submitted by growers. PCDL uses X-ray imaging to visually classify brood cell fungal pathogens including *Ascosphaera aggregata* and *Ascosphaera larvis*, insect parasites including imported chalcid wasps (*Monodontomerus obscurus*), cuckoo bees (*Epeoloides pilosula*), woodboring chalcid wasps (*Leucospis affinis*), long-tongued blister beetles (also known as sunflower beetles, *Nemognatha lutea*), Canadian chalcid wasps (*Pteromalus venustus*), and red-marked sapygids (*Sapyga pumila*), and predators/nest destroyers including American black flour beetles (*Tribolium audax*), giant flour beetles (*Tribolium brevicornis*), and checkered flower beetles (*Trichodes ornatus*). Brood cells are shipped directly to the PCDL from commercial growers as a loose aggregate. Samples used within this experiment were acquired from multiple growers from the Pacific Northwest. Grower information cannot be disclosed as part of a confidentiality agreement. Upon arrival, samples were transferred to a sterile incubator held at 27 ° C. Samples were used in both Faxitron and multiplex DNA analysis.

### Faxitron analysis

Using sterile laboratory techniques, individual brood cells from each grower sample were lined up on a piece of contact film. A total of five, 10-gram samples for each population were X-rayed using a Faxitron (Model 42855A) machine. Samples were X-rayed using Kodak Industrex-M radiographic film. Samples were processed at 20kVp for a 1-minute exposure time. Samples were visually analyzed using defined standards for fungal pathogens, invertebrate pests, dead larvae, pollen balls, and healthy larvae. After the brood cells were classified, they were removed from the sticky paper using sterilized tweezers and placed in sterile 1.5ml microcentrifuge tubes (visual classification of cells can be found in **Supplemental File S1**). To minimize any contamination within samples, all laboratory equipment was sterilized before and after contact with brood cells with 95% ethanol and a Bunsen burner.

### *Megachile rotundata* brood cell DNA extraction

DNA was extracted from individual cells using a modified CTAB extraction protocol. Individual brood cells were placed in 2ml DNAse/RNAse-free homogenate tubes (Biospec, OK, USA) with a single sterilized 6.4mm diameter glass bead (Biospec, OK, USA). Seven hundred and fifty μl of CTAB extraction buffer was added to each tube (OPS diagnostics, NJ, USA). Samples were homogenized for 2 minutes in a Mini-beadbeater-16 (Biospec, OK, USA). Tubes with homogenate were incubated at 60°C in a water bath for 30 minutes. Following the incubation period, samples were centrifuged for 10 minutes at 14,000 x g and the supernatant was transferred to new 1.5ml tubes. Five μl of RNase solution A (20mg/ml, Fisher Scientific, MA, USA) was added and incubated at 37°C for 20 minutes. Three hundred μl of chloroform/isoamyl alcohol (24:1) was then added to each sample and vortexed for 5 seconds, then centrifuged for 1 minute at 14,000g to separate the phases. The chloroform/isoamyl alcohol step was conducted twice. The upper aqueous phase was transferred to a new 1.5ml microcentrifuge tube. DNA was precipitated by adding 500μl cold isopropanol. Samples were left for 12 hours at −20 ° C. Samples were centrifuged at 14,000g for 10 minutes to pelletize DNA. Supernatant was decanted without disturbing the pellet and was subsequently washed with 1ml of ice-cold 70% ethanol, and the samples were vortexed and centrifuged at 14,000g for 10 minutes. Ethanol was decanted and excess ethanol was removed from the pellet with a pipettor. Samples were air dried in a sterile PCR cabinet for 15 min. DNA was dissolved in 100μl RNase/DNase-free H_2_O. DNA concentrations were determined using a Nanodrop 2000 (Thermo Fisher Scientific, MA, USA). Samples were stored at −20 C until multiplex processing.

### *Megachile rotundata* brood cell multiplex PCR and data analysis

Total DNA from each sample was sent to Floodlight Genomics LLC (TN, USA). Floodlight Genomics used an optimized Hi-Plex approach to amplify targets in a single multiplex PCR reaction [29]. Primers were designed to amplify both known fungal pathogens (*A. aggregata* and *A. larvis*, as they are the two species of *Ascosphaera* screened by the Parma Diagnostic Laboratory), known invertebrate pests, and unknown bacteria (**Supplemental File S2**). The primers were also intentionally designed over large regions of genes used in phylogenetic investigations including cytochrome C oxidase, 18S ribosomal RNA, 28S ribosomal RNA and bacterial V regions to be able to examine the overall taxonomic makeup of the *M. rotundata* brood cell. The sample-specific barcoded amplicons were sequenced on the Illumina HiSeq X platform according to the manufacturer’s directions. Floodlight Genomics delivered sample-specific short raw DNA sequence reads as FASTQ files. Annotation of the raw reads was performed with Geneious Bioinformatics Software (Auckland, New Zealand). Reads were grouped by both individual brood cell and as an aggregate of all brood cells. Raw reads were mapped to reference sequences at 100% stringency for classification. Raw reads were also assembled (as an aggregate of all samples) into biological contigs using Geneious (**Supplemental File S3**). Assembled contigs were uploaded into Blast2go (BioBam Bioinformatics, MA, USA) for analysis. The National Center for Biotechnology Information (NCBI) nucleotide database was downloaded in May of 2021. The database was uploaded into Blast2go to generate a refence database. Contigs were blasted against the reference database with a E value cut of 10^-10^. Taxonomic identification of contigs was determined using R version 4.0.4 and the *taxonomizr* package [30]. Phylogenic trees were generated based on NCBI scientific name through phyloT v2 database 2021.1 [31].

### *Ascosphaera* DNA extraction

An additional set of *M. rotundata* brood cells were classified as being infected with *Ascosphaera* (via X-ray diagnostics) and were placed within sterile 1.5ml centrifuge tubes. Brood cells were sterilized prior to DNA extraction by washing with a sterilization solution designed to sterilize and remove contaminants from the outside of the *M. rotundata* brood cells. The sterilization solution comprised 5% 190 proof Ethanol, 1% Tween 20 (ThermoFisher Scientific, Waltham, MA, USA), and 0.1% D-256 (Venco, St. Joseph, MO) and was generated in DNase/RNase-free water (ThermoFisher Scientific, Waltham, MA, USA). Aliquots of 1ml of sterilization solution were placed into 1.5 ml microcentrifuge tubes, and new aliquots were used for every brood cell. Brood cells were dipped in the sterilization solution aliquot using forceps for 30 seconds moving up and down within the solution. Cells were then washed with DNase/RNase-free water to remove any residues. Sterilized brood cells were then placed in sterile 2ml DNAse/RNAse-free homogenate tubes (Biospec, OK, USA) and allowed to completely dry before DNA extraction in a sterile PCR cabinet. DNA was extracted from individual cells using a DNeasy Plant Pro Kit (Qiagen, Hilden, Germany). The DNeasy Plant Pro Kit requires limited steps compared to the CTAB method, resulting in less chance to contaminate the purified DNA. DNA concentration was determined using a Nanodrop 2000. Samples were stored at −20°C until PCR and multiplex processing.

### *Ascosphaera* traditional PCR

To confirm the presence of *Ascophaera* from DNA isolations, a PCR amplification was conducted on a subset of the DNA extractions using primers designed by James and Skinner 2005 [28]. Three PCR reactions were conducted to amplify all *Ascosphaera* species, group 1 (*A. aggregata* and *A. proliperda*), and group 2 (*A. columbrina, A. variegata, A. larvis*, and *A. pollenicola*) (Primers can be found in **Supplemental file S2**). Specifically, 25μl PCR was conducted with GoTaq Green Master Mix (Promega Corporation, Madison, WI). Reaction conditions included 240s at 94°C as the initial denaturing step, followed by 40 cycles of 60s at 94°C for denaturation, 60s at 63°C for annealing, 60s at 72°C for extension and a final extension of 300s at 72°C. PCR amplification products were separated on a 1.5% agarose gel to confirm presence of corresponding DNA fragments.

### *Ascosphaera* multiplex PCR

Twenty IST1-5.8S-IST2 sequences from Anderson et al. 1998 [32] were downloaded from NCBI, the IST2 region of the sequences was aligned using Geneiuos Prime tree builder global alignment (**Figure 1**). From the alignment we determined that the IST2 region had significant sequence differences that could be used to classify *ascosphaera* species. Primers were designed to amplify the IST2 region of *Ascosphaera* (**Supplemental File S2**). Total DNA from 38 *Ascosphaera* samples were sent to Floodlight Genomics LLC. Floodlight Genomics used an optimized Hi-Plex approach to amplify targets in a single multiplex PCR reaction [29]. The sample-specific barcoded amplicons were sequenced on the Illumina HiSeq X platform according to the manufacturer’s directions. Floodlight Genomics delivered sample-specific short raw DNA sequence reads as FASTQ files. Annotation of the raw reads was performed with Geneious Bioinformatics Software (Auckland, New Zealand). Reads were treated as individual cells and mapped to reference sequences with a minimum overlap identity of 95% and reads only mapping to the single best match of the 20 reference sequences was allowed. A sample was considered infected with an *Ascosphaera* species if there were more than 1000 sequence hits. The high threshold of 1000 was used to eliminate the chance that a cell was not completely sterilized before extraction.

**Figure 1.**
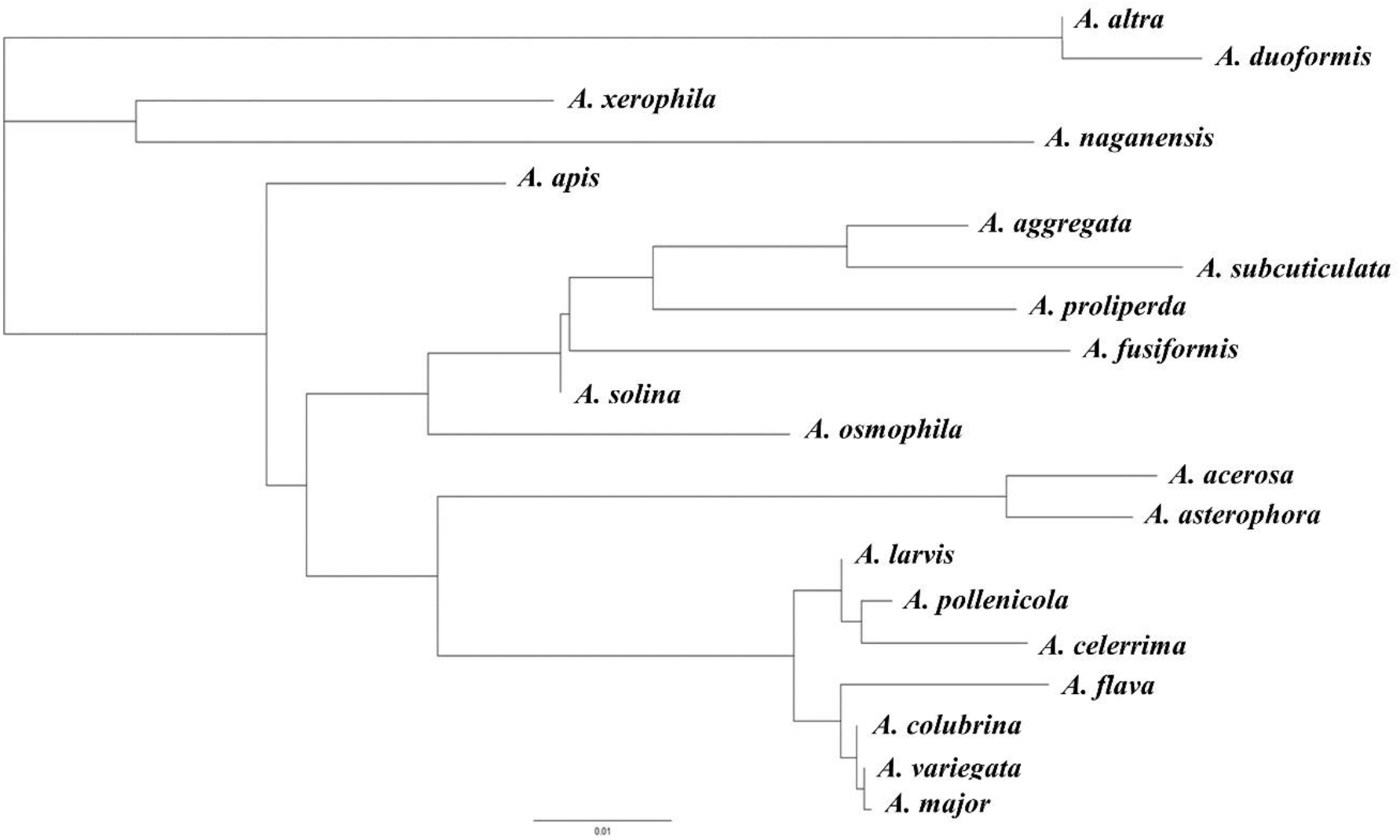
IST2 sequences alignment from *ascosphaera* species obtained from Anderson et al. 1998 [32]

## Results

### *Megachile rotundata* brood cell multiplex PCR

A total of 21,974,525 short Illumina reads were acquired from Floodlight Genomics and mapped to a set of reference sequences to known pathogens and parasitoids of *M. rotundata* cells. In total 607,769 reads were successfully mapped to a reference sequence (100% overlapping identity). Only *A. aggregata, A. larvis, T. ornatus*, and *T. brevicornis* were successfully identified within the samples using this method. *A. larvis* and *A. aggregata* were found within samples from each population analyzed. *T. ornatus* was classified in population 1 and *T. brevicornis* was classified in population 2. We noted that almost all samples had DNA which corresponded to both *A. aggregata* and *A.larvis.* This was most likely due to how samples were processed and shipped as aggregates, resulting in both fungal and invertebrate DNA contamination throughout the entire sample. While within regional samples it is possible to determine taxonomic classification, it was not possible within individual cells for this reason.

### BLAST assembly

To further explore the taxonomic makeup of the *M. rotundata* brood cells, we aggregated all reads into a single sample. A total of 51,056 contigs were assembled using Geneious Prime. The assembly was conducted using standard cutoffs and conditions provide by the software. The contigs were compared against the entire NCBI nucleotide database to investigate taxonomic composition of *M. rotundata* cells and explore new predators and pathogens of *M. rotundata* cells. This composition was conducted as an aggregate of all brood cells to examine the general makeup of these complex biological systems. In total there were 2,438 different species classified in the analysis (**Supplemental File S4**), including 13 archaeal, 1,716 bacterial, and 709 eukaryotic contigs which could be differentiated into biological classes (**Supplemental Figure S5**). The predominant orders of insects were Hymenoptera and Coleoptera (**Figure 2**). Contigs within the class Insecta belonged to multiple different parasitic wasps, bees, and nest destroying beetles. These include various parasitic wasp taxa (Chalcididae, Eupelminae, *Melittobia spp., Mesopolobus spp., Pteromalus spp., Torymus spp*., and *Pediobius spp*.), the cuckoo bee (*Coelioxys pieliana*), and predator/nest destroyers *Tribolium audax* and *Tribolium brevicornis*. Additional parasitic wasp taxa that infest aphids and other non-*M. rotundata* insects were also observed, including *Diaeretiella spp., Encarsia spp., Lysiphlebus spp*., and *Pauesia spp*.

**Figure 2.**
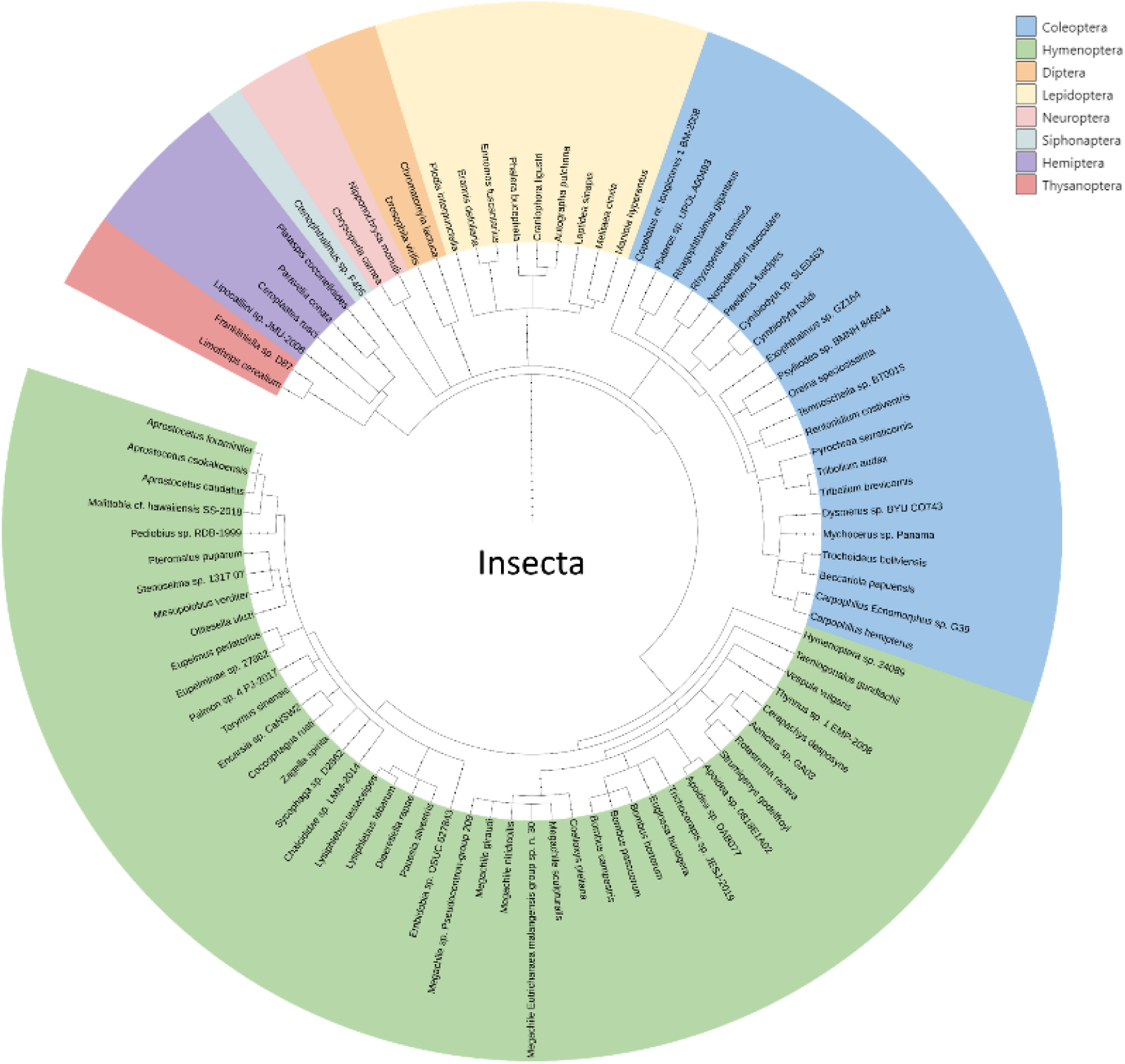
Insect species and orders of contigs found within *M. rotundata* brood cells

We further investigated the division of fungal ascomycetes found within *M. rotundata* brood cells. Members of the fungal division Ascomycota (**Figure 3**) are known as sac fungi and include multiple known pathogens of bees within the class Eurotiomycetes. The contig assembly revealed multiple different known human and insect pathogens including *Aspergillus* and 20 *Ascosphaera* species. *Aspergillus* species, which are known mammalian pathogens, are most likely a product of decomposing plant tissue within the brood cocoon. *Ascosphaera* species are known pathogens of insects and could be predating on and reproducing within the brood cells.

**Figure 3.**
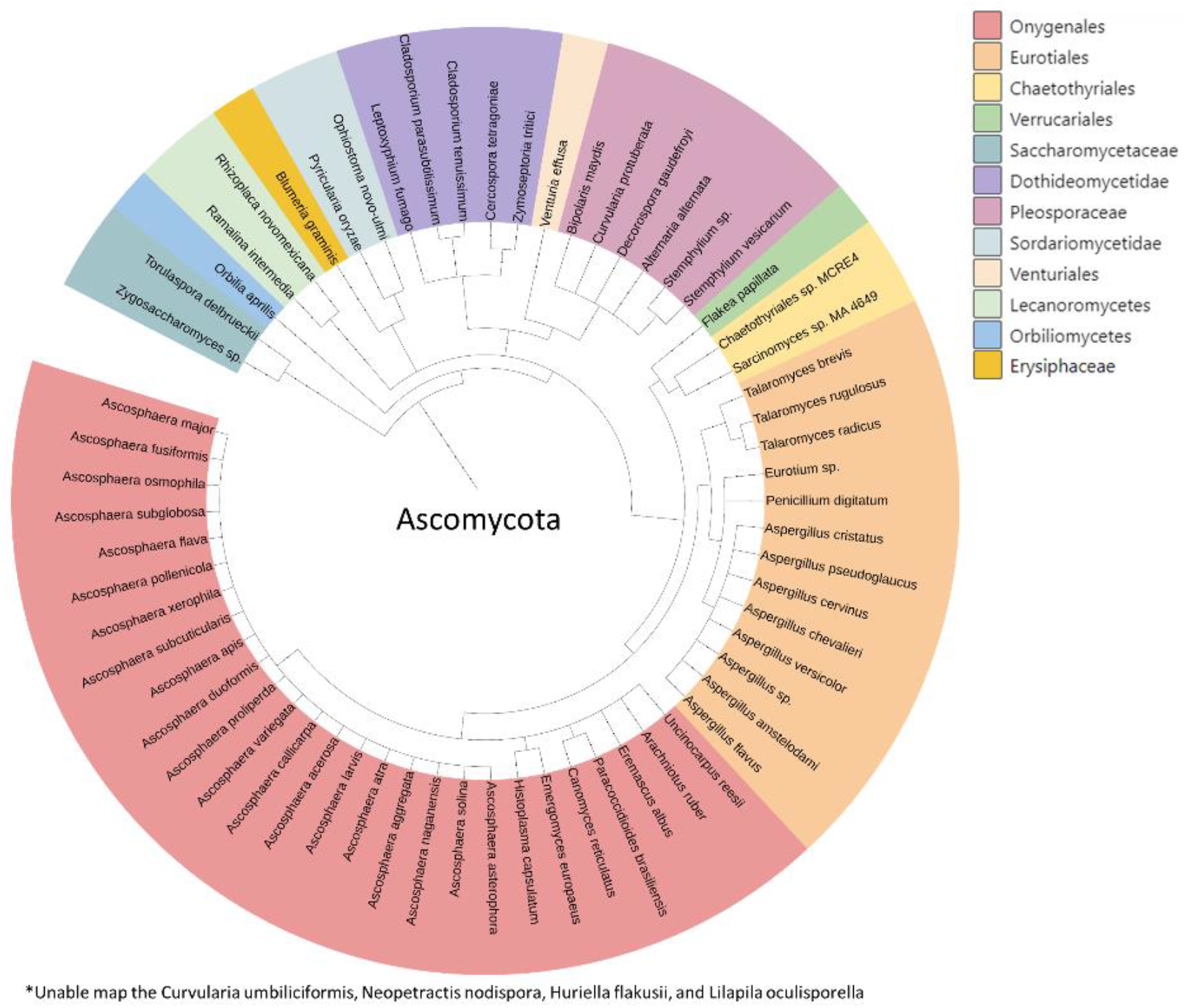
Fungi division Ascomycota designation of contigs found within *M. rotundata* brood cells

### *Ascosphaera* data analysis

PCR revealed the presence of co-infected of *M. rotundata* cells by multiple different *Ascosphaera* species. A subset of 18 *Ascosphaera* DNA extractions were PCR amplified to confirm the presence of *Ascosphaera* within the *M. rotundata* sample. For each *Ascosphaera* DNA extraction, three PCR amplifications were conducted. The first PCR amplification used non-specific primers which amplify most *Ascosphaera* species [28], which revealed that 13 out of the 18 samples were positive for *Ascosphaera* (**Figure 4a**). The second amplification which was specific for the amplification of *A. columbrina, A. variegate, A. larvis*, and *A. pollenicola* revealed that that 12 of the 18 samples were positive for one or more of these species (**Figure 4b**) and the final PCR amplification which was specific to *A. aggregata* and *A. proliperda* revealed 13 out of the 18 samples were positive for one or both of these species (**Figure 4a**). The PCR amplification also indicated that most samples (12 in total) had the presence of multiple different *Ascosphaera* species. The size of the bands from the second amplification also indicate that the primers may not only be specific to *Ascosphaera* for this amplification. A negative control was run with all PCR amplification and is depicted as the last well in each image, which had no amplification.

**Figure 4.**
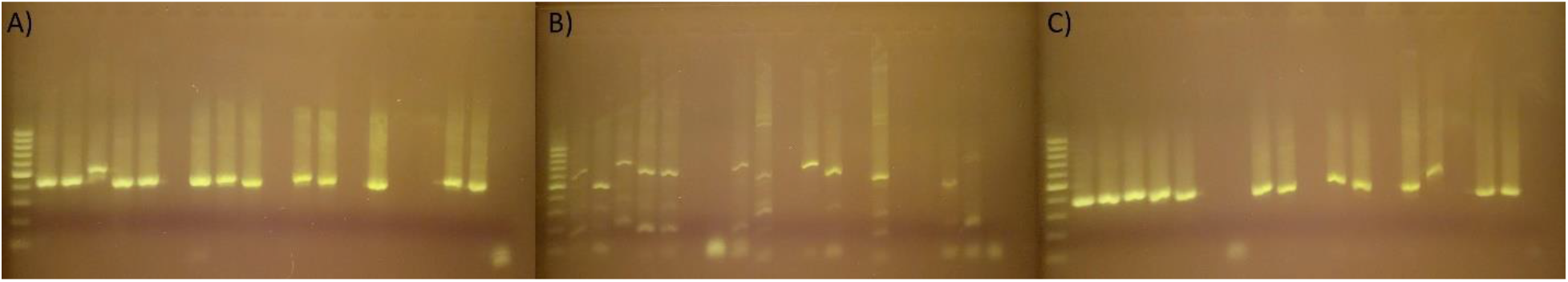
Amplification of *Ascosphaera* from *M. rotundata* cells on a 1.5% agarose gel with a 1kb DNA marker A) general *Ascosphaera* amplification, B) *A. columbrina, A. variegate, A. larvis*, and *A. pollenicola* specific PCR and C) *A. aggregata* and *A. proliperda*.

Deep sequencing of the IST2 region of *Ascosphaera* revealed the identification of the species of *Ascosphaera* infecting *M. rotundata*. The amplification of the IST2 region resulted in the amplification of *A. aggregata, A. atra, A. duoformis, A. naganensis, A. proliperda,* and *A. solina.* Only one sample was not co-infected with multiple different *ascosphaera* species which was only infected with *A. aggregata*. Further, *A. Larvis, A. acerosa, A. asterophora* was detected within the IST2 region amplification however were below the classified threshold for detection.

**Table 1.** Multiplex amplification of *M. rotundata* cells infested with *ascosphaera*

| Sample | *A. aggregata* |  | *A. atra* |  | *A. duoformis* |  | *A. naganensis* |  | *A. proliperda* |  | *A. solina* |  |
| --- | --- | --- | --- | --- | --- | --- | --- | --- | --- | --- | --- | --- |
|  | Reads | % of Reads | Reads | % of Reads | Reads | % of Reads | Reads | % of Reads | Reads | % of Reads | Reads | % of Reads |
| Cell 94 | 5820 | 100 | 0 | 0 | 0 | 0 | 0 | 0 | 0 | 0 | 0 | 0 |
| Cell 104 | 21782 | 85.44 | 0 | 0 | 0 | 0 | 3712 | 14.51 | 0 | 0 | 0 | 0 |
| Cell 129 | 16991 | 32.08 | 0 | 0 | 0 | 0 | 3081 | 5.81 | 25134 | 47.41 | 7766 | 14.64 |
| Cell 130 | 36120 | 81.74 | 0 | 0 | 0 | 0 | 5980 | 13.51 | 0 | 0 | 2090 | 4.72 |
| Cell 132 | 5564 | 25.69 | 0 | 0 | 0 | 0 | 1003 | 4.63 | 11613 | 53.55 | 3476 | 15.99 |
| Cell 136 | 15975 | 84.71 | 0 | 0 | 0 | 0 | 2884 | 15.22 | 0 | 0 | 0 | 0 |
| Cell 138 | 24325 | 85.32 | 0 | 0 | 0 | 0 | 4186 | 14.64 | 0 | 0 | 0 | 0 |
| Cell 197 | 0 | 0 | 46969 | 95.29 | 2321 | 4.70 | 0 | 0 | 0 | 0 | 0 | 0 |
| Cell 266 | 0 | 0 | 9321 | 55.75 | 5360 | 31.95 | 0 | 0 | 2038 | 12.13 | 0 | 0 |
| Cell 323 | 0 | 0 | 25708 | 8.47 | 2768 | 0.91 | 1311 | 0.43 | 266622 | 87.84 | 7123 | 2.35 |
| Cell 331 | 0 | 0 | 0 | 0 | 0 | 0 | 0 | 0 | 46261 | 96.36 | 1747 | 3.63 |
| Cell 360 | 0 | 0 | 0 | 0 | 0 | 0 | 1357 | 0.57 | 228693 | 96.30 | 7425 | 3.13 |
| Cell 364 | 11558 | 31.38 | 0 | 0 | 0 | 0 | 2005 | 5.44 | 19986 | 54.21 | 3284 | 8.89 |
| Cell 679 | 4696 | 67.37 | 0 | 0 | 0 | 0 | 0 | 0 | 1179 | 16.75 | 1095 | 15.52 |
| Cell 682 | 15333 | 55.77 | 1637 | 5.94 | 0 | 0 | 2381 | 8.64 | 5757 | 20.89 | 2385 | 8.65 |
| Cell 683 | 11330 | 51.78 | 5030 | 22.93 | 0 | 0 | 2012 | 9.16 | 1591 | 7.24 | 1918 | 8.73 |
| Cell 684 | 11778 | 22.80 | 33897 | 65.58 | 4051 | 7.83 | 1936 | 3.74 | 0 | 0 | 0 | 0 |
| Cell 685 | 2583 | 23.89 | 8227 | 75.94 | 0 | 0 | 0 | 0 | 0 | 0 | 0 | 0 |
| Cell 689 | 13910 | 65.87 | 2651 | 12.52 | 0 | 0 | 1789 | 8.44 | 0 | 0 | 2766 | 13.05 |
| Cell 691 | 25997 | 74.26 | 3500 | 9.98 | 1329 | 3.79 | 4182 | 11.92 | 0 | 0 | 0 | 0 |
| Cell 692 | 6645 | 76.84 | 0 | 0 | 0 | 0 | 0 | 0 | 0 | 0 | 2003 | 22.96 |
| Cell 693 | 13991 | 60.54 | 3758 | 16.22 | 1168 | 5.04 | 1965 | 8.47 | 0 | 0 | 2228 | 9.60 |
| Cell 698 | 18027 | 76.13 | 0 | 0 | 0 | 0 | 2465 | 10.38 | 0 | 0 | 3186 | 13.41 |
| Cell 724 | 17504 | 80.54 | 0 | 0 | 0 | 0 | 2513 | 11.52 | 0 | 0 | 1716 | 7.86 |
| Cell 733 | 7239 | 77.29 | 0 | 0 | 0 | 0 | 0 | 0 | 0 | 0 | 2127 | 22.52 |
| Cell 734 | 17893 | 66.09 | 0 | 0 | 0 | 0 | 2590 | 9.54 | 1389 | 5.12 | 5200 | 19.15 |
| Cell 754 | 8993 | 26.07 | 0 | 0 | 0 | 0 | 1075 | 3.11 | 18426 | 53.36 | 6008 | 17.37 |
| Total Reads per species | 308234 |  | 140698 |  | 16997 |  | 48427 |  | 628689 |  | 63543 |  |

## Discussion

The continued health and protection of *M. rotundata* is vital for the success of the alfalfa seed, alfalfa hay, and livestock industries. While *M. rotundata* larvae develop within their cocoon produced by the mother bee they can still be preyed upon by multiple different parasitoids, pathogens, and nest destroyers. Classification of the different pathogens, parasitoids, and predators has traditionally revolved around visual classifications and X-ray diagnostics. Within this investigation we set out to examine the taxonomic makeup of *M. rotundata* brood cells and developed a multiplex PCR assay to examine the different *Ascosphaera* species that can infect *M. rotundata* brood cells and classified multiple potential new pathogens using this technique. We also found that developing brood can be co-infected my multiple *Ascosphaera* species.

We examined cells for the presence of known parasites, predators, nest destroyers, and fungal pathogens by first compiling genomic sequences from NCBI based on Evan et al. 1980 [22], which provided current known parasitoids and pathogens of *M. rotundata* brood cells. From the sequences we designed PCR primers and conducted a multiplex PCR reaction to confirm and classify known pathogens, parasites, predators, and nest destroyers of *M. rotundata* [19] within our samples to better understand the taxonomic composition of *M. rotundata* brood cells. Approximately 600,000 reads were successfully mapped to compiled references sequences, and these reads revealed the presence of *A. aggregata, A. larvis, T. ornatus, and T. brevicornis* within our *M. rotundata* brood cell samples. After this mapping, there were over 21 million reads that remained unmapped. To explore unmapped reads and examine the taxonomic makeup of the *M. rotundata* cell, we chose to assemble the short reads into contigs and BLAST against the entire NCBI database.

The BLAST result revealed the diversity of the composition of *M. rotundata* cells, resulting in 2,438 different species classified. It is important to note that this investigation could not differentiate between environmental contaminants within the samples. Environmental DNA could have been picked up from the foraging mother *M. rotundata* from plant material as she provisioned her nest cells, including fecal material deposited by other insects. Understanding the limitations of this investigation, we chose to examine likely new fungal pathogens and insect predators and parasites. Within our BLAST investigation we noted the presence of more than 20 *ascosphaera* species. We further examined the presence of invertebrate pests and noted multiple species that have the potential to predate on insects, including bees. These included parasitic wasps, the cuckoo bee (*Coelioxys pieliana*) and predator/nest destroyers *Tribolium audax* and *Tribolium brevicornis*. However, with multiple species lacking sequence data in NCBI, it is hard to make a definitive identification, and an approach of this manner might be more useful to provide class and family identifications and generate direction for future investigations, including providing insight into future sequencing endeavors to produce more comprehensive data for taxonomic classification.

One interesting result from the BLAST analysis was abundance of different *Ascosphaera* species found associated with *M. rotundata* cells. To better classify the identity of the *Ascosphaera* species causing chalkbrood in *M. rotundata*, we decided to sequence the ITS2 region of the IST1-5.8S-IST2 nucleic acid sequence. Our findings suggest that *M. rotundata* brood cells can be infected by multiple different *Ascosphaera* species, including being co-infected by multiple *Ascosphaera* species at the same time. From this analysis we noted that most cells examined were co-infected with multiple different *Ascosphaera* species including, *A. aggregata, A. atra, A. duoformis, A. naganensis, A. proliperda*, and *A. solina*. We also noted the presence of *A. larvis, A. acerosa, A. asterophora* within the samples, however they were below the detection threshold cutoff. Most samples were infected with *A. aggregata*, which is the major concern for alfalfa seed growers. However, 5 of the samples had no *A. aggregata* detected within the samples (above the detection threshold) and were primarily infected with *A. atra* or *A. proliperda.* The findings suggest that other species of *Ascosphaera* besides *A. aggregata* could be contributing to chalkbrood within *M. rotundata*. Our study was limited to a subset of infected *M. rotundata* brood cells, and there is a possibility that other brood cells, particularly from different growing regions, could be infected with different *Ascosphaera* species which were not detected within this investigation. Further, the current genomic resources are limited regarding sequence data of *Ascosphaera* species, and we based our assessments of our IST2 multiplex reaction only on reference sequences from Anderson et al. 1998 [32] who sequenced different *Ascosphaera* species from pure cultures growing on slanted solid agar media. It is possible that our alignments may be matching to unclassified *Ascosphaera* species. This suggests the need for more in-depth genomic data for this agriculturally relevant species.

From our data, it is clear that multiple *Ascosphaera* species can infect *M. rotundata* cells, including simultaneous co-infection. More research should be conducted to determine how the interactions between different *Ascosphaera* species may be affecting their pathogenicity.

This investigation revealed the complex taxonomic diversity of *M. rotundata* brood cells, with more than 2,400 different known species found inside the brood cell including species of plants, fungi, bacteria, archaea and insects. We further identified multiple different species that have the potential to predate on *M. rotundata* that should be examined in more detail, including species of *Ascosphaera* that could cause the disease phenotype known as chalkbrood. Pollinators will continue to be a vital resource for alfalfa seed producers so the agricultural community and growers should be on the lookout for new and emerging pathogens and parasitoids of leafcutting bees. We hope this investigation has provided useful new methods and data to support that goal.

## Acknowledgements

This research was supported by funding from United States Department of Agriculture Alfalfa Pollinator Research Initiative grant number 58-2080-0-009 awarded to JC.

## Supporting information

**Supplemental File S1:** *Megachile rotundata* bee cells classification used in multiplex analysis

**Supplemental File S2:** *Megachile rotundata* cells Multiplex, Ascosphaera PCR, and Ascosphaera cells Multiplex primers

**Supplemental File S3:** *Megachile rotundata* bee contigs assembled by Geneious primer

**Supplemental File S4:** Taxonomic classification of contigs

**Supplemental File S5:** Taxonomic tree of class identification of *Megachile rotundata* bee cells

