## Supplemental Files for "Multiplex PCR reveals unique trends in pathogen and parasitoid infestations of alfalfa leafcutting brood cells"

**Supplemental File S1**: Leafcutting bee cell samples used in the multiplex analysis with visual diagnostic classifications conducted from X-ray analysis.

| Location | Sample Number | Faxitron Identification |
|---|---|---|
| Population 1 | 568 | Unknown |
| Population 1 | 569 | Unknown |
| Population 1 | 570 | Unknown |
| Population 1 | 571 | Unknown |
| Population 1 | 572 | Unknown |
| Population 1 | 573 | Unknown |
| Population 1 | 574 | Unknown |
| Population 1 | 575 | Unknown |
| Population 1 | 576 | Unknown |
| Population 1 | 577 | Unknown |
| Population 1 | 578 | Unknown |
| Population 1 | 579 | Unknown |
| Population 1 | 580 | Unknown |
| Population 1 | 581 | Unknown |
| Population 1 | 582 | Unknown |
| Population 1 | 583 | Control |
| Population 1 | 584 | Control |
| Population 1 | 585 | Control |
| Population 1 | 586 | Parasite |
| Population 1 | 587 | Parasite |
| Population 1 | 588 | Parasite |
| Population 1 | 589 | Parasite |
| Population 1 | 590 | Chalkbrood |
| Population 1 | 591 | Dead Larva |
| Population 1 | 592 | Chalkbrood |
| Population 1 | 593 | Parasite |
| Population 1 | 594 | Parasite |
| Population 1 | 595 | Dead Larva |
| Population 1 | 596 | Chalkbrood |
| Population 1 | 597 | Parasite |
| Population 1 | 598 | Chalkbrood |
| Population 1 | 599 | Parasite |
| Population 1 | 600 | Chalkbrood |
| Population 2 | 501 | Unknown |
| Population 2 | 502 | Unknown |
| Population 2 | 503 | Unknown |

| | | |
|---|---|---|
| Population 2 | 504 | Unknown |
| Population 2 | 505 | Unknown |
| Population 2 | 506 | Unknown |
| Population 2 | 507 | Unknown |
| Population 2 | 508 | Unknown |
| Population 2 | 509 | Unknown |
| Population 2 | 510 | Unknown |
| Population 2 | 511 | Unknown |
| Population 2 | 512 | Unknown |
| Population 2 | 513 | Unknown |
| Population 2 | 514 | Unknown |
| Population 2 | 515 | Unknown |
| Population 2 | 516 | Unknown |
| Population 2 | 517 | Unknown |
| Population 2 | 518 | Unknown |
| Population 2 | 519 | Unknown |
| Population 2 | 520 | Unknown |
| Population 2 | 521 | Unknown |
| Population 2 | 522 | Unknown |
| Population 2 | 523 | Unknown |
| Population 2 | 524 | Unknown |
| Population 2 | 525 | Unknown |
| Population 2 | 526 | Unknown |
| Population 2 | 527 | Unknown |
| Population 2 | 528 | Unknown |
| Population 2 | 529 | Unknown |
| Population 2 | 530 | Unknown |
| Population 2 | 531 | Control |
| Population 2 | 532 | Control |
| Population 2 | 533 | Control |
| Population 2 | 534 | Control |
| Population 2 | 535 | Control |
| Population 2 | 536 | Control |
| Population 2 | 537 | Control |
| Population 2 | 538 | Pollen Ball |
| Population 2 | 539 | Dead Larva |
| Population 2 | 540 | Dead Adult |
| Population 2 | 541 | Pollen Ball |
| Population 2 | 542 | Parasite |

| | | |
|---|---|---|
| Population 2 | 543 | Pollen Ball |
| Population 2 | 544 | Chalkbrood |
| Population 2 | 545 | Pollen Ball |
| Population 2 | 546 | Parasite |
| Population 2 | 547 | Dead Larva |
| Population 2 | 548 | Dead Adult |
| Population 2 | 549 | Chalkbrood |
| Population 2 | 550 | Pollen Ball |
| Population 2 | 551 | Parasite |
| Population 2 | 552 | Chalkbrood |
| Population 2 | 553 | Pollen Ball |
| Population 2 | 554 | Chalkbrood |
| Population 2 | 555 | Chalkbrood |
| Population 2 | 556 | Dead Larva |
| Population 2 | 557 | Pollen Ball |
| Population 2 | 558 | Dead Larva |
| Population 2 | 559 | Chalkbrood |
| Population 2 | 560 | Parasite |
| Population 2 | 561 | Dead Larva |
| Population 2 | 562 | Pollen Ball |
| Population 2 | 563 | Pollen Ball |
| Population 2 | 564 | Dead Larva |
| Population 2 | 565 | Chalkbrood |
| Population 2 | 566 | Dead Pupae |
| Population 2 | 567 | Chalkbrood |
| Population 3 | 401 | Unknown |
| Population 3 | 402 | Unknown |
| Population 3 | 403 | Unknown |
| Population 3 | 404 | Unknown |
| Population 3 | 405 | Unknown |
| Population 3 | 406 | Unknown |
| Population 3 | 407 | Unknown |
| Population 3 | 408 | Unknown |
| Population 3 | 409 | Unknown |
| Population 3 | 410 | Unknown |
| Population 3 | 411 | Unknown |
| Population 3 | 412 | Control |
| Population 3 | 413 | Control |
| Population 3 | 414 | Control |

| | | |
|---|---|---|
| Population 3 | 415 | Control |
| Population 3 | 416 | Control |
| Population 3 | 417 | Control |
| Population 3 | 418 | Control |
| Population 3 | 419 | Control |
| Population 3 | 420 | Control |
| Population 3 | 421 | Control |
| Population 3 | 422 | Parasite |
| Population 3 | 423 | Parasite |
| Population 3 | 424 | Parasite |
| Population 3 | 425 | Parasite |
| Population 3 | 426 | Parasite |
| Population 3 | 427 | Parasite |
| Population 3 | 428 | Parasite |
| Population 3 | 429 | Parasite |
| Population 3 | 430 | Parasite |
| Population 3 | 431 | Parasite |
| Population 3 | 432 | Parasite |
| Population 3 | 433 | Parasite |
| Population 3 | 434 | Parasite |
| Population 3 | 435 | Parasite |
| Population 3 | 436 | Parasite |
| Population 3 | 437 | Chalkbrood |
| Population 4 | 438 | Unknown |
| Population 4 | 439 | Unknown |
| Population 4 | 440 | Unknown |
| Population 4 | 441 | Unknown |
| Population 4 | 442 | Unknown |
| Population 4 | 443 | Unknown |
| Population 4 | 444 | Unknown |
| Population 4 | 445 | Unknown |
| Population 4 | 446 | Unknown |
| Population 4 | 447 | Unknown |
| Population 4 | 448 | Unknown |
| Population 4 | 449 | Unknown |
| Population 4 | 450 | Unknown |
| Population 4 | 451 | Unknown |
| Population 4 | 452 | Unknown |
| Population 4 | 453 | Unknown |

| | | |
|---|---|---|
| Population 4 | 454 | Unknown |
| Population 4 | 455 | Unknown |
| Population 4 | 456 | Unknown |
| Population 4 | 457 | Unknown |
| Population 4 | 458 | Unknown |
| Population 4 | 459 | Unknown |
| Population 4 | 460 | Unknown |
| Population 4 | 461 | Unknown |
| Population 4 | 462 | Unknown |
| Population 4 | 463 | Unknown |
| Population 4 | 464 | Control |
| Population 4 | 465 | Control |
| Population 4 | 466 | Control |
| Population 4 | 467 | Control |
| Population 4 | 468 | Control |
| Population 4 | 469 | Control |
| Population 4 | 470 | Control |
| Population 4 | 471 | Predator |
| Population 4 | 472 | Chalkbrood |
| Population 4 | 473 | Predator |
| Population 4 | 474 | Parasite |
| Population 4 | 475 | Chalkbrood |
| Population 4 | 476 | Chalkbrood |
| Population 4 | 477 | Chalkbrood |
| Population 4 | 478 | Chalkbrood |
| Population 4 | 479 | Pollen Ball |
| Population 4 | 480 | Chalkbrood |
| Population 4 | 481 | Parasite |
| Population 4 | 482 | Predator |
| Population 4 | 483 | Predator |
| Population 4 | 484 | Parasite |
| Population 4 | 485 | Parasite |
| Population 4 | 486 | Pollen Ball |
| Population 4 | 487 | Parasite |
| Population 4 | 488 | Pollen Ball |
| Population 4 | 489 | Pollen Ball |
| Population 4 | 490 | Predator |
| Population 4 | 491 | Parasite |
| Population 4 | 492 | Pollen Ball |

| | | |
|---|---|---|
| Population 4 | 493 | Parasite |
| Population 4 | 494 | Pollen Ball |
| Population 4 | 495 | Parasite |
| Population 4 | 496 | Chalkbrood |
| Population 4 | 497 | Chalkbrood |
| Population 4 | 498 | Pollen Ball |
| Population 4 | 499 | Parasite |
| Population 4 | 500 | Chalkbrood |

**Supplemental File S2**:

Primers and primer targets used in the multiplex amplification

| Primer set | Forward | Reverse | Target Amplicon | Target Species |
|---|---|---|---|---|
| 1 | CCCGCGAACGAAATAAATAGAG | AGCCACCTACGGTGCTCG | Microsatellite Loci for Varroa destructor | Varroa destructor |
| 2 | GCGCAAACTTAACGCTCG | TCAAGCCAGAGTGCTGCAG | Microsatellite Loci for Varroa destructor | Varroa destructor |
| 3 | ACTCCTACGGGGAGGCAGCA | ATTACCGCGGCTGCTGG | V3-V4 | Bacteria |
| 4 | AGCAGCCGCGGTAAT | AGGGTATCTAATCCT | V4-V5 | Bacteria |
| 5 | GAATTGACGGGGACCC | GGGTTGCGCTCGTTA | V6-V7 | Bacteria |
| 6 | GAATTGACGGGGGCCC | GGGTTGCGCTCGTTG | V6-V7 | Bacteria |
| 7 | GTACACACCGCCCGTC | AAGGAGGTGATCCAACCGCA | V8-V9 | Bacteria |
| 8 | GCACACACCGCCCGTC | AAGGAGGTGATCCAGCCGCA | V8-V9 | Bacteria |
| 9 | CGTCATTGCAACCCTCAAGC | AAGAACATGATTGATCTTGG | 18S ribosomal RNA gene | Ascosphaera aggregate |
| 10 | CGTCATTGCAACCCTCAAGC | AGGCCACAAGAGCGGAAGAC | 18S ribosomal RNA gene | Ascosphaera larvis |
| 11 | GTGAAGCGGCAAAAGCTCAA | GTCGAAGGAGCTCTACACGG | 5.8S ribosomal Spacer | Ascosphaera aggregate and Ascosphaera larvis |
| 12 | AACGTGTGTCTGTGCGG | CTTCGACTGGAGTTCGTTT | 5.8S ribosomal Spacer | Ascosphaera aggregate and Ascosphaera larvis |
| 13 | GGAGGATCATCCGTAGACCT | CCTGCTAGTACGGGTAGAGA | CO1 | Tribolium audax |
| 14 | CAGGATGAACTGTTTATCCACC A | TTGCAGTTAATAATATCGATCATG CT | CO1 | Pteromalus venustus |
| 15 | CTGGTTGAACTGTTTATCCCCC | TGCTCATACAAATAATGGAGTTCG G | CO1 | Trichodes ornatus |
| 16 | TCTGGAATTTCCTCTATTTTAGG TGC | ATTGCTCCAGCTAATACGGGT | CO1 | Epeoloides pilosula |
| 17 | ACATATTGCTGGTGTTTCATCAA | AAATAACAATATTGTAATAGCACC AGC | CO1 | Monodontomerus obscurus |
| 18 | ATCAACATACGACCTGCAGG | TAGGATCTCCACCTCCTGCA | CO1 | Nemognatha lutea |
| 19 | TCTTCATATTGCAGGAATTTCCT CA | GCACCAGCAAGAACAGGTAA | CO1 | Sapyga pumila |
| 20 | CCAGTTATAATAGGTGGATTTG GCA | CCAGTTCCTGTACCTGATCCA | CO1 | Leucospis affinis |
| 21 | CACAGTAGGAGGATTAACTGGG | ACATAGTGGAAATGTGCAACTACG | CO1 | Tribolium audax |
| 22 | CACAGTAGGAGGACTAACAGG AG | ACATAGTGGAAATGTGCAACTACA | CO1 | Tribolium brevicornis |
| 23 | TACTGTAGGGGGATTAACAG | ACATAATGAAAATGAGCTAC | CO1 | Monodontomerus obscurus |
| 24 | CCTAGTAGAACGTCGCGACC | TGACGCACACACATTCGAGA | 28S | Epeoloides pilosula |
| 25 | TCGTCTCGCTCGGATTTACG | CAATGACTCGCGCACATGTT | 28S | Leucospis affinis |
| 26 | AATGGTATACGGCCGCAGTC | CCCGAGAGTACCCAAAGCAG | 28S | Sapyga pumila |
| 27 | GTAGGACGTCGCGATCCTTT | CACTAGGTCCGATCGCCATC | 28S | Tribolium brevicornis |
| 28 | TCGAGCGACGGTTGATCATT | CCATCTGTCTACCGTCGAGC | 28S | Trichodes ornatus |

Primers for the amplification of ITS2

| Primer Name | Forward | Primer Name | Reverse |
|---|---|---|---|
| ITS2_FW1 | TCTGGTATTCCGGAGGGCATGC | ITS2_R1 | CGAAGCCCCATACGCTCGGGCAC |
| ITS2_FW2 | TCTGGTATTCCGGGGGGCATGC | ITS2_R2 | CAAAGCCCCATACGCTCGGGCAC |

| | | | |
|---|---|---|---|
| ITS2_FW3 | CCTGGTATTCCGGGGGGCATGC | ITS2_R3 | GTTCAAGTTCCATGCGCTCGGGCAC |
| | | ITS2_R4 | CTTCAAGGTCCATGCGCTCGGGCAC |
| | | ITS2_R5 | CGAAGCCCCATACGATCGGGCAC |
| | | ITS2_R6 | GAGAAAACCCATACGCTCGGGCAC |
| | | ITS2_R7 | CCACGAGAAGTTGGTATAGACCG |
| | | ITS2_R8 | CCACTGTAAAAACAAAAAAGAAC |
| | | ITS2_R9 | CCACTAGAAGTAAATGATG |
| | | ITS2_R10 | CCACGAGAAAAAGATGGTATCG |
| | | ITS2_R11 | CCACTGGAAGTAGTAGATGGTTG |
| | | ITS2_R12 | CCACAGAAAAAAAAATAGTTGACG |
| | | ITS2_R13 | CCACAGAAAAAAAATAATGGTTG |
| | | ITS2_R14 | CCACAGAAAACAATTGGTTGG |
| | | ITS2_R15 | CCACTAGATGTGATGATAGCTGG |
| | | ITS2_R16 | CCACAGAAAAAAATAATAGTTGG |
| | | ITS2_R17 | CCACAGAAAAAAATAATGGTTGGT |
| | | ITS2_R18 | CCACTAGAAGGTGTGGCAGTAG |
| | | ITS2_R19 | CCACTGGAGTAAAATAGGCCGTG |
| | | ITS2_R20 | CCACTGAGACGATGATTGATCTG |
| | | ITS2_R21 | CCACTGAAGATTGATCTGGGTTC |
| | | ITS2_R22 | CCACTGTTGAAAAGAACATG |
| | | ITS2_R23 | CCACTAGAAGTAAATTGGTTG |

Primers from James and Skinner 2005 used for the amplification of ascosphaera

| **Ascosphaera amplification** | **Forward** | **Reverse** |
|---|---|---|
| General ascosphaera primer | GCACTCCCACCCTTGTCTA | GAWCACGACGCCGTCACT |
| Group 1 primers | GTGCTTTCCCGGTACTCC | GAWCACGACGCCGTCACT |
| Group 2 primers | GCACTCCCACCCTTGTCTA | CCACAGAAAAAATAATAGTTGGTTCG |

**Supplemental File S4**: Taxonomic identification of contigs determined by taxonomizr R version 4.0.4

| Superkingdom | Phylum | Class | Order | Family | Genus | Species |
|---|---|---|---|---|---|---|
| Archaea | Euryarchaeota | Halobacteria | Halobacteriales | Halobacteriaceae | | uncultured haloarchaeon |
| Archaea | Euryarchaeota | Methanomicrobia | Methanosarcinales | Methanosarcinaceae | Methanosarcina | Methanosarcina mazei |
| Archaea | Crenarchaeota | Thermoprotei | Thermoproteales | Thermoproteaceae | Pyrobaculum | uncultured Pyrobaculum sp. |
| Archaea | Euryarchaeota | Halobacteria | Halobacteriales | Halobacteriaceae | Halobacterium | Halobacterium salinarum |
| Archaea | Euryarchaeota | Halobacteria | Halobacteriales | | | haloarchaeon 3A1-DGR |
| Archaea | Euryarchaeota | Halobacteria | Halobacteriales | | | haloarchaeon str. Nh.2 |
| Archaea | Euryarchaeota | Halobacteria | Haloferacales | Haloferacaceae | Haloplanus | Haloplanus rubicundus |
| Archaea | Euryarchaeota | Halobacteria | Haloferacales | Halorubraceae | Halorubrum | Halorubrum gandharaense |
| Archaea | Euryarchaeota | Methanobacteria | Methanobacteriales | Methanobacteriaceae | Methanobacterium | Methanobacterium bryantii |
| Archaea | Euryarchaeota | Methanobacteria | Methanobacteriales | Methanobacteriaceae | Methanobrevibacter | Methanobrevibacter sp. YE315 |
| Archaea | Euryarchaeota | Methanomicrobia | Methanomicrobiales | Methanocorpusculaceae | Methanocorpusculum | Methanocorpusculum labreanum |
| Archaea | Thaumarchaeota | Nitrososphaeria | Nitrososphaerales | Nitrososphaeraceae | Nitrososphaera | Candidatus Nitrososphaera evergladensis |
| Archaea | | | | | | uncultured archaeon |
| Bacteria | | | | | | uncultured bacterium |
| Bacteria | | | | | | uncultured rumen bacterium |
| Bacteria | Firmicutes | Bacilli | Lactobacillales | Lactobacillaceae | Apilactobacillus | Apilactobacillus bombintestini |
| Bacteria | Firmicutes | Bacilli | Lactobacillales | Lactobacillaceae | Apilactobacillus | Apilactobacillus ozensis |
| Bacteria | Proteobacteria | Gammaproteobacteria | Enterobacterales | Morganellaceae | Arsenophonus | Arsenophonus endosymbiont of Aphis craccivora |
| Bacteria | Proteobacteria | Alphaproteobacteria | Sphingomonadales | Sphingomonadaceae | Sphingomonas | Sphingomonas sp. PAMC26645 |
| Bacteria | Proteobacteria | Gammaproteobacteria | Enterobacterales | Enterobacteriaceae | Klebsiella | Klebsiella pneumoniae |
| Bacteria | Actinobacteria | Actinomycetia | | | | uncultured actinobacterium |
| Bacteria | Proteobacteria | Gammaproteobacteria | Enterobacterales | Erwiniaceae | Tatumella | Tatumella sp. TA1 |
| Bacteria | | | | | | bacterium |
| Bacteria | Firmicutes | Bacilli | Bacillales | Bacillaceae | Pradoshia | Pradoshia sp. D12 |
| Bacteria | Firmicutes | Bacilli | Lactobacillales | Lactobacillaceae | Fructobacillus | Fructobacillus fructosus |
| Bacteria | Cyanobacteria | | | | | uncultured cyanobacterium |
| Bacteria | Firmicutes | Bacilli | Lactobacillales | Lactobacillaceae | Apilactobacillus | Apilactobacillus micheneri |
| Bacteria | Firmicutes | Bacilli | Bacillales | Bacillaceae | Bacillus | Bacillus sp. (in: Bacteria) |
| Bacteria | Actinobacteria | Actinomycetia | Propionibacteriales | Nocardioidaceae | Nocardioides | Nocardioides sp. |
| Bacteria | Proteobacteria | Betaproteobacteria | Burkholderiales | Oxalobacteraceae | Massilia | Massilia flava |
| Bacteria | Proteobacteria | Gammaproteobacteria | | | | uncultured Gammaproteobacteria bacterium |
| Bacteria | | | | | | uncultured soil bacterium |
| Bacteria | Firmicutes | Bacilli | Lactobacillales | Lactobacillaceae | Apilactobacillus | Apilactobacillus kunkeei |

| | | | | | | |
|---|---|---|---|---|---|---|
| Bacteria | Firmicutes | Bacilli | | | | uncultured Bacilli bacterium |
| Bacteria | Firmicutes | Bacilli | Bacillales | Bacillaceae | Priestia | Priestia megaterium |
| Bacteria | Proteobacteria | Gammaproteobacteria | Enterobacterales | Morganellaceae | Arsenophonus | Arsenophonus endosymbiont of Cardiaspina fiscella |
| Bacteria | Proteobacteria | | | | | uncultured proteobacterium |
| Bacteria | Proteobacteria | Gammaproteobacteria | Pseudomonadales | Pseudomonadaceae | Pseudomonas | uncultured Pseudomonas sp. |
| Bacteria | Proteobacteria | Gammaproteobacteria | Enterobacterales | Bruguierivoracaceae | Sodalis | Sodalis secondary endosymbiont of Curculio hachijoensis |
| Bacteria | | | | | | uncultured endolithic bacterium |
| Bacteria | Firmicutes | Bacilli | Bacillales | Bacillaceae | Bacillus | Bacillus subtilis |
| Bacteria | Gemmatimonadetes | Gemmatimonadetes | Gemmatimonadales | Gemmatimonadaceae | Gemmatimonas | uncultured Gemmatimonas sp. |
| Bacteria | Firmicutes | Bacilli | Bacillales | Bacillaceae | Bacillus | Bacillus licheniformis |
| Bacteria | Proteobacteria | Gammaproteobacteria | Enterobacterales | Enterobacteriaceae | Escherichia | Escherichia coli |
| Bacteria | Proteobacteria | Gammaproteobacteria | Enterobacterales | Enterobacteriaceae | | endosymbiont of Columbicola elbeli |
| Bacteria | Acidobacteria | | | | | uncultured Acidobacteria bacterium |
| Bacteria | Proteobacteria | Alphaproteobacteria | Hyphomicrobiales | Methylobacteriaceae | Methylobacterium | Methylobacterium sp. |
| Bacteria | Proteobacteria | Alphaproteobacteria | Sphingomonadales | Sphingomonadaceae | Sphingomonas | Sphingomonas sp. |
| Bacteria | Proteobacteria | Gammaproteobacteria | Enterobacterales | Enterobacteriaceae | Salmonella | Salmonella enterica |
| Bacteria | Firmicutes | Clostridia | Eubacteriales | Clostridiaceae | Clostridium | Clostridium sp. |
| Bacteria | Firmicutes | Clostridia | Eubacteriales | | | uncultured Eubacteriales bacterium |
| Bacteria | Actinobacteria | Actinomycetia | Micrococcales | Micrococcaceae | Arthrobacter | Arthrobacter agilis |
| Bacteria | Actinobacteria | Actinomycetia | Streptomycetales | Streptomycetaceae | Streptomyces | Streptomyces sp. |
| Bacteria | Firmicutes | Bacilli | Bacillales | Bacillaceae | Bacillus | uncultured Bacillus sp. |
| Bacteria | Firmicutes | Bacilli | Lactobacillales | Lactobacillaceae | Apilactobacillus | Apilactobacillus timberlakei |
| Bacteria | Proteobacteria | Gammaproteobacteria | Enterobacterales | Erwiniaceae | Phaseolibacter | Phaseolibacter flectens |
| Bacteria | Proteobacteria | Gammaproteobacteria | Enterobacterales | Morganellaceae | Proteus | Proteus mirabilis |
| Bacteria | Proteobacteria | Gammaproteobacteria | Enterobacterales | Morganellaceae | Proteus | Proteus vulgaris |
| Bacteria | Actinobacteria | Actinomycetia | Geodermatophilales | Geodermatophilaceae | Blastococcus | Blastococcus sp. |
| Bacteria | Bacteroidetes | Cytophagia | Cytophagales | Hymenobacteraceae | Hymenobacter | Hymenobacter sp. |
| Bacteria | Firmicutes | Bacilli | Bacillales | | Exiguobacterium | Exiguobacterium acetylicum |
| Bacteria | Proteobacteria | Betaproteobacteria | Rhodocyclales | Zoogloeaceae | Zoogloea | uncultured proteobacterium OCS7 |
| Bacteria | Proteobacteria | Gammaproteobacteria | Enterobacterales | Yersiniaceae | Serratia | Serratia marcescens |
| Bacteria | Proteobacteria | Gammaproteobacteria | Orbales | Orbaceae | Gilliamella | Gilliamella apicola |
| Bacteria | Actinobacteria | Actinomycetia | Corynebacteriales | Nocardiaceae | Rhodococcus | Rhodococcus sp. |
| Bacteria | Actinobacteria | Actinomycetia | Micrococcales | Micrococcaceae | Arthrobacter | Arthrobacter sp. |
| Bacteria | Firmicutes | Bacilli | Bacillales | Bacillaceae | Bacillus | Bacillus velezensis |

| | | | | | | |
|---|---|---|---|---|---|---|
| Bacteria | Firmicutes | Bacilli | Lactobacillales | Lactobacillaceae | Lactobacillus | Lactobacillus sp. |
| Bacteria | Proteobacteria | Alphaproteobacteria | Sphingomonadales | Sphingomonadaceae | Sphingomonas | uncultured Sphingomonas sp. |
| Bacteria | Proteobacteria | Betaproteobacteria | Burkholderiales | Oxalobacteraceae | Massilia | Massilia sp. |
| Bacteria | Proteobacteria | Deltaproteobacteria | Myxococcales | | | uncultured Myxococcales bacterium |
| Bacteria | Proteobacteria | Gammaproteobacteria | Pseudomonadales | Pseudomonadaceae | Pseudomonas | Pseudomonas sp. |
| Bacteria | Actinobacteria | Actinomycetia | Corynebacteriales | Corynebacteriaceae | Corynebacterium | uncultured Corynebacterium sp. |
| Bacteria | Actinobacteria | Actinomycetia | Propionibacteriales | Propionibacteriaceae | Cutibacterium | Cutibacterium acnes |
| Bacteria | Firmicutes | Bacilli | Bacillales | Staphylococcaceae | Staphylococcus | Staphylococcus epidermidis |
| Bacteria | Firmicutes | Negativicutes | Veillonellales | Veillonellaceae | Veillonella | Veillonella dispar |
| Bacteria | Proteobacteria | Alphaproteobacteria | Rhodospirillales | Acetobacteraceae | Acetobacter | Acetobacter pasteurianus |
| Bacteria | Proteobacteria | Betaproteobacteria | | | | uncultured beta proteobacterium |
| Bacteria | Proteobacteria | Deltaproteobacteria | | | | uncultured delta proteobacterium |
| Bacteria | | | | | | bacterium YC-ZSS-LKJ118 |
| Bacteria | Chloroflexi | | | | | uncultured Chloroflexi bacterium |
| Bacteria | Firmicutes | Bacilli | Bacillales | Bacillaceae | Bacillus | Bacillus altitudinis |
| Bacteria | Firmicutes | Bacilli | Bacillales | Bacillaceae | Psychrobacillus | Psychrobacillus psychrodurans |
| Bacteria | Firmicutes | Bacilli | Bacillales | Planococcaceae | Sporosarcina | Sporosarcina luteola |
| Bacteria | Firmicutes | Bacilli | Bacillales | Staphylococcaceae | Staphylococcus | uncultured Staphylococcus sp. |
| Bacteria | Firmicutes | Tissierellia | Tissierellales | Peptoniphilaceae | Anaerococcus | Anaerococcus prevotii |
| Bacteria | Firmicutes | | | | | uncultured Firmicutes bacterium |
| Bacteria | Proteobacteria | Betaproteobacteria | Burkholderiales | | Aquabacterium | uncultured Aquabacterium sp. |
| Bacteria | Actinobacteria | Actinomycetia | Corynebacteriales | Corynebacteriaceae | Corynebacterium | Corynebacterium sp. |
| Bacteria | Actinobacteria | Actinomycetia | Corynebacteriales | Nocardiaceae | Rhodococcus | Rhodococcus pyridinivorans |
| Bacteria | Firmicutes | Bacilli | Bacillales | Bacillaceae | Bacillus | Bacillus safensis |
| Bacteria | Firmicutes | Bacilli | Bacillales | Paenibacillaceae | Paenibacillus | Paenibacillus xylanilyticus |
| Bacteria | Proteobacteria | Alphaproteobacteria | Hyphomicrobiales | Methylobacteriaceae | Methylobacterium | Methylobacterium sp. NI91 |
| Bacteria | Proteobacteria | Alphaproteobacteria | Sphingomonadales | Sphingomonadaceae | Sphingomonas | Sphingomonas panacisoli |
| Bacteria | Proteobacteria | Alphaproteobacteria | | | | uncultured Alphaproteobacteria bacterium |
| Bacteria | Proteobacteria | Gammaproteobacteria | Enterobacterales | Enterobacteriaceae | | Enterobacteriaceae endosymbiont of Donacia cincticornis |
| Bacteria | Proteobacteria | Gammaproteobacteria | Enterobacterales | Erwiniaceae | Buchnera | Buchnera aphidicola |
| Bacteria | Proteobacteria | Gammaproteobacteria | Enterobacterales | Morganellaceae | Arsenophonus | Arsenophonus nasoniae |
| Bacteria | Proteobacteria | Gammaproteobacteria | Pseudomonadales | Moraxellaceae | Acinetobacter | uncultured Acinetobacter sp. |
| Bacteria | | | | | | bacterium BFN5 |
| Bacteria | Actinobacteria | Actinomycetia | Corynebacteriales | Mycobacteriaceae | Mycobacterium | uncultured Mycobacterium sp. |

| | | | | | | |
|---|---|---|---|---|---|---|
| Bacteria | Actinobacteria | Actinomycetia | Micrococcales | Micrococcaceae | Micrococcus | Micrococcus yunnanensis |
| Bacteria | Actinobacteria | Thermoleophilia | Solirubrobacterales | Conexibacteraceae | Conexibacter | uncultured Conexibacter sp. |
| Bacteria | Firmicutes | Bacilli | Bacillales | Staphylococcaceae | Staphylococcus | Staphylococcus aureus |
| Bacteria | Firmicutes | Bacilli | Bacillales | | Exiguobacterium | Exiguobacterium sp. CNU020 |
| Bacteria | Firmicutes | Clostridia | Eubacteriales | Lachnospiraceae | Tyzzerella | [Clostridium] piliforme |
| Bacteria | Firmicutes | Clostridia | Eubacteriales | Lachnospiraceae | | uncultured Lachnospiraceae bacterium |
| Bacteria | Proteobacteria | Alphaproteobacteria | Hyphomicrobiales | Methylobacteriaceae | Methylobacterium | uncultured Methylobacterium sp. |
| Bacteria | Proteobacteria | Alphaproteobacteria | Hyphomicrobiales | Methylobacteriaceae | Microvirga | Microvirga sp. |
| Bacteria | Proteobacteria | Alphaproteobacteria | Hyphomicrobiales | | | uncultured Rhizobiales bacterium |
| Bacteria | Proteobacteria | Alphaproteobacteria | Rhodospirillales | Acetobacteraceae | Acetobacter | Acetobacter peroxydans |
| Bacteria | Proteobacteria | Alphaproteobacteria | Rhodospirillales | Acetobacteraceae | | uncultured Acetobacteraceae bacterium |
| Bacteria | Proteobacteria | Gammaproteobacteria | Enterobacterales | Enterobacteriaceae | Klebsiella | uncultured Klebsiella sp. |
| Bacteria | Proteobacteria | Gammaproteobacteria | Enterobacterales | Erwiniaceae | Pantoea | Pantoea agglomerans |
| Bacteria | Actinobacteria | Actinomycetia | Corynebacteriales | Mycobacteriaceae | Mycobacterium | Mycobacterium paragordonae |
| Bacteria | Actinobacteria | Actinomycetia | Corynebacteriales | Mycobacteriaceae | Mycolicibacterium | Mycolicibacterium monacense |
| Bacteria | Actinobacteria | Actinomycetia | Micrococcales | Micrococcaceae | Rothia | Rothia mucilaginosa |
| Bacteria | Actinobacteria | Actinomycetia | Nakamurellales | Nakamurellaceae | Nakamurella | Nakamurella sp. |
| Bacteria | Firmicutes | Bacilli | Bacillales | Bacillaceae | Bacillus | Bacillus amyloliquefaciens |
| Bacteria | Firmicutes | Bacilli | Bacillales | Planococcaceae | Planomicrobium | Planomicrobium glaciei |
| Bacteria | Firmicutes | Bacilli | Bacillales | Planococcaceae | Sporosarcina | uncultured Sporosarcina sp. |
| Bacteria | Firmicutes | Bacilli | Bacillales | Staphylococcaceae | Jeotgalicoccus | uncultured Jeotgalicoccus sp. |
| Bacteria | Firmicutes | Bacilli | Lactobacillales | Lactobacillaceae | Lentilactobacillus | Lentilactobacillus buchneri |
| Bacteria | Firmicutes | Bacilli | Lactobacillales | Streptococcaceae | Lactococcus | Lactococcus lactis |
| Bacteria | Firmicutes | Clostridia | Eubacteriales | Clostridiaceae | Clostridium | uncultured Clostridium sp. |
| Bacteria | Proteobacteria | Betaproteobacteria | Burkholderiales | Oxalobacteraceae | Janthinobacterium | uncultured Janthinobacterium sp. |
| Bacteria | Proteobacteria | Betaproteobacteria | Burkholderiales | Oxalobacteraceae | Massilia | Massilia aurea |
| Bacteria | Proteobacteria | Betaproteobacteria | Burkholderiales | Oxalobacteraceae | Massilia | Massilia timonae |
| Bacteria | Proteobacteria | Betaproteobacteria | Burkholderiales | | Xylophilus | Xylophilus ampelinus |
| Bacteria | Proteobacteria | Gammaproteobacteria | Enterobacterales | Erwiniaceae | Erwinia | Erwinia sp. |
| Bacteria | Proteobacteria | Gammaproteobacteria | Enterobacterales | Erwiniaceae | Pantoea | Pantoea sp. |
| Bacteria | Proteobacteria | Gammaproteobacteria | Enterobacterales | Morganellaceae | Arsenophonus | uncultured Arsenophonus sp. |
| Bacteria | Proteobacteria | Gammaproteobacteria | Pseudomonadales | Moraxellaceae | Moraxella | Moraxella osloensis |
| Bacteria | Proteobacteria | Gammaproteobacteria | Xanthomonadales | Xanthomonadaceae | Stenotrophomonas | Stenotrophomonas maltophilia |
| Bacteria | Proteobacteria | Gammaproteobacteria | Xanthomonadales | Xanthomonadaceae | Stenotrophomonas | Stenotrophomonas sp. |

| | | | | | | |
|---|---|---|---|---|---|---|
| Bacteria | Actinobacteria | Actinomycetia | Corynebacteriales | Corynebacteriaceae | Corynebacterium | Corynebacterium tuberculostearicum |
| Bacteria | Actinobacteria | Actinomycetia | Corynebacteriales | Mycobacteriaceae | Mycobacterium | Mycobacterium gordonae |
| Bacteria | Actinobacteria | Actinomycetia | Corynebacteriales | Mycobacteriaceae | Mycobacterium | Mycobacterium heidelbergense |
| Bacteria | Actinobacteria | Actinomycetia | Corynebacteriales | Mycobacteriaceae | Mycobacterium | Mycobacterium sp. |
| Bacteria | Actinobacteria | Actinomycetia | Corynebacteriales | Nocardiaceae | Rhodococcus | Rhodococcus fascians |
| Bacteria | Actinobacteria | Actinomycetia | Corynebacteriales | Nocardiaceae | Rhodococcus | Rhodococcus sp. THG-MD7 |
| Bacteria | Actinobacteria | Actinomycetia | Micrococcales | Microbacteriaceae | Agrococcus | Agrococcus sp. |
| Bacteria | Actinobacteria | Actinomycetia | Micrococcales | Microbacteriaceae | Microbacterium | Microbacterium caowuchunii |
| Bacteria | Actinobacteria | Actinomycetia | Micrococcales | Microbacteriaceae | Microbacterium | Microbacterium sp. |
| Bacteria | Actinobacteria | Actinomycetia | Micrococcales | Microbacteriaceae | Microbacterium | uncultured Microbacterium sp. |
| Bacteria | Actinobacteria | Actinomycetia | Micrococcales | Micrococcaceae | Kocuria | Kocuria sp. |
| Bacteria | Actinobacteria | Actinomycetia | Propionibacteriales | Nocardioidaceae | Marmoricola | Marmoricola sp. |
| Bacteria | Actinobacteria | Actinomycetia | Propionibacteriales | Nocardioidaceae | Nocardioides | Nocardioides anomalus |
| Bacteria | Actinobacteria | Actinomycetia | Streptomycetales | Streptomycetaceae | Streptomyces | Streptomyces olivochromogenes |
| Bacteria | Actinobacteria | Thermoleophilia | Solirubrobacterales | Patulibacteraceae | Patulibacter | uncultured Patulibacter sp. |
| Bacteria | Bacteroidetes | Flavobacteriia | Flavobacteriales | Flavobacteriaceae | Flavobacterium | Flavobacterium sp. |
| Bacteria | Bacteroidetes | Flavobacteriia | Flavobacteriales | Weeksellaceae | Chryseobacterium | Chryseobacterium sp. |
| Bacteria | Bacteroidetes | Sphingobacteriia | Sphingobacteriales | Sphingobacteriaceae | Pedobacter | Pedobacter sp. |
| Bacteria | Bacteroidetes | Sphingobacteriia | Sphingobacteriales | Sphingobacteriaceae | Sphingobacterium | uncultured Sphingobacterium sp. |
| Bacteria | Bacteroidetes | | | | | uncultured Bacteroidetes bacterium |
| Bacteria | Deinococcus-Thermus | Deinococci | Deinococcales | Deinococcaceae | Deinococcus | Deinococcus geothermalis |
| Bacteria | Firmicutes | Bacilli | Bacillales | Bacillaceae | Bacillus | Bacillus paralicheniformis |
| Bacteria | Firmicutes | Bacilli | Bacillales | Bacillaceae | Bacillus | Bacillus pumilus |
| Bacteria | Firmicutes | Bacilli | Bacillales | Bacillaceae | Bacillus | Bacillus sp. DSL-17 |
| Bacteria | Firmicutes | Bacilli | Bacillales | Bacillaceae | Niallia | Bacillus nealsonii |
| Bacteria | Firmicutes | Bacilli | Bacillales | Paenibacillaceae | Paenibacillus | Paenibacillus nuruki |
| Bacteria | Firmicutes | Bacilli | Bacillales | Paenibacillaceae | Paenibacillus | Paenibacillus sp. HW |
| Bacteria | Firmicutes | Bacilli | Bacillales | Planococcaceae | Sporosarcina | Sporosarcina sp. |
| Bacteria | Firmicutes | Bacilli | Bacillales | Staphylococcaceae | Salinicoccus | Salinicoccus sp. BAB 3246 |
| Bacteria | Firmicutes | Bacilli | Lactobacillales | Enterococcaceae | Enterococcus | Enterococcus faecalis |
| Bacteria | Firmicutes | Bacilli | Lactobacillales | Lactobacillaceae | Lactobacillus | uncultured Lactobacillus sp. |
| Bacteria | Firmicutes | Bacilli | Lactobacillales | Lactobacillaceae | Pediococcus | Pediococcus acidilactici |
| Bacteria | Firmicutes | Bacilli | Lactobacillales | Lactobacillaceae | Pediococcus | Pediococcus pentosaceus |
| Bacteria | Firmicutes | Bacilli | Lactobacillales | Streptococcaceae | Streptococcus | Streptococcus gordonii |
| Bacteria | Firmicutes | Bacilli | Lactobacillales | Streptococcaceae | Streptococcus | Streptococcus mutans |
| Bacteria | Firmicutes | Negativicutes | Veillonellales | Veillonellaceae | Veillonella | Veillonella sp. |

| | | | | | | |
|---|---|---|---|---|---|---|
| Bacteria | Proteobacteria | Alphaproteobacteria | Caulobacterales | Caulobacteraceae | Brevundimonas | Brevundimonas sp. |
| Bacteria | Proteobacteria | Alphaproteobacteria | Maricaulales | Maricaulaceae | Marinicauda | Marinicauda algicola |
| Bacteria | Proteobacteria | Alphaproteobacteria | Rhodospirillales | Acetobacteraceae | Bombella | Bombella sp. ESL0368 |
| Bacteria | Proteobacteria | Alphaproteobacteria | Rhodospirillales | Acetobacteraceae | Roseomonas | Roseomonas sp. |
| Bacteria | Proteobacteria | Alphaproteobacteria | Rhodospirillales | Rhodospirillaceae | Tistrella | uncultured Tistrella sp. |
| Bacteria | Proteobacteria | Alphaproteobacteria | Sphingomonadales | Sphingomonadaceae | Novosphingobium | Novosphingobium ginsenosidimutans |
| Bacteria | Proteobacteria | Alphaproteobacteria | Sphingomonadales | Sphingomonadaceae | Novosphingobium | Novosphingobium sp. |
| Bacteria | Proteobacteria | Gammaproteobacteria | Enterobacterales | Enterobacteriaceae | Citrobacter | Citrobacter freundii |
| Bacteria | Proteobacteria | Gammaproteobacteria | Enterobacterales | Enterobacteriaceae | Enterobacter | Enterobacter sp. |
| Bacteria | Proteobacteria | Gammaproteobacteria | Enterobacterales | Erwiniaceae | Pantoea | uncultured Pantoea sp. |
| Bacteria | Proteobacteria | Gammaproteobacteria | Enterobacterales | Morganellaceae | Providencia | Providencia sp. 1709051003 |
| Bacteria | Proteobacteria | Gammaproteobacteria | Pseudomonadales | Moraxellaceae | Acinetobacter | Acinetobacter sp. |
| Bacteria | Verrucomicrobia | | | | | uncultured Verrucomicrobia bacterium |
| Bacteria | | | | | | cotton phyllosphere bacterium K |
| Bacteria | Actinobacteria | Acidimicrobiia | Acidimicrobiales | | | uncultured Acidimicrobiales bacterium |
| Bacteria | Actinobacteria | Actinomycetia | Actinomycetales | | | Actinomycetales bacterium |
| Bacteria | Actinobacteria | Actinomycetia | Actinomycetales | | | uncultured Actinomycetales bacterium |
| Bacteria | Actinobacteria | Actinomycetia | Corynebacteriales | Corynebacteriaceae | Corynebacterium | Corynebacterium minutissimum |
| Bacteria | Actinobacteria | Actinomycetia | Corynebacteriales | Mycobacteriaceae | Mycobacterium | Mycobacterium marinum |
| Bacteria | Actinobacteria | Actinomycetia | Micrococcales | Bogoriellaceae | Georgenia | Georgenia satyanarayanai |
| Bacteria | Actinobacteria | Actinomycetia | Micrococcales | Dermabacteraceae | Brachybacterium | Brachybacterium saurashtrense |
| Bacteria | Actinobacteria | Actinomycetia | Micrococcales | Dermacoccaceae | Dermacoccus | Dermacoccus abyssi |
| Bacteria | Actinobacteria | Actinomycetia | Micrococcales | Microbacteriaceae | Frigoribacterium | Frigoribacterium sp. |
| Bacteria | Actinobacteria | Actinomycetia | Micrococcales | Microbacteriaceae | Rhodoluna | Rhodoluna limnophila |
| Bacteria | Actinobacteria | Actinomycetia | Pseudonocardiales | Pseudonocardiaceae | Actinomycetospora | Actinomycetospora sp. |
| Bacteria | Actinobacteria | Rubrobacteria | Rubrobacterales | Rubrobacteraceae | | uncultured Rubrobacteraceae bacterium |
| Bacteria | Bacteroidetes | Bacteroidia | Bacteroidales | Prevotellaceae | Prevotella | Prevotella melaninogenica |
| Bacteria | Chloroflexi | Caldilineae | Caldilineales | Caldilineaceae | | uncultured Caldilineaceae bacterium |
| Bacteria | Cyanobacteria | | Chroococcales | Geminocystaceae | Cyanobacterium | uncultured Cyanobacterium sp. |
| Bacteria | Cyanobacteria | | Oscillatoriales | Cyanothecaceae | Cyanothece | uncultured Cyanothece sp. |
| Bacteria | Cyanobacteria | | | | | uncultured Antarctic cyanobacterium |
| Bacteria | Deinococcus-Thermus | Deinococci | Deinococcales | Deinococcaceae | Deinococcus | Deinococcus sp. |

| | | | | | | |
|---|---|---|---|---|---|---|
| Bacteria | Deinococcus-Thermus | Deinococci | Deinococcales | | | uncultured Deinococcales bacterium |
| Bacteria | Firmicutes | Bacilli | Bacillales | Bacillaceae | Alkalihalobacillus | Alkalihalobacillus krulwichiae |
| Bacteria | Firmicutes | Bacilli | Bacillales | Bacillaceae | Bacillus | Bacillus tequilensis |
| Bacteria | Firmicutes | Bacilli | Bacillales | Bacillaceae | Bacillus | Bacillus thuringiensis |
| Bacteria | Firmicutes | Bacilli | Bacillales | Bacillaceae | Peribacillus | Peribacillus butanolivorans |
| Bacteria | Firmicutes | Bacilli | Bacillales | Bacillaceae | Peribacillus | Peribacillus simplex |
| Bacteria | Firmicutes | Bacilli | Bacillales | Bacillaceae | Priestia | Priestia aryabhattai |
| Bacteria | Firmicutes | Bacilli | Bacillales | Bacillaceae | Weizmannia | Weizmannia coagulans |
| Bacteria | Firmicutes | Bacilli | Bacillales | Planococcaceae | Sporosarcina | Sporosarcina globispora |
| Bacteria | Firmicutes | Bacilli | Bacillales | Staphylococcaceae | Mammaliicoccus | Mammaliicoccus fleurettii |
| Bacteria | Firmicutes | Bacilli | Bacillales | Staphylococcaceae | Staphylococcus | Staphylococcus equorum |
| Bacteria | Firmicutes | Bacilli | Bacillales | Staphylococcaceae | Staphylococcus | Staphylococcus saprophyticus |
| Bacteria | Firmicutes | Bacilli | Bacillales | Staphylococcaceae | Staphylococcus | Staphylococcus sp. |
| Bacteria | Firmicutes | Bacilli | Lactobacillales | Lactobacillaceae | Latilactobacillus | Latilactobacillus curvatus |
| Bacteria | Firmicutes | Bacilli | Lactobacillales | Lactobacillaceae | Leuconostoc | Leuconostoc mesenteroides |
| Bacteria | Firmicutes | Bacilli | Lactobacillales | Streptococcaceae | Lactococcus | Lactococcus garvieae |
| Bacteria | Firmicutes | Bacilli | Lactobacillales | Streptococcaceae | Streptococcus | Streptococcus sp. LPB0220 |
| Bacteria | Firmicutes | Bacilli | Lactobacillales | Streptococcaceae | Streptococcus | Streptococcus thermophilus |
| Bacteria | Firmicutes | Bacilli | Lactobacillales | Streptococcaceae | Streptococcus | uncultured Streptococcus sp. |
| Bacteria | Firmicutes | Clostridia | Eubacteriales | Clostridiaceae | Clostridium | Clostridium sp. MT10-4G |
| Bacteria | Firmicutes | Clostridia | Eubacteriales | Eubacteriaceae | Eubacterium | uncultured Eubacterium sp. |
| Bacteria | Firmicutes | Clostridia | Eubacteriales | Lachnospiraceae | Tyzzerella | [Clostridium] colinum |
| Bacteria | Firmicutes | Tissierellia | Tissierellales | Peptoniphilaceae | Parvimonas | Parvimonas micra |
| Bacteria | Gemmatimonadetes | | | | | uncultured Gemmatimonadetes bacterium |
| Bacteria | Proteobacteria | Alphaproteobacteria | Caulobacterales | Caulobacteraceae | Brevundimonas | uncultured Brevundimonas sp. |
| Bacteria | Proteobacteria | Alphaproteobacteria | Hyphomicrobiales | Rhizobiaceae | Rhizobium | Rhizobium sp. |
| Bacteria | Proteobacteria | Alphaproteobacteria | Rhodobacterales | Rhodobacteraceae | Paracoccus | Paracoccus sp. |
| Bacteria | Proteobacteria | Alphaproteobacteria | Rhodobacterales | Rhodobacteraceae | Paracoccus | Paracoccus xiamenensis |
| Bacteria | Proteobacteria | Alphaproteobacteria | Rhodospirillales | Acetobacteraceae | | Acetobacteraceae bacterium |
| Bacteria | Proteobacteria | Alphaproteobacteria | Sphingomonadales | Sphingomonadaceae | Sphingomonas | Sphingomonas sp. G8 |
| Bacteria | Proteobacteria | Betaproteobacteria | Burkholderiales | Burkholderiaceae | Paraburkholderia | Paraburkholderia tropica |
| Bacteria | Proteobacteria | Betaproteobacteria | Burkholderiales | Comamonadaceae | Hydrogenophaga | Hydrogenophaga sp. BPS33 |
| Bacteria | Proteobacteria | Betaproteobacteria | Burkholderiales | Oxalobacteraceae | Massilia | Massilia violaceinigra |
| Bacteria | Proteobacteria | Gammaproteobacteria | Enterobacterales | Enterobacteriaceae | Enterobacter | Enterobacter hormaechei |
| Bacteria | Proteobacteria | Gammaproteobacteria | Enterobacterales | Enterobacteriaceae | Escherichia | uncultured Escherichia sp. |
| Bacteria | Proteobacteria | Gammaproteobacteria | Enterobacterales | Enterobacteriaceae | Klebsiella | Klebsiella aerogenes |

| | | | | | | |
|---|---|---|---|---|---|---|
| Bacteria | Proteobacteria | Gammaproteobacteria | Enterobacterales | Enterobacteriaceae | Klebsiella | Klebsiella oxytoca |
| Bacteria | Proteobacteria | Gammaproteobacteria | Enterobacterales | Erwiniaceae | Pantoea | Pantoea stewartii |
| Bacteria | Proteobacteria | Gammaproteobacteria | Enterobacterales | Erwiniaceae | Tatumella | Tatumella terrea |
| Bacteria | Proteobacteria | Gammaproteobacteria | Enterobacterales | Morganellaceae | Morganella | Morganella morganii |
| Bacteria | Proteobacteria | Gammaproteobacteria | Enterobacterales | Pectobacteriaceae | Pectobacterium | Pectobacterium odoriferum |
| Bacteria | Proteobacteria | Gammaproteobacteria | Orbales | Orbaceae | Orbus | uncultured Orbus sp. |
| Bacteria | Proteobacteria | Gammaproteobacteria | Pseudomonadales | Moraxellaceae | Acinetobacter | Acinetobacter indicus |
| Bacteria | Proteobacteria | Gammaproteobacteria | Pseudomonadales | Moraxellaceae | Acinetobacter | Acinetobacter radioresistens |
| Bacteria | Proteobacteria | Gammaproteobacteria | Pseudomonadales | Moraxellaceae | Acinetobacter | Acinetobacter sp. 10FS3-1 |
| Bacteria | Proteobacteria | Gammaproteobacteria | Pseudomonadales | Pseudomonadaceae | Pseudomonas | Pseudomonas aeruginosa |
| Bacteria | Proteobacteria | Gammaproteobacteria | Pseudomonadales | Pseudomonadaceae | Pseudomonas | Pseudomonas brenneri |
| Bacteria | Proteobacteria | Gammaproteobacteria | Pseudomonadales | Pseudomonadaceae | Pseudomonas | Pseudomonas sp. BIOMIG1BAC |
| Bacteria | Proteobacteria | Gammaproteobacteria | Xanthomonadales | Rhodanobacteraceae | Rhodanobacter | uncultured Rhodanobacter sp. |
| Bacteria | Proteobacteria | Gammaproteobacteria | Xanthomonadales | Xanthomonadaceae | Luteimonas | Luteimonas sp. JM171 |
| Bacteria | Tenericutes | Mollicutes | Entomoplasmatales | Spiroplasmataceae | Spiroplasma | Spiroplasma apis |
| Bacteria | | | | | | bacterium enrichment culture |
| Bacteria | | | | | | cotton phyllosphere bacterium D |
| Bacteria | | | | | | uncultured marine bacterium |
| Bacteria | Acidobacteria | Acidobacteriia | Acidobacteriales | Acidobacteriaceae | Granulicella | Granulicella sp. WH15 |
| Bacteria | Acidobacteria | Acidobacteriia | Acidobacteriales | Acidobacteriaceae | | uncultured Acidobacteriaceae bacterium |
| Bacteria | Actinobacteria | Actinomycetia | Actinomycetales | Actinomycetaceae | Actinomyces | Actinomyces gaoshouyii |
| Bacteria | Actinobacteria | Actinomycetia | Actinomycetales | Actinomycetaceae | Actinomyces | Actinomyces sp. oral clone IO077 |
| Bacteria | Actinobacteria | Actinomycetia | Actinomycetales | Actinomycetaceae | Actinomyces | Actinomyces viscosus |
| Bacteria | Actinobacteria | Actinomycetia | Actinomycetales | Actinomycetaceae | Actinomyces | uncultured Actinomyces sp. |
| Bacteria | Actinobacteria | Actinomycetia | Actinomycetales | | | Actinomycetales bacterium SSCS15 |
| Bacteria | Actinobacteria | Actinomycetia | Actinomycetales | | | uncultured actinomycete |
| Bacteria | Actinobacteria | Actinomycetia | Corynebacteriales | Corynebacteriaceae | Corynebacterium | Corynebacterium aurimucosum |
| Bacteria | Actinobacteria | Actinomycetia | Corynebacteriales | Corynebacteriaceae | Corynebacterium | Corynebacterium glyciniphilum |
| Bacteria | Actinobacteria | Actinomycetia | Corynebacteriales | Corynebacteriaceae | Corynebacterium | Corynebacterium jeikeium |
| Bacteria | Actinobacteria | Actinomycetia | Corynebacteriales | Corynebacteriaceae | Corynebacterium | Corynebacterium kroppenstedtii |
| Bacteria | Actinobacteria | Actinomycetia | Corynebacteriales | Dietziaceae | Dietzia | Dietzia sp. |
| Bacteria | Actinobacteria | Actinomycetia | Corynebacteriales | Mycobacteriaceae | Mycobacterium | Mycobacterium helveticum |
| Bacteria | Actinobacteria | Actinomycetia | Corynebacteriales | Mycobacteriaceae | Mycolicibacterium | Mycolicibacterium hodleri |

| | | | | | | |
|---|---|---|---|---|---|---|
| Bacteria | Actinobacteria | Actinomycetia | Corynebacteriales | Mycobacteriaceae | Mycolicibacterium | Mycolicibacterium smegmatis |
| Bacteria | Actinobacteria | Actinomycetia | Corynebacteriales | Mycobacteriaceae | Mycolicibacterium | Mycolicibacterium sp. |
| Bacteria | Actinobacteria | Actinomycetia | Corynebacteriales | Nocardiaceae | Nocardia | Nocardia otitidiscaviarum |
| Bacteria | Actinobacteria | Actinomycetia | Corynebacteriales | Nocardiaceae | Rhodococcus | Rhodococcus sp. XE1 |
| Bacteria | Actinobacteria | Actinomycetia | Frankiales | Frankiaceae | Frankia | Candidatus Frankia nodulisporulans |
| Bacteria | Actinobacteria | Actinomycetia | Frankiales | Frankiaceae | Frankia | Frankia symbiont of Alnus alnobetula |
| Bacteria | Actinobacteria | Actinomycetia | Frankiales | | | uncultured Frankineae bacterium |
| Bacteria | Actinobacteria | Actinomycetia | Geodermatophilales | Geodermatophilaceae | Blastococcus | Blastococcus saxobsidens |
| Bacteria | Actinobacteria | Actinomycetia | Geodermatophilales | Geodermatophilaceae | Geodermatophilus | Geodermatophilus sp. |
| Bacteria | Actinobacteria | Actinomycetia | Geodermatophilales | Geodermatophilaceae | Modestobacter | Modestobacter sp. |
| Bacteria | Actinobacteria | Actinomycetia | Kineosporiales | Kineosporiaceae | Angustibacter | Angustibacter sp. |
| Bacteria | Actinobacteria | Actinomycetia | Kineosporiales | Kineosporiaceae | Kineococcus | Kineococcus sp. |
| Bacteria | Actinobacteria | Actinomycetia | Micrococcales | Brevibacteriaceae | Brevibacterium | Brevibacterium linens |
| Bacteria | Actinobacteria | Actinomycetia | Micrococcales | Brevibacteriaceae | Brevibacterium | Brevibacterium sp. CS2 |
| Bacteria | Actinobacteria | Actinomycetia | Micrococcales | Dermabacteraceae | Brachybacterium | Brachybacterium paraconglomeratum |
| Bacteria | Actinobacteria | Actinomycetia | Micrococcales | Dermabacteraceae | Brachybacterium | Brachybacterium sp. SGAir0954 |
| Bacteria | Actinobacteria | Actinomycetia | Micrococcales | Microbacteriaceae | Clavibacter | Clavibacter michiganensis |
| Bacteria | Actinobacteria | Actinomycetia | Micrococcales | Microbacteriaceae | Curtobacterium | Curtobacterium sp. |
| Bacteria | Actinobacteria | Actinomycetia | Micrococcales | Microbacteriaceae | Frigoribacterium | Frigoribacterium endophyticum |
| Bacteria | Actinobacteria | Actinomycetia | Micrococcales | Microbacteriaceae | Frondihabitans | Frondihabitans sp. |
| Bacteria | Actinobacteria | Actinomycetia | Micrococcales | Microbacteriaceae | Leifsonia | Leifsonia shinshuensis |
| Bacteria | Actinobacteria | Actinomycetia | Micrococcales | Microbacteriaceae | Microbacterium | Microbacterium sp. 4R-513 |
| Bacteria | Actinobacteria | Actinomycetia | Micrococcales | Microbacteriaceae | Salinibacterium | Salinibacterium sp. dk2585 |
| Bacteria | Actinobacteria | Actinomycetia | Micrococcales | Micrococcaceae | Arthrobacter | Arthrobacter pascens |
| Bacteria | Actinobacteria | Actinomycetia | Micrococcales | Micrococcaceae | Arthrobacter | Arthrobacter sp. AL16P05 |
| Bacteria | Actinobacteria | Actinomycetia | Micrococcales | Micrococcaceae | Kocuria | Kocuria rosea |
| Bacteria | Actinobacteria | Actinomycetia | Micrococcales | Micrococcaceae | Kocuria | Kocuria sp. PM0532155 |
| Bacteria | Actinobacteria | Actinomycetia | Micrococcales | Micrococcaceae | Micrococcus | Micrococcus luteus |
| Bacteria | Actinobacteria | Actinomycetia | Micrococcales | Micrococcaceae | Nesterenkonia | Nesterenkonia sp. MCCC 1A10687 |
| Bacteria | Actinobacteria | Actinomycetia | Micrococcales | Micrococcaceae | Rothia | Rothia aeria |
| Bacteria | Actinobacteria | Actinomycetia | Micrococcales | Micrococcaceae | Rothia | Rothia dentocariosa |
| Bacteria | Actinobacteria | Actinomycetia | Micrococcales | Micrococcaceae | Sinomonas | Sinomonas sp. No.22 |
| Bacteria | Actinobacteria | Actinomycetia | Micrococcales | Ornithinimicrobiaceae | Ornithinimicrobium | Ornithinimicrobium pratense |
| Bacteria | Actinobacteria | Actinomycetia | Micrococcales | Promicromonosporaceae | Cellulosimicrobium | Cellulosimicrobium funkei |

| | | | | | | |
|---|---|---|---|---|---|---|
| Bacteria | Actinobacteria | Actinomycetia | Micrococcales | Sanguibacteraceae | Sanguibacter | Sanguibacter sp. MN12-15 |
| Bacteria | Actinobacteria | Actinomycetia | Micromonosporales | Micromonosporaceae | Actinoplanes | Actinoplanes violaceus |
| Bacteria | Actinobacteria | Actinomycetia | Micromonosporales | Micromonosporaceae | Micromonospora | Micromonospora sp. |
| Bacteria | Actinobacteria | Actinomycetia | Propionibacteriales | Nocardioidaceae | Aeromicrobium | Aeromicrobium sp. |
| Bacteria | Actinobacteria | Actinomycetia | Propionibacteriales | Nocardioidaceae | Friedmanniella | Friedmanniella sp. |
| Bacteria | Actinobacteria | Actinomycetia | Propionibacteriales | Nocardioidaceae | Nocardioides | Nocardioides ganghwensis |
| Bacteria | Actinobacteria | Actinomycetia | Propionibacteriales | Nocardioidaceae | Nocardioides | Nocardioides plantarum |
| Bacteria | Actinobacteria | Actinomycetia | Propionibacteriales | Nocardioidaceae | Nocardioides | Nocardioides sp. dk884 |
| Bacteria | Actinobacteria | Actinomycetia | Propionibacteriales | Propionibacteriaceae | Cutibacterium | Cutibacterium granulosum |
| Bacteria | Actinobacteria | Actinomycetia | Propionibacteriales | Propionibacteriaceae | Propionibacterium | uncultured Propionibacterium sp. |
| Bacteria | Actinobacteria | Actinomycetia | Pseudonocardiales | Pseudonocardiaceae | Pseudonocardia | Pseudonocardia sp. |
| Bacteria | Actinobacteria | Actinomycetia | Streptomycetales | Streptomycetaceae | Streptomyces | Streptomyces tendae |
| Bacteria | Actinobacteria | Coriobacteriia | Coriobacteriales | Atopobiaceae | Atopobium | uncultured Atopobium sp. |
| Bacteria | Actinobacteria | Rubrobacteria | Rubrobacterales | | | uncultured Rubrobacterales bacterium |
| Bacteria | Actinobacteria | Thermoleophilia | Solirubrobacterales | Solirubrobacteraceae | Solirubrobacter | Solirubrobacter phytolaccae |
| Bacteria | Actinobacteria | Thermoleophilia | Solirubrobacterales | | | Solirubrobacterales bacterium |
| Bacteria | Bacteroidetes | Bacteroidia | Bacteroidales | Paludibacteraceae | Paludibacter | uncultured Paludibacter sp. |
| Bacteria | Bacteroidetes | Bacteroidia | Bacteroidales | Prevotellaceae | Prevotella | Prevotella nigrescens |
| Bacteria | Bacteroidetes | Bacteroidia | Bacteroidales | Prevotellaceae | Prevotella | uncultured Prevotella sp. |
| Bacteria | Bacteroidetes | Cytophagia | Cytophagales | Hymenobacteraceae | Hymenobacter | Hymenobacter baengnokdamensis |
| Bacteria | Bacteroidetes | Cytophagia | Cytophagales | Hymenobacteraceae | Hymenobacter | Hymenobacter busanensis |
| Bacteria | Bacteroidetes | Cytophagia | Cytophagales | Hymenobacteraceae | Hymenobacter | Hymenobacter jejuensis |
| Bacteria | Bacteroidetes | Cytophagia | Cytophagales | Hymenobacteraceae | Hymenobacter | Hymenobacter psychrotolerans |
| Bacteria | Bacteroidetes | Cytophagia | Cytophagales | Hymenobacteraceae | Siccationidurans | Siccationidurans occulans |
| Bacteria | Bacteroidetes | Sphingobacteriia | Sphingobacteriales | | | uncultured Sphingobacteriales bacterium |
| Bacteria | Bacteroidetes | | | | | Bacteroidetes bacterium |
| Bacteria | Candidatus Saccharibacteria | | | | | uncultured Candidatus Saccharibacteria bacterium |
| Bacteria | Chloroflexi | Anaerolineae | | | | uncultured Anaerolineae bacterium |
| Bacteria | Cyanobacteria | | Oscillatoriales | | | uncultured Oscillatoriales cyanobacterium |
| Bacteria | Cyanobacteria | | Spirulinales | Spirulinaceae | Halospirulina | uncultured Halospirulina sp. |
| Bacteria | Cyanobacteria | | Synechococcales | Synechococcaceae | Synechococcus | Synechococcus sp. CB0101 |
| Bacteria | Deinococcus-Thermus | Deinococci | Deinococcales | Deinococcaceae | Deinococcus | Deinococcus saudiensis |
| Bacteria | Deinococcus-Thermus | Deinococci | Deinococcales | Deinococcaceae | Deinococcus | uncultured Deinococcus sp. |

| | | | | | | |
|---|---|---|---|---|---|---|
| Bacteria | Firmicutes | Bacilli | Bacillales | Bacillaceae | Alkalihalobacillus | Alkalihalobacillus clausii |
| Bacteria | Firmicutes | Bacilli | Bacillales | Bacillaceae | Alkalihalobacillus | Alkalihalobacillus gibsonii |
| Bacteria | Firmicutes | Bacilli | Bacillales | Bacillaceae | Anoxybacillus | Anoxybacillus amylolyticus |
| Bacteria | Firmicutes | Bacilli | Bacillales | Bacillaceae | Bacillus | Bacillus cereus |
| Bacteria | Firmicutes | Bacilli | Bacillales | Bacillaceae | Bacillus | Bacillus siamensis |
| Bacteria | Firmicutes | Bacilli | Bacillales | Bacillaceae | Bacillus | Bacillus sp. N3536 |
| Bacteria | Firmicutes | Bacilli | Bacillales | Bacillaceae | Bacillus | Bacillus sp. S22720 |
| Bacteria | Firmicutes | Bacilli | Bacillales | Bacillaceae | Caldibacillus | Caldibacillus thermoamylovorans |
| Bacteria | Firmicutes | Bacilli | Bacillales | Bacillaceae | Priestia | Priestia endophytica |
| Bacteria | Firmicutes | Bacilli | Bacillales | Bacillaceae | Rossellomorea | Bacillus aquimaris |
| Bacteria | Firmicutes | Bacilli | Bacillales | Bacillaceae | Rossellomorea | Bacillus oryzaecorticis |
| Bacteria | Firmicutes | Bacilli | Bacillales | Bacillaceae | Terribacillus | Terribacillus goriensis |
| Bacteria | Firmicutes | Bacilli | Bacillales | Bacillaceae | Virgibacillus | Virgibacillus doumboii |
| Bacteria | Firmicutes | Bacilli | Bacillales | Listeriaceae | Listeria | Listeria monocytogenes |
| Bacteria | Firmicutes | Bacilli | Bacillales | Paenibacillaceae | Aneurinibacillus | Aneurinibacillus aneurinilyticus |
| Bacteria | Firmicutes | Bacilli | Bacillales | Paenibacillaceae | Paenibacillus | Paenibacillus sp. |
| Bacteria | Firmicutes | Bacilli | Bacillales | Planococcaceae | Planococcus | Planococcus massiliensis |
| Bacteria | Firmicutes | Bacilli | Bacillales | Planococcaceae | Planococcus | Planococcus sp. R-36970 |
| Bacteria | Firmicutes | Bacilli | Bacillales | Planococcaceae | Planomicrobium | Planomicrobium okeanokoites |
| Bacteria | Firmicutes | Bacilli | Bacillales | Planococcaceae | Planomicrobium | Planomicrobium sp. |
| Bacteria | Firmicutes | Bacilli | Bacillales | Planococcaceae | Planomicrobium | Planomicrobium sp. Y50 |
| Bacteria | Firmicutes | Bacilli | Bacillales | Planococcaceae | Sporosarcina | Sporosarcina aquimarina |
| Bacteria | Firmicutes | Bacilli | Bacillales | Staphylococcaceae | Jeotgalicoccus | Jeotgalicoccus psychrophilus |
| Bacteria | Firmicutes | Bacilli | Bacillales | Staphylococcaceae | Staphylococcus | Staphylococcus auricularis |
| Bacteria | Firmicutes | Bacilli | Bacillales | Staphylococcaceae | Staphylococcus | Staphylococcus caprae |
| Bacteria | Firmicutes | Bacilli | Bacillales | Staphylococcaceae | Staphylococcus | Staphylococcus haemolyticus |
| Bacteria | Firmicutes | Bacilli | Bacillales | Staphylococcaceae | Staphylococcus | Staphylococcus succinus |
| Bacteria | Firmicutes | Bacilli | Bacillales | | Exiguobacterium | Exiguobacterium undae |
| Bacteria | Firmicutes | Bacilli | Lactobacillales | Aerococcaceae | Suicoccus | Suicoccus acidiformans |
| Bacteria | Firmicutes | Bacilli | Lactobacillales | Enterococcaceae | Enterococcus | Enterococcus cecorum |
| Bacteria | Firmicutes | Bacilli | Lactobacillales | Enterococcaceae | Enterococcus | Enterococcus faecium |
| Bacteria | Firmicutes | Bacilli | Lactobacillales | Enterococcaceae | Enterococcus | Enterococcus gallinarum |
| Bacteria | Firmicutes | Bacilli | Lactobacillales | Enterococcaceae | Enterococcus | uncultured Enterococcus sp. |
| Bacteria | Firmicutes | Bacilli | Lactobacillales | Enterococcaceae | Tetragenococcus | Tetragenococcus halophilus |
| Bacteria | Firmicutes | Bacilli | Lactobacillales | Enterococcaceae | Vagococcus | Vagococcus fluvialis |
| Bacteria | Firmicutes | Bacilli | Lactobacillales | Lactobacillaceae | Fructobacillus | Fructobacillus durionis |
| Bacteria | Firmicutes | Bacilli | Lactobacillales | Lactobacillaceae | Lactobacillus | Lactobacillus helsingborgensis |
| Bacteria | Firmicutes | Bacilli | Lactobacillales | Lactobacillaceae | Latilactobacillus | Latilactobacillus sakei |

| | | | | | | |
|---|---|---|---|---|---|---|
| Bacteria | Firmicutes | Bacilli | Lactobacillales | Lactobacillaceae | Weissella | Weissella hellenica |
| Bacteria | Firmicutes | Bacilli | Lactobacillales | Streptococcaceae | Streptococcus | Streptococcus agalactiae |
| Bacteria | Firmicutes | Bacilli | Lactobacillales | Streptococcaceae | Streptococcus | Streptococcus equinus |
| Bacteria | Firmicutes | Bacilli | Lactobacillales | Streptococcaceae | Streptococcus | Streptococcus gwangjuense |
| Bacteria | Firmicutes | Bacilli | Lactobacillales | Streptococcaceae | Streptococcus | Streptococcus mitis |
| Bacteria | Firmicutes | Bacilli | Lactobacillales | Streptococcaceae | Streptococcus | Streptococcus ovis |
| Bacteria | Firmicutes | Bacilli | Lactobacillales | Streptococcaceae | Streptococcus | Streptococcus pneumoniae |
| Bacteria | Firmicutes | Bacilli | Lactobacillales | Streptococcaceae | Streptococcus | Streptococcus sanguinis |
| Bacteria | Firmicutes | Bacilli | Lactobacillales | Streptococcaceae | Streptococcus | Streptococcus sp. 6(2018) |
| Bacteria | Firmicutes | Clostridia | Eubacteriales | Clostridiaceae | Alkaliphilus | uncultured Alkaliphilus sp. |
| Bacteria | Firmicutes | Clostridia | Eubacteriales | Clostridiaceae | Clostridium | Clostridium butyricum |
| Bacteria | Firmicutes | Clostridia | Eubacteriales | Lachnospiraceae | Epulopiscium | uncultured Epulopiscium sp. |
| Bacteria | Firmicutes | Clostridia | Eubacteriales | Peptostreptococcaceae | | Peptostreptococcaceae bacterium oral taxon 383 |
| Bacteria | Firmicutes | Tissierellia | Tissierellales | Peptoniphilaceae | Anaerococcus | Anaerococcus obesiensis |
| Bacteria | Firmicutes | Tissierellia | Tissierellales | Peptoniphilaceae | Anaerococcus | Anaerococcus sp. mt242 |
| Bacteria | Firmicutes | Tissierellia | Tissierellales | Peptoniphilaceae | Finegoldia | Finegoldia magna |
| Bacteria | Fusobacteria | Fusobacteriia | Fusobacteriales | Fusobacteriaceae | Fusobacterium | Fusobacterium sp. |
| Bacteria | Planctomycetes | Planctomycetia | Gemmatales | Gemmataceae | Gemmata | Gemmata obscuriglobus |
| Bacteria | Planctomycetes | Planctomycetia | Gemmatales | Gemmataceae | Urbifossiella | Urbifossiella limnaea |
| Bacteria | Planctomycetes | Planctomycetia | Isosphaerales | Isosphaeraceae | Aquisphaera | Aquisphaera sp. JC650 |
| Bacteria | Planctomycetes | Planctomycetia | Isosphaerales | Isosphaeraceae | Aquisphaera | Aquisphaera sp. JC669 |
| Bacteria | Planctomycetes | Planctomycetia | Planctomycetales | | | uncultured planctomycete |
| Bacteria | Proteobacteria | Alphaproteobacteria | Caulobacterales | Caulobacteraceae | Caulobacter | Caulobacter mirabilis |
| Bacteria | Proteobacteria | Alphaproteobacteria | Caulobacterales | Caulobacteraceae | Caulobacter | uncultured Caulobacter sp. |
| Bacteria | Proteobacteria | Alphaproteobacteria | Caulobacterales | Caulobacteraceae | | uncultured Caulobacteraceae bacterium |
| Bacteria | Proteobacteria | Alphaproteobacteria | Hyphomicrobiales | Aurantimonadaceae | Aureimonas | Aureimonas sp. |
| Bacteria | Proteobacteria | Alphaproteobacteria | Hyphomicrobiales | Bradyrhizobiaceae | Nitrobacter | uncultured Nitrobacter sp. |
| Bacteria | Proteobacteria | Alphaproteobacteria | Hyphomicrobiales | Bradyrhizobiaceae | Tardiphaga | Tardiphaga sp. vice352 |
| Bacteria | Proteobacteria | Alphaproteobacteria | Hyphomicrobiales | Chelatococcaceae | Chelatococcus | Chelatococcus sp. |
| Bacteria | Proteobacteria | Alphaproteobacteria | Hyphomicrobiales | Devosiaceae | Devosia | Devosia sp. |
| Bacteria | Proteobacteria | Alphaproteobacteria | Hyphomicrobiales | Methylobacteriaceae | Methylobacterium | Methylobacterium sp. GOBB3-216 |
| Bacteria | Proteobacteria | Alphaproteobacteria | Hyphomicrobiales | Methylocystaceae | Methylocystis | Methylocystis heyeri |
| Bacteria | Proteobacteria | Alphaproteobacteria | Hyphomicrobiales | Rhizobiaceae | Agrobacterium | Agrobacterium larrymoorei |
| Bacteria | Proteobacteria | Alphaproteobacteria | Hyphomicrobiales | Rhizobiaceae | Neorhizobium | uncultured Neorhizobium sp. |
| Bacteria | Proteobacteria | Alphaproteobacteria | Hyphomicrobiales | Rhizobiaceae | Rhizobium | Rhizobium leguminosarum |

| | | | | | | |
|---|---|---|---|---|---|---|
| Bacteria | Proteobacteria | Alphaproteobacteria | Hyphomicrobiales | Rhizobiaceae | Rhizobium | Rhizobium rhizoryzae |
| Bacteria | Proteobacteria | Alphaproteobacteria | Hyphomicrobiales | Rhizobiaceae | Shinella | Shinella sp. |
| Bacteria | Proteobacteria | Alphaproteobacteria | Hyphomicrobiales | Xanthobacteraceae | Ancylobacter | Ancylobacter sp. |
| Bacteria | Proteobacteria | Alphaproteobacteria | Micropepsales | Micropepsaceae | Rhizomicrobium | uncultured Rhizomicrobium sp. |
| Bacteria | Proteobacteria | Alphaproteobacteria | Rhodobacterales | Rhodobacteraceae | Paracoccus | Paracoccus contaminans |
| Bacteria | Proteobacteria | Alphaproteobacteria | Rhodobacterales | Rhodobacteraceae | Paracoccus | Paracoccus lichenicola |
| Bacteria | Proteobacteria | Alphaproteobacteria | Rhodobacterales | Rhodobacteraceae | Paracoccus | Paracoccus pueri |
| Bacteria | Proteobacteria | Alphaproteobacteria | Rhodobacterales | Rhodobacteraceae | Paracoccus | uncultured Paracoccus sp. |
| Bacteria | Proteobacteria | Alphaproteobacteria | Rhodobacterales | Rhodobacteraceae | Polymorphum | Polymorphum gilvum |
| Bacteria | Proteobacteria | Alphaproteobacteria | Rhodobacterales | Rhodobacteraceae | Roseovarius | Roseovarius sp. |
| Bacteria | Proteobacteria | Alphaproteobacteria | Rhodobacterales | Rhodobacteraceae | | Rhodobacteraceae bacterium |
| Bacteria | Proteobacteria | Alphaproteobacteria | Rhodospirillales | Acetobacteraceae | Gluconacetobacter | Gluconacetobacter diazotrophicus |
| Bacteria | Proteobacteria | Alphaproteobacteria | Rhodospirillales | Acetobacteraceae | Gluconobacter | Gluconobacter sp. TMW 2.767 |
| Bacteria | Proteobacteria | Alphaproteobacteria | Rhodospirillales | Acetobacteraceae | Paracraurococcus | Paracraurococcus sp. |
| Bacteria | Proteobacteria | Alphaproteobacteria | Rhodospirillales | Azospirillaceae | Azospirillum | Azospirillum lipoferum |
| Bacteria | Proteobacteria | Alphaproteobacteria | Rhodospirillales | Rhodospirillaceae | Thalassospira | uncultured Thalassospira sp. |
| Bacteria | Proteobacteria | Alphaproteobacteria | Sphingomonadales | Erythrobacteraceae | Erythrobacter | Erythrobacter neustonensis |
| Bacteria | Proteobacteria | Alphaproteobacteria | Sphingomonadales | Sphingomonadaceae | Novosphingobium | Novosphingobium sp. Gsoil 351 |
| Bacteria | Proteobacteria | Alphaproteobacteria | Sphingomonadales | Sphingomonadaceae | Sphingomonas | Sphingomonas sp. 8AM |
| Bacteria | Proteobacteria | Alphaproteobacteria | Sphingomonadales | Sphingomonadaceae | Sphingomonas | Sphingomonas sp. RNS-1 |
| Bacteria | Proteobacteria | Alphaproteobacteria | Sphingomonadales | Sphingomonadaceae | Sphingopyxis | uncultured Sphingopyxis sp. |
| Bacteria | Proteobacteria | Alphaproteobacteria | Sphingomonadales | Sphingomonadaceae | | uncultured Sphingomonadaceae bacterium |
| Bacteria | Proteobacteria | Alphaproteobacteria | Sphingomonadales | | | uncultured Sphingomonadales bacterium |
| Bacteria | Proteobacteria | Betaproteobacteria | Burkholderiales | Burkholderiaceae | Burkholderia | Burkholderia cepacia |
| Bacteria | Proteobacteria | Betaproteobacteria | Burkholderiales | Comamonadaceae | Acidovorax | Acidovorax citrulli |
| Bacteria | Proteobacteria | Betaproteobacteria | Burkholderiales | Comamonadaceae | Delftia | Delftia sp. |
| Bacteria | Proteobacteria | Betaproteobacteria | Burkholderiales | Comamonadaceae | Delftia | Delftia tsuruhatensis |
| Bacteria | Proteobacteria | Betaproteobacteria | Burkholderiales | Comamonadaceae | Variovorax | Variovorax paradoxus |
| Bacteria | Proteobacteria | Betaproteobacteria | Burkholderiales | Oxalobacteraceae | Herbaspirillum | Herbaspirillum hiltneri |
| Bacteria | Proteobacteria | Betaproteobacteria | Burkholderiales | Oxalobacteraceae | Janthinobacterium | Janthinobacterium lividum |
| Bacteria | Proteobacteria | Betaproteobacteria | Burkholderiales | Oxalobacteraceae | Massilia | uncultured Massilia sp. |
| Bacteria | Proteobacteria | Betaproteobacteria | Burkholderiales | | Xylophilus | Xylophilus rhododendri |
| Bacteria | Proteobacteria | Betaproteobacteria | Neisseriales | Chromobacteriaceae | Gulbenkiania | Gulbenkiania mobilis |

| | | | | | | |
|---|---|---|---|---|---|---|
| Bacteria | Proteobacteria | Betaproteobacteria | Neisseriales | Chromobacteriaceae | Microvirgula | Microvirgula aerodenitrificans |
| Bacteria | Proteobacteria | Betaproteobacteria | Neisseriales | Neisseriaceae | Snodgrassella | Snodgrassella alvi |
| Bacteria | Proteobacteria | Betaproteobacteria | Nitrosomonadales | Methylophilaceae | Methylophilus | Methylophilus medardicus |
| Bacteria | Proteobacteria | Betaproteobacteria | Rhodocyclales | Azonexaceae | Dechloromonas | Dechloromonas sp. HYN0024 |
| Bacteria | Proteobacteria | Betaproteobacteria | Rhodocyclales | Rhodocyclaceae | | uncultured Rhodocyclaceae bacterium |
| Bacteria | Proteobacteria | Deltaproteobacteria | Myxococcales | Archangiaceae | Stigmatella | Stigmatella aurantiaca |
| Bacteria | Proteobacteria | Deltaproteobacteria | Myxococcales | Polyangiaceae | Sorangium | Sorangium cellulosum |
| Bacteria | Proteobacteria | Deltaproteobacteria | Myxococcales | | | Sorangiineae bacterium HB-1 |
| Bacteria | Proteobacteria | Epsilonproteobacteria | Campylobacterales | Campylobacteraceae | Sulfurospirillum | uncultured Sulfurospirillum sp. |
| Bacteria | Proteobacteria | Gammaproteobacteria | Enterobacterales | Bruguierivoracaceae | Sodalis | Sodalis praecaptivus |
| Bacteria | Proteobacteria | Gammaproteobacteria | Enterobacterales | Bruguierivoracaceae | Sodalis | uncultured Sodalis sp. |
| Bacteria | Proteobacteria | Gammaproteobacteria | Enterobacterales | Enterobacteriaceae | Atlantibacter | Atlantibacter hermannii |
| Bacteria | Proteobacteria | Gammaproteobacteria | Enterobacterales | Enterobacteriaceae | Candidatus Hamiltonella | Candidatus Hamiltonella defensa |
| Bacteria | Proteobacteria | Gammaproteobacteria | Enterobacterales | Enterobacteriaceae | Cronobacter | Cronobacter sakazakii |
| Bacteria | Proteobacteria | Gammaproteobacteria | Enterobacterales | Enterobacteriaceae | Klebsiella | Klebsiella pasteurii |
| Bacteria | Proteobacteria | Gammaproteobacteria | Enterobacterales | Enterobacteriaceae | Klebsiella | Klebsiella quasipneumoniae |
| Bacteria | Proteobacteria | Gammaproteobacteria | Enterobacterales | Enterobacteriaceae | | Enterobacteriaceae bacterium G7_4_2BCO2 |
| Bacteria | Proteobacteria | Gammaproteobacteria | Enterobacterales | Enterobacteriaceae | | uncultured Enterobacteriaceae bacterium |
| Bacteria | Proteobacteria | Gammaproteobacteria | Enterobacterales | Erwiniaceae | Erwinia | Erwinia persicina |
| Bacteria | Proteobacteria | Gammaproteobacteria | Enterobacterales | Erwiniaceae | Pantoea | Pantoea ananatis |
| Bacteria | Proteobacteria | Gammaproteobacteria | Enterobacterales | Morganellaceae | Arsenophonus | Arsenophonus endosymbiont of Aphis sp. |
| Bacteria | Proteobacteria | Gammaproteobacteria | Enterobacterales | Morganellaceae | Arsenophonus | Arsenophonus sp. Jabalpur2014 |
| Bacteria | Proteobacteria | Gammaproteobacteria | Enterobacterales | Morganellaceae | Arsenophonus | Arsenophonus sp. P-220 |
| Bacteria | Proteobacteria | Gammaproteobacteria | Enterobacterales | Morganellaceae | Providencia | Providencia vermicola |
| Bacteria | Proteobacteria | Gammaproteobacteria | Enterobacterales | Morganellaceae | Providencia | uncultured Providencia sp. |
| Bacteria | Proteobacteria | Gammaproteobacteria | Enterobacterales | Pectobacteriaceae | Dickeya | Dickeya chrysanthemi |
| Bacteria | Proteobacteria | Gammaproteobacteria | Enterobacterales | Yersiniaceae | Serratia | Serratia liquefaciens |
| Bacteria | Proteobacteria | Gammaproteobacteria | Enterobacterales | Yersiniaceae | Serratia | uncultured Serratia sp. |
| Bacteria | Proteobacteria | Gammaproteobacteria | Pasteurellales | Pasteurellaceae | Pasteurella | Pasteurella multocida |
| Bacteria | Proteobacteria | Gammaproteobacteria | Pseudomonadales | Moraxellaceae | Acinetobacter | Acinetobacter apis |
| Bacteria | Proteobacteria | Gammaproteobacteria | Pseudomonadales | Moraxellaceae | Acinetobacter | Acinetobacter lwoffii |
| Bacteria | Proteobacteria | Gammaproteobacteria | Pseudomonadales | Moraxellaceae | Acinetobacter | Acinetobacter towneri |

| | | | | | | |
|---|---|---|---|---|---|---|
| Bacteria | Proteobacteria | Gammaproteobacteria | Pseudomonadales | Pseudomonadaceae | Pseudomonas | Pseudomonas chengduensis |
| Bacteria | Proteobacteria | Gammaproteobacteria | Pseudomonadales | Pseudomonadaceae | Pseudomonas | Pseudomonas fluorescens |
| Bacteria | Proteobacteria | Gammaproteobacteria | Pseudomonadales | Pseudomonadaceae | Pseudomonas | Pseudomonas psychrophila |
| Bacteria | Proteobacteria | Gammaproteobacteria | Pseudomonadales | Pseudomonadaceae | Pseudomonas | Pseudomonas syringae |
| Bacteria | Proteobacteria | Gammaproteobacteria | Vibrionales | Vibrionaceae | Salinivibrio | Salinivibrio costicola |
| Bacteria | Proteobacteria | Gammaproteobacteria | Xanthomonadales | Rhodanobacteraceae | Rhodanobacter | Rhodanobacter glycinis |
| Bacteria | Proteobacteria | Gammaproteobacteria | Xanthomonadales | Xanthomonadaceae | Lysobacter | uncultured Lysobacter sp. |
| Bacteria | Proteobacteria | Gammaproteobacteria | Xanthomonadales | Xanthomonadaceae | Pseudoxanthomonas | uncultured Pseudoxanthomonas sp. |
| Bacteria | Proteobacteria | Gammaproteobacteria | Xanthomonadales | Xanthomonadaceae | Stenotrophomonas | uncultured Stenotrophomonas sp. |
| Bacteria | Proteobacteria | Gammaproteobacteria | Xanthomonadales | Xanthomonadaceae | Xanthomonas | Xanthomonas axonopodis |
| Bacteria | Tenericutes | Mollicutes | Mycoplasmatales | Mycoplasmataceae | Mycoplasma | Mycoplasma wenyonii |
| Bacteria | | | | | | bacterium daSW.36 |
| Bacteria | | | | | | soil bacterium b10 |
| Bacteria | Abditibacteriota | Abditibacteria | Abditibacteriales | Abitibacteriaceae | Abditibacterium | Abditibacterium utsteinense |
| Bacteria | Acidobacteria | Acidobacteriia | Acidobacteriales | Acidobacteriaceae | Edaphobacter | Edaphobacter modestus |
| Bacteria | Acidobacteria | Acidobacteriia | Acidobacteriales | Acidobacteriaceae | Edaphobacter | Edaphobacter sp. IMSNU JC2954 |
| Bacteria | Acidobacteria | Acidobacteriia | Acidobacteriales | Acidobacteriaceae | Terriglobus | Terriglobus roseus |
| Bacteria | Acidobacteria | Acidobacteriia | Acidobacteriales | Acidobacteriaceae | Terriglobus | Terriglobus sp. |
| Bacteria | Acidobacteria | Acidobacteriia | Bryobacterales | Solibacteraceae | Candidatus Solibacter | uncultured Candidatus Solibacter sp. |
| Bacteria | Acidobacteria | Blastocatellia | Blastocatellales | Blastocatellaceae | Blastocatella | uncultured Blastocatella sp. |
| Bacteria | Acidobacteria | Blastocatellia | | | Chloracidobacterium | uncultured Chloracidobacterium sp. |
| Bacteria | Acidobacteria | Holophagae | | | | uncultured Holophagae bacterium |
| Bacteria | Actinobacteria | Acidimicrobiia | Acidimicrobiales | Acidimicrobiaceae | Acidimicrobium | uncultured Acidimicrobium sp. |
| Bacteria | Actinobacteria | Acidimicrobiia | Acidimicrobiales | Acidimicrobiaceae | Aciditerrimonas | uncultured Aciditerrimonas sp. |
| Bacteria | Actinobacteria | Acidimicrobiia | Acidimicrobiales | Iamiaceae | Iamia | uncultured Iamia sp. |
| Bacteria | Actinobacteria | Acidimicrobiia | Acidimicrobiales | Ilumatobacteraceae | Ilumatobacter | uncultured Ilumatobacter sp. |
| Bacteria | Actinobacteria | Actinomycetia | Acidothermales | Acidothermaceae | | uncultured Acidothermaceae bacterium |
| Bacteria | Actinobacteria | Actinomycetia | Actinomycetales | Actinomycetaceae | Actinomyces | Actinomyces israelii |
| Bacteria | Actinobacteria | Actinomycetia | Actinomycetales | Actinomycetaceae | Actinomyces | Actinomyces naeslundii |
| Bacteria | Actinobacteria | Actinomycetia | Actinomycetales | Actinomycetaceae | Actinomyces | Actinomyces sp. 432 |
| Bacteria | Actinobacteria | Actinomycetia | Actinomycetales | Actinomycetaceae | Actinomyces | Actinomyces sp. A11 |
| Bacteria | Actinobacteria | Actinomycetia | Actinomycetales | Actinomycetaceae | Actinomyces | Actinomyces sp. oral taxon 171 |
| Bacteria | Actinobacteria | Actinomycetia | Actinomycetales | Actinomycetaceae | Actinomyces | Actinomyces sp. oral taxon 414 |
| Bacteria | Actinobacteria | Actinomycetia | Actinomycetales | Actinomycetaceae | Actinomyces | Actinomyces sp. oral taxon 848 |

| | | | | | | |
|---|---|---|---|---|---|---|
| Bacteria | Actinobacteria | Actinomycetia | Actinomycetales | Actinomycetaceae | Pauljensenia | Pauljensenia hongkongensis |
| Bacteria | Actinobacteria | Actinomycetia | Actinomycetales | Actinomycetaceae | Schaalia | Schaalia radingae |
| Bacteria | Actinobacteria | Actinomycetia | Bifidobacteriales | Bifidobacteriaceae | Bifidobacterium | Bifidobacterium dentium |
| Bacteria | Actinobacteria | Actinomycetia | Bifidobacteriales | Bifidobacteriaceae | Bifidobacterium | Bifidobacterium longum |
| Bacteria | Actinobacteria | Actinomycetia | Candidatus Nanopelagicales | Candidatus Nanopelagicaceae | Candidatus Planktophila | Candidatus Planktophila vernalis |
| Bacteria | Actinobacteria | Actinomycetia | Catenulisporales | Catenulisporaceae | Catenulispora | Catenulispora sp. |
| Bacteria | Actinobacteria | Actinomycetia | Corynebacteriales | Corynebacteriaceae | Corynebacterium | Corynebacterium accolens |
| Bacteria | Actinobacteria | Actinomycetia | Corynebacteriales | Corynebacteriaceae | Corynebacterium | Corynebacterium camporealensis |
| Bacteria | Actinobacteria | Actinomycetia | Corynebacteriales | Corynebacteriaceae | Corynebacterium | Corynebacterium endometrii |
| Bacteria | Actinobacteria | Actinomycetia | Corynebacteriales | Corynebacteriaceae | Corynebacterium | Corynebacterium glaucum |
| Bacteria | Actinobacteria | Actinomycetia | Corynebacteriales | Corynebacteriaceae | Corynebacterium | Corynebacterium glucuronolyticum |
| Bacteria | Actinobacteria | Actinomycetia | Corynebacteriales | Corynebacteriaceae | Corynebacterium | Corynebacterium imitans |
| Bacteria | Actinobacteria | Actinomycetia | Corynebacteriales | Corynebacteriaceae | Corynebacterium | Corynebacterium mucifaciens |
| Bacteria | Actinobacteria | Actinomycetia | Corynebacteriales | Corynebacteriaceae | Corynebacterium | Corynebacterium nuruki |
| Bacteria | Actinobacteria | Actinomycetia | Corynebacteriales | Corynebacteriaceae | Corynebacterium | Corynebacterium propinquum |
| Bacteria | Actinobacteria | Actinomycetia | Corynebacteriales | Corynebacteriaceae | Corynebacterium | Corynebacterium provencense |
| Bacteria | Actinobacteria | Actinomycetia | Corynebacteriales | Corynebacteriaceae | Corynebacterium | Corynebacterium sanguinis |
| Bacteria | Actinobacteria | Actinomycetia | Corynebacteriales | Corynebacteriaceae | Corynebacterium | Corynebacterium segmentosum |
| Bacteria | Actinobacteria | Actinomycetia | Corynebacteriales | Corynebacteriaceae | Corynebacterium | Corynebacterium sp. ATCC 6931 |
| Bacteria | Actinobacteria | Actinomycetia | Corynebacteriales | Corynebacteriaceae | Corynebacterium | Corynebacterium sp. LMM-1652 |
| Bacteria | Actinobacteria | Actinomycetia | Corynebacteriales | Corynebacteriaceae | Corynebacterium | Corynebacterium sp. Marseille-P3884 |
| Bacteria | Actinobacteria | Actinomycetia | Corynebacteriales | Corynebacteriaceae | Corynebacterium | Corynebacterium stationis |
| Bacteria | Actinobacteria | Actinomycetia | Corynebacteriales | Corynebacteriaceae | Corynebacterium | Corynebacterium suranareeae |
| Bacteria | Actinobacteria | Actinomycetia | Corynebacteriales | Corynebacteriaceae | Corynebacterium | Corynebacterium xerosis |
| Bacteria | Actinobacteria | Actinomycetia | Corynebacteriales | Corynebacteriaceae | | Corynebacteriaceae bacterium 'ARUP UnID 227' |
| Bacteria | Actinobacteria | Actinomycetia | Corynebacteriales | Dietziaceae | Dietzia | Dietzia cinnamea |
| Bacteria | Actinobacteria | Actinomycetia | Corynebacteriales | Dietziaceae | Dietzia | Dietzia kunjamensis |
| Bacteria | Actinobacteria | Actinomycetia | Corynebacteriales | Dietziaceae | Dietzia | Dietzia maris |
| Bacteria | Actinobacteria | Actinomycetia | Corynebacteriales | Dietziaceae | Dietzia | Dietzia sp. Marseille-Q0999 |
| Bacteria | Actinobacteria | Actinomycetia | Corynebacteriales | Gordoniaceae | Gordonia | Gordonia amarae |
| Bacteria | Actinobacteria | Actinomycetia | Corynebacteriales | Gordoniaceae | Gordonia | Gordonia sp. WQ-01 |
| Bacteria | Actinobacteria | Actinomycetia | Corynebacteriales | Mycobacteriaceae | Hoyosella | Hoyosella subflava |
| Bacteria | Actinobacteria | Actinomycetia | Corynebacteriales | Mycobacteriaceae | Mycobacterium | Mycobacterium avium |

| | | | | | | |
|---|---|---|---|---|---|---|
| Bacteria | Actinobacteria | Actinomycetia | Corynebacteriales | Mycobacteriaceae | Mycobacterium | Mycobacterium conspicuum |
| Bacteria | Actinobacteria | Actinomycetia | Corynebacteriales | Mycobacteriaceae | Mycobacterium | Mycobacterium intracellulare |
| Bacteria | Actinobacteria | Actinomycetia | Corynebacteriales | Mycobacteriaceae | Mycobacterium | Mycobacterium sp. 12/13.4 AW |
| Bacteria | Actinobacteria | Actinomycetia | Corynebacteriales | Mycobacteriaceae | Mycobacterium | Mycobacterium sp. DWMJ-1330B1 |
| Bacteria | Actinobacteria | Actinomycetia | Corynebacteriales | Mycobacteriaceae | Mycobacterium | Mycobacterium sp. FI-07148 |
| Bacteria | Actinobacteria | Actinomycetia | Corynebacteriales | Mycobacteriaceae | Mycobacterium | Mycobacterium sp. N160CC |
| Bacteria | Actinobacteria | Actinomycetia | Corynebacteriales | Mycobacteriaceae | Mycolicibacterium | Mycolicibacterium arabiense |
| Bacteria | Actinobacteria | Actinomycetia | Corynebacteriales | Mycobacteriaceae | Mycolicibacterium | Mycolicibacterium chitae |
| Bacteria | Actinobacteria | Actinomycetia | Corynebacteriales | Mycobacteriaceae | Mycolicibacterium | Mycolicibacterium confluentis |
| Bacteria | Actinobacteria | Actinomycetia | Corynebacteriales | Mycobacteriaceae | Mycolicibacterium | Mycolicibacterium pallens |
| Bacteria | Actinobacteria | Actinomycetia | Corynebacteriales | Mycobacteriaceae | Mycolicibacterium | Mycolicibacterium parafortuitum |
| Bacteria | Actinobacteria | Actinomycetia | Corynebacteriales | Mycobacteriaceae | Mycolicibacterium | Mycolicibacterium sediminis |
| Bacteria | Actinobacteria | Actinomycetia | Corynebacteriales | Mycobacteriaceae | Mycolicibacterium | Mycolicibacterium tusciae |
| Bacteria | Actinobacteria | Actinomycetia | Corynebacteriales | Nocardiaceae | Nocardia | Nocardia asteroides |
| Bacteria | Actinobacteria | Actinomycetia | Corynebacteriales | Nocardiaceae | Nocardia | Nocardia brasiliensis |
| Bacteria | Actinobacteria | Actinomycetia | Corynebacteriales | Nocardiaceae | Nocardia | Nocardia globerula |
| Bacteria | Actinobacteria | Actinomycetia | Corynebacteriales | Nocardiaceae | Nocardia | Nocardia salmonicida |
| Bacteria | Actinobacteria | Actinomycetia | Corynebacteriales | Nocardiaceae | Nocardia | Nocardia seriolae |
| Bacteria | Actinobacteria | Actinomycetia | Corynebacteriales | Nocardiaceae | Nocardia | Nocardia sp. |
| Bacteria | Actinobacteria | Actinomycetia | Corynebacteriales | Nocardiaceae | Nocardia | Nocardia sp. DSM 46070 |
| Bacteria | Actinobacteria | Actinomycetia | Corynebacteriales | Nocardiaceae | Nocardia | Nocardia stercoris |
| Bacteria | Actinobacteria | Actinomycetia | Corynebacteriales | Nocardiaceae | Nocardia | uncultured Nocardia sp. |
| Bacteria | Actinobacteria | Actinomycetia | Corynebacteriales | Nocardiaceae | Rhodococcus | Rhodococcus gannanensis |
| Bacteria | Actinobacteria | Actinomycetia | Corynebacteriales | Nocardiaceae | Rhodococcus | Rhodococcus hoagii |
| Bacteria | Actinobacteria | Actinomycetia | Corynebacteriales | Nocardiaceae | Rhodococcus | Rhodococcus sp. COL-34 |
| Bacteria | Actinobacteria | Actinomycetia | Corynebacteriales | Nocardiaceae | Rhodococcus | Rhodococcus sp. P1Y |
| Bacteria | Actinobacteria | Actinomycetia | Corynebacteriales | Nocardiaceae | Rhodococcus | Rhodococcus sp. PAMC28707 |
| Bacteria | Actinobacteria | Actinomycetia | Corynebacteriales | Nocardiaceae | Rhodococcus | uncultured Rhodococcus sp. |
| Bacteria | Actinobacteria | Actinomycetia | Corynebacteriales | Nocardiaceae | Williamsia | Williamsia sp. |
| Bacteria | Actinobacteria | Actinomycetia | Corynebacteriales | Nocardiaceae | Williamsia | Williamsia sp. PE32 |
| Bacteria | Actinobacteria | Actinomycetia | Corynebacteriales | Tsukamurellaceae | Tsukamurella | Tsukamurella strandjordii |
| Bacteria | Actinobacteria | Actinomycetia | Cryptosporangiales | Cryptosporangiaceae | Cryptosporangium | Cryptosporangium sp. |
| Bacteria | Actinobacteria | Actinomycetia | Cryptosporangiales | Cryptosporangiaceae | Cryptosporangium | Cryptosporangium sp. YIM 75710 |
| Bacteria | Actinobacteria | Actinomycetia | Geodermatophilales | Geodermatophilaceae | Blastococcus | Blastococcus sp. BMG 8361 |

| | | | | | | |
|---|---|---|---|---|---|---|
| Bacteria | Actinobacteria | Actinomycetia | Geodermatophilales | Geodermatophilaceae | Blastococcus | Blastococcus sp. I12A-02939 |
| Bacteria | Actinobacteria | Actinomycetia | Geodermatophilales | Geodermatophilaceae | Blastococcus | Blastococcus sp. Marseille-P5729 |
| Bacteria | Actinobacteria | Actinomycetia | Geodermatophilales | Geodermatophilaceae | Blastococcus | Blastococcus sp. TPS166 |
| Bacteria | Actinobacteria | Actinomycetia | Geodermatophilales | Geodermatophilaceae | Blastococcus | Candidatus Blastococcus massiliensis |
| Bacteria | Actinobacteria | Actinomycetia | Geodermatophilales | Geodermatophilaceae | Geodermatophilus | Geodermatophilus africanus |
| Bacteria | Actinobacteria | Actinomycetia | Geodermatophilales | Geodermatophilaceae | Geodermatophilus | Geodermatophilus siccatus |
| Bacteria | Actinobacteria | Actinomycetia | Geodermatophilales | Geodermatophilaceae | Klenkia | Klenkia marina |
| Bacteria | Actinobacteria | Actinomycetia | Geodermatophilales | Geodermatophilaceae | Modestobacter | Modestobacter multiseptatus |
| Bacteria | Actinobacteria | Actinomycetia | Geodermatophilales | Geodermatophilaceae | Modestobacter | Modestobacter sp. 1G52 |
| Bacteria | Actinobacteria | Actinomycetia | Geodermatophilales | Geodermatophilaceae | Modestobacter | Modestobacter sp. I12A-02575 |
| Bacteria | Actinobacteria | Actinomycetia | Geodermatophilales | Geodermatophilaceae | Modestobacter | Modestobacter versicolor |
| Bacteria | Actinobacteria | Actinomycetia | Geodermatophilales | Geodermatophilaceae | Modestobacter | uncultured Modestobacter sp. |
| Bacteria | Actinobacteria | Actinomycetia | Geodermatophilales | Geodermatophilaceae | | Geodermatophilaceae bacterium |
| Bacteria | Actinobacteria | Actinomycetia | Jatrophihabitantales | Jatrophihabitantaceae | Jatrophihabitans | Jatrophihabitans sp. |
| Bacteria | Actinobacteria | Actinomycetia | Jatrophihabitantales | Jatrophihabitantaceae | Jatrophihabitans | Jatrophihabitans sp. GAS493 |
| Bacteria | Actinobacteria | Actinomycetia | Kineosporiales | Kineosporiaceae | Angustibacter | Angustibacter sp. LNNU114164 |
| Bacteria | Actinobacteria | Actinomycetia | Kineosporiales | Kineosporiaceae | Pseudokineococcus | Pseudokineococcus lusitanus |
| Bacteria | Actinobacteria | Actinomycetia | Micrococcales | Beutenbergiaceae | Serinibacter | Serinibacter sp. |
| Bacteria | Actinobacteria | Actinomycetia | Micrococcales | Brevibacteriaceae | Brevibacterium | Brevibacterium antarcticum |
| Bacteria | Actinobacteria | Actinomycetia | Micrococcales | Brevibacteriaceae | Brevibacterium | Brevibacterium paucivorans |
| Bacteria | Actinobacteria | Actinomycetia | Micrococcales | Brevibacteriaceae | Brevibacterium | Brevibacterium siliguriense |
| Bacteria | Actinobacteria | Actinomycetia | Micrococcales | Brevibacteriaceae | Brevibacterium | Brevibacterium sp. |
| Bacteria | Actinobacteria | Actinomycetia | Micrococcales | Brevibacteriaceae | Brevibacterium | Brevibacterium sp. YZ-1 |
| Bacteria | Actinobacteria | Actinomycetia | Micrococcales | Cellulomonadaceae | Actinotalea | Actinotalea sp. |
| Bacteria | Actinobacteria | Actinomycetia | Micrococcales | Cellulomonadaceae | Cellulomonas | Cellulomonas hominis |
| Bacteria | Actinobacteria | Actinomycetia | Micrococcales | Cellulomonadaceae | Cellulomonas | Cellulomonas sp. I12A-02579 |
| Bacteria | Actinobacteria | Actinomycetia | Micrococcales | Cellulomonadaceae | Cellulomonas | Cellulomonas sp. JZ18 |
| Bacteria | Actinobacteria | Actinomycetia | Micrococcales | Cellulomonadaceae | | Cellulomonadaceae bacterium A4c |
| Bacteria | Actinobacteria | Actinomycetia | Micrococcales | Dermabacteraceae | Brachybacterium | Brachybacterium ginsengisoli |
| Bacteria | Actinobacteria | Actinomycetia | Micrococcales | Dermabacteraceae | Brachybacterium | Brachybacterium muris |
| Bacteria | Actinobacteria | Actinomycetia | Micrococcales | Dermabacteraceae | Dermabacter | Dermabacter vaginalis |
| Bacteria | Actinobacteria | Actinomycetia | Micrococcales | Dermacoccaceae | Dermacoccus | Dermacoccus nishinomiyaensis |
| Bacteria | Actinobacteria | Actinomycetia | Micrococcales | Intrasporangiaceae | Janibacter | Janibacter limosus |
| Bacteria | Actinobacteria | Actinomycetia | Micrococcales | Intrasporangiaceae | Janibacter | Janibacter massiliensis |

| | | | | | | |
|---|---|---|---|---|---|---|
| Bacteria | Actinobacteri a | Actinomycetia | Micrococcales | Intrasporangiaceae | Ornithinicoccus | Ornithinicoccus sp. TUT1253 |
| Bacteria | Actinobacteri a | Actinomycetia | Micrococcales | Microbacteriaceae | Agrococcus | Agrococcus baldri |
| Bacteria | Actinobacteri a | Actinomycetia | Micrococcales | Microbacteriaceae | Agrococcus | Agrococcus carbonis |
| Bacteria | Actinobacteri a | Actinomycetia | Micrococcales | Microbacteriaceae | Agrococcus | Agrococcus sp. Marseille-P2731 |
| Bacteria | Actinobacteri a | Actinomycetia | Micrococcales | Microbacteriaceae | Agrococcus | Agrococcus sp. SGAir0287 |
| Bacteria | Actinobacteri a | Actinomycetia | Micrococcales | Microbacteriaceae | Agromyces | Agromyces atrinae |
| Bacteria | Actinobacteri a | Actinomycetia | Micrococcales | Microbacteriaceae | Agromyces | Agromyces laixinhei |
| Bacteria | Actinobacteri a | Actinomycetia | Micrococcales | Microbacteriaceae | Agromyces | Agromyces mangrovi |
| Bacteria | Actinobacteri a | Actinomycetia | Micrococcales | Microbacteriaceae | Amnibacterium | Amnibacterium setariae |
| Bacteria | Actinobacteri a | Actinomycetia | Micrococcales | Microbacteriaceae | Amnibacterium | Amnibacterium sp. |
| Bacteria | Actinobacteri a | Actinomycetia | Micrococcales | Microbacteriaceae | Amnibacterium | Amnibacterium sp. VN08A0400 |
| Bacteria | Actinobacteri a | Actinomycetia | Micrococcales | Microbacteriaceae | Aquiluna | Aquiluna borgnonia |
| Bacteria | Actinobacteri a | Actinomycetia | Micrococcales | Microbacteriaceae | Chryseoglobus | Chryseoglobus sp. |
| Bacteria | Actinobacteri a | Actinomycetia | Micrococcales | Microbacteriaceae | Cryobacterium | Cryobacterium arcticum |
| Bacteria | Actinobacteri a | Actinomycetia | Micrococcales | Microbacteriaceae | Cryobacterium | uncultured Cryobacterium sp. |
| Bacteria | Actinobacteri a | Actinomycetia | Micrococcales | Microbacteriaceae | Curtobacterium | Curtobacterium flaccumfaciens |
| Bacteria | Actinobacteri a | Actinomycetia | Micrococcales | Microbacteriaceae | Curtobacterium | Curtobacterium herbarum |
| Bacteria | Actinobacteri a | Actinomycetia | Micrococcales | Microbacteriaceae | Curtobacterium | Curtobacterium luteum |
| Bacteria | Actinobacteri a | Actinomycetia | Micrococcales | Microbacteriaceae | Curtobacterium | Curtobacterium oceanosedimentum |
| Bacteria | Actinobacteri a | Actinomycetia | Micrococcales | Microbacteriaceae | Curtobacterium | Curtobacterium sp. I12A-01512 |
| Bacteria | Actinobacteri a | Actinomycetia | Micrococcales | Microbacteriaceae | Curtobacterium | Curtobacterium sp. VKM Ac-2060 |
| Bacteria | Actinobacteri a | Actinomycetia | Micrococcales | Microbacteriaceae | Frigoribacterium | Frigoribacterium sp. NBH87 |
| Bacteria | Actinobacteri a | Actinomycetia | Micrococcales | Microbacteriaceae | Glaciihabitans | Glaciihabitans sp. INWT7 |
| Bacteria | Actinobacteri a | Actinomycetia | Micrococcales | Microbacteriaceae | Herbiconiux | Herbiconiux sp. |
| Bacteria | Actinobacteri a | Actinomycetia | Micrococcales | Microbacteriaceae | Humibacter | Humibacter sp. WJ7-1 |
| Bacteria | Actinobacteri a | Actinomycetia | Micrococcales | Microbacteriaceae | Leifsonia | Leifsonia sp. |
| Bacteria | Actinobacteri a | Actinomycetia | Micrococcales | Microbacteriaceae | Leifsonia | Leifsonia xyli |
| Bacteria | Actinobacteri a | Actinomycetia | Micrococcales | Microbacteriaceae | Leifsonia | uncultured Leifsonia sp. |
| Bacteria | Actinobacteri a | Actinomycetia | Micrococcales | Microbacteriaceae | Marinisubtilis | Marinisubtilis pacificus |
| Bacteria | Actinobacteri a | Actinomycetia | Micrococcales | Microbacteriaceae | Marisediminicol a | Marisediminicola antarctica |
| Bacteria | Actinobacteri a | Actinomycetia | Micrococcales | Microbacteriaceae | Microbacterium | Microbacterium amylolyticum |
| Bacteria | Actinobacteri a | Actinomycetia | Micrococcales | Microbacteriaceae | Microbacterium | Microbacterium azadirachtae |
| Bacteria | Actinobacteri a | Actinomycetia | Micrococcales | Microbacteriaceae | Microbacterium | Microbacterium kitamiense |
| Bacteria | Actinobacteri a | Actinomycetia | Micrococcales | Microbacteriaceae | Microbacterium | Microbacterium lushaniae |

| | | | | | | |
|---|---|---|---|---|---|---|
| Bacteria | Actinobacteri a | Actinomycetia | Micrococcales | Microbacteriaceae | Microbacterium | Microbacterium luticocti |
| Bacteria | Actinobacteri a | Actinomycetia | Micrococcales | Microbacteriaceae | Microbacterium | Microbacterium mangrovi |
| Bacteria | Actinobacteri a | Actinomycetia | Micrococcales | Microbacteriaceae | Microbacterium | Microbacterium maritypicum |
| Bacteria | Actinobacteri a | Actinomycetia | Micrococcales | Microbacteriaceae | Microbacterium | Microbacterium murale |
| Bacteria | Actinobacteri a | Actinomycetia | Micrococcales | Microbacteriaceae | Microbacterium | Microbacterium oryzae |
| Bacteria | Actinobacteri a | Actinomycetia | Micrococcales | Microbacteriaceae | Microbacterium | Microbacterium oxydans |
| Bacteria | Actinobacteri a | Actinomycetia | Micrococcales | Microbacteriaceae | Microbacterium | Microbacterium paraoxydans |
| Bacteria | Actinobacteri a | Actinomycetia | Micrococcales | Microbacteriaceae | Microbacterium | Microbacterium saccharophilum |
| Bacteria | Actinobacteri a | Actinomycetia | Micrococcales | Microbacteriaceae | Microbacterium | Microbacterium sp. EK6 |
| Bacteria | Actinobacteri a | Actinomycetia | Micrococcales | Microbacteriaceae | Microbacterium | Microbacterium sp. LC549 |
| Bacteria | Actinobacteri a | Actinomycetia | Micrococcales | Microbacteriaceae | Microbacterium | Microbacterium sp. oral clone AV005b |
| Bacteria | Actinobacteri a | Actinomycetia | Micrococcales | Microbacteriaceae | Microbacterium | Microbacterium trichothecenolyticum |
| Bacteria | Actinobacteri a | Actinomycetia | Micrococcales | Microbacteriaceae | Microbacterium | Microbacterium yannicii |
| Bacteria | Actinobacteri a | Actinomycetia | Micrococcales | Microbacteriaceae | Microcella | Microcella alkaliphila |
| Bacteria | Actinobacteri a | Actinomycetia | Micrococcales | Microbacteriaceae | Microcella | Microcella putealis |
| Bacteria | Actinobacteri a | Actinomycetia | Micrococcales | Microbacteriaceae | Mycetocola | Mycetocola manganoxydans |
| Bacteria | Actinobacteri a | Actinomycetia | Micrococcales | Microbacteriaceae | Mycetocola | Mycetocola sp. JXN-3 |
| Bacteria | Actinobacteri a | Actinomycetia | Micrococcales | Microbacteriaceae | Protaetiibacter | Protaetiibacter larvae |
| Bacteria | Actinobacteri a | Actinomycetia | Micrococcales | Microbacteriaceae | Rathayibacter | Rathayibacter sp. VKM Ac-2759 |
| Bacteria | Actinobacteri a | Actinomycetia | Micrococcales | Microbacteriaceae | Rathayibacter | Rathayibacter sp. VKM Ac-2804 |
| Bacteria | Actinobacteri a | Actinomycetia | Micrococcales | Microbacteriaceae | Salinibacterium | Salinibacterium sp. C2W-9 |
| Bacteria | Actinobacteri a | Actinomycetia | Micrococcales | Microbacteriaceae | Salinibacterium | Salinibacterium sp. MH F10 |
| Bacteria | Actinobacteri a | Actinomycetia | Micrococcales | Microbacteriaceae | Subtercola | Subtercola sp. N1-13 |
| Bacteria | Actinobacteri a | Actinomycetia | Micrococcales | Microbacteriaceae | | uncultured Microbacteriaceae bacterium |
| Bacteria | Actinobacteri a | Actinomycetia | Micrococcales | Micrococcaceae | Arthrobacter | Arthrobacter citreus |
| Bacteria | Actinobacteri a | Actinomycetia | Micrococcales | Micrococcaceae | Arthrobacter | Arthrobacter cryoconiti |
| Bacteria | Actinobacteri a | Actinomycetia | Micrococcales | Micrococcaceae | Arthrobacter | Arthrobacter dokdonellae |
| Bacteria | Actinobacteri a | Actinomycetia | Micrococcales | Micrococcaceae | Arthrobacter | Arthrobacter flavus |
| Bacteria | Actinobacteri a | Actinomycetia | Micrococcales | Micrococcaceae | Arthrobacter | Arthrobacter halodurans |
| Bacteria | Actinobacteri a | Actinomycetia | Micrococcales | Micrococcaceae | Arthrobacter | Arthrobacter pityocampae |
| Bacteria | Actinobacteri a | Actinomycetia | Micrococcales | Micrococcaceae | Arthrobacter | Arthrobacter ruber |
| Bacteria | Actinobacteri a | Actinomycetia | Micrococcales | Micrococcaceae | Arthrobacter | Arthrobacter sedimenti |
| Bacteria | Actinobacteri a | Actinomycetia | Micrococcales | Micrococcaceae | Arthrobacter | Arthrobacter sp. 6C-1 |

| | | | | | | |
|---|---|---|---|---|---|---|
| Bacteria | Actinobacteria | Actinomycetia | Micrococcales | Micrococcaceae | Arthrobacter | Arthrobacter sp. 9V |
| Bacteria | Actinobacteria | Actinomycetia | Micrococcales | Micrococcaceae | Arthrobacter | Arthrobacter sp. A2-54 |
| Bacteria | Actinobacteria | Actinomycetia | Micrococcales | Micrococcaceae | Arthrobacter | Arthrobacter sp. BAR24 |
| Bacteria | Actinobacteria | Actinomycetia | Micrococcales | Micrococcaceae | Arthrobacter | Arthrobacter sp. BMG5743 |
| Bacteria | Actinobacteria | Actinomycetia | Micrococcales | Micrococcaceae | Arthrobacter | Arthrobacter sp. MN05-02 |
| Bacteria | Actinobacteria | Actinomycetia | Micrococcales | Micrococcaceae | Auritidibacter | Auritidibacter sp. NML130574 |
| Bacteria | Actinobacteria | Actinomycetia | Micrococcales | Micrococcaceae | Citricoccus | Citricoccus parietis |
| Bacteria | Actinobacteria | Actinomycetia | Micrococcales | Micrococcaceae | Citricoccus | Citrococcus sp. |
| Bacteria | Actinobacteria | Actinomycetia | Micrococcales | Micrococcaceae | Glutamicibacter | Glutamicibacter creatinolyticus |
| Bacteria | Actinobacteria | Actinomycetia | Micrococcales | Micrococcaceae | Glutamicibacter | Glutamicibacter protophormiae |
| Bacteria | Actinobacteria | Actinomycetia | Micrococcales | Micrococcaceae | Kocuria | Kocuria indica |
| Bacteria | Actinobacteria | Actinomycetia | Micrococcales | Micrococcaceae | Micrococcus | Micrococcus endophyticus |
| Bacteria | Actinobacteria | Actinomycetia | Micrococcales | Micrococcaceae | Micrococcus | Micrococcus sp. |
| Bacteria | Actinobacteria | Actinomycetia | Micrococcales | Micrococcaceae | Nesterenkonia | Nesterenkonia lutea |
| Bacteria | Actinobacteria | Actinomycetia | Micrococcales | Micrococcaceae | Nesterenkonia | Nesterenkonia sp. AC84 |
| Bacteria | Actinobacteria | Actinomycetia | Micrococcales | Micrococcaceae | Paenarthrobacter | Paenarthrobacter nicotinovorans |
| Bacteria | Actinobacteria | Actinomycetia | Micrococcales | Micrococcaceae | Paenarthrobacter | Paenarthrobacter ureafaciens |
| Bacteria | Actinobacteria | Actinomycetia | Micrococcales | Micrococcaceae | Pseudarthrobacter | Pseudarthrobacter sp. |
| Bacteria | Actinobacteria | Actinomycetia | Micrococcales | Micrococcaceae | Pseudarthrobacter | Pseudarthrobacter sp. BIM B-2242 |
| Bacteria | Actinobacteria | Actinomycetia | Micrococcales | Micrococcaceae | Pseudarthrobacter | Pseudarthrobacter sp. YJ56 |
| Bacteria | Actinobacteria | Actinomycetia | Micrococcales | Micrococcaceae | Zafaria | Zafaria cholistanensis |
| Bacteria | Actinobacteria | Actinomycetia | Micrococcales | Micrococcaceae | | Micrococcaceae bacterium |
| Bacteria | Actinobacteria | Actinomycetia | Micrococcales | Ornithinimicrobiaceae | Ornithinimicrobium | Ornithinimicrobium cerasi |
| Bacteria | Actinobacteria | Actinomycetia | Micrococcales | Ornithinimicrobiaceae | Ornithinimicrobium | Ornithinimicrobium flavum |
| Bacteria | Actinobacteria | Actinomycetia | Micrococcales | Promicromonosporaceae | Cellulosimicrobium | Cellulosimicrobium cellulans |
| Bacteria | Actinobacteria | Actinomycetia | Micrococcales | Promicromonosporaceae | Promicromonospora | Promicromonospora thailandica |
| Bacteria | Actinobacteria | Actinomycetia | Micrococcales | Sanguibacteraceae | Sanguibacter | Sanguibacter inulinus |
| Bacteria | Actinobacteria | Actinomycetia | Micrococcales | | | uncultured Micrococcineae bacterium |
| Bacteria | Actinobacteria | Actinomycetia | Micromonosporales | Micromonosporaceae | Actinoplanes | Actinoplanes couchii |
| Bacteria | Actinobacteria | Actinomycetia | Micromonosporales | Micromonosporaceae | Actinoplanes | Actinoplanes rhizophilus |
| Bacteria | Actinobacteria | Actinomycetia | Micromonosporales | Micromonosporaceae | Actinoplanes | Actinoplanes sp. |
| Bacteria | Actinobacteria | Actinomycetia | Micromonosporales | Micromonosporaceae | Actinoplanes | Actinoplanes sp. OR16 |
| Bacteria | Actinobacteria | Actinomycetia | Micromonosporales | Micromonosporaceae | Micromonospora | Micromonospora craniellae |

| | | | | | | |
|---|---|---|---|---|---|---|
| Bacteria | Actinobacteria | Actinomycetia | Micromonosporales | Micromonosporaceae | Micromonospora | Micromonospora endophytica (Xie et al. 2001) Li et al. 2019 |
| Bacteria | Actinobacteria | Actinomycetia | Micromonosporales | Micromonosporaceae | Micromonospora | Micromonospora sp. MP38-C7 |
| Bacteria | Actinobacteria | Actinomycetia | Micromonosporales | Micromonosporaceae | Micromonospora | Micromonospora sp. NEAU-zk4 |
| Bacteria | Actinobacteria | Actinomycetia | Micromonosporales | Micromonosporaceae | Micromonospora | Micromonospora terminaliae |
| Bacteria | Actinobacteria | Actinomycetia | Micromonosporales | Micromonosporaceae | Plantactinospora | Plantactinospora sp. ACT125 |
| Bacteria | Actinobacteria | Actinomycetia | Micromonosporales | Micromonosporaceae | Plantactinospora | Plantactinospora sp. BB1 |
| Bacteria | Actinobacteria | Actinomycetia | Micromonosporales | Micromonosporaceae | Salinispora | Salinispora sp. NPS-14029 |
| Bacteria | Actinobacteria | Actinomycetia | Micromonosporales | Micromonosporaceae | | uncultured Micromonosporaceae bacterium |
| Bacteria | Actinobacteria | Actinomycetia | Motilibacterales | Motilibacteraceae | Motilibacter | Motilibacter aurantiacus |
| Bacteria | Actinobacteria | Actinomycetia | Nakamurellales | Nakamurellaceae | Nakamurella | Nakamurella sp. s14-144 |
| Bacteria | Actinobacteria | Actinomycetia | Propionibacteriales | Kribbellaceae | Kribbella | Kribbella sp. |
| Bacteria | Actinobacteria | Actinomycetia | Propionibacteriales | Nocardioidaceae | Aeromicrobium | Aeromicrobium erythreum |
| Bacteria | Actinobacteria | Actinomycetia | Propionibacteriales | Nocardioidaceae | Aeromicrobium | Aeromicrobium yanjiei |
| Bacteria | Actinobacteria | Actinomycetia | Propionibacteriales | Nocardioidaceae | Aeromicrobium | uncultured Aeromicrobium sp. |
| Bacteria | Actinobacteria | Actinomycetia | Propionibacteriales | Nocardioidaceae | Friedmanniella | Friedmanniella luteola |
| Bacteria | Actinobacteria | Actinomycetia | Propionibacteriales | Nocardioidaceae | Marmoricola | Marmoricola scoriae |
| Bacteria | Actinobacteria | Actinomycetia | Propionibacteriales | Nocardioidaceae | Nocardioides | Nocardioides albus |
| Bacteria | Actinobacteria | Actinomycetia | Propionibacteriales | Nocardioidaceae | Nocardioides | Nocardioides alkalitolerans |
| Bacteria | Actinobacteria | Actinomycetia | Propionibacteriales | Nocardioidaceae | Nocardioides | Nocardioides cynanchi |
| Bacteria | Actinobacteria | Actinomycetia | Propionibacteriales | Nocardioidaceae | Nocardioides | Nocardioides euryhalodurans |
| Bacteria | Actinobacteria | Actinomycetia | Propionibacteriales | Nocardioidaceae | Nocardioides | Nocardioides hwasunensis |
| Bacteria | Actinobacteria | Actinomycetia | Propionibacteriales | Nocardioidaceae | Nocardioides | Nocardioides jensenii |
| Bacteria | Actinobacteria | Actinomycetia | Propionibacteriales | Nocardioidaceae | Nocardioides | Nocardioides kribbensis |
| Bacteria | Actinobacteria | Actinomycetia | Propionibacteriales | Nocardioidaceae | Nocardioides | Nocardioides lentus |
| Bacteria | Actinobacteria | Actinomycetia | Propionibacteriales | Nocardioidaceae | Nocardioides | Nocardioides litorisoli |
| Bacteria | Actinobacteria | Actinomycetia | Propionibacteriales | Nocardioidaceae | Nocardioides | Nocardioides pakistanensis |
| Bacteria | Actinobacteria | Actinomycetia | Propionibacteriales | Nocardioidaceae | Nocardioides | Nocardioides seonyuensis |
| Bacteria | Actinobacteria | Actinomycetia | Propionibacteriales | Nocardioidaceae | Nocardioides | Nocardioides sp. dk3136 |
| Bacteria | Actinobacteria | Actinomycetia | Propionibacteriales | Nocardioidaceae | Nocardioides | Nocardioides sp. LNNU 3343 |
| Bacteria | Actinobacteria | Actinomycetia | Propionibacteriales | Nocardioidaceae | Nocardioides | uncultured Nocardioides sp. |
| Bacteria | Actinobacteria | Actinomycetia | Propionibacteriales | Propionibacteriaceae | Aestuariimicrobium | Aestuariimicrobium sp. |
| Bacteria | Actinobacteria | Actinomycetia | Propionibacteriales | Propionibacteriaceae | Cutibacterium | Cutibacterium avidum |
| Bacteria | Actinobacteria | Actinomycetia | Propionibacteriales | Propionibacteriaceae | Microlunatus | Microlunatus aurantiacus |

| | | | | | | |
|---|---|---|---|---|---|---|
| Bacteria | Actinobacteria | Actinomycetia | Propionibacteriales | Propionibacteriaceae | Microlunatus | Microlunatus lacustris |
| Bacteria | Actinobacteria | Actinomycetia | Propionibacteriales | Propionibacteriaceae | Microlunatus | Microlunatus sagamiharensis |
| Bacteria | Actinobacteria | Actinomycetia | Propionibacteriales | Propionibacteriaceae | Microlunatus | Microlunatus speluncae |
| Bacteria | Actinobacteria | Actinomycetia | Propionibacteriales | Propionibacteriaceae | Propionibacterium | Propionibacterium sp. |
| Bacteria | Actinobacteria | Actinomycetia | Propionibacteriales | Propionibacteriaceae | Raineyella | Raineyella sp. CBA3103 |
| Bacteria | Actinobacteria | Actinomycetia | Propionibacteriales | Propionibacteriaceae | Tessaracoccus | Tessaracoccus flavus |
| Bacteria | Actinobacteria | Actinomycetia | Propionibacteriales | Propionibacteriaceae | Tessaracoccus | Tessaracoccus profundi |
| Bacteria | Actinobacteria | Actinomycetia | Propionibacteriales | Propionibacteriaceae | Tessaracoccus | Tessaracoccus rhinocerotis |
| Bacteria | Actinobacteria | Actinomycetia | Propionibacteriales | Propionibacteriaceae | Tessaracoccus | Tessaracoccus sp. |
| Bacteria | Actinobacteria | Actinomycetia | Propionibacteriales | Propionibacteriaceae | Tessaracoccus | Tessaracoccus sp. AMV11 |
| Bacteria | Actinobacteria | Actinomycetia | Pseudonocardiales | Pseudonocardiaceae | Actinomycetospora | Actinomycetospora callitridis |
| Bacteria | Actinobacteria | Actinomycetia | Pseudonocardiales | Pseudonocardiaceae | Actinomycetospora | Actinomycetospora sp. I14A-01100 |
| Bacteria | Actinobacteria | Actinomycetia | Pseudonocardiales | Pseudonocardiaceae | Actinomycetospora | Actinomycetospora sp. J12910 |
| Bacteria | Actinobacteria | Actinomycetia | Pseudonocardiales | Pseudonocardiaceae | Actinomycetospora | Actinomycetospora sp. L1879 |
| Bacteria | Actinobacteria | Actinomycetia | Pseudonocardiales | Pseudonocardiaceae | Actinomycetospora | Actinomycetospora sp. S19-30 |
| Bacteria | Actinobacteria | Actinomycetia | Pseudonocardiales | Pseudonocardiaceae | Actinophytocola | Actinophytocola sp. J33806 |
| Bacteria | Actinobacteria | Actinomycetia | Pseudonocardiales | Pseudonocardiaceae | Actinosynnema | Actinosynnema pretiosum |
| Bacteria | Actinobacteria | Actinomycetia | Pseudonocardiales | Pseudonocardiaceae | Allokutzneria | Allokutzneria sp. |
| Bacteria | Actinobacteria | Actinomycetia | Pseudonocardiales | Pseudonocardiaceae | Prauserella | Prauserella sediminis |
| Bacteria | Actinobacteria | Actinomycetia | Pseudonocardiales | Pseudonocardiaceae | Pseudonocardia | Pseudonocardia broussonetiae |
| Bacteria | Actinobacteria | Actinomycetia | Pseudonocardiales | Pseudonocardiaceae | Pseudonocardia | Pseudonocardia yuanmonensis |
| Bacteria | Actinobacteria | Actinomycetia | Pseudonocardiales | Pseudonocardiaceae | Pseudonocardia | uncultured Pseudonocardia sp. |
| Bacteria | Actinobacteria | Actinomycetia | Pseudonocardiales | Pseudonocardiaceae | Saccharopolyspora | Saccharopolyspora sp. |
| Bacteria | Actinobacteria | Actinomycetia | Pseudonocardiales | Pseudonocardiaceae | Saccharopolyspora | Saccharopolyspora taberi |
| Bacteria | Actinobacteria | Actinomycetia | Pseudonocardiales | Pseudonocardiaceae | Saccharothrix | Saccharothrix syringae |
| Bacteria | Actinobacteria | Actinomycetia | Pseudonocardiales | | | uncultured Pseudonocardineae bacterium |
| Bacteria | Actinobacteria | Actinomycetia | Streptomycetales | Streptomycetaceae | Kitasatospora | Kitasatospora sp. |
| Bacteria | Actinobacteria | Actinomycetia | Streptomycetales | Streptomycetaceae | Streptomyces | Streptomyces ambofaciens |
| Bacteria | Actinobacteria | Actinomycetia | Streptomycetales | Streptomycetaceae | Streptomyces | Streptomyces cacaoi |
| Bacteria | Actinobacteria | Actinomycetia | Streptomycetales | Streptomycetaceae | Streptomyces | Streptomyces griseocarneus |
| Bacteria | Actinobacteria | Actinomycetia | Streptomycetales | Streptomycetaceae | Streptomyces | Streptomyces griseus |
| Bacteria | Actinobacteria | Actinomycetia | Streptomycetales | Streptomycetaceae | Streptomyces | Streptomyces indiaensis |
| Bacteria | Actinobacteria | Actinomycetia | Streptomycetales | Streptomycetaceae | Streptomyces | Streptomyces sp. CMB-StM0423 |

| | | | | | | |
|---|---|---|---|---|---|---|
| Bacteria | Actinobacteria | Actinomycetia | Streptomycetales | Streptomycetaceae | Streptomyces | Streptomyces sp. CN48 |
| Bacteria | Actinobacteria | Actinomycetia | Streptomycetales | Streptomycetaceae | Streptomyces | Streptomyces sp. DR-R71 |
| Bacteria | Actinobacteria | Actinomycetia | Streptomycetales | Streptomycetaceae | Streptomyces | Streptomyces sp. JB150 |
| Bacteria | Actinobacteria | Actinomycetia | Streptomycetales | Streptomycetaceae | Streptomyces | Streptomyces spinoverrucosus |
| Bacteria | Actinobacteria | Actinomycetia | Streptomycetales | Streptomycetaceae | Streptomyces | Streptomyces thermosacchari |
| Bacteria | Actinobacteria | Actinomycetia | Streptomycetales | Streptomycetaceae | Streptomyces | Streptomyces turgidiscabies |
| Bacteria | Actinobacteria | Actinomycetia | Streptomycetales | Streptomycetaceae | Streptomyces | Streptomyces vinaceus |
| Bacteria | Actinobacteria | Actinomycetia | Streptomycetales | Streptomycetaceae | Streptomyces | Streptomyces xiangluensis |
| Bacteria | Actinobacteria | Actinomycetia | Streptomycetales | Streptomycetaceae | Streptomyces | uncultured Streptomyces sp. |
| Bacteria | Actinobacteria | Actinomycetia | Streptomycetales | Streptomycetaceae | | Streptomycetaceae bacterium |
| Bacteria | Actinobacteria | Actinomycetia | Streptosporangiales | Nocardiopsaceae | Nocardiopsis | Nocardiopsis dassonvillei |
| Bacteria | Actinobacteria | Actinomycetia | Streptosporangiales | Nocardiopsaceae | Nocardiopsis | Nocardiopsis sp. |
| Bacteria | Actinobacteria | Actinomycetia | Streptosporangiales | Streptosporangiaceae | Nonomuraea | Nonomuraea sp. 410B12 |
| Bacteria | Actinobacteria | Actinomycetia | Streptosporangiales | Streptosporangiaceae | Nonomuraea | Nonomuraea terrinata |
| Bacteria | Actinobacteria | Actinomycetia | Streptosporangiales | Thermomonosporaceae | Actinoallomurus | Actinoallomurus bryophytorum |
| Bacteria | Actinobacteria | Actinomycetia | Streptosporangiales | Thermomonosporaceae | Actinomadura | Actinomadura sp. DS-MS-114 |
| Bacteria | Actinobacteria | Actinomycetia | | | | Actinobacteridae bacterium Control.7 |
| Bacteria | Actinobacteria | Coriobacteriia | Coriobacteriales | Atopobiaceae | Lancefieldella | Lancefieldella parvula |
| Bacteria | Actinobacteria | Coriobacteriia | Coriobacteriales | Atopobiaceae | Lancefieldella | Lancefieldella rimae |
| Bacteria | Actinobacteria | Coriobacteriia | Coriobacteriales | Atopobiaceae | Olsenella | Olsenella congonensis |
| Bacteria | Actinobacteria | Coriobacteriia | Coriobacteriales | Atopobiaceae | Olsenella | Olsenella uli |
| Bacteria | Actinobacteria | Coriobacteriia | Coriobacteriales | Atopobiaceae | Olsenella | Olsenella umbonata |
| Bacteria | Actinobacteria | Coriobacteriia | Coriobacteriales | Atopobiaceae | Olsenella | uncultured Olsenella sp. |
| Bacteria | Actinobacteria | Coriobacteriia | Coriobacteriales | Coriobacteriaceae | Collinsella | Collinsella intestinalis |
| Bacteria | Actinobacteria | Coriobacteriia | Coriobacteriales | | | Coriobacteriales bacterium DNF00809 |
| Bacteria | Actinobacteria | Coriobacteriia | Eggerthellales | Eggerthellaceae | Slackia | uncultured Slackia sp. |
| Bacteria | Actinobacteria | Rubrobacteria | Gaiellales | Gaiellaceae | Gaiella | uncultured Gaiella sp. |
| Bacteria | Actinobacteria | Rubrobacteria | Rubrobacterales | Baekduiaceae | Baekduia | Baekduia soli |
| Bacteria | Actinobacteria | Rubrobacteria | | | | uncultured Rubrobacteria bacterium |
| Bacteria | Actinobacteria | Thermoleophilia | Solirubrobacterales | Conexibacteraceae | Conexibacter | Conexibacter arvalis |
| Bacteria | Actinobacteria | Thermoleophilia | Solirubrobacterales | Conexibacteraceae | Conexibacter | Conexibacter stalactiti |
| Bacteria | Actinobacteria | Thermoleophilia | Solirubrobacterales | Patulibacteraceae | Patulibacter | Patulibacter americanus |
| Bacteria | Actinobacteria | Thermoleophilia | Solirubrobacterales | Patulibacteraceae | Patulibacter | Patulibacter ginsengiterrae |
| Bacteria | Actinobacteria | Thermoleophilia | Solirubrobacterales | Patulibacteraceae | Patulibacter | Patulibacter sp. |

| | | | | | | |
|---|---|---|---|---|---|---|
| Bacteria | Actinobacteria | Thermoleophilia | Solirubrobacterales | Patulibacteraceae | Patulibacter | Patulibacter sp. M68C4Ba |
| Bacteria | Actinobacteria | Thermoleophilia | Solirubrobacterales | Patulibacteraceae | Patulibacter | Patulibacter sp. R16 |
| Bacteria | Actinobacteria | Thermoleophilia | Solirubrobacterales | Patulibacteraceae | Patulibacter | Patulibacter sp. RG55-125 |
| Bacteria | Actinobacteria | Thermoleophilia | Solirubrobacterales | Solirubrobacteraceae | Solirubrobacter | Solirubrobacter sp. |
| Bacteria | Actinobacteria | Thermoleophilia | | | | uncultured Thermoleophilia bacterium |
| Bacteria | Armatimonadetes | Chthonomonadetes | Chthonomonadales | Chthonomonadaceae | Chthonomonas | Chthonomonas calidirosea |
| Bacteria | Bacteroidetes | Bacteroidia | Bacteroidales | Muribaculaceae | Duncaniella | Duncaniella sp. B8 |
| Bacteria | Bacteroidetes | Bacteroidia | Bacteroidales | Porphyromonadaceae | | uncultured Porphyromonadaceae bacterium |
| Bacteria | Bacteroidetes | Bacteroidia | Bacteroidales | Prevotellaceae | Alloprevotella | Alloprevotella sp. feline oral taxon 114 |
| Bacteria | Bacteroidetes | Bacteroidia | Bacteroidales | Prevotellaceae | Prevotella | Prevotella copri |
| Bacteria | Bacteroidetes | Bacteroidia | Bacteroidales | Prevotellaceae | Prevotella | Prevotella multiformis |
| Bacteria | Bacteroidetes | Bacteroidia | Bacteroidales | Prevotellaceae | Prevotella | Prevotella oralis |
| Bacteria | Bacteroidetes | Bacteroidia | Bacteroidales | Prevotellaceae | Prevotella | Prevotella sp. |
| Bacteria | Bacteroidetes | Bacteroidia | Bacteroidales | Prevotellaceae | Prevotella | Prevotella sp. oral clone FU048 |
| Bacteria | Bacteroidetes | Bacteroidia | Bacteroidales | Rikenellaceae | Alistipes | Alistipes putredinis |
| Bacteria | Bacteroidetes | Bacteroidia | Bacteroidales | | Phocaeicola | Phocaeicola vulgatus |
| Bacteria | Bacteroidetes | Bacteroidia | Bacteroidales | | | Bacteroidales bacterium |
| Bacteria | Bacteroidetes | Chitinophagia | Chitinophagales | Chitinophagaceae | Chitinophaga | Chitinophaga sp. |
| Bacteria | Bacteroidetes | Chitinophagia | Chitinophagales | Chitinophagaceae | Chitinophaga | Chitinophaga terrae Kim and Jung 2007 |
| Bacteria | Bacteroidetes | Chitinophagia | Chitinophagales | Chitinophagaceae | Sediminibacterium | uncultured Sediminibacterium sp. |
| Bacteria | Bacteroidetes | Chitinophagia | Chitinophagales | Chitinophagaceae | Segetibacter | uncultured Segetibacter sp. |
| Bacteria | Bacteroidetes | Chitinophagia | Chitinophagales | Chitinophagaceae | Taibaiella | Taibaiella sp. |
| Bacteria | Bacteroidetes | Cytophagia | Cytophagales | Cytophagaceae | Cytophaga | Cytophaga hutchinsonii |
| Bacteria | Bacteroidetes | Cytophagia | Cytophagales | Cytophagaceae | Dyadobacter | Dyadobacter alkalitolerans |
| Bacteria | Bacteroidetes | Cytophagia | Cytophagales | Cytophagaceae | Dyadobacter | Dyadobacter fermentans |
| Bacteria | Bacteroidetes | Cytophagia | Cytophagales | Cytophagaceae | Spirosoma | Spirosoma agri |
| Bacteria | Bacteroidetes | Cytophagia | Cytophagales | Hymenobacteraceae | Adhaeribacter | Adhaeribacter radiodurans |
| Bacteria | Bacteroidetes | Cytophagia | Cytophagales | Hymenobacteraceae | Adhaeribacter | Adhaeribacter sp. |
| Bacteria | Bacteroidetes | Cytophagia | Cytophagales | Hymenobacteraceae | Adhaeribacter | Adhaeribacter swui |
| Bacteria | Bacteroidetes | Cytophagia | Cytophagales | Hymenobacteraceae | Hymenobacter | Hymenobacter bucti |
| Bacteria | Bacteroidetes | Cytophagia | Cytophagales | Hymenobacteraceae | Hymenobacter | Hymenobacter gelipurpurascens |
| Bacteria | Bacteroidetes | Cytophagia | Cytophagales | Hymenobacteraceae | Hymenobacter | Hymenobacter oligotrophus |
| Bacteria | Bacteroidetes | Cytophagia | Cytophagales | Hymenobacteraceae | Hymenobacter | Hymenobacter perfusus |
| Bacteria | Bacteroidetes | Cytophagia | Cytophagales | Hymenobacteraceae | Hymenobacter | Hymenobacter polaris |
| Bacteria | Bacteroidetes | Cytophagia | Cytophagales | Hymenobacteraceae | Hymenobacter | Hymenobacter rivuli |

| | | | | | | |
|---|---|---|---|---|---|---|
| Bacteria | Bacteroidetes | Cytophagia | Cytophagales | Hymenobacteraceae | Hymenobacter | Hymenobacter russus |
| Bacteria | Bacteroidetes | Cytophagia | Cytophagales | Hymenobacteraceae | Hymenobacter | Hymenobacter sp. DG01 |
| Bacteria | Bacteroidetes | Cytophagia | Cytophagales | Hymenobacteraceae | Hymenobacter | Hymenobacter sp. L2A12 |
| Bacteria | Bacteroidetes | Cytophagia | Cytophagales | Hymenobacteraceae | Hymenobacter | Hymenobacter sp. MD1 |
| Bacteria | Bacteroidetes | Cytophagia | Cytophagales | Hymenobacteraceae | Hymenobacter | Hymenobacter sp. S2-20-2 |
| Bacteria | Bacteroidetes | Cytophagia | Cytophagales | Hymenobacteraceae | Hymenobacter | Hymenobacter terrae |
| Bacteria | Bacteroidetes | Cytophagia | Cytophagales | Hymenobacteraceae | Hymenobacter | uncultured Hymenobacter sp. |
| Bacteria | Bacteroidetes | Cytophagia | Cytophagales | Hymenobacteraceae | Parahymenobacter | Parahymenobacter deserti |
| Bacteria | Bacteroidetes | Cytophagia | Cytophagales | Hymenobacteraceae | Parahymenobacter | Parahymenobacter ocellatus |
| Bacteria | Bacteroidetes | Cytophagia | Cytophagales | Hymenobacteraceae | Pontibacter | Pontibacter pudoricolor |
| Bacteria | Bacteroidetes | Cytophagia | Cytophagales | Hymenobacteraceae | Pontibacter | Pontibacter sp. |
| Bacteria | Bacteroidetes | Cytophagia | Cytophagales | Hymenobacteraceae | Rufibacter | Rufibacter roseus |
| Bacteria | Bacteroidetes | Cytophagia | Cytophagales | Hymenobacteraceae | Rufibacter | Rufibacter sp. |
| Bacteria | Bacteroidetes | Flavobacteriia | Flavobacteriales | Cryomorphaceae | | uncultured Cryomorphaceae bacterium |
| Bacteria | Bacteroidetes | Flavobacteriia | Flavobacteriales | Flavobacteriaceae | Aequorivita | Aequorivita sp. |
| Bacteria | Bacteroidetes | Flavobacteriia | Flavobacteriales | Flavobacteriaceae | Flavobacterium | Flavobacterium hauense |
| Bacteria | Bacteroidetes | Flavobacteriia | Flavobacteriales | Flavobacteriaceae | Flavobacterium | Flavobacterium resistens |
| Bacteria | Bacteroidetes | Flavobacteriia | Flavobacteriales | Flavobacteriaceae | Flavobacterium | Flavobacterium sp. AM20-97 |
| Bacteria | Bacteroidetes | Flavobacteriia | Flavobacteriales | Flavobacteriaceae | Flavobacterium | Flavobacterium sp. THG-DN6.19 |
| Bacteria | Bacteroidetes | Flavobacteriia | Flavobacteriales | Flavobacteriaceae | Flavobacterium | Flavobacterium sp. WBUAFS-WPGT3 |
| Bacteria | Bacteroidetes | Flavobacteriia | Flavobacteriales | Flavobacteriaceae | Flavobacterium | Flavobacterium succinicans |
| Bacteria | Bacteroidetes | Flavobacteriia | Flavobacteriales | Flavobacteriaceae | Flavobacterium | Flavobacterium tiangeerense |
| Bacteria | Bacteroidetes | Flavobacteriia | Flavobacteriales | Flavobacteriaceae | Salegentibacter | uncultured Salegentibacter sp. |
| Bacteria | Bacteroidetes | Flavobacteriia | Flavobacteriales | Flavobacteriaceae | Salinimicrobium | Salinimicrobium sp. |
| Bacteria | Bacteroidetes | Flavobacteriia | Flavobacteriales | Flavobacteriaceae | Xanthomarina | Xanthomarina spongicola |
| Bacteria | Bacteroidetes | Flavobacteriia | Flavobacteriales | Weeksellaceae | Chryseobacterium | Chryseobacterium aahli |
| Bacteria | Bacteroidetes | Flavobacteriia | Flavobacteriales | Weeksellaceae | Chryseobacterium | Chryseobacterium daecheongense |
| Bacteria | Bacteroidetes | Flavobacteriia | Flavobacteriales | Weeksellaceae | Chryseobacterium | Chryseobacterium indologenes |
| Bacteria | Bacteroidetes | Flavobacteriia | Flavobacteriales | Weeksellaceae | Epilithonimonas | Epilithonimonas bovis |
| Bacteria | Bacteroidetes | Sphingobacteriia | Sphingobacteriales | Sphingobacteriaceae | Mucilaginibacter | Mucilaginibacter sp. |
| Bacteria | Bacteroidetes | Sphingobacteriia | Sphingobacteriales | Sphingobacteriaceae | Mucilaginibacter | Mucilaginibacter sp. E22 |
| Bacteria | Bacteroidetes | Sphingobacteriia | Sphingobacteriales | Sphingobacteriaceae | Mucilaginibacter | Mucilaginibacter sp. S20-104 |
| Bacteria | Bacteroidetes | Sphingobacteriia | Sphingobacteriales | Sphingobacteriaceae | Mucilaginibacter | Mucilaginibacter xinganensis |
| Bacteria | Bacteroidetes | Sphingobacteriia | Sphingobacteriales | Sphingobacteriaceae | Mucilaginibacter | Mucilaginibacter yixingensis |

| | | | | | | |
|---|---|---|---|---|---|---|
| Bacteria | Bacteroidetes | Sphingobacteriia | Sphingobacteriales | Sphingobacteriaceae | Parapedobacter | uncultured Parapedobacter sp. |
| Bacteria | Bacteroidetes | Sphingobacteriia | Sphingobacteriales | Sphingobacteriaceae | Pedobacter | Pedobacter sp. S8-2 |
| Bacteria | Bacteroidetes | Sphingobacteriia | Sphingobacteriales | Sphingobacteriaceae | Sphingobacterium | Sphingobacterium faecium |
| Bacteria | Bacteroidetes | Sphingobacteriia | Sphingobacteriales | Sphingobacteriaceae | Sphingobacterium | Sphingobacterium kitahiroshimense |
| Bacteria | Bacteroidetes | Sphingobacteriia | Sphingobacteriales | Sphingobacteriaceae | Sphingobacterium | Sphingobacterium multivorum |
| Bacteria | Bacteroidetes | | | | | Bacteroidetes bacterium swS02 |
| Bacteria | Balneolaeota | Balneolia | Balneolales | Balneolaceae | Aliifodinibius | Aliifodinibius salicampi |
| Bacteria | candidate division CPR3 | | | | | candidate division CPR3 bacterium GW2011_GWF2_35_18 |
| Bacteria | Chloroflexi | Chloroflexia | Chloroflexales | Chloroflexaceae | Chloroflexus | uncultured Chloroflexus sp. |
| Bacteria | Chloroflexi | Chloroflexia | Kallotenuales | Kallotenuaceae | | Kallotenuaceae bacterium |
| Bacteria | Chloroflexi | Dehalococcoidia | Dehalococcoidales | Dehalococcoidaceae | Dehalococcoides | Dehalococcoides mccartyi |
| Bacteria | Chloroflexi | Ktedonobacteria | Ktedonobacterales | Thermosporotrichaceae | Thermosporothrix | Thermosporothrix sp. COM3 |
| Bacteria | Chloroflexi | Ktedonobacteria | Thermogemmatisporales | Thermogemmatisporaceae | Thermogemmatispora | Thermogemmatispora argillosa |
| Bacteria | Chloroflexi | Thermomicrobia | Sphaerobacterales | Sphaerobacteraceae | Sphaerobacter | uncultured Sphaerobacter sp. |
| Bacteria | Chloroflexi | Thermomicrobia | | | | uncultured Thermomicrobia bacterium |
| Bacteria | Chloroflexi | | | | | uncultured sludge bacterium S47 |
| Bacteria | Cyanobacteria | Gloeobacteria | Gloeobacterales | Gloeobacteraceae | Gloeobacter | Gloeobacter kilaueensis |
| Bacteria | Cyanobacteria | | Chroococcales | Microcystaceae | Microcystis | Microcystis flos-aquae |
| Bacteria | Cyanobacteria | | Chroococcales | Microcystaceae | Microcystis | Microcystis sp. KH11 |
| Bacteria | Cyanobacteria | | Nostocales | Aphanizomenonaceae | Dolichospermum | Dolichospermum planctonicum |
| Bacteria | Cyanobacteria | | Nostocales | Nostocaceae | Anabaena | Anabaena sp. YBS01 |
| Bacteria | Cyanobacteria | | Nostocales | Nostocaceae | Nostoc | Nostoc sp. CENA239 |
| Bacteria | Cyanobacteria | | Nostocales | Nostocaceae | Nostoc | Nostoc sp. F19 |
| Bacteria | Cyanobacteria | | Oscillatoriales | Microcoleaceae | Microcoleus | Microcoleus pseudautumnalis |
| Bacteria | Cyanobacteria | | Oscillatoriales | Microcoleaceae | Microcoleus | Microcoleus steenstrupii |
| Bacteria | Cyanobacteria | | Oscillatoriales | Microcoleaceae | Planktothricoides | uncultured Planktothricoides sp. |
| Bacteria | Cyanobacteria | | Oscillatoriales | Oscillatoriaceae | Phormidium | Phormidium sp. YACCYB597 |
| Bacteria | Cyanobacteria | | Synechococcales | Leptolyngbyaceae | Euryhalinema | Euryhalinema mangrovii |
| Bacteria | Cyanobacteria | | Synechococcales | Leptolyngbyaceae | Leptolyngbya | Leptolyngbya sp. 37.1 |
| Bacteria | Cyanobacteria | | Synechococcales | Leptolyngbyaceae | Stenomitos | Stenomitos sp. L25 |
| Bacteria | Cyanobacteria | | Synechococcales | Oculatellaceae | Oculatella | Oculatella atacamensis |
| Bacteria | Cyanobacteria | | Synechococcales | Synechococcaceae | Synechococcus | uncultured Synechococcus sp. |
| Bacteria | Cyanobacteria | | Synechococcales | Trichocoleusaceae | Trichocoleus | Trichocoleus desertorum |
| Bacteria | Cyanobacteria | | | | | uncultured marine cyanobacterium |
| Bacteria | Deinococcus-Thermus | Deinococci | Deinococcales | Deinococcaceae | Deinococcus | Deinococcus actinosclerus |

| | | | | | | |
|---|---|---|---|---|---|---|
| Bacteria | Deinococcus-Thermus | Deinococci | Deinococcales | Deinococcaceae | Deinococcus | Deinococcus grandis |
| Bacteria | Deinococcus-Thermus | Deinococci | Deinococcales | Deinococcaceae | Deinococcus | Deinococcus puniceus |
| Bacteria | Deinococcus-Thermus | Deinococci | Deinococcales | Deinococcaceae | Deinococcus | Deinococcus radiopugnans |
| Bacteria | Deinococcus-Thermus | Deinococci | Deinococcales | Deinococcaceae | Deinococcus | Deinococcus ruber |
| Bacteria | Deinococcus-Thermus | Deinococci | Deinococcales | Deinococcaceae | Deinococcus | Deinococcus soli Cha et al. 2016 |
| Bacteria | Deinococcus-Thermus | Deinococci | Deinococcales | Deinococcaceae | Deinococcus | Deinococcus sp. A2-51 |
| Bacteria | Deinococcus-Thermus | Deinococci | Deinococcales | Deinococcaceae | Deinococcus | Deinococcus sp. MN12-1 |
| Bacteria | Deinococcus-Thermus | Deinococci | Deinococcales | Deinococcaceae | Deinococcus | Deinococcus sp. NW-56 |
| Bacteria | Deinococcus-Thermus | Deinococci | Deinococcales | Deinococcaceae | Deinococcus | Deinococcus sp. R-36479 |
| Bacteria | Deinococcus-Thermus | Deinococci | Thermales | Thermaceae | Thermus | Thermus sp. |
| Bacteria | Deinococcus-Thermus | Deinococci | Thermales | Thermaceae | Thermus | Thermus thermophilus |
| Bacteria | Deinococcus-Thermus | | | | | uncultured Thermus/Deinococcus group bacterium |
| Bacteria | Firmicutes | Bacilli | Bacillales | Alicyclobacillaceae | Tumebacillus | Tumebacillus algifaecis |
| Bacteria | Firmicutes | Bacilli | Bacillales | Bacillaceae | Aeribacillus | Aeribacillus pallidus |
| Bacteria | Firmicutes | Bacilli | Bacillales | Bacillaceae | Alkalibacillus | Alkalibacillus sp. MGR8 |
| Bacteria | Firmicutes | Bacilli | Bacillales | Bacillaceae | Alkalihalobacillus | Alkalihalobacillus halodurans |
| Bacteria | Firmicutes | Bacilli | Bacillales | Bacillaceae | Alkalihalobacillus | Alkalihalobacillus miscanthi |
| Bacteria | Firmicutes | Bacilli | Bacillales | Bacillaceae | Alkalihalobacillus | Alkalihalobacillus wakoensis |
| Bacteria | Firmicutes | Bacilli | Bacillales | Bacillaceae | Anoxybacillus | Anoxybacillus flavithermus |
| Bacteria | Firmicutes | Bacilli | Bacillales | Bacillaceae | Bacillus | [Brevibacterium] frigoritolerans |
| Bacteria | Firmicutes | Bacilli | Bacillales | Bacillaceae | Bacillus | Bacillus alkalitolerans |
| Bacteria | Firmicutes | Bacilli | Bacillales | Bacillaceae | Bacillus | Bacillus mycoides |
| Bacteria | Firmicutes | Bacilli | Bacillales | Bacillaceae | Bacillus | Bacillus smithii |
| Bacteria | Firmicutes | Bacilli | Bacillales | Bacillaceae | Bacillus | Bacillus sp. 6:21 |
| Bacteria | Firmicutes | Bacilli | Bacillales | Bacillaceae | Bacillus | Bacillus sp. AN6-4 |
| Bacteria | Firmicutes | Bacilli | Bacillales | Bacillaceae | Bacillus | Bacillus sp. BD59S |
| Bacteria | Firmicutes | Bacilli | Bacillales | Bacillaceae | Bacillus | Bacillus sp. CS25 |
| Bacteria | Firmicutes | Bacilli | Bacillales | Bacillaceae | Bacillus | Bacillus sp. DM2 |
| Bacteria | Firmicutes | Bacilli | Bacillales | Bacillaceae | Bacillus | Bacillus sp. G1DM-7 |
| Bacteria | Firmicutes | Bacilli | Bacillales | Bacillaceae | Bacillus | Bacillus sp. G5-6b |
| Bacteria | Firmicutes | Bacilli | Bacillales | Bacillaceae | Bacillus | Bacillus sp. HTJR2 |
| Bacteria | Firmicutes | Bacilli | Bacillales | Bacillaceae | Bacillus | Bacillus sp. KH172YL63 |
| Bacteria | Firmicutes | Bacilli | Bacillales | Bacillaceae | Bacillus | Bacillus sp. KP12 |
| Bacteria | Firmicutes | Bacilli | Bacillales | Bacillaceae | Bacillus | Bacillus sp. Lgg5.4 |
| Bacteria | Firmicutes | Bacilli | Bacillales | Bacillaceae | Bacillus | Bacillus sp. LS56 |
| Bacteria | Firmicutes | Bacilli | Bacillales | Bacillaceae | Bacillus | Bacillus sp. Lzh-5 |
| Bacteria | Firmicutes | Bacilli | Bacillales | Bacillaceae | Bacillus | Bacillus sp. MH602 |

| | | | | | | |
|---|---|---|---|---|---|---|
| Bacteria | Firmicutes | Bacilli | Bacillales | Bacillaceae | Bacillus | Bacillus sp. S22224 |
| Bacteria | Firmicutes | Bacilli | Bacillales | Bacillaceae | Bacillus | Bacillus sp. S3 |
| Bacteria | Firmicutes | Bacilli | Bacillales | Bacillaceae | Bacillus | Bacillus sp. S3SS555 |
| Bacteria | Firmicutes | Bacilli | Bacillales | Bacillaceae | Bacillus | Bacillus sp. SB12.1 |
| Bacteria | Firmicutes | Bacilli | Bacillales | Bacillaceae | Bacillus | Bacillus sp. SB49 |
| Bacteria | Firmicutes | Bacilli | Bacillales | Bacillaceae | Bacillus | Bacillus sp. wens01 |
| Bacteria | Firmicutes | Bacilli | Bacillales | Bacillaceae | Bacillus | Bacillus toyonensis |
| Bacteria | Firmicutes | Bacilli | Bacillales | Bacillaceae | Bacillus | Bacillus tropicus |
| Bacteria | Firmicutes | Bacilli | Bacillales | Bacillaceae | Cerasibacillus | Cerasibacillus terrae |
| Bacteria | Firmicutes | Bacilli | Bacillales | Bacillaceae | Domibacillus | Domibacillus sp. |
| Bacteria | Firmicutes | Bacilli | Bacillales | Bacillaceae | Geobacillus | Geobacillus sp. E263 |
| Bacteria | Firmicutes | Bacilli | Bacillales | Bacillaceae | Geobacillus | Geobacillus stearothermophilus |
| Bacteria | Firmicutes | Bacilli | Bacillales | Bacillaceae | Geobacillus | Geobacillus thermoleovorans |
| Bacteria | Firmicutes | Bacilli | Bacillales | Bacillaceae | Gracilibacillus | Gracilibacillus sp. SCU50 |
| Bacteria | Firmicutes | Bacilli | Bacillales | Bacillaceae | Halobacillus | Halobacillus litoralis |
| Bacteria | Firmicutes | Bacilli | Bacillales | Bacillaceae | Halobacillus | Halobacillus sp. Marseille-Q1614 |
| Bacteria | Firmicutes | Bacilli | Bacillales | Bacillaceae | Halobacillus | Halobacillus sp. YIM-kkny2 |
| Bacteria | Firmicutes | Bacilli | Bacillales | Bacillaceae | Lysinibacillus | Lysinibacillus sp. |
| Bacteria | Firmicutes | Bacilli | Bacillales | Bacillaceae | Lysinibacillus | Lysinibacillus sphaericus |
| Bacteria | Firmicutes | Bacilli | Bacillales | Bacillaceae | Margalitia | Bacillus camelliae |
| Bacteria | Firmicutes | Bacilli | Bacillales | Bacillaceae | Mesobacillus | Mesobacillus subterraneus |
| Bacteria | Firmicutes | Bacilli | Bacillales | Bacillaceae | Metabacillus | Metabacillus niabensis |
| Bacteria | Firmicutes | Bacilli | Bacillales | Bacillaceae | Neobacillus | Neobacillus niacini |
| Bacteria | Firmicutes | Bacilli | Bacillales | Bacillaceae | Oceanobacillus | Oceanobacillus halophilum |
| Bacteria | Firmicutes | Bacilli | Bacillales | Bacillaceae | Oceanobacillus | Oceanobacillus sp. |
| Bacteria | Firmicutes | Bacilli | Bacillales | Bacillaceae | Oceanobacillus | Oceanobacillus sp. R-27606 |
| Bacteria | Firmicutes | Bacilli | Bacillales | Bacillaceae | Oceanobacillus | Oceanobacillus zhaokaii |
| Bacteria | Firmicutes | Bacilli | Bacillales | Bacillaceae | Ornithinibacillus | Ornithinibacillus sp. R-27028 |
| Bacteria | Firmicutes | Bacilli | Bacillales | Bacillaceae | Parageobacillus | Parageobacillus caldoxylosilyticus |
| Bacteria | Firmicutes | Bacilli | Bacillales | Bacillaceae | Peribacillus | Peribacillus psychrosaccharolyticus |
| Bacteria | Firmicutes | Bacilli | Bacillales | Bacillaceae | Pontibacillus | Pontibacillus sp. HMF3514 |
| Bacteria | Firmicutes | Bacilli | Bacillales | Bacillaceae | Priestia | Priestia flexa |
| Bacteria | Firmicutes | Bacilli | Bacillales | Bacillaceae | Psychrobacillus | Psychrobacillus soli |
| Bacteria | Firmicutes | Bacilli | Bacillales | Bacillaceae | Psychrobacillus | Psychrobacillus sp. |
| Bacteria | Firmicutes | Bacilli | Bacillales | Bacillaceae | Pueribacillus | Pueribacillus theae |
| Bacteria | Firmicutes | Bacilli | Bacillales | Bacillaceae | Rossellomorea | Bacillus marisflavi |
| Bacteria | Firmicutes | Bacilli | Bacillales | Bacillaceae | Salirhabdus | Salirhabdus sp. Marseille-P4669 |
| Bacteria | Firmicutes | Bacilli | Bacillales | Bacillaceae | Schinkia | Calidifontibacillus azotoformans |

| | | | | | | |
|---|---|---|---|---|---|---|
| Bacteria | Firmicutes | Bacilli | Bacillales | Bacillaceae | Terribacillus | Terribacillus sp. J31 |
| Bacteria | Firmicutes | Bacilli | Bacillales | Bacillaceae | Virgibacillus | Virgibacillus natechei |
| Bacteria | Firmicutes | Bacilli | Bacillales | Bacillaceae | Virgibacillus | Virgibacillus proomii |
| Bacteria | Firmicutes | Bacilli | Bacillales | Listeriaceae | Brochothrix | Brochothrix sp. |
| Bacteria | Firmicutes | Bacilli | Bacillales | Paenibacillaceae | Brevibacillus | Brevibacillus agri |
| Bacteria | Firmicutes | Bacilli | Bacillales | Paenibacillaceae | Cohnella | Cohnella candidum |
| Bacteria | Firmicutes | Bacilli | Bacillales | Paenibacillaceae | Oxalophagus | Oxalophagus oxalicus |
| Bacteria | Firmicutes | Bacilli | Bacillales | Paenibacillaceae | Paenibacillus | Paenibacillus amylolyticus |
| Bacteria | Firmicutes | Bacilli | Bacillales | Paenibacillaceae | Paenibacillus | Paenibacillus bovis |
| Bacteria | Firmicutes | Bacilli | Bacillales | Paenibacillaceae | Paenibacillus | Paenibacillus campinasensis |
| Bacteria | Firmicutes | Bacilli | Bacillales | Paenibacillaceae | Paenibacillus | Paenibacillus lutimineralis |
| Bacteria | Firmicutes | Bacilli | Bacillales | Paenibacillaceae | Paenibacillus | Paenibacillus panacisoli |
| Bacteria | Firmicutes | Bacilli | Bacillales | Paenibacillaceae | Paenibacillus | Paenibacillus psychroresistens |
| Bacteria | Firmicutes | Bacilli | Bacillales | Paenibacillaceae | Paenibacillus | Paenibacillus sp. 32O-W |
| Bacteria | Firmicutes | Bacilli | Bacillales | Paenibacillaceae | Paenibacillus | Paenibacillus sp. AS15 |
| Bacteria | Firmicutes | Bacilli | Bacillales | Paenibacillaceae | Paenibacillus | Paenibacillus sp. FZW.41 |
| Bacteria | Firmicutes | Bacilli | Bacillales | Paenibacillaceae | Paenibacillus | Paenibacillus taichungensis |
| Bacteria | Firmicutes | Bacilli | Bacillales | Paenibacillaceae | Paenibacillus | Paenibacillus tundrae |
| Bacteria | Firmicutes | Bacilli | Bacillales | Paenibacillaceae | Paenibacillus | uncultured Paenibacillus sp. |
| Bacteria | Firmicutes | Bacilli | Bacillales | Paenibacillaceae | Saccharibacillus | Saccharibacillus brassicae |
| Bacteria | Firmicutes | Bacilli | Bacillales | Paenibacillaceae | Saccharibacillus | Saccharibacillus sp. WB 17 |
| Bacteria | Firmicutes | Bacilli | Bacillales | Paenibacillaceae | Thermobacillus | Thermobacillus composti |
| Bacteria | Firmicutes | Bacilli | Bacillales | Paenibacillaceae | Thermobacillus | Thermobacillus sp. enrichment culture clone 10 |
| Bacteria | Firmicutes | Bacilli | Bacillales | Planococcaceae | Bhargavaea | Bhargavaea sp. Marseille-Q1000 |
| Bacteria | Firmicutes | Bacilli | Bacillales | Planococcaceae | Caryophanon | Caryophanon latum |
| Bacteria | Firmicutes | Bacilli | Bacillales | Planococcaceae | Caryophanon | Caryophanon tenue |
| Bacteria | Firmicutes | Bacilli | Bacillales | Planococcaceae | Kurthia | Kurthia sp. 11kri321 |
| Bacteria | Firmicutes | Bacilli | Bacillales | Planococcaceae | Metalysinibacillus | Metalysinibacillus saudimassiliensis |
| Bacteria | Firmicutes | Bacilli | Bacillales | Planococcaceae | Planococcus | Planococcus donghaensis |
| Bacteria | Firmicutes | Bacilli | Bacillales | Planococcaceae | Planococcus | Planococcus halocryophilus |
| Bacteria | Firmicutes | Bacilli | Bacillales | Planococcaceae | Planococcus | Planococcus plakortidis |
| Bacteria | Firmicutes | Bacilli | Bacillales | Planococcaceae | Planococcus | Planococcus sp. (in: Bacteria) |
| Bacteria | Firmicutes | Bacilli | Bacillales | Planococcaceae | Planococcus | Planococcus sp. M3B-2 |
| Bacteria | Firmicutes | Bacilli | Bacillales | Planococcaceae | Planomicrobium | Planomicrobium chinense |
| Bacteria | Firmicutes | Bacilli | Bacillales | Planococcaceae | Planomicrobium | Planomicrobium flavidum |
| Bacteria | Firmicutes | Bacilli | Bacillales | Planococcaceae | Planomicrobium | Planomicrobium sp. CPCC 101110 |
| Bacteria | Firmicutes | Bacilli | Bacillales | Planococcaceae | Planomicrobium | uncultured Planomicrobium sp. |
| Bacteria | Firmicutes | Bacilli | Bacillales | Planococcaceae | Solibacillus | Solibacillus silvestris |

| | | | | | | |
|---|---|---|---|---|---|---|
| Bacteria | Firmicutes | Bacilli | Bacillales | Planococcaceae | Sporosarcina | Sporosarcina saromensis |
| Bacteria | Firmicutes | Bacilli | Bacillales | Planococcaceae | Ureibacillus | uncultured Ureibacillus sp. |
| Bacteria | Firmicutes | Bacilli | Bacillales | Planococcaceae | Ureibacillus | Ureibacillus defluvii |
| Bacteria | Firmicutes | Bacilli | Bacillales | Planococcaceae | Ureibacillus | Ureibacillus thermosphaericus |
| Bacteria | Firmicutes | Bacilli | Bacillales | Sporolactobacillaceae | Tuberibacillus | Tuberibacillus calidus |
| Bacteria | Firmicutes | Bacilli | Bacillales | Staphylococcaceae | Macrococcus | Macrococcus canis |
| Bacteria | Firmicutes | Bacilli | Bacillales | Staphylococcaceae | Macrococcus | Macrococcus caseolyticus |
| Bacteria | Firmicutes | Bacilli | Bacillales | Staphylococcaceae | Mammaliicoccus | Mammaliicoccus sciuri |
| Bacteria | Firmicutes | Bacilli | Bacillales | Staphylococcaceae | Nosocomiicoccus | Nosocomiicoccus ampullae |
| Bacteria | Firmicutes | Bacilli | Bacillales | Staphylococcaceae | Salinicoccus | Salinicoccus halitifaciens |
| Bacteria | Firmicutes | Bacilli | Bacillales | Staphylococcaceae | Salinicoccus | Salinicoccus halodurans |
| Bacteria | Firmicutes | Bacilli | Bacillales | Staphylococcaceae | Salinicoccus | Salinicoccus salsiraiae |
| Bacteria | Firmicutes | Bacilli | Bacillales | Staphylococcaceae | Staphylococcus | Staphylococcus arlettae |
| Bacteria | Firmicutes | Bacilli | Bacillales | Staphylococcaceae | Staphylococcus | Staphylococcus debuckii |
| Bacteria | Firmicutes | Bacilli | Bacillales | Staphylococcaceae | Staphylococcus | Staphylococcus hominis |
| Bacteria | Firmicutes | Bacilli | Bacillales | Staphylococcaceae | Staphylococcus | Staphylococcus lugdunensis |
| Bacteria | Firmicutes | Bacilli | Bacillales | Staphylococcaceae | Staphylococcus | Staphylococcus nepalensis |
| Bacteria | Firmicutes | Bacilli | Bacillales | Staphylococcaceae | Staphylococcus | Staphylococcus pasteuri |
| Bacteria | Firmicutes | Bacilli | Bacillales | Staphylococcaceae | Staphylococcus | Staphylococcus pettenkoferi |
| Bacteria | Firmicutes | Bacilli | Bacillales | Staphylococcaceae | Staphylococcus | Staphylococcus sp. BAB-5227 |
| Bacteria | Firmicutes | Bacilli | Bacillales | Staphylococcaceae | | Staphylococcaceae bacterium K2F20 |
| Bacteria | Firmicutes | Bacilli | Bacillales | Thermoactinomycetaceae | Hazenella | Hazenella coriacea |
| Bacteria | Firmicutes | Bacilli | Bacillales | Thermoactinomycetaceae | Kroppenstedtia | uncultured Kroppenstedtia sp. |
| Bacteria | Firmicutes | Bacilli | Bacillales | Thermoactinomycetaceae | Thermoactinomyces | Thermoactinomyces sp. |
| Bacteria | Firmicutes | Bacilli | Bacillales | | Exiguobacterium | Exiguobacterium sp. A19 |
| Bacteria | Firmicutes | Bacilli | Bacillales | | Exiguobacterium | Exiguobacterium sp. U13-1 |
| Bacteria | Firmicutes | Bacilli | Bacillales | | Exiguobacterium | uncultured Exiguobacterium sp. |
| Bacteria | Firmicutes | Bacilli | Bacillales | | Gemella | Gemella haemolysans |
| Bacteria | Firmicutes | Bacilli | Bacillales | | Gemella | Gemella palaticanis |
| Bacteria | Firmicutes | Bacilli | Bacillales | | Gemella | Gemella sanguinis |
| Bacteria | Firmicutes | Bacilli | Lactobacillales | Aerococcaceae | Abiotrophia | Abiotrophia defectiva |
| Bacteria | Firmicutes | Bacilli | Lactobacillales | Aerococcaceae | Abiotrophia | uncultured Abiotrophia sp. |
| Bacteria | Firmicutes | Bacilli | Lactobacillales | Aerococcaceae | Aerococcus | Aerococcus christensenii |
| Bacteria | Firmicutes | Bacilli | Lactobacillales | Aerococcaceae | Vaginisenegalia | Vaginisenegalia massiliensis |
| Bacteria | Firmicutes | Bacilli | Lactobacillales | Carnobacteriaceae | Carnobacterium | Carnobacterium inhibens |
| Bacteria | Firmicutes | Bacilli | Lactobacillales | Carnobacteriaceae | Carnobacterium | Carnobacterium sp. |
| Bacteria | Firmicutes | Bacilli | Lactobacillales | Carnobacteriaceae | Carnobacterium | Carnobacterium sp. 7196 |

| | | | | | | |
|---|---|---|---|---|---|---|
| Bacteria | Firmicutes | Bacilli | Lactobacillales | Carnobacteriaceae | Dolosigranulum | Dolosigranulum pigrum |
| Bacteria | Firmicutes | Bacilli | Lactobacillales | Carnobacteriaceae | Granulicatella | Granulicatella adiacens |
| Bacteria | Firmicutes | Bacilli | Lactobacillales | Carnobacteriaceae | Jeotgalibaca | Jeotgalibaca arthritidis |
| Bacteria | Firmicutes | Bacilli | Lactobacillales | Carnobacteriaceae | Marinilactibacillus | Marinilactibacillus sp. E-109 |
| Bacteria | Firmicutes | Bacilli | Lactobacillales | Carnobacteriaceae | | uncultured Carnobacteriaceae bacterium |
| Bacteria | Firmicutes | Bacilli | Lactobacillales | Enterococcaceae | Enterococcus | Enterococcus hirae |
| Bacteria | Firmicutes | Bacilli | Lactobacillales | Enterococcaceae | Enterococcus | Enterococcus italicus |
| Bacteria | Firmicutes | Bacilli | Lactobacillales | Enterococcaceae | Enterococcus | Enterococcus rotai |
| Bacteria | Firmicutes | Bacilli | Lactobacillales | Enterococcaceae | Enterococcus | Enterococcus sp. |
| Bacteria | Firmicutes | Bacilli | Lactobacillales | Enterococcaceae | Melissococcus | Melissococcus plutonius |
| Bacteria | Firmicutes | Bacilli | Lactobacillales | Lactobacillaceae | Dellaglioa | Dellaglioa algida |
| Bacteria | Firmicutes | Bacilli | Lactobacillales | Lactobacillaceae | Fructobacillus | Fructobacillus tropaeoli |
| Bacteria | Firmicutes | Bacilli | Lactobacillales | Lactobacillaceae | Furfurilactobacillus | Furfurilactobacillus curtus |
| Bacteria | Firmicutes | Bacilli | Lactobacillales | Lactobacillaceae | Lacticaseibacillus | Lacticaseibacillus casei |
| Bacteria | Firmicutes | Bacilli | Lactobacillales | Lactobacillaceae | Lacticaseibacillus | Lacticaseibacillus manihotivorans |
| Bacteria | Firmicutes | Bacilli | Lactobacillales | Lactobacillaceae | Lactiplantibacillus | Lactiplantibacillus plantarum |
| Bacteria | Firmicutes | Bacilli | Lactobacillales | Lactobacillaceae | Lactobacillus | Lactobacillus apis |
| Bacteria | Firmicutes | Bacilli | Lactobacillales | Lactobacillaceae | Lactobacillus | Lactobacillus crispatus |
| Bacteria | Firmicutes | Bacilli | Lactobacillales | Lactobacillaceae | Lactobacillus | Lactobacillus delbrueckii |
| Bacteria | Firmicutes | Bacilli | Lactobacillales | Lactobacillaceae | Lactobacillus | Lactobacillus gallinarum |
| Bacteria | Firmicutes | Bacilli | Lactobacillales | Lactobacillaceae | Lactobacillus | Lactobacillus johnsonii |
| Bacteria | Firmicutes | Bacilli | Lactobacillales | Lactobacillaceae | Lactobacillus | Lactobacillus sp. 18BSM |
| Bacteria | Firmicutes | Bacilli | Lactobacillales | Lactobacillaceae | Lactobacillus | Lactobacillus sp. ljubL1r |
| Bacteria | Firmicutes | Bacilli | Lactobacillales | Lactobacillaceae | Lactobacillus | Lactobacillus sp. oral taxon 424 |
| Bacteria | Firmicutes | Bacilli | Lactobacillales | Lactobacillaceae | Lapidilactobacillus | Lapidilactobacillus dextrinicus |
| Bacteria | Firmicutes | Bacilli | Lactobacillales | Lactobacillaceae | Lentilactobacillus | Lentilactobacillus kefiri |
| Bacteria | Firmicutes | Bacilli | Lactobacillales | Lactobacillaceae | Lentilactobacillus | Lentilactobacillus sunkii |
| Bacteria | Firmicutes | Bacilli | Lactobacillales | Lactobacillaceae | Leuconostoc | Leuconostoc citreum |
| Bacteria | Firmicutes | Bacilli | Lactobacillales | Lactobacillaceae | Leuconostoc | Leuconostoc fallax |
| Bacteria | Firmicutes | Bacilli | Lactobacillales | Lactobacillaceae | Leuconostoc | Leuconostoc gelidum |
| Bacteria | Firmicutes | Bacilli | Lactobacillales | Lactobacillaceae | Leuconostoc | Leuconostoc pseudomesenteroides |
| Bacteria | Firmicutes | Bacilli | Lactobacillales | Lactobacillaceae | Ligilactobacillus | Ligilactobacillus ceti |
| Bacteria | Firmicutes | Bacilli | Lactobacillales | Lactobacillaceae | Ligilactobacillus | Ligilactobacillus murinus |
| Bacteria | Firmicutes | Bacilli | Lactobacillales | Lactobacillaceae | Limosilactobacillus | Limosilactobacillus vaginalis |
| Bacteria | Firmicutes | Bacilli | Lactobacillales | Lactobacillaceae | Liquorilactobacillus | Liquorilactobacillus mali |
| Bacteria | Firmicutes | Bacilli | Lactobacillales | Lactobacillaceae | | uncultured Lactobacillaceae bacterium |
| Bacteria | Firmicutes | Bacilli | Lactobacillales | Streptococcaceae | Lactococcus | Lactococcus petauri |

| | | | | | | |
|---|---|---|---|---|---|---|
| Bacteria | Firmicutes | Bacilli | Lactobacillales | Streptococcaceae | Lactococcus | Lactococcus raffinolactis |
| Bacteria | Firmicutes | Bacilli | Lactobacillales | Streptococcaceae | Streptococcus | Streptococcus acidominimus |
| Bacteria | Firmicutes | Bacilli | Lactobacillales | Streptococcaceae | Streptococcus | Streptococcus chosunense |
| Bacteria | Firmicutes | Bacilli | Lactobacillales | Streptococcaceae | Streptococcus | Streptococcus criceti |
| Bacteria | Firmicutes | Bacilli | Lactobacillales | Streptococcaceae | Streptococcus | Streptococcus himalayensis |
| Bacteria | Firmicutes | Bacilli | Lactobacillales | Streptococcaceae | Streptococcus | Streptococcus infantis |
| Bacteria | Firmicutes | Bacilli | Lactobacillales | Streptococcaceae | Streptococcus | Streptococcus intermedius |
| Bacteria | Firmicutes | Bacilli | Lactobacillales | Streptococcaceae | Streptococcus | Streptococcus oralis |
| Bacteria | Firmicutes | Bacilli | Lactobacillales | Streptococcaceae | Streptococcus | Streptococcus pseudopneumoniae |
| Bacteria | Firmicutes | Bacilli | Lactobacillales | Streptococcaceae | Streptococcus | Streptococcus salivarius |
| Bacteria | Firmicutes | Bacilli | Lactobacillales | Streptococcaceae | Streptococcus | Streptococcus sp. |
| Bacteria | Firmicutes | Bacilli | Lactobacillales | Streptococcaceae | Streptococcus | Streptococcus sp. DAT741 |
| Bacteria | Firmicutes | Bacilli | Lactobacillales | Streptococcaceae | Streptococcus | Streptococcus sp. oral clone ASCE01 |
| Bacteria | Firmicutes | Bacilli | Lactobacillales | Streptococcaceae | Streptococcus | Streptococcus sp. oral clone BW009 |
| Bacteria | Firmicutes | Bacilli | Lactobacillales | Streptococcaceae | Streptococcus | Streptococcus sp. oral clone CH016 |
| Bacteria | Firmicutes | Bacilli | Lactobacillales | Streptococcaceae | Streptococcus | Streptococcus suis |
| Bacteria | Firmicutes | Bacilli | Lactobacillales | Streptococcaceae | Streptococcus | Streptococcus uberis |
| Bacteria | Firmicutes | Clostridia | Eubacteriales | Clostridiaceae | Alkaliphilus | Alkaliphilus oremlandii |
| Bacteria | Firmicutes | Clostridia | Eubacteriales | Clostridiaceae | Clostridium | Clostridium argentinense |
| Bacteria | Firmicutes | Clostridia | Eubacteriales | Clostridiaceae | Clostridium | Clostridium diolis |
| Bacteria | Firmicutes | Clostridia | Eubacteriales | Clostridiaceae | Clostridium | Clostridium intestinale |
| Bacteria | Firmicutes | Clostridia | Eubacteriales | Clostridiaceae | Clostridium | Clostridium sardiniense |
| Bacteria | Firmicutes | Clostridia | Eubacteriales | Clostridiaceae | Clostridium | Clostridium sp. K13-19 |
| Bacteria | Firmicutes | Clostridia | Eubacteriales | Clostridiaceae | Clostridium | Clostridium sp. R6 |
| Bacteria | Firmicutes | Clostridia | Eubacteriales | Clostridiaceae | Clostridium | Clostridium sp. SY8519 |
| Bacteria | Firmicutes | Clostridia | Eubacteriales | Clostridiaceae | Clostridium | Clostridium transplantifaecale |
| Bacteria | Firmicutes | Clostridia | Eubacteriales | Clostridiaceae | Geosporobacter | Geosporobacter sp. |
| Bacteria | Firmicutes | Clostridia | Eubacteriales | Clostridiaceae | Hungatella | Hungatella hathewayi |
| Bacteria | Firmicutes | Clostridia | Eubacteriales | Clostridiales Family XII. Incertae Sedis | Fusibacter | Fusibacter paucivorans |
| Bacteria | Firmicutes | Clostridia | Eubacteriales | Clostridiales Family XIII. Incertae Sedis | Mogibacterium | Mogibacterium diversum |
| Bacteria | Firmicutes | Clostridia | Eubacteriales | Clostridiales Family XIII. Incertae Sedis | | [Eubacterium] sulci |
| Bacteria | Firmicutes | Clostridia | Eubacteriales | Clostridiales Family XVI. Incertae Sedis | Carboxydocella | Carboxydocella thermautotrophica |
| Bacteria | Firmicutes | Clostridia | Eubacteriales | Defluviitaleaceae | Defluviitalea | Defluviitalea sp. GRX3 |
| Bacteria | Firmicutes | Clostridia | Eubacteriales | Eubacteriaceae | Eubacterium | Eubacterium oxidoreducens |
| Bacteria | Firmicutes | Clostridia | Eubacteriales | Eubacteriaceae | Eubacterium | Eubacterium sp. |
| Bacteria | Firmicutes | Clostridia | Eubacteriales | Eubacteriaceae | Eubacterium | Eubacterium sp. C124b |

| | | | | | | |
|---|---|---|---|---|---|---|
| Bacteria | Firmicutes | Clostridia | Eubacteriales | Eubacteriaceae | Eubacterium | Eubacterium sp. oral clone GI038 |
| Bacteria | Firmicutes | Clostridia | Eubacteriales | Eubacteriaceae | Eubacterium | Eubacterium sp. oral clone JN088 |
| Bacteria | Firmicutes | Clostridia | Eubacteriales | Lachnospiraceae | Anaerobutyricum | Anaerobutyricum hallii |
| Bacteria | Firmicutes | Clostridia | Eubacteriales | Lachnospiraceae | Anaerocolumna | Anaerocolumna sp. CTTW |
| Bacteria | Firmicutes | Clostridia | Eubacteriales | Lachnospiraceae | Anaerostipes | Anaerostipes hadrus |
| Bacteria | Firmicutes | Clostridia | Eubacteriales | Lachnospiraceae | Anaerotignum | Anaerotignum lactatifermentans |
| Bacteria | Firmicutes | Clostridia | Eubacteriales | Lachnospiraceae | Blautia | Blautia producta |
| Bacteria | Firmicutes | Clostridia | Eubacteriales | Lachnospiraceae | Blautia | Blautia sp. SC05B48 |
| Bacteria | Firmicutes | Clostridia | Eubacteriales | Lachnospiraceae | Blautia | uncultured Blautia sp. |
| Bacteria | Firmicutes | Clostridia | Eubacteriales | Lachnospiraceae | Catonella | Catonella sp. feline oral taxon 009 |
| Bacteria | Firmicutes | Clostridia | Eubacteriales | Lachnospiraceae | Lachnoanaerobaculum | Lachnoanaerobaculum gingivalis |
| Bacteria | Firmicutes | Clostridia | Eubacteriales | Lachnospiraceae | Lachnoanaerobaculum | Lachnoanaerobaculum sp. Marseille-Q4761 |
| Bacteria | Firmicutes | Clostridia | Eubacteriales | Lachnospiraceae | Lachnoclostridium | Lachnoclostridium sp. |
| Bacteria | Firmicutes | Clostridia | Eubacteriales | Lachnospiraceae | Lachnoclostridium | Lachnoclostridium sp. Marseille-P6806 |
| Bacteria | Firmicutes | Clostridia | Eubacteriales | Lachnospiraceae | Lachnospira | Lachnospira eligens |
| Bacteria | Firmicutes | Clostridia | Eubacteriales | Lachnospiraceae | Roseburia | Roseburia intestinalis |
| Bacteria | Firmicutes | Clostridia | Eubacteriales | Lachnospiraceae | | [Eubacterium] rectale |
| Bacteria | Firmicutes | Clostridia | Eubacteriales | Lachnospiraceae | | Lachnospiraceae bacterium |
| Bacteria | Firmicutes | Clostridia | Eubacteriales | Lachnospiraceae | | Lachnospiraceae bacterium GAM79 |
| Bacteria | Firmicutes | Clostridia | Eubacteriales | Lachnospiraceae | | Lachnospiraceae bacterium KM106-2 |
| Bacteria | Firmicutes | Clostridia | Eubacteriales | Lachnospiraceae | | Lachnospiraceae bacterium MP1D12 |
| Bacteria | Firmicutes | Clostridia | Eubacteriales | Lachnospiraceae | | Lachnospiraceae bacterium sunii NSJ-8 |
| Bacteria | Firmicutes | Clostridia | Eubacteriales | Oscillospiraceae | Acetivibrio | Acetivibrio thermocellus |
| Bacteria | Firmicutes | Clostridia | Eubacteriales | Oscillospiraceae | Faecalibacterium | Faecalibacterium prausnitzii |
| Bacteria | Firmicutes | Clostridia | Eubacteriales | Oscillospiraceae | Fastidiosipila | Fastidiosipila sanguinis |
| Bacteria | Firmicutes | Clostridia | Eubacteriales | Oscillospiraceae | Flavonifractor | Flavonifractor plautii |
| Bacteria | Firmicutes | Clostridia | Eubacteriales | Oscillospiraceae | Oscillibacter | Oscillibacter sp. |
| Bacteria | Firmicutes | Clostridia | Eubacteriales | Oscillospiraceae | | [Clostridium] cellulosi |
| Bacteria | Firmicutes | Clostridia | Eubacteriales | Oscillospiraceae | | uncultured Oscillospiraceae bacterium |
| Bacteria | Firmicutes | Clostridia | Eubacteriales | Peptococcaceae | Desulfohalotomaculum | Desulfohalotomaculum peckii |
| Bacteria | Firmicutes | Clostridia | Eubacteriales | Peptococcaceae | | Peptococcaceae bacterium DCMF |
| Bacteria | Firmicutes | Clostridia | Eubacteriales | Peptostreptococcaceae | Clostridioides | Clostridioides difficile |
| Bacteria | Firmicutes | Clostridia | Eubacteriales | Peptostreptococcaceae | Intestinibacter | uncultured Intestinibacter sp. |
| Bacteria | Firmicutes | Clostridia | Eubacteriales | Peptostreptococcaceae | Paeniclostridium | Paeniclostridium ghonii |
| Bacteria | Firmicutes | Clostridia | Eubacteriales | Peptostreptococcaceae | Peptostreptococcus | Peptostreptococcus sp. |

| | | | | | | |
|---|---|---|---|---|---|---|
| Bacteria | Firmicutes | Clostridia | Eubacteriales | Peptostreptococcaceae | Peptostreptococcus | Peptostreptococcus sp. MDA2346-2 |
| Bacteria | Firmicutes | Clostridia | Eubacteriales | Peptostreptococcaceae | Peptostreptococcus | Peptostreptococcus sp. oral clone FL008 |
| Bacteria | Firmicutes | Clostridia | Eubacteriales | Peptostreptococcaceae | Romboutsia | Romboutsia lituseburensis |
| Bacteria | Firmicutes | Clostridia | Eubacteriales | Peptostreptococcaceae | Romboutsia | Romboutsia sp. Marseille-P6047 |
| Bacteria | Firmicutes | Clostridia | Eubacteriales | Peptostreptococcaceae | Sporacetigenium | Sporacetigenium mesophilum |
| Bacteria | Firmicutes | Clostridia | Eubacteriales | Peptostreptococcaceae | | Peptostreptococcaceae bacterium canine oral taxon 155 |
| Bacteria | Firmicutes | Clostridia | Eubacteriales | Peptostreptococcaceae | | Peptostreptococcaceae bacterium oral taxon 929 |
| Bacteria | Firmicutes | Clostridia | Eubacteriales | | | Clostridiales bacterium |
| Bacteria | Firmicutes | Clostridia | Eubacteriales | | | Clostridiales bacterium NS4-1 |
| Bacteria | Firmicutes | Clostridia | Halanaerobiales | Halanaerobiaceae | | uncultured Halanaerobiaceae bacterium |
| Bacteria | Firmicutes | Clostridia | | | | uncultured Clostridia bacterium |
| Bacteria | Firmicutes | Erysipelotrichia | Erysipelotrichales | Erysipelotrichaceae | Bulleidia | uncultured Bulleidia sp. |
| Bacteria | Firmicutes | Negativicutes | Acidaminococcales | Acidaminococcaceae | Phascolarctobacterium | Phascolarctobacterium faecium |
| Bacteria | Firmicutes | Negativicutes | Selenomonadales | Selenomonadaceae | Megamonas | Megamonas funiformis |
| Bacteria | Firmicutes | Negativicutes | Selenomonadales | Selenomonadaceae | Selenomonas | Selenomonas massiliensis |
| Bacteria | Firmicutes | Negativicutes | Selenomonadales | Selenomonadaceae | Selenomonas | Selenomonas sp. oral taxon 136 |
| Bacteria | Firmicutes | Negativicutes | Selenomonadales | Selenomonadaceae | Selenomonas | Selenomonas sputigena |
| Bacteria | Firmicutes | Negativicutes | Selenomonadales | Selenomonadaceae | Selenomonas | uncultured Selenomonas sp. |
| Bacteria | Firmicutes | Negativicutes | Selenomonadales | Sporomusaceae | Propionispora | Propionispora hippei |
| Bacteria | Firmicutes | Negativicutes | Veillonellales | Veillonellaceae | Dialister | Dialister invisus |
| Bacteria | Firmicutes | Negativicutes | Veillonellales | Veillonellaceae | Dialister | Dialister pneumosintes |
| Bacteria | Firmicutes | Negativicutes | Veillonellales | Veillonellaceae | Negativicoccus | Negativicoccus massiliensis |
| Bacteria | Firmicutes | Negativicutes | Veillonellales | Veillonellaceae | Veillonella | Veillonella magna |
| Bacteria | Firmicutes | Negativicutes | Veillonellales | Veillonellaceae | Veillonella | Veillonella parvula |
| Bacteria | Firmicutes | Negativicutes | Veillonellales | Veillonellaceae | Veillonella | Veillonella rodentium |
| Bacteria | Firmicutes | Tissierellia | Tissierellales | Peptoniphilaceae | Anaerococcus | Anaerococcus mediterraneensis |
| Bacteria | Firmicutes | Tissierellia | Tissierellales | Peptoniphilaceae | Anaerococcus | Anaerococcus sp. enrichment culture clone MRHull-Fe-04H |
| Bacteria | Firmicutes | Tissierellia | Tissierellales | Peptoniphilaceae | Anaerococcus | Anaerococcus vaginimassiliensis |
| Bacteria | Firmicutes | Tissierellia | Tissierellales | Peptoniphilaceae | Helcococcus | Helcococcus massiliensis |
| Bacteria | Firmicutes | Tissierellia | Tissierellales | Peptoniphilaceae | Peptoniphilus | Peptoniphilus harei |
| Bacteria | Firmicutes | Tissierellia | Tissierellales | Peptoniphilaceae | Peptoniphilus | Peptoniphilus ivorii |
| Bacteria | Firmicutes | Tissierellia | Tissierellales | Peptoniphilaceae | Peptoniphilus | Peptoniphilus sp. S467 |
| Bacteria | Firmicutes | Tissierellia | Tissierellales | Tissierellaceae | Gudongella | Gudongella oleilytica |
| Bacteria | Firmicutes | Tissierellia | Tissierellales | Tissierellaceae | Tepidimicrobium | Tepidimicrobium sp. |

| | | | | | | |
|---|---|---|---|---|---|---|
| Bacteria | Firmicutes | Tissierellia | Tissierellales | Tissierellaceae | Tepidimicrobium | Tepidimicrobium sp. GRC5 |
| Bacteria | Firmicutes | Tissierellia | Tissierellales | Tissierellaceae | Tissierella | uncultured Tissierella sp. |
| Bacteria | Firmicutes | Tissierellia | | | Ezakiella | Ezakiella massiliensis |
| Bacteria | Firmicutes | | | | | uncultured bacterium SJA-118 |
| Bacteria | Fusobacteria | Fusobacteriia | Fusobacteriales | Fusobacteriaceae | Fusobacterium | Fusobacterium nucleatum |
| Bacteria | Fusobacteria | Fusobacteriia | Fusobacteriales | Fusobacteriaceae | Fusobacterium | Fusobacterium pseudoperiodonticum |
| Bacteria | Fusobacteria | Fusobacteriia | Fusobacteriales | Fusobacteriaceae | Fusobacterium | uncultured Fusobacterium sp. |
| Bacteria | Fusobacteria | Fusobacteriia | Fusobacteriales | Leptotrichiaceae | Leptotrichia | Leptotrichia hongkongensis |
| Bacteria | Gemmatimonadetes | Gemmatimonadetes | Gemmatimonadales | Gemmatimonadaceae | Gemmatirosa | Gemmatirosa kalamazoonesis |
| Bacteria | Nitrospirae | Nitrospira | Nitrospirales | Nitrospiraceae | | uncultured Nitrospiraceae bacterium |
| Bacteria | Planctomycetes | Phycisphaerae | Tepidisphaerales | Tepidisphaeraceae | Humisphaera | Humisphaera borealis |
| Bacteria | Planctomycetes | Phycisphaerae | | | | uncultured Phycisphaerae bacterium |
| Bacteria | Planctomycetes | Planctomycetia | Gemmatales | Gemmataceae | Gemmata | uncultured Gemmata sp. |
| Bacteria | Planctomycetes | Planctomycetia | Isosphaerales | Isosphaeraceae | Aquisphaera | Aquisphaera giovannonii |
| Bacteria | Planctomycetes | Planctomycetia | Isosphaerales | Isosphaeraceae | Paludisphaera | Paludisphaera borealis |
| Bacteria | Planctomycetes | Planctomycetia | Isosphaerales | Isosphaeraceae | Singulisphaera | uncultured Singulisphaera sp. |
| Bacteria | Planctomycetes | Planctomycetia | Pirellulales | Lacipirellulaceae | Bythopirellula | Bythopirellula goksoyri |
| Bacteria | Planctomycetes | Planctomycetia | Pirellulales | Lacipirellulaceae | Lacipirellula | Lacipirellula parvula |
| Bacteria | Planctomycetes | Planctomycetia | Pirellulales | Pirellulaceae | Anatilimnocola | Anatilimnocola aggregata |
| Bacteria | Planctomycetes | Planctomycetia | Pirellulales | Pirellulaceae | Novipirellula | Novipirellula galeiformis |
| Bacteria | Planctomycetes | Planctomycetia | Planctomycetales | Planctomycetaceae | Caulifigura | Caulifigura coniformis |
| Bacteria | Planctomycetes | Planctomycetia | Planctomycetales | Planctomycetaceae | Planctomyces | Planctomyces sp. SH-PL14 |
| Bacteria | Planctomycetes | Planctomycetia | Planctomycetales | Planctomycetaceae | | Gemmata-like str. JW3-8s0 |
| Bacteria | Planctomycetes | Planctomycetia | Planctomycetales | | | uncultured sludge bacterium A1 |
| Bacteria | Planctomycetes | Planctomycetia | Planctomycetales | | | uncultured sludge bacterium A19 |
| Bacteria | Planctomycetes | Planctomycetia | Planctomycetales | | | uncultured sludge bacterium A35 |
| Bacteria | Planctomycetes | | | | | Planctomycetes bacterium |
| Bacteria | Proteobacteria | Alphaproteobacteria | Caulobacterales | Caulobacteraceae | Asticcacaulis | Asticcacaulis sp. |
| Bacteria | Proteobacteria | Alphaproteobacteria | Caulobacterales | Caulobacteraceae | Brevundimonas | Brevundimonas alba |
| Bacteria | Proteobacteria | Alphaproteobacteria | Caulobacterales | Caulobacteraceae | Brevundimonas | Brevundimonas diminuta |
| Bacteria | Proteobacteria | Alphaproteobacteria | Caulobacterales | Caulobacteraceae | Brevundimonas | Brevundimonas lenta |
| Bacteria | Proteobacteria | Alphaproteobacteria | Caulobacterales | Caulobacteraceae | Brevundimonas | Brevundimonas mediterranea |
| Bacteria | Proteobacteria | Alphaproteobacteria | Caulobacterales | Caulobacteraceae | Brevundimonas | Brevundimonas sp. 'scallop' |
| Bacteria | Proteobacteria | Alphaproteobacteria | Caulobacterales | Caulobacteraceae | Brevundimonas | Brevundimonas sp. M20 |

| | | | | | | |
|---|---|---|---|---|---|---|
| Bacteria | Proteobacteria | Alphaproteobac teria | Caulobacterales | Caulobacteraceae | Brevundimonas | Brevundimonas vancanneytii |
| Bacteria | Proteobacteria | Alphaproteobac teria | Caulobacterales | Caulobacteraceae | Caulobacter | Caulobacter flavus |
| Bacteria | Proteobacteria | Alphaproteobac teria | Caulobacterales | Caulobacteraceae | Caulobacter | Caulobacter soli |
| Bacteria | Proteobacteria | Alphaproteobac teria | Caulobacterales | Caulobacteraceae | Caulobacter | Caulobacter sp. HGR25 |
| Bacteria | Proteobacteria | Alphaproteobac teria | Caulobacterales | Caulobacteraceae | Caulobacter | Caulobacter zeae |
| Bacteria | Proteobacteria | Alphaproteobac teria | Caulobacterales | Caulobacteraceae | Phenylobacteriu m | Phenylobacterium muchangponense |
| Bacteria | Proteobacteria | Alphaproteobac teria | Caulobacterales | Caulobacteraceae | Phenylobacteriu m | Phenylobacterium parvum |
| Bacteria | Proteobacteria | Alphaproteobac teria | Caulobacterales | Caulobacteraceae | Phenylobacteriu m | uncultured Phenylobacterium sp. |
| Bacteria | Proteobacteria | Alphaproteobac teria | Caulobacterales | Caulobacteraceae | | Caulobacteraceae bacterium |
| Bacteria | Proteobacteria | Alphaproteobac teria | Hyphomicrobiale s | Amorphaceae | Amorphus | Amorphus sp. |
| Bacteria | Proteobacteria | Alphaproteobac teria | Hyphomicrobiale s | Aurantimonadacea e | Aurantimonas | Aurantimonas endophytica |
| Bacteria | Proteobacteria | Alphaproteobac teria | Hyphomicrobiale s | Aurantimonadacea e | Aureimonas | Aureimonas altamirensis |
| Bacteria | Proteobacteria | Alphaproteobac teria | Hyphomicrobiale s | Aurantimonadacea e | Aureimonas | Aureimonas leprariae |
| Bacteria | Proteobacteria | Alphaproteobac teria | Hyphomicrobiale s | Aurantimonadacea e | Aureimonas | Aureimonas sp. AU12 |
| Bacteria | Proteobacteria | Alphaproteobac teria | Hyphomicrobiale s | Aurantimonadacea e | Aureimonas | Aureimonas sp. N4 |
| Bacteria | Proteobacteria | Alphaproteobac teria | Hyphomicrobiale s | Aurantimonadacea e | Jiella | Jiella sp. |
| Bacteria | Proteobacteria | Alphaproteobac teria | Hyphomicrobiale s | Aurantimonadacea e | Martelella | Martelella mediterranea |
| Bacteria | Proteobacteria | Alphaproteobac teria | Hyphomicrobiale s | Beijerinckiaceae | Methylocella | Methylocella tundrae |
| Bacteria | Proteobacteria | Alphaproteobac teria | Hyphomicrobiale s | Boseaceae | Bosea | Bosea sp. ANAM02 |
| Bacteria | Proteobacteria | Alphaproteobac teria | Hyphomicrobiale s | Boseaceae | Bosea | Bosea sp. LC526 |
| Bacteria | Proteobacteria | Alphaproteobac teria | Hyphomicrobiale s | Bradyrhizobiaceae | Bradyrhizobium | Bradyrhizobium erythrophlei |
| Bacteria | Proteobacteria | Alphaproteobac teria | Hyphomicrobiale s | Bradyrhizobiaceae | Bradyrhizobium | Bradyrhizobium guangzhouense |
| Bacteria | Proteobacteria | Alphaproteobac teria | Hyphomicrobiale s | Bradyrhizobiaceae | Bradyrhizobium | Bradyrhizobium sp. |
| Bacteria | Proteobacteria | Alphaproteobac teria | Hyphomicrobiale s | Bradyrhizobiaceae | Bradyrhizobium | Bradyrhizobium sp. 6(2017) |
| Bacteria | Proteobacteria | Alphaproteobac teria | Hyphomicrobiale s | Bradyrhizobiaceae | Bradyrhizobium | Bradyrhizobium sp. PZHK5 |
| Bacteria | Proteobacteria | Alphaproteobac teria | Hyphomicrobiale s | Bradyrhizobiaceae | Rhodopseudomo nas | Rhodopseudomonas pseudopalustris |
| Bacteria | Proteobacteria | Alphaproteobac teria | Hyphomicrobiale s | Bradyrhizobiaceae | Rhodopseudomo nas | Rhodopseudomonas sp. |
| Bacteria | Proteobacteria | Alphaproteobac teria | Hyphomicrobiale s | Bradyrhizobiaceae | Tardiphaga | Tardiphaga robiniae |
| Bacteria | Proteobacteria | Alphaproteobac teria | Hyphomicrobiale s | Bradyrhizobiaceae | | star-like microcolonies |
| Bacteria | Proteobacteria | Alphaproteobac teria | Hyphomicrobiale s | Bradyrhizobiaceae | | uncultured Bradyrhizobiaceae bacterium |
| Bacteria | Proteobacteria | Alphaproteobac teria | Hyphomicrobiale s | Brucellaceae | Brucella | [Ochrobactrum] quorumnocens |
| Bacteria | Proteobacteria | Alphaproteobac teria | Hyphomicrobiale s | Brucellaceae | Brucella | Brucella anthropi |
| Bacteria | Proteobacteria | Alphaproteobac teria | Hyphomicrobiale s | Brucellaceae | Ochrobactrum | Ochrobactrum sp. Marseille-Q0166 |

| | | | | | | |
|---|---|---|---|---|---|---|
| Bacteria | Proteobacteria | Alphaproteobacteria | Hyphomicrobiales | Devosiaceae | Devosia | Devosia sp. 1566 |
| Bacteria | Proteobacteria | Alphaproteobacteria | Hyphomicrobiales | Devosiaceae | Youhaiella | Youhaiella tibetensis |
| Bacteria | Proteobacteria | Alphaproteobacteria | Hyphomicrobiales | Hyphomicrobiaceae | Methyloceanibacter | Methyloceanibacter caenitepidi |
| Bacteria | Proteobacteria | Alphaproteobacteria | Hyphomicrobiales | Hyphomicrobiaceae | Pedomicrobium | Pedomicrobium sp. AN4-4 |
| Bacteria | Proteobacteria | Alphaproteobacteria | Hyphomicrobiales | Kaistiaceae | Kaistia | Kaistia sp. 32K |
| Bacteria | Proteobacteria | Alphaproteobacteria | Hyphomicrobiales | Lichenihabitantaceae | Lichenihabitans | Lichenihabitans psoromatis |
| Bacteria | Proteobacteria | Alphaproteobacteria | Hyphomicrobiales | Methylobacteriaceae | Methylobacterium | Methylobacterium currus |
| Bacteria | Proteobacteria | Alphaproteobacteria | Hyphomicrobiales | Methylobacteriaceae | Methylobacterium | Methylobacterium gregans |
| Bacteria | Proteobacteria | Alphaproteobacteria | Hyphomicrobiales | Methylobacteriaceae | Methylobacterium | Methylobacterium mesophilicum |
| Bacteria | Proteobacteria | Alphaproteobacteria | Hyphomicrobiales | Methylobacteriaceae | Methylobacterium | Methylobacterium sp. 17Sr1-1 |
| Bacteria | Proteobacteria | Alphaproteobacteria | Hyphomicrobiales | Methylobacteriaceae | Methylobacterium | Methylobacterium sp. FyFern_25 |
| Bacteria | Proteobacteria | Alphaproteobacteria | Hyphomicrobiales | Methylobacteriaceae | Methylorubrum | Methylorubrum populi |
| Bacteria | Proteobacteria | Alphaproteobacteria | Hyphomicrobiales | Methylobacteriaceae | Methylorubrum | Methylorubrum zatmanii |
| Bacteria | Proteobacteria | Alphaproteobacteria | Hyphomicrobiales | Methylobacteriaceae | Microvirga | Microvirga brassicacearum |
| Bacteria | Proteobacteria | Alphaproteobacteria | Hyphomicrobiales | Methylobacteriaceae | Microvirga | Microvirga lotononidis |
| Bacteria | Proteobacteria | Alphaproteobacteria | Hyphomicrobiales | Methylobacteriaceae | Microvirga | Microvirga sp. MH5-10 |
| Bacteria | Proteobacteria | Alphaproteobacteria | Hyphomicrobiales | Methylobacteriaceae | | uncultured Methylobacteriaceae bacterium |
| Bacteria | Proteobacteria | Alphaproteobacteria | Hyphomicrobiales | Methylocystaceae | Hansschlegelia | Hansschlegelia quercus |
| Bacteria | Proteobacteria | Alphaproteobacteria | Hyphomicrobiales | Methylocystaceae | Methylosinus | Methylosinus sp. C49 |
| Bacteria | Proteobacteria | Alphaproteobacteria | Hyphomicrobiales | Parvibaculaceae | Kaustia | Kaustia mangrovi |
| Bacteria | Proteobacteria | Alphaproteobacteria | Hyphomicrobiales | Phyllobacteriaceae | Aminobacter | Aminobacter sp. MSH1 |
| Bacteria | Proteobacteria | Alphaproteobacteria | Hyphomicrobiales | Phyllobacteriaceae | Chelativorans | Chelativorans alearense |
| Bacteria | Proteobacteria | Alphaproteobacteria | Hyphomicrobiales | Phyllobacteriaceae | Mesorhizobium | Mesorhizobium oceanicum |
| Bacteria | Proteobacteria | Alphaproteobacteria | Hyphomicrobiales | Phyllobacteriaceae | Mesorhizobium | Mesorhizobium sp. 8 |
| Bacteria | Proteobacteria | Alphaproteobacteria | Hyphomicrobiales | Phyllobacteriaceae | Mesorhizobium | Mesorhizobium sp. AA22 |
| Bacteria | Proteobacteria | Alphaproteobacteria | Hyphomicrobiales | Phyllobacteriaceae | Nitratireductor | Nitratireductor kimnyeongensis |
| Bacteria | Proteobacteria | Alphaproteobacteria | Hyphomicrobiales | Phyllobacteriaceae | Phyllobacterium | Phyllobacterium loti |
| Bacteria | Proteobacteria | Alphaproteobacteria | Hyphomicrobiales | Phyllobacteriaceae | Phyllobacterium | Phyllobacterium zundukense |
| Bacteria | Proteobacteria | Alphaproteobacteria | Hyphomicrobiales | Phyllobacteriaceae | Tianweitania | Tianweitania sp. |
| Bacteria | Proteobacteria | Alphaproteobacteria | Hyphomicrobiales | Phyllobacteriaceae | | uncultured Phyllobacteriaceae bacterium |
| Bacteria | Proteobacteria | Alphaproteobacteria | Hyphomicrobiales | Rhizobiaceae | Agrobacterium | Agrobacterium sp. CPI243 |
| Bacteria | Proteobacteria | Alphaproteobacteria | Hyphomicrobiales | Rhizobiaceae | Agrobacterium | Agrobacterium sp. MA01 |
| Bacteria | Proteobacteria | Alphaproteobacteria | Hyphomicrobiales | Rhizobiaceae | Ensifer | uncultured Ensifer sp. |

| | | | | | | |
|---|---|---|---|---|---|---|
| Bacteria | Proteobacteria | Alphaproteobacteria | Hyphomicrobiales | Rhizobiaceae | Neorhizobium | Neorhizobium sp. SOG26 |
| Bacteria | Proteobacteria | Alphaproteobacteria | Hyphomicrobiales | Rhizobiaceae | Rhizobium | Rhizobium pseudoryzae |
| Bacteria | Proteobacteria | Alphaproteobacteria | Hyphomicrobiales | Rhizobiaceae | Rhizobium | Rhizobium sp. Q54 |
| Bacteria | Proteobacteria | Alphaproteobacteria | Hyphomicrobiales | Rhizobiaceae | Rhizobium | uncultured Rhizobium sp. |
| Bacteria | Proteobacteria | Alphaproteobacteria | Hyphomicrobiales | Rhizobiaceae | Sinorhizobium | Sinorhizobium meliloti |
| Bacteria | Proteobacteria | Alphaproteobacteria | Hyphomicrobiales | Rhizobiaceae | Sinorhizobium | Sinorhizobium sp. RAC02 |
| Bacteria | Proteobacteria | Alphaproteobacteria | Hyphomicrobiales | Rhizobiaceae | | arsenite-oxidising bacterium NT-25 |
| Bacteria | Proteobacteria | Alphaproteobacteria | Hyphomicrobiales | Rhizobiaceae | | Rhizobiaceae bacterium CRRU44 |
| Bacteria | Proteobacteria | Alphaproteobacteria | Hyphomicrobiales | Stappiaceae | Labrenzia | Labrenzia aggregata |
| Bacteria | Proteobacteria | Alphaproteobacteria | Hyphomicrobiales | Stappiaceae | Pannonibacter | Pannonibacter phragmitetus |
| Bacteria | Proteobacteria | Alphaproteobacteria | Hyphomicrobiales | Stappiaceae | Stappia | Stappia indica |
| Bacteria | Proteobacteria | Alphaproteobacteria | Hyphomicrobiales | Xanthobacteraceae | Labrys | Labrys sp. KNU-23 |
| Bacteria | Proteobacteria | Alphaproteobacteria | Hyphomicrobiales | Xanthobacteraceae | Xanthobacter | Xanthobacter sp. W31 |
| Bacteria | Proteobacteria | Alphaproteobacteria | Hyphomicrobiales | | Candidatus Tokpelaia | Candidatus Tokpelaia hoelldoblerii |
| Bacteria | Proteobacteria | Alphaproteobacteria | Hyphomicrobiales | | Nordella | Nordella sp. HKS 07 |
| Bacteria | Proteobacteria | Alphaproteobacteria | Hyphomicrobiales | | | Rhizobiales bacterium IZ6 |
| Bacteria | Proteobacteria | Alphaproteobacteria | Hyphomicrobiales | | | Rhizobiales bacterium RSB |
| Bacteria | Proteobacteria | Alphaproteobacteria | Rhodobacterales | Rhodobacteraceae | Acidimangrovimonas | Acidimangrovimonas sediminis |
| Bacteria | Proteobacteria | Alphaproteobacteria | Rhodobacterales | Rhodobacteraceae | Aestuarium | Aestuarium zhoushanense |
| Bacteria | Proteobacteria | Alphaproteobacteria | Rhodobacterales | Rhodobacteraceae | Epibacterium | Epibacterium mobile |
| Bacteria | Proteobacteria | Alphaproteobacteria | Rhodobacterales | Rhodobacteraceae | Paracoccus | Paracoccus homiensis |
| Bacteria | Proteobacteria | Alphaproteobacteria | Rhodobacterales | Rhodobacteraceae | Paracoccus | Paracoccus jeotgali |
| Bacteria | Proteobacteria | Alphaproteobacteria | Rhodobacterales | Rhodobacteraceae | Paracoccus | Paracoccus liaowanqingii |
| Bacteria | Proteobacteria | Alphaproteobacteria | Rhodobacterales | Rhodobacteraceae | Paracoccus | Paracoccus sp. DFMS01 |
| Bacteria | Proteobacteria | Alphaproteobacteria | Rhodobacterales | Rhodobacteraceae | Paracoccus | Paracoccus yeei |
| Bacteria | Proteobacteria | Alphaproteobacteria | Rhodobacterales | Rhodobacteraceae | Pseudorhodobacter | Pseudorhodobacter sp. |
| Bacteria | Proteobacteria | Alphaproteobacteria | Rhodobacterales | Rhodobacteraceae | Pseudorhodobacter | Pseudorhodobacter sp. 7B-639 |
| Bacteria | Proteobacteria | Alphaproteobacteria | Rhodobacterales | Rhodobacteraceae | Rubellimicrobium | Rubellimicrobium rubrum |
| Bacteria | Proteobacteria | Alphaproteobacteria | Rhodobacterales | Rhodobacteraceae | Rubellimicrobium | Rubellimicrobium sp. |
| Bacteria | Proteobacteria | Alphaproteobacteria | Rhodobacterales | Rhodobacteraceae | Sulfitobacter | Sulfitobacter sp. SK011 |
| Bacteria | Proteobacteria | Alphaproteobacteria | Rhodobacterales | Rhodobacteraceae | Tabrizicola | Tabrizicola piscis |
| Bacteria | Proteobacteria | Alphaproteobacteria | Rhodobacterales | Rhodobacteraceae | Tabrizicola | Tabrizicola sp. |
| Bacteria | Proteobacteria | Alphaproteobacteria | Rhodobacterales | | | uncultured Rhodobacterales bacterium |

| | | | | | | |
|---|---|---|---|---|---|---|
| Bacteria | Proteobacteria | Alphaproteobacteria | Rhodospirillales | Acetobacteraceae | Acetobacter | Acetobacter ghanensis |
| Bacteria | Proteobacteria | Alphaproteobacteria | Rhodospirillales | Acetobacteraceae | Acidiphilium | uncultured Acidiphilium sp. |
| Bacteria | Proteobacteria | Alphaproteobacteria | Rhodospirillales | Acetobacteraceae | Ameyamaea | Ameyamaea sp. Amy1 |
| Bacteria | Proteobacteria | Alphaproteobacteria | Rhodospirillales | Acetobacteraceae | Asaia | Asaia bogorensis |
| Bacteria | Proteobacteria | Alphaproteobacteria | Rhodospirillales | Acetobacteraceae | Commensalibacter | Commensalibacter sp. ESL0284 |
| Bacteria | Proteobacteria | Alphaproteobacteria | Rhodospirillales | Acetobacteraceae | Entomobacter | Entomobacter blattae |
| Bacteria | Proteobacteria | Alphaproteobacteria | Rhodospirillales | Acetobacteraceae | Gluconobacter | Gluconobacter sp. |
| Bacteria | Proteobacteria | Alphaproteobacteria | Rhodospirillales | Acetobacteraceae | Gluconobacter | Gluconobacter thailandicus |
| Bacteria | Proteobacteria | Alphaproteobacteria | Rhodospirillales | Acetobacteraceae | Komagataeibacter | Komagataeibacter xylinus |
| Bacteria | Proteobacteria | Alphaproteobacteria | Rhodospirillales | Acetobacteraceae | Neoasaia | Neoasaia sp. |
| Bacteria | Proteobacteria | Alphaproteobacteria | Rhodospirillales | Acetobacteraceae | Paracraurococcus | uncultured Paracraurococcus sp. |
| Bacteria | Proteobacteria | Alphaproteobacteria | Rhodospirillales | Acetobacteraceae | Roseococcus | Roseococcus sp. |
| Bacteria | Proteobacteria | Alphaproteobacteria | Rhodospirillales | Acetobacteraceae | Roseomonas | Roseomonas aquatica |
| Bacteria | Proteobacteria | Alphaproteobacteria | Rhodospirillales | Acetobacteraceae | Roseomonas | Roseomonas cervicalis |
| Bacteria | Proteobacteria | Alphaproteobacteria | Rhodospirillales | Acetobacteraceae | Roseomonas | Roseomonas globiformis |
| Bacteria | Proteobacteria | Alphaproteobacteria | Rhodospirillales | Acetobacteraceae | Roseomonas | Roseomonas mucosa |
| Bacteria | Proteobacteria | Alphaproteobacteria | Rhodospirillales | Acetobacteraceae | Roseomonas | Roseomonas nepalensis |
| Bacteria | Proteobacteria | Alphaproteobacteria | Rhodospirillales | Acetobacteraceae | Roseomonas | Roseomonas sp. 1318 |
| Bacteria | Proteobacteria | Alphaproteobacteria | Rhodospirillales | Acetobacteraceae | Roseomonas | Roseomonas sp. 546 |
| Bacteria | Proteobacteria | Alphaproteobacteria | Rhodospirillales | Acetobacteraceae | Roseomonas | Roseomonas sp. CPCC 101081 |
| Bacteria | Proteobacteria | Alphaproteobacteria | Rhodospirillales | Acetobacteraceae | Saccharibacter | Saccharibacter floricola |
| Bacteria | Proteobacteria | Alphaproteobacteria | Rhodospirillales | Acetobacteraceae | Stella | Stella sp. ATCC 35155 |
| Bacteria | Proteobacteria | Alphaproteobacteria | Rhodospirillales | Acetobacteraceae | Stella | uncultured Stella sp. |
| Bacteria | Proteobacteria | Alphaproteobacteria | Rhodospirillales | Azospirillaceae | Azospirillum | Azospirillum sp. Azo super |
| Bacteria | Proteobacteria | Alphaproteobacteria | Rhodospirillales | Azospirillaceae | Rhodocista | uncultured Rhodocista sp. |
| Bacteria | Proteobacteria | Alphaproteobacteria | Rhodospirillales | Azospirillaceae | Skermanella | Skermanella mucosa |
| Bacteria | Proteobacteria | Alphaproteobacteria | Rhodospirillales | Azospirillaceae | Skermanella | Skermanella pratensis |
| Bacteria | Proteobacteria | Alphaproteobacteria | Rhodospirillales | Azospirillaceae | Skermanella | Skermanella sp. |
| Bacteria | Proteobacteria | Alphaproteobacteria | Rhodospirillales | Azospirillaceae | Skermanella | Skermanella sp. Py-2-1 |
| Bacteria | Proteobacteria | Alphaproteobacteria | Rhodospirillales | Azospirillaceae | Skermanella | uncultured Skermanella sp. |
| Bacteria | Proteobacteria | Alphaproteobacteria | Rhodospirillales | Geminicoccaceae | Candidatus Halyseomicrobium | Candidatus Halyseomicrobium bavaricum |
| Bacteria | Proteobacteria | Alphaproteobacteria | Rhodospirillales | Rhodospirillaceae | Ferruginivarius | Ferruginivarius sediminum |
| Bacteria | Proteobacteria | Alphaproteobacteria | Rhodospirillales | Rhodospirillaceae | Hypericibacter | Hypericibacter terrae |

| | | | | | | |
|---|---|---|---|---|---|---|
| Bacteria | Proteobacteria | Alphaproteobacteria | Rhodospirillales | Rhodospirillaceae | Phaeospirillum | Phaeospirillum sp. JA815 |
| Bacteria | Proteobacteria | Alphaproteobacteria | Rhodospirillales | | Reyranella | Reyranella sp. |
| Bacteria | Proteobacteria | Alphaproteobacteria | Rickettsiales | Anaplasmataceae | Wolbachia | Wolbachia endosymbiont of Icosta sp. 2 ES-2018 |
| Bacteria | Proteobacteria | Alphaproteobacteria | Sphingomonadales | Erythrobacteraceae | Altererythrobacter | Altererythrobacter sp. TH136 |
| Bacteria | Proteobacteria | Alphaproteobacteria | Sphingomonadales | Erythrobacteraceae | Croceibacterium | Croceibacterium mercuriale |
| Bacteria | Proteobacteria | Alphaproteobacteria | Sphingomonadales | Erythrobacteraceae | Qipengyuania | Qipengyuania sediminis |
| Bacteria | Proteobacteria | Alphaproteobacteria | Sphingomonadales | Sphingomonadaceae | Blastomonas | uncultured Blastomonas sp. |
| Bacteria | Proteobacteria | Alphaproteobacteria | Sphingomonadales | Sphingomonadaceae | Novosphingobium | Novosphingobium fuchskuhlense |
| Bacteria | Proteobacteria | Alphaproteobacteria | Sphingomonadales | Sphingomonadaceae | Novosphingobium | Novosphingobium panipatense |
| Bacteria | Proteobacteria | Alphaproteobacteria | Sphingomonadales | Sphingomonadaceae | Novosphingobium | Novosphingobium pokkalii |
| Bacteria | Proteobacteria | Alphaproteobacteria | Sphingomonadales | Sphingomonadaceae | Novosphingobium | Novosphingobium sp. enrichment culture clone Van61 |
| Bacteria | Proteobacteria | Alphaproteobacteria | Sphingomonadales | Sphingomonadaceae | Novosphingobium | Novosphingobium sp. YIM78034 |
| Bacteria | Proteobacteria | Alphaproteobacteria | Sphingomonadales | Sphingomonadaceae | Novosphingobium | uncultured Novosphingobium sp. |
| Bacteria | Proteobacteria | Alphaproteobacteria | Sphingomonadales | Sphingomonadaceae | Sphingobium | Sphingobium amiense |
| Bacteria | Proteobacteria | Alphaproteobacteria | Sphingomonadales | Sphingomonadaceae | Sphingobium | Sphingobium sp. CAP-1 |
| Bacteria | Proteobacteria | Alphaproteobacteria | Sphingomonadales | Sphingomonadaceae | Sphingobium | Sphingobium sp. W2.09-52 |
| Bacteria | Proteobacteria | Alphaproteobacteria | Sphingomonadales | Sphingomonadaceae | Sphingobium | Sphingobium yanoikuyae |
| Bacteria | Proteobacteria | Alphaproteobacteria | Sphingomonadales | Sphingomonadaceae | Sphingobium | uncultured Sphingobium sp. |
| Bacteria | Proteobacteria | Alphaproteobacteria | Sphingomonadales | Sphingomonadaceae | Sphingomonas | Sphingomonas formosensis |
| Bacteria | Proteobacteria | Alphaproteobacteria | Sphingomonadales | Sphingomonadaceae | Sphingomonas | Sphingomonas insulae |
| Bacteria | Proteobacteria | Alphaproteobacteria | Sphingomonadales | Sphingomonadaceae | Sphingomonas | Sphingomonas olei |
| Bacteria | Proteobacteria | Alphaproteobacteria | Sphingomonadales | Sphingomonadaceae | Sphingomonas | Sphingomonas paucimobilis |
| Bacteria | Proteobacteria | Alphaproteobacteria | Sphingomonadales | Sphingomonadaceae | Sphingomonas | Sphingomonas sp. 2378 |
| Bacteria | Proteobacteria | Alphaproteobacteria | Sphingomonadales | Sphingomonadaceae | Sphingomonas | Sphingomonas sp. 3R-14 |
| Bacteria | Proteobacteria | Alphaproteobacteria | Sphingomonadales | Sphingomonadaceae | Sphingomonas | Sphingomonas sp. LK11 |
| Bacteria | Proteobacteria | Alphaproteobacteria | Sphingomonadales | Sphingomonadaceae | Sphingomonas | Sphingomonas sp. MK52 |
| Bacteria | Proteobacteria | Alphaproteobacteria | Sphingomonadales | Sphingomonadaceae | Sphingomonas | Sphingomonas sp. PDD-23b-6 |
| Bacteria | Proteobacteria | Alphaproteobacteria | Sphingomonadales | Sphingomonadaceae | Sphingopyxis | Sphingopyxis sp. |
| Bacteria | Proteobacteria | Alphaproteobacteria | Sphingomonadales | Sphingomonadaceae | Sphingopyxis | Sphingopyxis sp. PAMC25046 |
| Bacteria | Proteobacteria | Alphaproteobacteria | Sphingomonadales | Sphingomonadaceae | Sphingorhabdus | Sphingorhabdus lacus |
| Bacteria | Proteobacteria | Alphaproteobacteria | Sphingomonadales | Sphingomonadaceae | Zymomonas | Zymomonas mobilis |
| Bacteria | Proteobacteria | Alphaproteobacteria | Sphingomonadales | Sphingosinicellaceae | Sphingosinicella | Sphingosinicella sp. BN140058 |
| Bacteria | Proteobacteria | Alphaproteobacteria | | | | alpha proteobacterium P73 |

| | | | | | | |
|---|---|---|---|---|---|---|
| Bacteria | Proteobacteria | Alphaproteobacteria | | | | alpha proteobacterium XYY-2015 |
| Bacteria | Proteobacteria | Alphaproteobacteria | | | | Alphaproteobacteria bacterium |
| Bacteria | Proteobacteria | Alphaproteobacteria | | | | uncultured sludge bacterium H46 |
| Bacteria | Proteobacteria | Betaproteobacteria | Burkholderiales | Alcaligenaceae | Achromobacter | Achromobacter sp. |
| Bacteria | Proteobacteria | Betaproteobacteria | Burkholderiales | Alcaligenaceae | Achromobacter | Achromobacter xylosoxidans |
| Bacteria | Proteobacteria | Betaproteobacteria | Burkholderiales | Alcaligenaceae | Alcaligenes | Alcaligenes faecalis |
| Bacteria | Proteobacteria | Betaproteobacteria | Burkholderiales | Alcaligenaceae | Alcaligenes | uncultured Alcaligenes sp. |
| Bacteria | Proteobacteria | Betaproteobacteria | Burkholderiales | Alcaligenaceae | Bordetella | Bordetella sp. ScyaBb-1 |
| Bacteria | Proteobacteria | Betaproteobacteria | Burkholderiales | Alcaligenaceae | Candidimonas | Candidimonas sp. |
| Bacteria | Proteobacteria | Betaproteobacteria | Burkholderiales | Alcaligenaceae | Derxia | uncultured Derxia sp. |
| Bacteria | Proteobacteria | Betaproteobacteria | Burkholderiales | Alcaligenaceae | Orrella | Orrella sp. |
| Bacteria | Proteobacteria | Betaproteobacteria | Burkholderiales | Alcaligenaceae | Pigmentiphaga | Pigmentiphaga sp. H8 |
| Bacteria | Proteobacteria | Betaproteobacteria | Burkholderiales | Burkholderiaceae | Burkholderia | Burkholderia contaminans |
| Bacteria | Proteobacteria | Betaproteobacteria | Burkholderiales | Burkholderiaceae | Burkholderia | Burkholderia seminalis |
| Bacteria | Proteobacteria | Betaproteobacteria | Burkholderiales | Burkholderiaceae | Burkholderia | Burkholderia sp. symbiont of Dicranocephalus albipes |
| Bacteria | Proteobacteria | Betaproteobacteria | Burkholderiales | Burkholderiaceae | Burkholderia | Burkholderia sp. THE68 |
| Bacteria | Proteobacteria | Betaproteobacteria | Burkholderiales | Burkholderiaceae | Burkholderia | uncultured Burkholderia sp. |
| Bacteria | Proteobacteria | Betaproteobacteria | Burkholderiales | Burkholderiaceae | Caballeronia | Caballeronia sp. SBC2 |
| Bacteria | Proteobacteria | Betaproteobacteria | Burkholderiales | Burkholderiaceae | Caballeronia | Caballeronia udeis |
| Bacteria | Proteobacteria | Betaproteobacteria | Burkholderiales | Burkholderiaceae | Candidatus Protistobacter | Candidatus Protistobacter heckmanni |
| Bacteria | Proteobacteria | Betaproteobacteria | Burkholderiales | Burkholderiaceae | Cupriavidus | Cupriavidus gilardii |
| Bacteria | Proteobacteria | Betaproteobacteria | Burkholderiales | Burkholderiaceae | Cupriavidus | Cupriavidus pinatubonensis |
| Bacteria | Proteobacteria | Betaproteobacteria | Burkholderiales | Burkholderiaceae | Lautropia | Lautropia mirabilis |
| Bacteria | Proteobacteria | Betaproteobacteria | Burkholderiales | Burkholderiaceae | Lautropia | Lautropia sp. canine oral taxon 060 |
| Bacteria | Proteobacteria | Betaproteobacteria | Burkholderiales | Burkholderiaceae | Limnobacter | Limnobacter thiooxidans |
| Bacteria | Proteobacteria | Betaproteobacteria | Burkholderiales | Burkholderiaceae | Pandoraea | Pandoraea pnomenusa |
| Bacteria | Proteobacteria | Betaproteobacteria | Burkholderiales | Burkholderiaceae | Paraburkholderia | Paraburkholderia bannensis |
| Bacteria | Proteobacteria | Betaproteobacteria | Burkholderiales | Comamonadaceae | Acidovorax | Acidovorax antarcticus |
| Bacteria | Proteobacteria | Betaproteobacteria | Burkholderiales | Comamonadaceae | Acidovorax | Acidovorax sp. |
| Bacteria | Proteobacteria | Betaproteobacteria | Burkholderiales | Comamonadaceae | Acidovorax | Acidovorax sp. enrichment culture clone Van23 |
| Bacteria | Proteobacteria | Betaproteobacteria | Burkholderiales | Comamonadaceae | Acidovorax | uncultured Acidovorax sp. |
| Bacteria | Proteobacteria | Betaproteobacteria | Burkholderiales | Comamonadaceae | Brachymonas | uncultured Brachymonas sp. |
| Bacteria | Proteobacteria | Betaproteobacteria | Burkholderiales | Comamonadaceae | Comamonas | Comamonas thiooxydans |

| | | | | | | |
|---|---|---|---|---|---|---|
| Bacteria | Proteobacteria | Betaproteobacteria | Burkholderiales | Comamonadaceae | Curvibacter | Curvibacter sp. NFH12 |
| Bacteria | Proteobacteria | Betaproteobacteria | Burkholderiales | Comamonadaceae | Curvibacter | Curvibacter sp. W2.09-301r |
| Bacteria | Proteobacteria | Betaproteobacteria | Burkholderiales | Comamonadaceae | Hydrogenophaga | uncultured Hydrogenophaga sp. |
| Bacteria | Proteobacteria | Betaproteobacteria | Burkholderiales | Comamonadaceae | Hylemonella | Hylemonella sp. 32AD14 |
| Bacteria | Proteobacteria | Betaproteobacteria | Burkholderiales | Comamonadaceae | Limnohabitans | Limnohabitans sp. |
| Bacteria | Proteobacteria | Betaproteobacteria | Burkholderiales | Comamonadaceae | Ottowia | Ottowia oryzae |
| Bacteria | Proteobacteria | Betaproteobacteria | Burkholderiales | Comamonadaceae | Polaromonas | Polaromonas sp. GM1 |
| Bacteria | Proteobacteria | Betaproteobacteria | Burkholderiales | Comamonadaceae | Polaromonas | Polaromonas sp. Pch-P |
| Bacteria | Proteobacteria | Betaproteobacteria | Burkholderiales | Comamonadaceae | Polaromonas | uncultured Polaromonas sp. |
| Bacteria | Proteobacteria | Betaproteobacteria | Burkholderiales | Comamonadaceae | Pseudacidovorax | Pseudacidovorax intermedius |
| Bacteria | Proteobacteria | Betaproteobacteria | Burkholderiales | Comamonadaceae | Simplicispira | Simplicispira suum |
| Bacteria | Proteobacteria | Betaproteobacteria | Burkholderiales | Comamonadaceae | Variovorax | Variovorax sp. enrichment culture clone Van40 |
| Bacteria | Proteobacteria | Betaproteobacteria | Burkholderiales | Comamonadaceae | Variovorax | Variovorax sp. WDL1 |
| Bacteria | Proteobacteria | Betaproteobacteria | Burkholderiales | Comamonadaceae | Xenophilus | Xenophilus aerolatus |
| Bacteria | Proteobacteria | Betaproteobacteria | Burkholderiales | Oxalobacteraceae | Collimonas | Collimonas arenae |
| Bacteria | Proteobacteria | Betaproteobacteria | Burkholderiales | Oxalobacteraceae | Herbaspirillum | Herbaspirillum chlorophenolicum |
| Bacteria | Proteobacteria | Betaproteobacteria | Burkholderiales | Oxalobacteraceae | Herbaspirillum | Herbaspirillum lusitanum |
| Bacteria | Proteobacteria | Betaproteobacteria | Burkholderiales | Oxalobacteraceae | Herbaspirillum | Herbaspirillum seropedicae |
| Bacteria | Proteobacteria | Betaproteobacteria | Burkholderiales | Oxalobacteraceae | Herbaspirillum | uncultured Herbaspirillum sp. |
| Bacteria | Proteobacteria | Betaproteobacteria | Burkholderiales | Oxalobacteraceae | Janthinobacterium | Janthinobacterium agaricidamnosum |
| Bacteria | Proteobacteria | Betaproteobacteria | Burkholderiales | Oxalobacteraceae | Janthinobacterium | Janthinobacterium sp. |
| Bacteria | Proteobacteria | Betaproteobacteria | Burkholderiales | Oxalobacteraceae | Janthinobacterium | Janthinobacterium sp. Marseille-P9896 |
| Bacteria | Proteobacteria | Betaproteobacteria | Burkholderiales | Oxalobacteraceae | Massilia | Massilia alkalitolerans |
| Bacteria | Proteobacteria | Betaproteobacteria | Burkholderiales | Oxalobacteraceae | Massilia | Massilia brevitalea |
| Bacteria | Proteobacteria | Betaproteobacteria | Burkholderiales | Oxalobacteraceae | Massilia | Massilia putida |
| Bacteria | Proteobacteria | Betaproteobacteria | Burkholderiales | Oxalobacteraceae | Massilia | Massilia suwonensis |
| Bacteria | Proteobacteria | Betaproteobacteria | Burkholderiales | Oxalobacteraceae | Noviherbaspirillum | Noviherbaspirillum sp. UKPF54 |
| Bacteria | Proteobacteria | Betaproteobacteria | Burkholderiales | Oxalobacteraceae | Oxalicibacterium | Oxalicibacterium solurbis |
| Bacteria | Proteobacteria | Betaproteobacteria | Burkholderiales | Oxalobacteraceae | Oxalobacter | uncultured Oxalobacter sp. |
| Bacteria | Proteobacteria | Betaproteobacteria | Burkholderiales | Oxalobacteraceae | Undibacterium | Undibacterium sp. YM2 |
| Bacteria | Proteobacteria | Betaproteobacteria | Burkholderiales | Oxalobacteraceae | | Oxalobacteraceae bacterium |
| Bacteria | Proteobacteria | Betaproteobacteria | Burkholderiales | | Aquabacterium | Aquabacterium citratiphilum |
| Bacteria | Proteobacteria | Betaproteobacteria | Burkholderiales | | Aquabacterium | Aquabacterium sp. |

| | | | | | | |
|---|---|---|---|---|---|---|
| Bacteria | Proteobacteria | Betaproteobacteria | Burkholderiales | | Mitsuaria | Mitsuaria sp. 7 |
| Bacteria | Proteobacteria | Betaproteobacteria | Burkholderiales | | Rhizobacter | Rhizobacter dauci |
| Bacteria | Proteobacteria | Betaproteobacteria | Burkholderiales | | Xylophilus | Xylophilus sp. |
| Bacteria | Proteobacteria | Betaproteobacteria | Burkholderiales | | | [Polyangium] brachysporum |
| Bacteria | Proteobacteria | Betaproteobacteria | Neisseriales | Chromobacteriaceae | Aquaspirillum | Aquaspirillum putridiconchylium |
| Bacteria | Proteobacteria | Betaproteobacteria | Neisseriales | Chromobacteriaceae | Aquaspirillum | Aquaspirillum sp. LM1 |
| Bacteria | Proteobacteria | Betaproteobacteria | Neisseriales | Chromobacteriaceae | Pseudogulbenkiania | Pseudogulbenkiania sp. NH8B |
| Bacteria | Proteobacteria | Betaproteobacteria | Neisseriales | Neisseriaceae | Kingella | Kingella bonacorsii |
| Bacteria | Proteobacteria | Betaproteobacteria | Neisseriales | Neisseriaceae | Neisseria | Neisseria sp. feline oral taxon 145 |
| Bacteria | Proteobacteria | Betaproteobacteria | Neisseriales | Neisseriaceae | Neisseria | Neisseria sp. oral taxon 014 |
| Bacteria | Proteobacteria | Betaproteobacteria | Neisseriales | Neisseriaceae | | Neisseriaceae bacterium |
| Bacteria | Proteobacteria | Betaproteobacteria | Nitrosomonadales | Gallionellaceae | Candidatus Nitrotoga | Candidatus Nitrotoga sp. AM1P |
| Bacteria | Proteobacteria | Betaproteobacteria | Nitrosomonadales | Methylophilaceae | Candidatus Methylopumilus | Candidatus Methylopumilus turicensis |
| Bacteria | Proteobacteria | Betaproteobacteria | Nitrosomonadales | Methylophilaceae | Methylobacillus | Methylobacillus flagellatus |
| Bacteria | Proteobacteria | Betaproteobacteria | Nitrosomonadales | Nitrosomonadaceae | Nitrosomonas | Nitrosomonas ureae |
| Bacteria | Proteobacteria | Betaproteobacteria | Nitrosomonadales | Nitrosomonadaceae | Nitrosospira | Nitrosospira briensis |
| Bacteria | Proteobacteria | Betaproteobacteria | Nitrosomonadales | Nitrosomonadaceae | Nitrosospira | Nitrosospira sp. NRS527 |
| Bacteria | Proteobacteria | Betaproteobacteria | Nitrosomonadales | Nitrosomonadaceae | Nitrosospira | uncultured Nitrosospira sp. |
| Bacteria | Proteobacteria | Betaproteobacteria | Nitrosomonadales | Sterolibacteriaceae | Denitratisoma | uncultured Denitratisoma sp. |
| Bacteria | Proteobacteria | Betaproteobacteria | Rhodocyclales | Azonexaceae | Dechloromonas | Dechloromonas sp. |
| Bacteria | Proteobacteria | Betaproteobacteria | Rhodocyclales | Rhodocyclaceae | Azospira | Azospira sp. I09 |
| Bacteria | Proteobacteria | Betaproteobacteria | Rhodocyclales | Rhodocyclaceae | Azospira | uncultured Azospira sp. |
| Bacteria | Proteobacteria | Betaproteobacteria | Rhodocyclales | Rhodocyclaceae | Propionivibrio | uncultured Propionivibrio sp. |
| Bacteria | Proteobacteria | Betaproteobacteria | Rhodocyclales | Zoogloeaceae | Azoarcus | Azoarcus sp. M9-3-2 |
| Bacteria | Proteobacteria | Betaproteobacteria | Rhodocyclales | Zoogloeaceae | Azoarcus | Azoarcus sp. PH002 |
| Bacteria | Proteobacteria | Betaproteobacteria | | | | uncultured bacterium SJA-21 |
| Bacteria | Proteobacteria | Deltaproteobacteria | Desulfovibrionales | Desulfovibrionaceae | Desulfovibrio | Desulfovibrio desulfuricans |
| Bacteria | Proteobacteria | Deltaproteobacteria | Desulfovibrionales | Desulfovibrionaceae | | uncultured Desulfovibrionaceae bacterium |
| Bacteria | Proteobacteria | Deltaproteobacteria | Myxococcales | Archangiaceae | Archangium | Archangium violaceum |
| Bacteria | Proteobacteria | Deltaproteobacteria | Myxococcales | Myxococcaceae | Corallococcus | Corallococcus coralloides |
| Bacteria | Proteobacteria | Deltaproteobacteria | Myxococcales | Myxococcaceae | Corallococcus | Corallococcus macrosporus |
| Bacteria | Proteobacteria | Deltaproteobacteria | Myxococcales | Polyangiaceae | Chondromyces | Chondromyces crocatus |
| Bacteria | Proteobacteria | Deltaproteobacteria | Myxococcales | Polyangiaceae | Chondromyces | Chondromyces pediculatus |

| | | | | | | |
|---|---|---|---|---|---|---|
| Bacteria | Proteobacteria | Deltaproteobacteria | Myxococcales | Polyangiaceae | Sorangium | Sorangium sp. |
| Bacteria | Proteobacteria | Deltaproteobacteria | Myxococcales | Polyangiaceae | Sorangium | uncultured Sorangium sp. |
| Bacteria | Proteobacteria | Deltaproteobacteria | Myxococcales | Polyangiaceae | | uncultured Polyangiaceae bacterium |
| Bacteria | Proteobacteria | Deltaproteobacteria | Myxococcales | | Enhygromyxa | uncultured Enhygromyxa sp. |
| Bacteria | Proteobacteria | Deltaproteobacteria | Myxococcales | | | Myxococcales bacterium |
| Bacteria | Proteobacteria | Epsilonproteobacteria | Campylobacterales | Campylobacteraceae | Campylobacter | Campylobacter gracilis |
| Bacteria | Proteobacteria | Epsilonproteobacteria | Campylobacterales | Sulfurovaceae | Sulfurovum | Sulfurovum lithotrophicum |
| Bacteria | Proteobacteria | Gammaproteobacteria | Aeromonadales | Aeromonadaceae | Aeromonas | Aeromonas veronii |
| Bacteria | Proteobacteria | Gammaproteobacteria | Aeromonadales | Aeromonadaceae | Aeromonas | uncultured Aeromonas sp. |
| Bacteria | Proteobacteria | Gammaproteobacteria | Alteromonadales | Alteromonadaceae | Paraglaciecola | Paraglaciecola sp. L3A3 |
| Bacteria | Proteobacteria | Gammaproteobacteria | Alteromonadales | Colwelliaceae | Thalassomonas | uncultured Thalassomonas sp. |
| Bacteria | Proteobacteria | Gammaproteobacteria | Alteromonadales | Shewanellaceae | Shewanella | Shewanella seohaensis |
| Bacteria | Proteobacteria | Gammaproteobacteria | Alteromonadales | Shewanellaceae | Shewanella | Shewanella sp. |
| Bacteria | Proteobacteria | Gammaproteobacteria | Chromatiales | Chromatiaceae | Candidatus Thiosymbion | Candidatus Thiosymbion oneisti |
| Bacteria | Proteobacteria | Gammaproteobacteria | Chromatiales | Chromatiaceae | Phaeochromatium | Phaeochromatium fluminis |
| Bacteria | Proteobacteria | Gammaproteobacteria | Chromatiales | Chromatiaceae | Rheinheimera | Rheinheimera baltica |
| Bacteria | Proteobacteria | Gammaproteobacteria | Chromatiales | Chromatiaceae | Rheinheimera | uncultured Rheinheimera sp. |
| Bacteria | Proteobacteria | Gammaproteobacteria | Chromatiales | Ectothiorhodospiraceae | Candidatus Macondimonas | Candidatus Macondimonas diazotrophica |
| Bacteria | Proteobacteria | Gammaproteobacteria | Enterobacterales | Bruguierivoracaceae | Sodalis | Candidatus Sodalis pierantonius |
| Bacteria | Proteobacteria | Gammaproteobacteria | Enterobacterales | Bruguierivoracaceae | Sodalis | Sodalis endosymbiont of Cardiaspina maniformis |
| Bacteria | Proteobacteria | Gammaproteobacteria | Enterobacterales | Bruguierivoracaceae | Sodalis | Sodalis endosymbiont of Microlynchia galapagoensis |
| Bacteria | Proteobacteria | Gammaproteobacteria | Enterobacterales | Bruguierivoracaceae | Sodalis | Sodalis glossinidius |
| Bacteria | Proteobacteria | Gammaproteobacteria | Enterobacterales | Enterobacteriaceae | Atlantibacter | Atlantibacter sp. |
| Bacteria | Proteobacteria | Gammaproteobacteria | Enterobacterales | Enterobacteriaceae | Buttiauxella | Buttiauxella agrestis |
| Bacteria | Proteobacteria | Gammaproteobacteria | Enterobacterales | Enterobacteriaceae | Candidatus Kleidoceria | Candidatus Kleidoceria schneideri |
| Bacteria | Proteobacteria | Gammaproteobacteria | Enterobacterales | Enterobacteriaceae | Cedecea | Cedecea lapagei |
| Bacteria | Proteobacteria | Gammaproteobacteria | Enterobacterales | Enterobacteriaceae | Citrobacter | Citrobacter farmeri |
| Bacteria | Proteobacteria | Gammaproteobacteria | Enterobacterales | Enterobacteriaceae | Cronobacter | Cronobacter malonaticus |
| Bacteria | Proteobacteria | Gammaproteobacteria | Enterobacterales | Enterobacteriaceae | Enterobacter | Enterobacter asburiae |
| Bacteria | Proteobacteria | Gammaproteobacteria | Enterobacterales | Enterobacteriaceae | Enterobacter | Enterobacter cloacae |
| Bacteria | Proteobacteria | Gammaproteobacteria | Enterobacterales | Enterobacteriaceae | Enterobacter | Enterobacter cloacae complex sp. |
| Bacteria | Proteobacteria | Gammaproteobacteria | Enterobacterales | Enterobacteriaceae | Enterobacter | Enterobacter sichuanensis |
| Bacteria | Proteobacteria | Gammaproteobacteria | Enterobacterales | Enterobacteriaceae | Escherichia | Escherichia albertii |

| | | | | | | |
|---|---|---|---|---|---|---|
| Bacteria | Proteobacteria | Gammaproteobacteria | Enterobacterales | Enterobacteriaceae | Franconibacter | Franconibacter sp. |
| Bacteria | Proteobacteria | Gammaproteobacteria | Enterobacterales | Enterobacteriaceae | Klebsiella | Klebsiella sp. |
| Bacteria | Proteobacteria | Gammaproteobacteria | Enterobacterales | Enterobacteriaceae | Klebsiella | Klebsiella variicola |
| Bacteria | Proteobacteria | Gammaproteobacteria | Enterobacterales | Enterobacteriaceae | Kluyvera | Kluyvera cryocrescens |
| Bacteria | Proteobacteria | Gammaproteobacteria | Enterobacterales | Enterobacteriaceae | Kosakonia | Kosakonia cowanii |
| Bacteria | Proteobacteria | Gammaproteobacteria | Enterobacterales | Enterobacteriaceae | Leclercia | Leclercia sp. Colony189 |
| Bacteria | Proteobacteria | Gammaproteobacteria | Enterobacterales | Enterobacteriaceae | Raoultella | Raoultella sp. |
| Bacteria | Proteobacteria | Gammaproteobacteria | Enterobacterales | Enterobacteriaceae | Salmonella | Salmonella bongori |
| Bacteria | Proteobacteria | Gammaproteobacteria | Enterobacterales | Enterobacteriaceae | Salmonella | Salmonella sp. |
| Bacteria | Proteobacteria | Gammaproteobacteria | Enterobacterales | Enterobacteriaceae | Shigella | Shigella boydii |
| Bacteria | Proteobacteria | Gammaproteobacteria | Enterobacterales | Enterobacteriaceae | Shigella | Shigella dysenteriae |
| Bacteria | Proteobacteria | Gammaproteobacteria | Enterobacterales | Enterobacteriaceae | Shigella | Shigella sp. |
| Bacteria | Proteobacteria | Gammaproteobacteria | Enterobacterales | Enterobacteriaceae | Siccibacter | Siccibacter colletis |
| Bacteria | Proteobacteria | Gammaproteobacteria | Enterobacterales | Enterobacteriaceae | Siccibacter | Siccibacter turicensis |
| Bacteria | Proteobacteria | Gammaproteobacteria | Enterobacterales | Enterobacteriaceae | | endosymbiont of Columbicola clayae |
| Bacteria | Proteobacteria | Gammaproteobacteria | Enterobacterales | Enterobacteriaceae | | Enterobacteriaceae bacterium |
| Bacteria | Proteobacteria | Gammaproteobacteria | Enterobacterales | Enterobacteriaceae | | secondary endosymbiont of Heteropsylla texana |
| Bacteria | Proteobacteria | Gammaproteobacteria | Enterobacterales | Erwiniaceae | Erwinia | Erwinia gerundensis |
| Bacteria | Proteobacteria | Gammaproteobacteria | Enterobacterales | Erwiniaceae | Erwinia | Erwinia mediterraneensis |
| Bacteria | Proteobacteria | Gammaproteobacteria | Enterobacterales | Erwiniaceae | Erwinia | Erwinia rhapontici |
| Bacteria | Proteobacteria | Gammaproteobacteria | Enterobacterales | Erwiniaceae | Erwinia | Erwinia sp. J780 |
| Bacteria | Proteobacteria | Gammaproteobacteria | Enterobacterales | Erwiniaceae | Erwinia | Erwinia sp. LPPA 963 |
| Bacteria | Proteobacteria | Gammaproteobacteria | Enterobacterales | Erwiniaceae | Erwinia | Erwinia sp. QL-Z3 |
| Bacteria | Proteobacteria | Gammaproteobacteria | Enterobacterales | Erwiniaceae | Erwinia | Erwinia uzenensis |
| Bacteria | Proteobacteria | Gammaproteobacteria | Enterobacterales | Erwiniaceae | Erwinia | uncultured Erwinia sp. |
| Bacteria | Proteobacteria | Gammaproteobacteria | Enterobacterales | Erwiniaceae | Izhakiella | Izhakiella capsodis |
| Bacteria | Proteobacteria | Gammaproteobacteria | Enterobacterales | Erwiniaceae | Mixta | Mixta gaviniae |
| Bacteria | Proteobacteria | Gammaproteobacteria | Enterobacterales | Erwiniaceae | Pantoea | Pantoea alhagi |
| Bacteria | Proteobacteria | Gammaproteobacteria | Enterobacterales | Erwiniaceae | Pantoea | Pantoea allii |
| Bacteria | Proteobacteria | Gammaproteobacteria | Enterobacterales | Erwiniaceae | Pantoea | Pantoea sp. Sc1 |
| Bacteria | Proteobacteria | Gammaproteobacteria | Enterobacterales | Erwiniaceae | Pantoea | Pantoea vagans |
| Bacteria | Proteobacteria | Gammaproteobacteria | Enterobacterales | Erwiniaceae | Pantoea | Pantoea wallisii |
| Bacteria | Proteobacteria | Gammaproteobacteria | Enterobacterales | Erwiniaceae | Rosenbergiella | Rosenbergiella nectarea |
| Bacteria | Proteobacteria | Gammaproteobacteria | Enterobacterales | Erwiniaceae | Tatumella | Tatumella punctata |

| | | | | | | |
|---|---|---|---|---|---|---|
| Bacteria | Proteobacteria | Gammaproteobacteria | Enterobacterales | Hafniaceae | Edwardsiella | Edwardsiella sp. EA181011 |
| Bacteria | Proteobacteria | Gammaproteobacteria | Enterobacterales | Hafniaceae | Edwardsiella | Edwardsiella tarda |
| Bacteria | Proteobacteria | Gammaproteobacteria | Enterobacterales | Hafniaceae | Hafnia | Hafnia alvei |
| Bacteria | Proteobacteria | Gammaproteobacteria | Enterobacterales | Morganellaceae | Arsenophonus | Arsenophonus endosymbiont of Aleyrodes elevatus |
| Bacteria | Proteobacteria | Gammaproteobacteria | Enterobacterales | Morganellaceae | Arsenophonus | Arsenophonus endosymbiont of Trialeurodes vaporariorum |
| Bacteria | Proteobacteria | Gammaproteobacteria | Enterobacterales | Morganellaceae | Moellerella | Moellerella wisconsensis |
| Bacteria | Proteobacteria | Gammaproteobacteria | Enterobacterales | Morganellaceae | Proteus | Proteus alimentorum |
| Bacteria | Proteobacteria | Gammaproteobacteria | Enterobacterales | Morganellaceae | Providencia | Providencia rettgeri |
| Bacteria | Proteobacteria | Gammaproteobacteria | Enterobacterales | Morganellaceae | Providencia | Providencia sp. KN12 |
| Bacteria | Proteobacteria | Gammaproteobacteria | Enterobacterales | Morganellaceae | Providencia | Providencia stuartii |
| Bacteria | Proteobacteria | Gammaproteobacteria | Enterobacterales | Morganellaceae | Xenorhabdus | Xenorhabdus bovienii |
| Bacteria | Proteobacteria | Gammaproteobacteria | Enterobacterales | Pectobacteriaceae | Pectobacterium | Pectobacterium sp. CW5 |
| Bacteria | Proteobacteria | Gammaproteobacteria | Enterobacterales | Yersiniaceae | Gibbsiella | Gibbsiella quercinecans |
| Bacteria | Proteobacteria | Gammaproteobacteria | Enterobacterales | Yersiniaceae | Rahnella | Rahnella aquatilis |
| Bacteria | Proteobacteria | Gammaproteobacteria | Enterobacterales | Yersiniaceae | Rouxiella | Rouxiella badensis |
| Bacteria | Proteobacteria | Gammaproteobacteria | Enterobacterales | Yersiniaceae | Serratia | Serratia proteamaculans |
| Bacteria | Proteobacteria | Gammaproteobacteria | Enterobacterales | Yersiniaceae | Serratia | Serratia sp. |
| Bacteria | Proteobacteria | Gammaproteobacteria | Enterobacterales | Yersiniaceae | Serratia | Serratia ureilytica |
| Bacteria | Proteobacteria | Gammaproteobacteria | Enterobacterales | Yersiniaceae | Yersinia | Yersinia aldovae |
| Bacteria | Proteobacteria | Gammaproteobacteria | Enterobacterales | Yersiniaceae | Yersinia | Yersinia pestis |
| Bacteria | Proteobacteria | Gammaproteobacteria | Enterobacterales | Yersiniaceae | Yersinia | Yersinia ruckeri |
| Bacteria | Proteobacteria | Gammaproteobacteria | Enterobacterales | | | uncultured Enterobacterales bacterium |
| Bacteria | Proteobacteria | Gammaproteobacteria | Legionellales | Coxiellaceae | Rickettsiella | Candidatus Rickettsiella viridis |
| Bacteria | Proteobacteria | Gammaproteobacteria | Legionellales | Legionellaceae | Legionella | Legionella spiritensis |
| Bacteria | Proteobacteria | Gammaproteobacteria | Methylococcales | Methylococcaceae | Candidatus Methylospira | Candidatus Methylospira mobilis |
| Bacteria | Proteobacteria | Gammaproteobacteria | Methylococcales | Methylothermaceae | | Methylomonas clara |
| Bacteria | Proteobacteria | Gammaproteobacteria | Nevskiales | Sinobacteraceae | Nevskia | Nevskia sp. |
| Bacteria | Proteobacteria | Gammaproteobacteria | Nevskiales | Sinobacteraceae | Sinimarinibacterium | Sinimarinibacterium sp. NLF-5-8 |
| Bacteria | Proteobacteria | Gammaproteobacteria | Nevskiales | Sinobacteraceae | Solimonas | Solimonas sp. K1W22B-7 |
| Bacteria | Proteobacteria | Gammaproteobacteria | Oceanospirillales | Alcanivoracaceae | Alcanivorax | Alcanivorax sp. N3-2A |
| Bacteria | Proteobacteria | Gammaproteobacteria | Oceanospirillales | Halomonadaceae | Carnimonas | Carnimonas nigrificans |
| Bacteria | Proteobacteria | Gammaproteobacteria | Oceanospirillales | Halomonadaceae | Halomonas | Halomonas sp. BC-M4-5 |

| | | | | | | |
|---|---|---|---|---|---|---|
| Bacteria | Proteobacteria | Gammaproteobacteria | Oceanospirillales | Halomonadaceae | Zymobacter | Zymobacter palmae |
| Bacteria | Proteobacteria | Gammaproteobacteria | Oceanospirillales | Pleioneaceae | Aliikangiella | Aliikangiella coralliicola |
| Bacteria | Proteobacteria | Gammaproteobacteria | Orbales | Orbaceae | Frischella | Frischella japonica |
| Bacteria | Proteobacteria | Gammaproteobacteria | Orbales | Orbaceae | Frischella | Frischella perrara |
| Bacteria | Proteobacteria | Gammaproteobacteria | Pasteurellales | Pasteurellaceae | Actinobacillus | Actinobacillus indolicus |
| Bacteria | Proteobacteria | Gammaproteobacteria | Pasteurellales | Pasteurellaceae | Aggregatibacter | Aggregatibacter actinomycetemcomitans |
| Bacteria | Proteobacteria | Gammaproteobacteria | Pasteurellales | Pasteurellaceae | Avibacterium | Avibacterium volantium |
| Bacteria | Proteobacteria | Gammaproteobacteria | Pasteurellales | Pasteurellaceae | Basfia | [Mannheimia] succiniciproducens |
| Bacteria | Proteobacteria | Gammaproteobacteria | Pasteurellales | Pasteurellaceae | Haemophilus | Haemophilus influenzae |
| Bacteria | Proteobacteria | Gammaproteobacteria | Pasteurellales | Pasteurellaceae | Haemophilus | Haemophilus parainfluenzae |
| Bacteria | Proteobacteria | Gammaproteobacteria | Pasteurellales | Pasteurellaceae | Haemophilus | Haemophilus quentini |
| Bacteria | Proteobacteria | Gammaproteobacteria | Pasteurellales | Pasteurellaceae | Haemophilus | Haemophilus sp. 'paraurethrae' |
| Bacteria | Proteobacteria | Gammaproteobacteria | Pasteurellales | Pasteurellaceae | Pasteurella | Pasteurella skyensis |
| Bacteria | Proteobacteria | Gammaproteobacteria | Pseudomonadales | Moraxellaceae | Acinetobacter | Acinetobacter courvalinii |
| Bacteria | Proteobacteria | Gammaproteobacteria | Pseudomonadales | Moraxellaceae | Acinetobacter | Acinetobacter rhizosphaerae |
| Bacteria | Proteobacteria | Gammaproteobacteria | Pseudomonadales | Moraxellaceae | Acinetobacter | Acinetobacter schindleri |
| Bacteria | Proteobacteria | Gammaproteobacteria | Pseudomonadales | Moraxellaceae | Acinetobacter | Acinetobacter sp. asd01 |
| Bacteria | Proteobacteria | Gammaproteobacteria | Pseudomonadales | Moraxellaceae | Acinetobacter | Acinetobacter sp. Lhl-4r |
| Bacteria | Proteobacteria | Gammaproteobacteria | Pseudomonadales | Moraxellaceae | Acinetobacter | Acinetobacter sp. Marseille-Q1620 |
| Bacteria | Proteobacteria | Gammaproteobacteria | Pseudomonadales | Moraxellaceae | Moraxella | Moraxella sp. |
| Bacteria | Proteobacteria | Gammaproteobacteria | Pseudomonadales | Moraxellaceae | Psychrobacter | uncultured Psychrobacter sp. |
| Bacteria | Proteobacteria | Gammaproteobacteria | Pseudomonadales | Pseudomonadaceae | Entomomonas | Entomomonas moraniae |
| Bacteria | Proteobacteria | Gammaproteobacteria | Pseudomonadales | Pseudomonadaceae | Pseudomonas | Pseudomonas balearica |
| Bacteria | Proteobacteria | Gammaproteobacteria | Pseudomonadales | Pseudomonadaceae | Pseudomonas | Pseudomonas fuscovaginae |
| Bacteria | Proteobacteria | Gammaproteobacteria | Pseudomonadales | Pseudomonadaceae | Pseudomonas | Pseudomonas hutmensis |
| Bacteria | Proteobacteria | Gammaproteobacteria | Pseudomonadales | Pseudomonadaceae | Pseudomonas | Pseudomonas lutea |
| Bacteria | Proteobacteria | Gammaproteobacteria | Pseudomonadales | Pseudomonadaceae | Pseudomonas | Pseudomonas monteilii |
| Bacteria | Proteobacteria | Gammaproteobacteria | Pseudomonadales | Pseudomonadaceae | Pseudomonas | Pseudomonas nitritireducens |
| Bacteria | Proteobacteria | Gammaproteobacteria | Pseudomonadales | Pseudomonadaceae | Pseudomonas | Pseudomonas nitroreducens |
| Bacteria | Proteobacteria | Gammaproteobacteria | Pseudomonadales | Pseudomonadaceae | Pseudomonas | Pseudomonas paralactis |
| Bacteria | Proteobacteria | Gammaproteobacteria | Pseudomonadales | Pseudomonadaceae | Pseudomonas | Pseudomonas pelagia |
| Bacteria | Proteobacteria | Gammaproteobacteria | Pseudomonadales | Pseudomonadaceae | Pseudomonas | Pseudomonas putida |
| Bacteria | Proteobacteria | Gammaproteobacteria | Pseudomonadales | Pseudomonadaceae | Pseudomonas | Pseudomonas rhizosphaerae |
| Bacteria | Proteobacteria | Gammaproteobacteria | Pseudomonadales | Pseudomonadaceae | Pseudomonas | Pseudomonas sp. AA-T01 |

| | | | | | | |
|---|---|---|---|---|---|---|
| Bacteria | Proteobacteria | Gammaproteobacteria | Pseudomonadales | Pseudomonadaceae | Pseudomonas | Pseudomonas sp. B3100 |
| Bacteria | Proteobacteria | Gammaproteobacteria | Pseudomonadales | Pseudomonadaceae | Pseudomonas | Pseudomonas sp. FB6 |
| Bacteria | Proteobacteria | Gammaproteobacteria | Pseudomonadales | Pseudomonadaceae | Pseudomonas | Pseudomonas sp. LAB-08 |
| Bacteria | Proteobacteria | Gammaproteobacteria | Pseudomonadales | Pseudomonadaceae | Pseudomonas | Pseudomonas sp. LAMKW06 |
| Bacteria | Proteobacteria | Gammaproteobacteria | Pseudomonadales | Pseudomonadaceae | Pseudomonas | Pseudomonas sp. OIL-1 |
| Bacteria | Proteobacteria | Gammaproteobacteria | Pseudomonadales | Pseudomonadaceae | Pseudomonas | Pseudomonas sp. SSCT60 |
| Bacteria | Proteobacteria | Gammaproteobacteria | Pseudomonadales | Pseudomonadaceae | Pseudomonas | Pseudomonas sp. VT1B |
| Bacteria | Proteobacteria | Gammaproteobacteria | Pseudomonadales | Pseudomonadaceae | Pseudomonas | Pseudomonas stutzeri |
| Bacteria | Proteobacteria | Gammaproteobacteria | Pseudomonadales | Pseudomonadaceae | Pseudomonas | Pseudomonas syringae group genomosp. 3 |
| Bacteria | Proteobacteria | Gammaproteobacteria | Pseudomonadales | Pseudomonadaceae | Pseudomonas | Pseudomonas taiwanensis |
| Bacteria | Proteobacteria | Gammaproteobacteria | Pseudomonadales | Pseudomonadaceae | Pseudomonas | Pseudomonas tolaasii |
| Bacteria | Proteobacteria | Gammaproteobacteria | Pseudomonadales | Pseudomonadaceae | Pseudomonas | Pseudomonas viridiflava |
| Bacteria | Proteobacteria | Gammaproteobacteria | Thiotrichales | Thiotrichaceae | Achromatium | Achromatium minus |
| Bacteria | Proteobacteria | Gammaproteobacteria | Thiotrichales | Thiotrichaceae | Achromatium | Achromatium oxaliferum |
| Bacteria | Proteobacteria | Gammaproteobacteria | Vibrionales | Vibrionaceae | Photobacterium | Photobacterium sp. |
| Bacteria | Proteobacteria | Gammaproteobacteria | Vibrionales | Vibrionaceae | Vibrio | uncultured Vibrio sp. |
| Bacteria | Proteobacteria | Gammaproteobacteria | Vibrionales | Vibrionaceae | Vibrio | Vibrio astriarenae |
| Bacteria | Proteobacteria | Gammaproteobacteria | Vibrionales | Vibrionaceae | Vibrio | Vibrio parahaemolyticus |
| Bacteria | Proteobacteria | Gammaproteobacteria | Xanthomonadales | Rhodanobacteraceae | Dokdonella | Dokdonella koreensis |
| Bacteria | Proteobacteria | Gammaproteobacteria | Xanthomonadales | Rhodanobacteraceae | Dyella | Dyella caseinilytica |
| Bacteria | Proteobacteria | Gammaproteobacteria | Xanthomonadales | Rhodanobacteraceae | Luteibacter | Luteibacter rhizovicinus |
| Bacteria | Proteobacteria | Gammaproteobacteria | Xanthomonadales | Rhodanobacteraceae | Luteibacter | Luteibacter sp. |
| Bacteria | Proteobacteria | Gammaproteobacteria | Xanthomonadales | Xanthomonadaceae | Luteimonas | Luteimonas granuli |
| Bacteria | Proteobacteria | Gammaproteobacteria | Xanthomonadales | Xanthomonadaceae | Luteimonas | Luteimonas sp. LNNU24178 |
| Bacteria | Proteobacteria | Gammaproteobacteria | Xanthomonadales | Xanthomonadaceae | Luteimonas | Luteimonas sp. XBU10 |
| Bacteria | Proteobacteria | Gammaproteobacteria | Xanthomonadales | Xanthomonadaceae | Lysobacter | Lysobacter lycopersici |
| Bacteria | Proteobacteria | Gammaproteobacteria | Xanthomonadales | Xanthomonadaceae | Lysobacter | Lysobacter sp. |
| Bacteria | Proteobacteria | Gammaproteobacteria | Xanthomonadales | Xanthomonadaceae | Pseudoxanthomonas | Pseudoxanthomonas suwonensis |
| Bacteria | Proteobacteria | Gammaproteobacteria | Xanthomonadales | Xanthomonadaceae | Stenotrophomonas | Stenotrophomonas acidaminiphila |
| Bacteria | Proteobacteria | Gammaproteobacteria | Xanthomonadales | Xanthomonadaceae | Stenotrophomonas | Stenotrophomonas chelatiphaga |
| Bacteria | Proteobacteria | Gammaproteobacteria | Xanthomonadales | Xanthomonadaceae | Thermomonas | uncultured Thermomonas sp. |
| Bacteria | Proteobacteria | Gammaproteobacteria | Xanthomonadales | Xanthomonadaceae | Xanthomonas | Xanthomonas citri |
| Bacteria | Proteobacteria | Gammaproteobacteria | Xanthomonadales | Xanthomonadaceae | Xanthomonas | Xanthomonas phaseoli |
| Bacteria | Proteobacteria | Gammaproteobacteria | Xanthomonadales | Xanthomonadaceae | Xylella | uncultured Xylella sp. |

| | | | | | | |
|---|---|---|---|---|---|---|
| Bacteria | Proteobacteria | Gammaproteobacteria | Xanthomonadales | Xanthomonadaceae | | uncultured Xanthomonadaceae bacterium |
| Bacteria | Proteobacteria | Gammaproteobacteria | | | Candidatus Endonucleariobacter | Candidatus Endonucleariobacter sp. (ex Euplotes eurystomus) |
| Bacteria | Proteobacteria | Gammaproteobacteria | | | Gallaecimonas | Gallaecimonas sp. MA-8 |
| Bacteria | Proteobacteria | Gammaproteobacteria | | | | gamma proteobacterium PEB0183 |
| Bacteria | Proteobacteria | Gammaproteobacteria | | | | Gammaproteobacteria bacterium |
| Bacteria | Proteobacteria | Oligoflexia | Bdellovibrionales | Bdellovibrionaceae | Bdellovibrio | Bdellovibrio sp. |
| Bacteria | Proteobacteria | Oligoflexia | Bdellovibrionales | Bdellovibrionaceae | Bdellovibrio | Bdellovibrio sp. ZAP7 |
| Bacteria | Proteobacteria | Oligoflexia | Bdellovibrionales | Bdellovibrionaceae | Bdellovibrio | uncultured Bdellovibrio sp. |
| Bacteria | Proteobacteria | Oligoflexia | Silvanigrellales | Silvanigrellaceae | Silvanigrella | Silvanigrella sp. |
| Bacteria | Spirochaetes | Spirochaetia | Spirochaetales | Spirochaetaceae | Treponema | Treponema sp. Marseille-Q4523 |
| Bacteria | Spirochaetes | Spirochaetia | Spirochaetales | Spirochaetaceae | Treponema | Treponema succinifaciens |
| Bacteria | Spirochaetes | Spirochaetia | Spirochaetales | | | uncultured spirochete Kf401 |
| Bacteria | Tenericutes | Mollicutes | Acholeplasmatales | Acholeplasmataceae | Candidatus Phytoplasma | 'Sophora alopecuroides' yellows phytoplasma |
| Bacteria | Tenericutes | Mollicutes | Entomoplasmatales | Entomoplasmataceae | Mesoplasma | Mesoplasma coleopterae |
| Bacteria | Tenericutes | Mollicutes | Entomoplasmatales | Entomoplasmataceae | Mesoplasma | Mesoplasma florum |
| Bacteria | Tenericutes | Mollicutes | Entomoplasmatales | Spiroplasmataceae | Spiroplasma | Spiroplasma sp. Ar-1357 |
| Bacteria | Tenericutes | Mollicutes | Mycoplasmatales | Mycoplasmataceae | Mycoplasma | Candidatus Mycoplasma haemobos |
| Bacteria | Tenericutes | Mollicutes | Mycoplasmatales | Mycoplasmataceae | Mycoplasma | Candidatus Mycoplasma haemocervae |
| Bacteria | Tenericutes | Mollicutes | Mycoplasmatales | Mycoplasmataceae | Mycoplasma | Mycoplasma capricolum |
| Bacteria | Thermotogae | Thermotogae | Mesoaciditogales | Mesoaciditogaceae | Athalassotoga | Athalassotoga saccharophila |
| Bacteria | Verrucomicrobia | Opitutae | Opitutales | Opitutaceae | Lacunisphaera | Lacunisphaera limnophila |
| Bacteria | Verrucomicrobia | Opitutae | Opitutales | Opitutaceae | Opitutus | uncultured Opitutus sp. |
| Bacteria | Verrucomicrobia | Spartobacteria | | | Candidatus Xiphinematobacter | Candidatus Xiphinematobacter sp. Idaho Grape |
| Bacteria | Verrucomicrobia | Verrucomicrobiae | Verrucomicrobiales | Akkermansiaceae | | Akkermansiaceae bacterium |
| Bacteria | Verrucomicrobia | Verrucomicrobiae | Verrucomicrobiales | Verrucomicrobiaceae | Luteolibacter | Luteolibacter luteus |
| Bacteria | | | | | Vermiphilus | Vermiphilus pyriformis |
| Bacteria | | | | | | bacterium 52-L049658-122-015-D07 |
| Bacteria | | | | | | bacterium 87-L049658-122-015-G11 |
| Bacteria | | | | | | bacterium A182 |
| Bacteria | | | | | | bacterium AM0361 |
| Bacteria | | | | | | bacterium AW17 |
| Bacteria | | | | | | bacterium CSMCRI-1085 |
| Bacteria | | | | | | bacterium GD56SAN75% |
| Bacteria | | | | | | bacterium JP74 |

| | | | | | | |
|---|---|---|---|---|---|---|
| Bacteria | | | | | | bacterium KU12 |
| Bacteria | | | | | | bacterium L200B.191 |
| Bacteria | | | | | | bacterium NLAE-zl-C293 |
| Bacteria | | | | | | bacterium NLAE-zl-H137 |
| Bacteria | | | | | | bacterium PII.17 |
| Bacteria | | | | | | bacterium SU5 |
| Bacteria | | | | | | bacterium W132B.89 |
| Bacteria | | | | | | endophytic bacterium |
| Bacteria | | | | | | endosymbiont of Acanthamoeba sp. |
| Bacteria | | | | | | swine manure bacterium RT-21 |
| Bacteria | | | | | | swine manure bacterium RT-6B |
| Bacteria | | | | | | uncultured bacterium 126 |
| Bacteria | | | | | | uncultured bacterium Ak20-3 |
| Bacteria | | | | | | uncultured bacterium ctg7180000001448 |
| Bacteria | | | | | | uncultured bacterium FC3 |
| Bacteria | | | | | | uncultured compost bacterium |
| Bacteria | | | | | | uncultured endophytic bacterium |
| Bacteria | | | | | | uncultured freshwater bacterium |
| Bacteria | | | | | | uncultured rumen bacterium 4C0d-5 |
| Bacteria | | | | | | uncultured SB1 group bacterium |
| Bacteria | | | | | | uncultured sediment bacterium |
| Bacteria | | | | | | unidentified rumen bacterium RC28 |
| Eukaryota | Streptophyta | Magnoliopsida | Poales | Poaceae | Zea | Zea mays |
| Eukaryota | Ascomycota | Eurotiomycetes | Onygenales | Ascosphaeraceae | Ascosphaera | Ascosphaera aggregata |
| Eukaryota | Streptophyta | Magnoliopsida | Fabales | Fabaceae | Trigonella | Trigonella foenum-graecum |
| Eukaryota | | | | | | uncultured phototrophic eukaryote |
| Eukaryota | Streptophyta | Magnoliopsida | Fabales | Fabaceae | Medicago | Medicago arborea |
| Eukaryota | Streptophyta | Magnoliopsida | Asterales | Asteraceae | Praxelis | Praxelis clematidea |
| Eukaryota | Streptophyta | Magnoliopsida | Asterales | Asteraceae | Erigeron | Erigeron canadensis |
| Eukaryota | | | | | | uncultured eukaryote |
| Eukaryota | Ascomycota | Eurotiomycetes | Onygenales | Ascosphaeraceae | Ascosphaera | Ascosphaera asterophora |
| Eukaryota | Streptophyta | Magnoliopsida | Fabales | Fabaceae | Medicago | Medicago truncatula |
| Eukaryota | Streptophyta | Magnoliopsida | Asparagales | Asparagaceae | Asparagus | Asparagus filicinus |
| Eukaryota | Streptophyta | Magnoliopsida | Fabales | Fabaceae | Medicago | Medicago sativa |
| Eukaryota | Arthropoda | Insecta | Hemiptera | Coccidae | Ceroplastes | Ceroplastes rusci |
| Eukaryota | Streptophyta | Magnoliopsida | Liliales | Liliaceae | Lilium | Lilium washingtonianum |
| Eukaryota | Ascomycota | Eurotiomycetes | Onygenales | Ascosphaeraceae | Ascosphaera | Ascosphaera proliperda |
| Eukaryota | Streptophyta | Magnoliopsida | Rosales | Rhamnaceae | Rhamnus | Rhamnus taquetii |

| | | | | | | |
|---|---|---|---|---|---|---|
| Eukaryota | Ascomycota | Eurotiomycetes | Onygenales | Ascosphaeraceae | Ascosphaera | Ascosphaera atra |
| Eukaryota | Streptophyta | Magnoliopsida | Cucurbitales | Cucurbitaceae | Citrullus | Citrullus mucosospermus |
| Eukaryota | Streptophyta | Magnoliopsida | Poales | Poaceae | Setaria | Setaria viridis |
| Eukaryota | Ascomycota | Eurotiomycetes | Onygenales | Ascosphaeraceae | Ascosphaera | Ascosphaera flava |
| Eukaryota | Streptophyta | Magnoliopsida | Brassicales | Brassicaceae | Brassica | Brassica rapa |
| Eukaryota | Streptophyta | Magnoliopsida | Dipsacales | Adoxaceae | Sambucus | Sambucus nigra |
| Eukaryota | Arthropoda | Insecta | Hymenoptera | Eulophidae | Aprostocetus | Aprostocetus foraminifer |
| Eukaryota | Arthropoda | Insecta | Hymenoptera | Megachilidae | Megachile | Megachile giraudi |
| Eukaryota | Streptophyta | Polypodiopsida | Cyatheales | Plagiogyriaceae | Plagiogyria | Plagiogyria euphlebia |
| Eukaryota | Streptophyta | Magnoliopsida | Fabales | Fabaceae | Cicer | Cicer arietinum |
| Eukaryota | Streptophyta | Magnoliopsida | Fabales | Fabaceae | Medicago | Medicago pironae |
| Eukaryota | Streptophyta | Magnoliopsida | Sapindales | Sapindaceae | Aesculus | Aesculus chinensis |
| Eukaryota | Streptophyta | Magnoliopsida | Caryophyllales | Chenopodiaceae | Chenopodium | Chenopodium quinoa |
| Eukaryota | Streptophyta | Magnoliopsida | Solanales | Convolvulaceae | Ipomoea | Ipomoea triloba |
| Eukaryota | Streptophyta | Magnoliopsida | Apiales | Apiaceae | Ligusticum | Ligusticum hispidum |
| Eukaryota | Streptophyta | Magnoliopsida | Ranunculales | Papaveraceae | Meconopsis | Meconopsis integrifolia |
| Eukaryota | Streptophyta | Magnoliopsida | Lamiales | Oleaceae | Ligustrum | Ligustrum quihoui |
| Eukaryota | | | | | | uncultured fungus |
| Eukaryota | Ascomycota | Eurotiomycetes | Onygenales | Ascosphaeraceae | Ascosphaera | Ascosphaera solina |
| Eukaryota | Arthropoda | Insecta | Hymenoptera | Apidae | Bombus | Bombus pascuorum |
| Eukaryota | Arthropoda | Insecta | Hymenoptera | Eulophidae | Aprostocetus | Aprostocetus caudatus |
| Eukaryota | Ascomycota | Eurotiomycetes | Eurotiales | Aspergillaceae | Aspergillus | Aspergillus chevalieri |
| Eukaryota | Ascomycota | Eurotiomycetes | Onygenales | Ascosphaeraceae | Ascosphaera | Ascosphaera naganensis |
| Eukaryota | Streptophyta | Magnoliopsida | Brassicales | Brassicaceae | Brassica | Brassica oleracea |
| Eukaryota | Streptophyta | Magnoliopsida | Rosales | Cannabaceae | Humulus | Humulus lupulus |
| Eukaryota | Streptophyta | | | | | uncultured Streptophyta |
| Eukaryota | Ascomycota | Eurotiomycetes | Onygenales | Ascosphaeraceae | Ascosphaera | Ascosphaera acerosa |
| Eukaryota | Streptophyta | Magnoliopsida | Solanales | Solanaceae | Solanum | Solanum aethiopicum |
| Eukaryota | Mucoromycota | Glomeromycetes | Glomerales | Glomeraceae | Glomus | Glomus sp. |
| Eukaryota | Streptophyta | Magnoliopsida | Brassicales | Brassicaceae | Arabidopsis | Arabidopsis thaliana |
| Eukaryota | Streptophyta | Magnoliopsida | Brassicales | Brassicaceae | Raphanus | Raphanus sativus |
| Eukaryota | Ascomycota | Eurotiomycetes | Onygenales | Ascosphaeraceae | Ascosphaera | Ascosphaera subglobosa |
| Eukaryota | Streptophyta | Magnoliopsida | Lamiales | Verbenaceae | Lippia | Lippia origanoides |
| Eukaryota | Streptophyta | Magnoliopsida | Poales | Poaceae | Digitaria | Digitaria exilis |
| Eukaryota | Streptophyta | Magnoliopsida | Solanales | Solanaceae | Solanum | Solanum pinnatisectum |
| Eukaryota | Streptophyta | Magnoliopsida | Solanales | Solanaceae | Solanum | Solanum tuberosum |
| Eukaryota | Streptophyta | Magnoliopsida | Zygophyllales | Zygophyllaceae | Zygophyllum | Zygophyllum fabago |
| Eukaryota | Arthropoda | Insecta | Hymenoptera | Megachilidae | Megachile | Megachile nitidicollis |
| Eukaryota | Streptophyta | Magnoliopsida | Asterales | Asteraceae | Paraprenanthes | Paraprenanthes diversifolia |
| Eukaryota | Streptophyta | Magnoliopsida | Brassicales | Brassicaceae | Brassica | Brassica napus |
| Eukaryota | Streptophyta | Magnoliopsida | Fabales | Fabaceae | Medicago | Medicago monspeliaca |

| | | | | | | |
|---|---|---|---|---|---|---|
| Eukaryota | Streptophyta | Magnoliopsida | Fabales | Fabaceae | Vigna | Vigna angularis |
| Eukaryota | Arthropoda | Insecta | Hymenoptera | Megachilidae | Megachile | Megachile (Eutricharaea) malangensis group sp. n. 30 |
| Eukaryota | Chordata | Mammalia | Rodentia | Muridae | Mus | Mus musculus |
| Eukaryota | Streptophyta | Magnoliopsida | Fagales | Betulaceae | Betula | Betula pendula |
| Eukaryota | Streptophyta | Magnoliopsida | Lamiales | Acanthaceae | Rungia | Rungia pectinata |
| Eukaryota | Streptophyta | Magnoliopsida | Poales | Poaceae | Lolium | Lolium perenne |
| Eukaryota | Streptophyta | Magnoliopsida | Ranunculales | Berberidaceae | Epimedium | Epimedium brevicornu |
| Eukaryota | Streptophyta | Magnoliopsida | Solanales | Solanaceae | Solanum | Solanum lycopersicum |
| Eukaryota | Streptophyta | Magnoliopsida | Fabales | Fabaceae | Melilotus | Melilotus albus |
| Eukaryota | Arthropoda | Insecta | Coleoptera | Cerylonidae | Mychocerus | Mychocerus sp. Panama |
| Eukaryota | Ascomycota | Eurotiomycetes | Onygenales | Ascosphaeraceae | Ascosphaera | Ascosphaera duoformis |
| Eukaryota | Streptophyta | Bryopsida | Dicranales | Fissidentaceae | Fissidens | Fissidens nobilis |
| Eukaryota | Streptophyta | Cycadopsida | Cycadales | Cycadaceae | Cycas | Cycas szechuanensis |
| Eukaryota | Streptophyta | Gnetopsida | Ephedrales | Ephedraceae | Ephedra | Ephedra saxatilis |
| Eukaryota | Streptophyta | Magnoliopsida | Apiales | Apiaceae | Chamaesium | Chamaesium thalictrifolium |
| Eukaryota | Streptophyta | Magnoliopsida | Caryophyllales | Aizoaceae | Sesuvium | Sesuvium portulacastrum |
| Eukaryota | Streptophyta | Magnoliopsida | Lamiales | Scrophulariaceae | Scrophularia | Scrophularia takesimensis |
| Eukaryota | Streptophyta | Magnoliopsida | Nymphaeales | Nymphaeaceae | Nymphaea | Nymphaea colorata |
| Eukaryota | Streptophyta | Magnoliopsida | Rosales | Moraceae | Artocarpus | Artocarpus altilis |
| Eukaryota | Streptophyta | Magnoliopsida | Rosales | Rosaceae | Prunus | Prunus dulcis |
| Eukaryota | Streptophyta | Magnoliopsida | Solanales | Convolvulaceae | Cuscuta | Cuscuta approximata |
| Eukaryota | Streptophyta | Magnoliopsida | | | | Magnoliophyta environmental sample |
| Eukaryota | Arthropoda | Insecta | Hymenoptera | Apidae | Bombus | Bombus hortorum |
| Eukaryota | Arthropoda | Insecta | Hymenoptera | Eulophidae | Melittobia | Melittobia cf. hawaiiensis SS-2018 |
| Eukaryota | Mucoromycota | | | | | uncultured Glomeromycotina |
| Eukaryota | Streptophyta | Magnoliopsida | Apiales | Araliaceae | Panax | Panax trifolius |
| Eukaryota | Streptophyta | Magnoliopsida | Asterales | Asteraceae | Marshallia | Marshallia ramosa |
| Eukaryota | Streptophyta | Magnoliopsida | Cucurbitales | Cucurbitaceae | Cucumis | Cucumis melo |
| Eukaryota | Streptophyta | Magnoliopsida | Cucurbitales | Cucurbitaceae | Cucumis | Cucumis sativus |
| Eukaryota | Streptophyta | Magnoliopsida | Fabales | Fabaceae | Vigna | Vigna unguiculata |
| Eukaryota | Streptophyta | Magnoliopsida | Malvales | Muntingiaceae | Muntingia | Muntingia calabura |
| Eukaryota | Streptophyta | Magnoliopsida | Poales | Bromeliaceae | Ananas | Ananas comosus |
| Eukaryota | Arthropoda | Insecta | Hymenoptera | Apidae | Bombus | Bombus campestris |
| Eukaryota | Arthropoda | Insecta | Hymenoptera | Megachilidae | Megachile | Megachile sp. Pseudocentron-group 209 |
| Eukaryota | Arthropoda | Insecta | Hymenoptera | Pteromalidae | Otitesella | Otitesella uluzi |
| Eukaryota | Ascomycota | Dothideomycetes | Pleosporales | Pleosporaceae | Stemphylium | Stemphylium vesicarium |
| Eukaryota | Ascomycota | Eurotiomycetes | Eurotiales | Trichocomaceae | Talaromyces | Talaromyces rugulosus |
| Eukaryota | Ascomycota | Sordariomycetes | Magnaporthales | Pyriculariaceae | Pyricularia | Pyricularia oryzae |
| Eukaryota | Chlorophyta | Ulvophyceae | Ulvales | Kornmanniaceae | Lithotrichon | Lithotrichon fluminensis |

| | | | | | | |
|---|---|---|---|---|---|---|
| Eukaryota | Mucoromycota | Glomeromycetes | Glomerales | Glomeraceae | | uncultured Glomus |
| Eukaryota | Streptophyta | Magnoliopsida | Apiales | Araliaceae | Panax | Panax sp. 'sinensis' |
| Eukaryota | Streptophyta | Magnoliopsida | Asparagales | Asphodelaceae | Eremurus | Eremurus robustus |
| Eukaryota | Streptophyta | Magnoliopsida | Asparagales | Orchidaceae | Corallorhiza | Corallorhiza mertensiana |
| Eukaryota | Streptophyta | Magnoliopsida | Asparagales | Orchidaceae | Hexalectris | Hexalectris arizonica |
| Eukaryota | Streptophyta | Magnoliopsida | Brassicales | Caricaceae | Carica | Carica papaya |
| Eukaryota | Streptophyta | Magnoliopsida | Caryophyllales | Dioncophyllaceae | Dioncophyllum | Dioncophyllum thollonii |
| Eukaryota | Streptophyta | Magnoliopsida | Ericales | Pentaphylacaceae | Anneslea | Anneslea fragrans |
| Eukaryota | Streptophyta | Magnoliopsida | Fabales | Fabaceae | Arachis | Arachis hypogaea |
| Eukaryota | Streptophyta | Magnoliopsida | Fabales | Fabaceae | Castanospermum | Castanospermum australe |
| Eukaryota | Streptophyta | Magnoliopsida | Fabales | Fabaceae | Ladeania | Ladeania lanceolata |
| Eukaryota | Streptophyta | Magnoliopsida | Fabales | Fabaceae | Lotus | Lotus japonicus |
| Eukaryota | Streptophyta | Magnoliopsida | Fabales | Fabaceae | Medicago | Medicago biflora |
| Eukaryota | Streptophyta | Magnoliopsida | Fabales | Fabaceae | Medicago | Medicago minima |
| Eukaryota | Streptophyta | Magnoliopsida | Fabales | Fabaceae | Vigna | Vigna radiata |
| Eukaryota | Streptophyta | Magnoliopsida | Fabales | Fabaceae | | Phaseoleae environmental sample |
| Eukaryota | Streptophyta | Magnoliopsida | Liliales | Liliaceae | Fritillaria | Fritillaria maximowiczii |
| Eukaryota | Streptophyta | Magnoliopsida | Malvales | Malvaceae | Gossypium | Gossypium raimondii |
| Eukaryota | Streptophyta | Magnoliopsida | Malvales | Malvaceae | Gossypium | Gossypium turneri |
| Eukaryota | Streptophyta | Magnoliopsida | Ranunculales | Ranunculaceae | Aquilegia | Aquilegia rockii |
| Eukaryota | Streptophyta | Magnoliopsida | Sapindales | Nitrariaceae | Peganum | Peganum harmala |
| Eukaryota | Streptophyta | Magnoliopsida | Saxifragales | Saxifragaceae | Saniculiphyllum | Saniculiphyllum guangxiense |
| Eukaryota | Streptophyta | Magnoliopsida | Solanales | Convolvulaceae | Convolvulus | Convolvulus arvensis |
| Eukaryota | Streptophyta | Magnoliopsida | Solanales | Convolvulaceae | Ipomoea | Ipomoea trifida |
| Eukaryota | Streptophyta | Magnoliopsida | Zingiberales | Marantaceae | Stromanthe | Stromanthe stromanthoides |
| Eukaryota | Streptophyta | Magnoliopsida | Zygophyllales | Zygophyllaceae | Tetraena | Tetraena mongolica |
| Eukaryota | Arthropoda | Insecta | Coleoptera | Trogossitidae | Temnoscheila | Temnoscheila sp. BT0015 |
| Eukaryota | Arthropoda | Insecta | Hymenoptera | Megachilidae | Megachile | Megachile sculpturalis |
| Eukaryota | Arthropoda | Insecta | Hymenoptera | Scelionidae | Embidobia | Embidobia sp. OSUC 627843 |
| Eukaryota | Arthropoda | Insecta | Hymenoptera | | | Apoidea sp. DAB077 |
| Eukaryota | Arthropoda | Insecta | Hymenoptera | | | Hymenoptera sp. 24089 |
| Eukaryota | Arthropoda | Ostracoda | Myodocopida | Cypridinidae | Heterodesmus | Heterodesmus apriculus |
| Eukaryota | Ascomycota | Eurotiomycetes | Eurotiales | Aspergillaceae | Aspergillus | Aspergillus amstelodami |
| Eukaryota | Ascomycota | Eurotiomycetes | Eurotiales | Aspergillaceae | Aspergillus | Aspergillus sp. |
| Eukaryota | Ascomycota | Eurotiomycetes | Onygenales | Ascosphaeraceae | Ascosphaera | Ascosphaera apis |
| Eukaryota | Chordata | Actinopteri | Gadiformes | Gadidae | Gadus | Gadus morhua |
| Eukaryota | Chordata | Actinopteri | Salmoniformes | Salmonidae | Coregonus | Coregonus sp. 'balchen' |
| Eukaryota | Chordata | Mammalia | Artiodactyla | Suidae | Sus | Sus scrofa |
| Eukaryota | Mucoromycota | Glomeromycetes | Glomerales | Glomeraceae | | uncultured Funneliformis |
| Eukaryota | Mucoromycota | | | | | Mucoromycotina sp. |

| | | | | | | |
|---|---|---|---|---|---|---|
| Eukaryota | Streptophyta | Magnoliopsida | Asterales | Asteraceae | Pseudognaphalium | Pseudognaphalium sp. RS-2020 |
| Eukaryota | Streptophyta | Magnoliopsida | Asterales | Campanulaceae | Codonopsis | Codonopsis lanceolata |
| Eukaryota | Streptophyta | Magnoliopsida | Caryophyllales | Chenopodiaceae | Halimione | Halimione portulacoides |
| Eukaryota | Streptophyta | Magnoliopsida | Cucurbitales | Begoniaceae | Begonia | Begonia guangxiensis |
| Eukaryota | Streptophyta | Magnoliopsida | Cucurbitales | Cucurbitaceae | Cucurbita | Cucurbita pepo |
| Eukaryota | Streptophyta | Magnoliopsida | Dilleniales | Dilleniaceae | Dillenia | Dillenia indica |
| Eukaryota | Streptophyta | Magnoliopsida | Dipsacales | Caprifoliaceae | Abelia | Abelia uniflora |
| Eukaryota | Streptophyta | Magnoliopsida | Ericales | Balsaminaceae | Impatiens | Impatiens guizhouensis |
| Eukaryota | Streptophyta | Magnoliopsida | Ericales | Ebenaceae | Diospyros | Diospyros virginiana |
| Eukaryota | Streptophyta | Magnoliopsida | Ericales | Theaceae | Stewartia | Stewartia sinii |
| Eukaryota | Streptophyta | Magnoliopsida | Fabales | Fabaceae | Glycyrrhiza | Glycyrrhiza inflata |
| Eukaryota | Streptophyta | Magnoliopsida | Fabales | Fabaceae | Lespedeza | Lespedeza bicolor |
| Eukaryota | Streptophyta | Magnoliopsida | Fabales | Fabaceae | Medicago | Medicago arabica |
| Eukaryota | Streptophyta | Magnoliopsida | Fabales | Fabaceae | Medicago | Medicago radiata |
| Eukaryota | Streptophyta | Magnoliopsida | Fabales | Fabaceae | Melilotus | Melilotus officinalis |
| Eukaryota | Streptophyta | Magnoliopsida | Fabales | Fabaceae | Robinia | Robinia pseudoacacia |
| Eukaryota | Streptophyta | Magnoliopsida | Fabales | Fabaceae | Trifolium | Trifolium repens |
| Eukaryota | Streptophyta | Magnoliopsida | Fabales | Fabaceae | Vicia | Vicia villosa |
| Eukaryota | Streptophyta | Magnoliopsida | Lamiales | Lamiaceae | Teucrium | Teucrium mascatense |
| Eukaryota | Streptophyta | Magnoliopsida | Lamiales | Oleaceae | Olea | Olea europaea |
| Eukaryota | Streptophyta | Magnoliopsida | Lamiales | Oleaceae | Priogymnanthus | Priogymnanthus hasslerianus |
| Eukaryota | Streptophyta | Magnoliopsida | Malpighiales | Rhizophoraceae | Rhizophora | Rhizophora x lamarckii |
| Eukaryota | Streptophyta | Magnoliopsida | Malpighiales | Violaceae | Viola | Viola websteri |
| Eukaryota | Streptophyta | Magnoliopsida | Petrosaviales | Petrosaviaceae | Japonolirion | Japonolirion osense |
| Eukaryota | Streptophyta | Magnoliopsida | Poales | Poaceae | Coix | Coix lacryma-jobi |
| Eukaryota | Streptophyta | Magnoliopsida | Poales | Poaceae | Zea | Zea luxurians |
| Eukaryota | Streptophyta | Magnoliopsida | Rosales | Cannabaceae | Pteroceltis | Pteroceltis tatarinowii |
| Eukaryota | Streptophyta | Magnoliopsida | Santalales | Santalaceae | Santalum | Santalum boninense |
| Eukaryota | Streptophyta | Magnoliopsida | Solanales | Solanaceae | Capsicum | Capsicum chacoense |
| Eukaryota | Streptophyta | Magnoliopsida | Solanales | Solanaceae | Solanum | Solanum pennellii |
| Eukaryota | Streptophyta | Pinopsida | | Pinaceae | Pinus | Pinus yunnanensis |
| Eukaryota | Arthropoda | Insecta | Coleoptera | Laemophloeidae | Dysmerus | Dysmerus sp. BYU CO743 |
| Eukaryota | Arthropoda | Insecta | Hymenoptera | Aphelinidae | Encarsia | Encarsia sp. CaNSW2 |
| Eukaryota | Arthropoda | Insecta | Hymenoptera | Eupelmidae | Eupelmus | Eupelmus pedatorius |
| Eukaryota | Arthropoda | Insecta | Lepidoptera | Noctuidae | Autographa | Autographa pulchrina |
| Eukaryota | Arthropoda | Insecta | Lepidoptera | Pyralidae | Plodia | Plodia interpunctella |
| Eukaryota | Ascomycota | Dothideomycetes | Capnodiales | Capnodiaceae | Leptoxyphium | Leptoxyphium fumago |
| Eukaryota | Ascomycota | Eurotiomycetes | Eurotiales | Aspergillaceae | Aspergillus | Aspergillus cristatus |
| Eukaryota | Ascomycota | Eurotiomycetes | Eurotiales | Aspergillaceae | Aspergillus | Aspergillus flavus |
| Eukaryota | Ascomycota | Eurotiomycetes | Onygenales | Ajellomycetaceae | Histoplasma | Histoplasma capsulatum |

| | | | | | | |
|---|---|---|---|---|---|---|
| Eukaryota | Ascomycota | Eurotiomycetes | Onygenales | Ascosphaeraceae | Ascosphaera | Ascosphaera larvis |
| Eukaryota | Ascomycota | Eurotiomycetes | Onygenales | Ascosphaeraceae | Ascosphaera | Ascosphaera subcuticularis |
| Eukaryota | Ascomycota | Eurotiomycetes | Onygenales | | Paracoccidioides | Paracoccidioides brasiliensis |
| Eukaryota | Basidiomycota | Agaricomycetes | Boletales | | | uncultured Boletaceae |
| Eukaryota | Chlorophyta | | | | | uncultured Chlorophyta |
| Eukaryota | Chordata | Actinopteri | Cypriniformes | Danionidae | Danio | Danio rerio |
| Eukaryota | Chordata | Aves | Passeriformes | Turdidae | Erithacus | Erithacus rubecula |
| Eukaryota | Chordata | Mammalia | Carnivora | Canidae | Canis | Canis lupus |
| Eukaryota | Streptophyta | Magnoliopsida | Alismatales | Zosteraceae | Zostera | Zostera marina |
| Eukaryota | Streptophyta | Magnoliopsida | Arecales | Arecaceae | Borassus | Borassus flabellifer |
| Eukaryota | Streptophyta | Magnoliopsida | Asparagales | Amaryllidaceae | Allium | Allium cepa |
| Eukaryota | Streptophyta | Magnoliopsida | Asparagales | Orchidaceae | Dendrobium | Dendrobium huoshanense |
| Eukaryota | Streptophyta | Magnoliopsida | Asparagales | Orchidaceae | Paphiopedilum | Paphiopedilum gratrixianum |
| Eukaryota | Streptophyta | Magnoliopsida | Asterales | Asteraceae | Artemisia | Artemisia capillaris |
| Eukaryota | Streptophyta | Magnoliopsida | Asterales | Asteraceae | Lactuca | Lactuca sativa |
| Eukaryota | Streptophyta | Magnoliopsida | Asterales | Asteraceae | Pseudognaphalium | Pseudognaphalium sandwicensium |
| Eukaryota | Streptophyta | Magnoliopsida | Asterales | Asteraceae | Taraxacum | Taraxacum officinale |
| Eukaryota | Streptophyta | Magnoliopsida | Brassicales | Brassicaceae | Chorispora | Chorispora tenella |
| Eukaryota | Streptophyta | Magnoliopsida | Brassicales | Brassicaceae | Phoenicaulis | Phoenicaulis cheiranthoides |
| Eukaryota | Streptophyta | Magnoliopsida | Brassicales | Brassicaceae | Physaria | Physaria ludoviciana |
| Eukaryota | Streptophyta | Magnoliopsida | Brassicales | Brassicaceae | Sinapis | Sinapis alba |
| Eukaryota | Streptophyta | Magnoliopsida | Brassicales | Brassicaceae | Sisymbrium | Sisymbrium orientale |
| Eukaryota | Streptophyta | Magnoliopsida | Brassicales | Resedaceae | Ochradenus | Ochradenus baccatus |
| Eukaryota | Streptophyta | Magnoliopsida | Caryophyllales | Amaranthaceae | Celosia | Celosia cristata |
| Eukaryota | Streptophyta | Magnoliopsida | Caryophyllales | Ancistrocladaceae | Ancistrocladus | Ancistrocladus tectorius |
| Eukaryota | Streptophyta | Magnoliopsida | Caryophyllales | Asteropeiaceae | Asteropeia | Asteropeia rhopaloides |
| Eukaryota | Streptophyta | Magnoliopsida | Caryophyllales | Cactaceae | Mammillaria | Mammillaria albiflora |
| Eukaryota | Streptophyta | Magnoliopsida | Caryophyllales | Polygonaceae | Fallopia | Fallopia multiflora |
| Eukaryota | Streptophyta | Magnoliopsida | Cornales | Cornaceae | Cornus | Cornus sanguinea |
| Eukaryota | Streptophyta | Magnoliopsida | Cornales | Hydrostachyaceae | Hydrostachys | Hydrostachys polymorpha |
| Eukaryota | Streptophyta | Magnoliopsida | Cucurbitales | Cucurbitaceae | Citrullus | Citrullus lanatus |
| Eukaryota | Streptophyta | Magnoliopsida | Cucurbitales | Cucurbitaceae | Momordica | Momordica charantia |
| Eukaryota | Streptophyta | Magnoliopsida | Ericales | Ericaceae | Vaccinium | Vaccinium macrocarpon |
| Eukaryota | Streptophyta | Magnoliopsida | Ericales | Pentaphylacaceae | Euryodendron | Euryodendron excelsum |
| Eukaryota | Streptophyta | Magnoliopsida | Fabales | Fabaceae | Caragana | Caragana kozlowii |
| Eukaryota | Streptophyta | Magnoliopsida | Fabales | Fabaceae | Faidherbia | Faidherbia albida |
| Eukaryota | Streptophyta | Magnoliopsida | Fabales | Fabaceae | Hedysarum | Hedysarum semenovii |
| Eukaryota | Streptophyta | Magnoliopsida | Fabales | Fabaceae | Lupinus | Lupinus angustifolius |
| Eukaryota | Streptophyta | Magnoliopsida | Fabales | Fabaceae | Medicago | Medicago orbicularis |
| Eukaryota | Streptophyta | Magnoliopsida | Fabales | Fabaceae | Medicago | Medicago papillosa |

| | | | | | | |
|---|---|---|---|---|---|---|
| Eukaryota | Streptophyta | Magnoliopsida | Fabales | Fabaceae | Sarcodum | Sarcodum scandens |
| Eukaryota | Streptophyta | Magnoliopsida | Fabales | Fabaceae | Trifolium | Trifolium pratense |
| Eukaryota | Streptophyta | Magnoliopsida | Fabales | Fabaceae | Trifolium | Trifolium resupinatum |
| Eukaryota | Streptophyta | Magnoliopsida | Fabales | Fabaceae | Vachellia | Vachellia nilotica |
| Eukaryota | Streptophyta | Magnoliopsida | Fabales | Fabaceae | Vicia | Vicia sativa |
| Eukaryota | Streptophyta | Magnoliopsida | Fabales | Fabaceae | Wisteria | Wisteria sinensis |
| Eukaryota | Streptophyta | Magnoliopsida | Fagales | Betulaceae | Corylus | Corylus avellana |
| Eukaryota | Streptophyta | Magnoliopsida | Fagales | Fagaceae | Quercus | Quercus bawanglingensis |
| Eukaryota | Streptophyta | Magnoliopsida | Gentianales | Apocynaceae | Vincetoxicum | Vincetoxicum shaanxiense |
| Eukaryota | Streptophyta | Magnoliopsida | Gentianales | Rubiaceae | Coffea | Coffea arabica |
| Eukaryota | Streptophyta | Magnoliopsida | Gentianales | Rubiaceae | Rondeletia | Rondeletia odorata |
| Eukaryota | Streptophyta | Magnoliopsida | Gentianales | Rubiaceae | Theligonum | Theligonum cynocrambe |
| Eukaryota | Streptophyta | Magnoliopsida | Lamiales | Lamiaceae | Anisomeles | Anisomeles indica |
| Eukaryota | Streptophyta | Magnoliopsida | Lamiales | Lamiaceae | Salvia | Salvia przewalskii |
| Eukaryota | Streptophyta | Magnoliopsida | Lamiales | Lamiaceae | Tectona | Tectona grandis |
| Eukaryota | Streptophyta | Magnoliopsida | Malpighiales | Euphorbiaceae | Ricinus | Ricinus communis |
| Eukaryota | Streptophyta | Magnoliopsida | Malpighiales | Picrodendraceae | Podocalyx | Podocalyx loranthoides |
| Eukaryota | Streptophyta | Magnoliopsida | Malpighiales | Violaceae | Viola | Viola pedatifida |
| Eukaryota | Streptophyta | Magnoliopsida | Malvales | Malvaceae | Malva | Malva sylvestris |
| Eukaryota | Streptophyta | Magnoliopsida | Malvales | Thymelaeaceae | Daphne | Daphne genkwa |
| Eukaryota | Streptophyta | Magnoliopsida | Myrtales | Lythraceae | Cuphea | Cuphea hyssopifolia |
| Eukaryota | Streptophyta | Magnoliopsida | Myrtales | Melastomataceae | Medinilla | Medinilla magnifica |
| Eukaryota | Streptophyta | Magnoliopsida | Myrtales | Onagraceae | Epilobium | Epilobium ulleungensis |
| Eukaryota | Streptophyta | Magnoliopsida | Poales | Eriocaulaceae | Syngonanthus | Syngonanthus chrysanthus |
| Eukaryota | Streptophyta | Magnoliopsida | Poales | Poaceae | Oryza | Oryza sativa |
| Eukaryota | Streptophyta | Magnoliopsida | Ranunculales | Lardizabalaceae | Archakebia | Archakebia apetala |
| Eukaryota | Streptophyta | Magnoliopsida | Ranunculales | Menispermaceae | Sinomenium | Sinomenium acutum |
| Eukaryota | Streptophyta | Magnoliopsida | Rosales | Cannabaceae | Cannabis | Cannabis sativa |
| Eukaryota | Streptophyta | Magnoliopsida | Rosales | Cannabaceae | Trema | Trema orientale |
| Eukaryota | Streptophyta | Magnoliopsida | Rosales | Rosaceae | Sorbus | Sorbus setschwanensis |
| Eukaryota | Streptophyta | Magnoliopsida | Rosales | Ulmaceae | Ampelocera | Ampelocera longissima |
| Eukaryota | Streptophyta | Magnoliopsida | Rosales | Urticaceae | Rousselia | Rousselia humilis |
| Eukaryota | Streptophyta | Magnoliopsida | Solanales | Convolvulaceae | Cressa | Cressa cretica |
| Eukaryota | Streptophyta | Magnoliopsida | Solanales | Convolvulaceae | Cuscuta | Cuscuta mexicana |
| Eukaryota | | | | | | fungal sp. |
| Eukaryota | Arthropoda | Arachnida | Sarcoptiformes | Oppiidae | Medioppia | Medioppia subpectinata |
| Eukaryota | Arthropoda | Arachnida | Trombidiformes | Johnstonianidae | Diplothrombium | Diplothrombium sp. HP-Hyd192 |
| Eukaryota | Arthropoda | Insecta | Coleoptera | Bostrichidae | Rhyzopertha | Rhyzopertha dominica |
| Eukaryota | Arthropoda | Insecta | Coleoptera | Chrysomelidae | Oreina | Oreina speciosissima |
| Eukaryota | Arthropoda | Insecta | Coleoptera | Chrysomelidae | Psylliodes | Psylliodes sp. BMNH 846644 |

| | | | | | | |
|---|---|---|---|---|---|---|
| Eukaryota | Arthropoda | Insecta | Coleoptera | Curculionidae | Exophthalmus | Exophthalmus sp. GZ164 |
| Eukaryota | Arthropoda | Insecta | Coleoptera | Dytiscidae | Copelatus | Copelatus nr. longicornis 1 BM-2008 |
| Eukaryota | Arthropoda | Insecta | Coleoptera | Endomychidae | Beccariola | Beccariola papuensis |
| Eukaryota | Arthropoda | Insecta | Coleoptera | Endomychidae | Trochoideus | Trochoideus boliviensis |
| Eukaryota | Arthropoda | Insecta | Coleoptera | Hydrophilidae | Cymbiodyta | Cymbiodyta sp. SLE0463 |
| Eukaryota | Arthropoda | Insecta | Coleoptera | Hydrophilidae | Cymbiodyta | Cymbiodyta toddi |
| Eukaryota | Arthropoda | Insecta | Coleoptera | Lycidae | Plateros | Plateros sp. UPOL A00493 |
| Eukaryota | Arthropoda | Insecta | Coleoptera | Nitidulidae | Carpophilus | Carpophilus (Ecnomorphus) sp. G39 |
| Eukaryota | Arthropoda | Insecta | Coleoptera | Nitidulidae | Carpophilus | Carpophilus hemipterus |
| Eukaryota | Arthropoda | Insecta | Coleoptera | Nosodendridae | Nosodendron | Nosodendron fasciculare |
| Eukaryota | Arthropoda | Insecta | Coleoptera | Pyrochroidae | Pyrochroa | Pyrochroa serraticornis |
| Eukaryota | Arthropoda | Insecta | Coleoptera | Rentoniidae | Rentonidium | Rentonidium costiventris |
| Eukaryota | Arthropoda | Insecta | Coleoptera | Rhagophthalmidae | Rhagophthalmus | Rhagophthalmus giganteus |
| Eukaryota | Arthropoda | Insecta | Coleoptera | Staphylinidae | Paederus | Paederus fuscipes |
| Eukaryota | Arthropoda | Insecta | Coleoptera | Tenebrionidae | Tribolium | Tribolium audax |
| Eukaryota | Arthropoda | Insecta | Coleoptera | Tenebrionidae | Tribolium | Tribolium brevicornis |
| Eukaryota | Arthropoda | Insecta | Diptera | Agromyzidae | Chromatomyia | Chromatomyia lactuca |
| Eukaryota | Arthropoda | Insecta | Diptera | Drosophilidae | Drosophila | Drosophila virilis |
| Eukaryota | Arthropoda | Insecta | Hemiptera | Nogodinidae | | Lipocalliini sp. JMU-2006 |
| Eukaryota | Arthropoda | Insecta | Hemiptera | Plataspidae | Plataspis | Plataspis coccinelloides |
| Eukaryota | Arthropoda | Insecta | Hemiptera | Veliidae | Paravelia | Paravelia conata |
| Eukaryota | Arthropoda | Insecta | Hymenoptera | Agaonidae | Sycophaga | Sycophaga sp. D2662 |
| Eukaryota | Arthropoda | Insecta | Hymenoptera | Aphelinidae | Coccophagus | Coccophagus rusti |
| Eukaryota | Arthropoda | Insecta | Hymenoptera | Apidae | Euglossa | Euglossa bursigera |
| Eukaryota | Arthropoda | Insecta | Hymenoptera | Apidae | Trichocerapis | Trichocerapis sp. JESJ-2019 |
| Eukaryota | Arthropoda | Insecta | Hymenoptera | Braconidae | Diaeretiella | Diaeretiella rapae |
| Eukaryota | Arthropoda | Insecta | Hymenoptera | Braconidae | Lysiphlebus | Lysiphlebus fabarum |
| Eukaryota | Arthropoda | Insecta | Hymenoptera | Braconidae | Lysiphlebus | Lysiphlebus testaceipes |
| Eukaryota | Arthropoda | Insecta | Hymenoptera | Braconidae | Pauesia | Pauesia silvestris |
| Eukaryota | Arthropoda | Insecta | Hymenoptera | Chalcididae | | Chalcididae sp. LMM-2014 |
| Eukaryota | Arthropoda | Insecta | Hymenoptera | Eulophidae | Aprostocetus | Aprostocetus csokakoensis |
| Eukaryota | Arthropoda | Insecta | Hymenoptera | Eulophidae | Pediobius | Pediobius sp. RDB-1999 |
| Eukaryota | Arthropoda | Insecta | Hymenoptera | Eupelmidae | | Eupelminae sp. 27862 |
| Eukaryota | Arthropoda | Insecta | Hymenoptera | Formicidae | Aenictus | Aenictus sp. GA03 |
| Eukaryota | Arthropoda | Insecta | Hymenoptera | Formicidae | Cerapachys | Cerapachys desposyne |
| Eukaryota | Arthropoda | Insecta | Hymenoptera | Formicidae | Rotastruma | Rotastruma recava |
| Eukaryota | Arthropoda | Insecta | Hymenoptera | Formicidae | Strumigenys | Strumigenys godeffroyi |
| Eukaryota | Arthropoda | Insecta | Hymenoptera | Megachilidae | Coelioxys | Coelioxys pieliana |
| Eukaryota | Arthropoda | Insecta | Hymenoptera | Pteromalidae | Mesopolobus | Mesopolobus verditer |
| Eukaryota | Arthropoda | Insecta | Hymenoptera | Pteromalidae | Pteromalus | Pteromalus puparum |

| | | | | | | |
|---|---|---|---|---|---|---|
| Eukaryota | Arthropoda | Insecta | Hymenoptera | Pteromalidae | Stenoselma | Stenoselma sp. 1317_07 |
| Eukaryota | Arthropoda | Insecta | Hymenoptera | Thynnidae | Thynnus | Thynnus sp. 1 EMP-2008 |
| Eukaryota | Arthropoda | Insecta | Hymenoptera | Torymidae | Palmon | Palmon sp. 4 PJ-2017 |
| Eukaryota | Arthropoda | Insecta | Hymenoptera | Torymidae | Torymus | Torymus sinensis |
| Eukaryota | Arthropoda | Insecta | Hymenoptera | Trichogrammatidae | Zagella | Zagella spirita |
| Eukaryota | Arthropoda | Insecta | Hymenoptera | Trigonalidae | Taeniogonalos | Taeniogonalos gundlachii |
| Eukaryota | Arthropoda | Insecta | Hymenoptera | Vespidae | Vespula | Vespula vulgaris |
| Eukaryota | Arthropoda | Insecta | Hymenoptera | | | Apoidea sp. 0818E1A02 |
| Eukaryota | Arthropoda | Insecta | Lepidoptera | Geometridae | Ennomos | Ennomos fuscantarius |
| Eukaryota | Arthropoda | Insecta | Lepidoptera | Geometridae | Erannis | Erannis defoliaria |
| Eukaryota | Arthropoda | Insecta | Lepidoptera | Noctuidae | Craniophora | Craniophora ligustri |
| Eukaryota | Arthropoda | Insecta | Lepidoptera | Notodontidae | Phalera | Phalera bucephala |
| Eukaryota | Arthropoda | Insecta | Lepidoptera | Nymphalidae | Maniola | Maniola hyperantus |
| Eukaryota | Arthropoda | Insecta | Lepidoptera | Nymphalidae | Melitaea | Melitaea cinxia |
| Eukaryota | Arthropoda | Insecta | Lepidoptera | Pieridae | Leptidea | Leptidea sinapis |
| Eukaryota | Arthropoda | Insecta | Neuroptera | Chrysopidae | Chrysoperla | Chrysoperla carnea |
| Eukaryota | Arthropoda | Insecta | Neuroptera | Chrysopidae | Nipponochrysa | Nipponochrysa moriutii |
| Eukaryota | Arthropoda | Insecta | Siphonaptera | Ctenophthalmidae | Ctenophthalmus | Ctenophthalmus sp. F405 |
| Eukaryota | Arthropoda | Insecta | Thysanoptera | Thripidae | Frankliniella | Frankliniella sp. D87 |
| Eukaryota | Arthropoda | Insecta | Thysanoptera | Thripidae | Limothrips | Limothrips cerealium |
| Eukaryota | Arthropoda | Malacostraca | Decapoda | Palaemonidae | Macrobrachium | Macrobrachium nipponense |
| Eukaryota | Ascomycota | Dothideomycetes | Cladosporiales | Cladosporiaceae | Cladosporium | Cladosporium parasubtilissimum |
| Eukaryota | Ascomycota | Dothideomycetes | Cladosporiales | Cladosporiaceae | Cladosporium | Cladosporium tenuissimum |
| Eukaryota | Ascomycota | Dothideomycetes | Mycosphaerellales | Mycosphaerellaceae | Cercospora | Cercospora tetragoniae |
| Eukaryota | Ascomycota | Dothideomycetes | Mycosphaerellales | Mycosphaerellaceae | Zymoseptoria | Zymoseptoria tritici |
| Eukaryota | Ascomycota | Dothideomycetes | Pleosporales | Pleosporaceae | Alternaria | Alternaria alternata |
| Eukaryota | Ascomycota | Dothideomycetes | Pleosporales | Pleosporaceae | Bipolaris | Bipolaris maydis |
| Eukaryota | Ascomycota | Dothideomycetes | Pleosporales | Pleosporaceae | Curvularia | Curvularia protuberata |
| Eukaryota | Ascomycota | Dothideomycetes | Pleosporales | Pleosporaceae | Curvularia | Curvularia umbiliciformis |
| Eukaryota | Ascomycota | Dothideomycetes | Pleosporales | Pleosporaceae | Decorospora | Decorospora gaudefroyi |
| Eukaryota | Ascomycota | Dothideomycetes | Pleosporales | Pleosporaceae | Stemphylium | Stemphylium sp. |
| Eukaryota | Ascomycota | Dothideomycetes | Venturiales | Venturiaceae | Venturia | Venturia effusa |
| Eukaryota | Ascomycota | Eurotiomycetes | Chaetothyriales | | Sarcinomyces | Sarcinomyces sp. MA 4649 |
| Eukaryota | Ascomycota | Eurotiomycetes | Chaetothyriales | | | Chaetothyriales sp. MCRE4 |
| Eukaryota | Ascomycota | Eurotiomycetes | Eurotiales | Aspergillaceae | Aspergillus | Aspergillus cervinus |
| Eukaryota | Ascomycota | Eurotiomycetes | Eurotiales | Aspergillaceae | Aspergillus | Aspergillus pseudoglaucus |
| Eukaryota | Ascomycota | Eurotiomycetes | Eurotiales | Aspergillaceae | Aspergillus | Aspergillus versicolor |
| Eukaryota | Ascomycota | Eurotiomycetes | Eurotiales | Aspergillaceae | Eurotium | Eurotium sp. |

| | | | | | | |
|---|---|---|---|---|---|---|
| Eukaryota | Ascomycota | Eurotiomycetes | Eurotiales | Aspergillaceae | Penicillium | Penicillium digitatum |
| Eukaryota | Ascomycota | Eurotiomycetes | Eurotiales | Trichocomaceae | Talaromyces | Talaromyces brevis |
| Eukaryota | Ascomycota | Eurotiomycetes | Eurotiales | Trichocomaceae | Talaromyces | Talaromyces radicus |
| Eukaryota | Ascomycota | Eurotiomycetes | Onygenales | Ajellomycetaceae | Emergomyces | Emergomyces europaeus |
| Eukaryota | Ascomycota | Eurotiomycetes | Onygenales | Ascosphaeraceae | Ascosphaera | Ascosphaera callicarpa |
| Eukaryota | Ascomycota | Eurotiomycetes | Onygenales | Ascosphaeraceae | Ascosphaera | Ascosphaera fusiformis |
| Eukaryota | Ascomycota | Eurotiomycetes | Onygenales | Ascosphaeraceae | Ascosphaera | Ascosphaera major |
| Eukaryota | Ascomycota | Eurotiomycetes | Onygenales | Ascosphaeraceae | Ascosphaera | Ascosphaera osmophila |
| Eukaryota | Ascomycota | Eurotiomycetes | Onygenales | Ascosphaeraceae | Ascosphaera | Ascosphaera pollenicola |
| Eukaryota | Ascomycota | Eurotiomycetes | Onygenales | Ascosphaeraceae | Ascosphaera | Ascosphaera variegata |
| Eukaryota | Ascomycota | Eurotiomycetes | Onygenales | Ascosphaeraceae | Ascosphaera | Ascosphaera xerophila |
| Eukaryota | Ascomycota | Eurotiomycetes | Onygenales | Eremascaceae | Eremascus | Eremascus albus |
| Eukaryota | Ascomycota | Eurotiomycetes | Onygenales | Gymnoascaceae | Arachniotus | Arachniotus ruber |
| Eukaryota | Ascomycota | Eurotiomycetes | Onygenales | Onygenaceae | Uncinocarpus | Uncinocarpus reesii |
| Eukaryota | Ascomycota | Eurotiomycetes | Onygenales | | Canomyces | Canomyces reticulatus |
| Eukaryota | Ascomycota | Eurotiomycetes | Verrucariales | Verrucariaceae | Flakea | Flakea papillata |
| Eukaryota | Ascomycota | Lecanoromycetes | Lecanorales | Lecanoraceae | Rhizoplaca | Rhizoplaca novomexicana |
| Eukaryota | Ascomycota | Lecanoromycetes | Lecanorales | Ramalinaceae | Ramalina | Ramalina intermedia |
| Eukaryota | Ascomycota | Lecanoromycetes | Ostropales | Gyalectaceae | Neopetractis | Neopetractis nodispora |
| Eukaryota | Ascomycota | Lecanoromycetes | Teloschistales | Teloschistaceae | Huriella | Huriella flakusii |
| Eukaryota | Ascomycota | Leotiomycetes | Erysiphales | Erysiphaceae | Blumeria | Blumeria graminis |
| Eukaryota | Ascomycota | Orbiliomycetes | Orbiliales | Orbiliaceae | Lilapila | Lilapila oculisporella |
| Eukaryota | Ascomycota | Orbiliomycetes | Orbiliales | Orbiliaceae | Orbilia | Orbilia aprilis |
| Eukaryota | Ascomycota | Saccharomycetes | Saccharomycetales | Saccharomycetaceae | Torulaspora | Torulaspora delbrueckii |
| Eukaryota | Ascomycota | Saccharomycetes | Saccharomycetales | Saccharomycetaceae | Zygosaccharomyces | Zygosaccharomyces sp. |
| Eukaryota | Ascomycota | Sordariomycetes | Ophiostomatales | Ophiostomataceae | Ophiostoma | Ophiostoma novo-ulmi |
| Eukaryota | Bacillariophyta | Bacillariophyceae | Naviculales | Phaeodactylaceae | Phaeodactylum | Phaeodactylum tricornutum |
| Eukaryota | Bacillariophyta | Coscinodiscophyceae | Melosirales | Melosiraceae | Melosira | Melosira varians |
| Eukaryota | Basidiomycota | Malasseziomycetes | Malasseziales | Malasseziaceae | Malassezia | Malassezia restricta |
| Eukaryota | Basidiomycota | Moniliellomycetes | Moniliellales | Moniliellaceae | Moniliella | Moniliella sp. |
| Eukaryota | Chlorophyta | Chlorophyceae | Chlamydomonadales | Chlamydomonadaceae | Chlamydomonas | Chlamydomonas globosa |
| Eukaryota | Chlorophyta | Chlorophyceae | Chlamydomonadales | Chlorococcaceae | Chlorococcum | Chlorococcum tatrense |
| Eukaryota | Chlorophyta | Pedinophyceae | Pedinomonadales | Pedinomonadaceae | Pedinomonas | Pedinomonas tuberculata |
| Eukaryota | Chlorophyta | Trebouxiophyceae | Chlorellales | Chlorellaceae | Pseudochlorella | Pseudochlorella pringsheimii |
| Eukaryota | Chlorophyta | Trebouxiophyceae | Prasiolales | Prasiolaceae | Prasiola | Prasiola crispa |
| Eukaryota | Chlorophyta | Trebouxiophyceae | Trebouxiales | Trebouxiaceae | Lobosphaera | Lobosphaera incisa |
| Eukaryota | Chlorophyta | Trebouxiophyceae | Trebouxiales | Trebouxiaceae | Symbiochloris | Symbiochloris handae |

| | | | | | | |
|---|---|---|---|---|---|---|
| Eukaryota | Chlorophyta | Ulvophyceae | Ulvales | Kornmanniaceae | Pseudendoclonium | Pseudendoclonium sp. SAG 2051 |
| Eukaryota | Chordata | Actinopteri | Anabantiformes | Anabantidae | Anabas | Anabas testudineus |
| Eukaryota | Chordata | Actinopteri | Perciformes | Bovichtidae | Cottoperca | Cottoperca gobio |
| Eukaryota | Chordata | Actinopteri | Perciformes | Channichthyidae | Pseudochaenichthys | Pseudochaenichthys georgianus |
| Eukaryota | Chordata | Actinopteri | Perciformes | Serranidae | Epinephelus | Epinephelus fuscoguttatus |
| Eukaryota | Chordata | Actinopteri | Perciformes | Serranidae | Plectropomus | Plectropomus leopardus |
| Eukaryota | Chordata | Actinopteri | Salmoniformes | Salmonidae | Salmo | Salmo trutta |
| Eukaryota | Chordata | Mammalia | Primates | Hominidae | Homo | Homo sapiens |
| Eukaryota | Chordata | Mammalia | Rodentia | Cricetidae | Onychomys | Onychomys torridus |
| Eukaryota | Chordata | Mammalia | Rodentia | Sciuridae | Sciurus | Sciurus carolinensis |
| Eukaryota | Chytridiomycota | Chytridiomycetes | Rhizophydiales | Rhizophydiaceae | Rhizophydium | Rhizophydium patellarium |
| Eukaryota | Echinodermata | Echinoidea | Camarodonta | Strongylocentrotidae | Strongylocentrotus | Strongylocentrotus purpuratus |
| Eukaryota | Mollusca | Gastropoda | Lepetellida | Fissurellidae | Emarginula | Emarginula sp. MNHN IM 2013-18121 |
| Eukaryota | Mollusca | Gastropoda | Pleurobranchida | Pleurobranchidae | Berthella | Berthella californica |
| Eukaryota | Mucoromycota | Endogonomycetes | Endogonales | Endogonaceae | Jimgerdemannia | Jimgerdemannia flammicorona |
| Eukaryota | Mucoromycota | Glomeromycetes | Diversisporales | Acaulosporaceae | | uncultured Acaulospora |
| Eukaryota | Mucoromycota | Glomeromycetes | Diversisporales | Gigasporaceae | Dentiscutata | Dentiscutata cerradensis |
| Eukaryota | Mucoromycota | Glomeromycetes | Diversisporales | Gigasporaceae | | uncultured Gigasporaceae |
| Eukaryota | Mucoromycota | Glomeromycetes | Glomerales | Claroideoglomeraceae | Claroideoglomus | Claroideoglomus sp. |
| Eukaryota | Mucoromycota | Glomeromycetes | Glomerales | Claroideoglomeraceae | | uncultured Claroideoglomus |
| Eukaryota | Mucoromycota | Glomeromycetes | Paraglomerales | Paraglomeraceae | Paraglomus | Paraglomus sp. |
| Eukaryota | Nematoda | Chromadorea | Rhabditida | Strongyloididae | Strongyloides | Strongyloides venezuelensis |
| Eukaryota | Oomycota | | Albuginales | Albuginaceae | Albugo | Albugo candida |
| Eukaryota | Oomycota | | Peronosporales | Peronosporaceae | Pseudoperonospora | Pseudoperonospora humuli |
| Eukaryota | Oomycota | | Pythiales | Pythiaceae | Phytopythium | Phytopythium vexans |
| Eukaryota | Platyhelminthes | Cestoda | Diphyllobothriidea | Diphyllobothriidae | Spirometra | Spirometra erinaceieuropaei |
| Eukaryota | Platyhelminthes | Rhabditophora | Macrostomida | Macrostomidae | Bradynectes | Bradynectes sterreri |
| Eukaryota | Rhodophyta | Bangiophyceae | Cyanidiales | Cyanidiaceae | Cyanidioschyzon | Cyanidioschyzon merolae |
| Eukaryota | Rhodophyta | Florideophyceae | Ceramiales | Ceramiaceae | Antithamnionella | Antithamnionella ternifolia |
| Eukaryota | Rhodophyta | Florideophyceae | Ceramiales | Dasyaceae | Dasya | Dasya binghamiae |
| Eukaryota | Streptophyta | Bryopsida | Archidiales | Archidiaceae | Archidium | Archidium alternifolium |
| Eukaryota | Streptophyta | Bryopsida | Bryales | Mniaceae | Pohlia | Pohlia nutans |
| Eukaryota | Streptophyta | Haplomitriopsida | Calobryales | Haplomitriaceae | Haplomitrium | Haplomitrium blumei |
| Eukaryota | Streptophyta | Jungermanniopsida | Metzgeriales | Aneuraceae | Aneura | Aneura pinguis |
| Eukaryota | Streptophyta | Lycopodiopsida | Lycopodiales | Lycopodiaceae | Lycopodium | Lycopodium clavatum |
| Eukaryota | Streptophyta | Lycopodiopsida | Selaginellales | Selaginellaceae | Selaginella | Selaginella sanguinolenta |

| | | | | | | |
|---|---|---|---|---|---|---|
| Eukaryota | Streptophyta | Magnoliopsida | Acorales | Acoraceae | Acorus | Acorus calamus |
| Eukaryota | Streptophyta | Magnoliopsida | Acorales | Acoraceae | Acorus | Acorus tatarinowii |
| Eukaryota | Streptophyta | Magnoliopsida | Alismatales | Alismataceae | Alisma | Alisma plantago-aquatica |
| Eukaryota | Streptophyta | Magnoliopsida | Alismatales | Araceae | Epipremnum | Epipremnum aureum |
| Eukaryota | Streptophyta | Magnoliopsida | Alismatales | Araceae | Spirodela | Spirodela intermedia |
| Eukaryota | Streptophyta | Magnoliopsida | Alismatales | Araceae | Spirodela | Spirodela polyrhiza |
| Eukaryota | Streptophyta | Magnoliopsida | Alismatales | Araceae | Symplocarpus | Symplocarpus nipponicus |
| Eukaryota | Streptophyta | Magnoliopsida | Apiales | Apiaceae | Chamaesium | Chamaesium viridiflorum |
| Eukaryota | Streptophyta | Magnoliopsida | Apiales | Apiaceae | Hansenia | Hansenia oviformis |
| Eukaryota | Streptophyta | Magnoliopsida | Apiales | Apiaceae | Sanicula | Sanicula chinensis |
| Eukaryota | Streptophyta | Magnoliopsida | Apiales | Araliaceae | Panax | Panax ginseng |
| Eukaryota | Streptophyta | Magnoliopsida | Apiales | Araliaceae | Panax | Panax notoginseng |
| Eukaryota | Streptophyta | Magnoliopsida | Apiales | Araliaceae | Panax | Panax vietnamensis |
| Eukaryota | Streptophyta | Magnoliopsida | Apiales | Torricelliaceae | Melanophylla | Melanophylla alnifolia |
| Eukaryota | Streptophyta | Magnoliopsida | Apiales | Torricelliaceae | Melanophylla | Melanophylla modestei |
| Eukaryota | Streptophyta | Magnoliopsida | Arecales | Arecaceae | Arenga | Arenga pinnata |
| Eukaryota | Streptophyta | Magnoliopsida | Arecales | Arecaceae | Phoenix | Phoenix dactylifera |
| Eukaryota | Streptophyta | Magnoliopsida | Asparagales | Amaryllidaceae | Allium | Allium ovalifolium |
| Eukaryota | Streptophyta | Magnoliopsida | Asparagales | Asparagaceae | Dracaena | Dracaena terniflora |
| Eukaryota | Streptophyta | Magnoliopsida | Asparagales | Asphodelaceae | Kniphofia | Kniphofia uvaria |
| Eukaryota | Streptophyta | Magnoliopsida | Asparagales | Orchidaceae | Anoectochilus | Anoectochilus emeiensis |
| Eukaryota | Streptophyta | Magnoliopsida | Asparagales | Orchidaceae | Cypripedium | Cypripedium macranthos |
| Eukaryota | Streptophyta | Magnoliopsida | Asparagales | Orchidaceae | Dendrobium | Dendrobium flexicaule |
| Eukaryota | Streptophyta | Magnoliopsida | Asparagales | Orchidaceae | Dendrobium | Dendrobium moniliforme |
| Eukaryota | Streptophyta | Magnoliopsida | Asparagales | Orchidaceae | Epipactis | Epipactis thunbergii |
| Eukaryota | Streptophyta | Magnoliopsida | Asparagales | Orchidaceae | Gastrodia | Gastrodia elata |
| Eukaryota | Streptophyta | Magnoliopsida | Asparagales | Orchidaceae | Hexalectris | Hexalectris warnockii |
| Eukaryota | Streptophyta | Magnoliopsida | Asparagales | Orchidaceae | Oberonia | Oberonia japonica |
| Eukaryota | Streptophyta | Magnoliopsida | Asparagales | Orchidaceae | Platanthera | Platanthera chlorantha |
| Eukaryota | Streptophyta | Magnoliopsida | Asterales | Asteraceae | Ageratum | Ageratum conyzoides |
| Eukaryota | Streptophyta | Magnoliopsida | Asterales | Asteraceae | Cirsium | Cirsium canescens |
| Eukaryota | Streptophyta | Magnoliopsida | Asterales | Asteraceae | Cirsium | Cirsium setidens |
| Eukaryota | Streptophyta | Magnoliopsida | Asterales | Asteraceae | Marshallia | Marshallia legrandii |
| Eukaryota | Streptophyta | Magnoliopsida | Asterales | Asteraceae | Pseudognaphalium | Pseudognaphalium luteoalbum |
| Eukaryota | Streptophyta | Magnoliopsida | Asterales | Asteraceae | Solidago | Solidago missouriensis |
| Eukaryota | Streptophyta | Magnoliopsida | Asterales | Asteraceae | | Atractylodes environmental sample |
| Eukaryota | Streptophyta | Magnoliopsida | Asterales | Campanulaceae | Campanula | Campanula rotundifolia |
| Eukaryota | Streptophyta | Magnoliopsida | Asterales | Goodeniaceae | Brunonia | Brunonia australis |
| Eukaryota | Streptophyta | Magnoliopsida | Austrobaileyales | Schisandraceae | Illicium | Illicium verum |
| Eukaryota | Streptophyta | Magnoliopsida | Austrobaileyales | Schisandraceae | Kadsura | Kadsura coccinea |
| Eukaryota | Streptophyta | Magnoliopsida | Austrobaileyales | Schisandraceae | Schisandra | Schisandra sphenanthera |

| | | | | | | |
|---|---|---|---|---|---|---|
| Eukaryota | Streptophyta | Magnoliopsida | Brassicales | Brassicaceae | Aethionema | Aethionema arabicum |
| Eukaryota | Streptophyta | Magnoliopsida | Brassicales | Brassicaceae | Arabidopsis | Arabidopsis arenosa |
| Eukaryota | Streptophyta | Magnoliopsida | Brassicales | Brassicaceae | Biscutella | Biscutella vincentina |
| Eukaryota | Streptophyta | Magnoliopsida | Brassicales | Brassicaceae | Bunias | Bunias orientalis |
| Eukaryota | Streptophyta | Magnoliopsida | Brassicales | Brassicaceae | Capsella | Capsella bursa-pastoris |
| Eukaryota | Streptophyta | Magnoliopsida | Brassicales | Brassicaceae | Christolea | Christolea crassifolia |
| Eukaryota | Streptophyta | Magnoliopsida | Brassicales | Brassicaceae | Cochlearia | Cochlearia aestuaria |
| Eukaryota | Streptophyta | Magnoliopsida | Brassicales | Brassicaceae | Cochlearia | Cochlearia officinalis |
| Eukaryota | Streptophyta | Magnoliopsida | Brassicales | Brassicaceae | Erysimum | Erysimum angustatum |
| Eukaryota | Streptophyta | Magnoliopsida | Brassicales | Brassicaceae | Eutrema | Eutrema yunnanense |
| Eukaryota | Streptophyta | Magnoliopsida | Brassicales | Brassicaceae | Heldreichia | Heldreichia bupleurifolia |
| Eukaryota | Streptophyta | Magnoliopsida | Brassicales | Brassicaceae | Lunaria | Lunaria annua |
| Eukaryota | Streptophyta | Magnoliopsida | Brassicales | Brassicaceae | Matthiola | Matthiola longipetala |
| Eukaryota | Streptophyta | Magnoliopsida | Brassicales | Brassicaceae | Pachycladon | Pachycladon fastigiatum |
| Eukaryota | Streptophyta | Magnoliopsida | Brassicales | Salvadoraceae | Salvadora | Salvadora angustifolia |
| Eukaryota | Streptophyta | Magnoliopsida | Brassicales | Tropaeolaceae | Tropaeolum | Tropaeolum majus |
| Eukaryota | Streptophyta | Magnoliopsida | Buxales | Buxaceae | Pachysandra | Pachysandra terminalis |
| Eukaryota | Streptophyta | Magnoliopsida | Caryophyllales | Amaranthaceae | Amaranthus | Amaranthus caudatus |
| Eukaryota | Streptophyta | Magnoliopsida | Caryophyllales | Amaranthaceae | Ptilotus | Ptilotus polystachyus |
| Eukaryota | Streptophyta | Magnoliopsida | Caryophyllales | Cactaceae | Carnegiea | Carnegiea gigantea |
| Eukaryota | Streptophyta | Magnoliopsida | Caryophyllales | Cactaceae | Pereskiopsis | Pereskiopsis diguetii |
| Eukaryota | Streptophyta | Magnoliopsida | Caryophyllales | Chenopodiaceae | Beta | Beta vulgaris |
| Eukaryota | Streptophyta | Magnoliopsida | Caryophyllales | Chenopodiaceae | Dysphania | Dysphania botrys |
| Eukaryota | Streptophyta | Magnoliopsida | Caryophyllales | Chenopodiaceae | Salsola | Salsola montana |
| Eukaryota | Streptophyta | Magnoliopsida | Caryophyllales | Chenopodiaceae | Spinacia | Spinacia oleracea |
| Eukaryota | Streptophyta | Magnoliopsida | Caryophyllales | Drosophyllaceae | Drosophyllum | Drosophyllum lusitanicum |
| Eukaryota | Streptophyta | Magnoliopsida | Caryophyllales | Limeaceae | Limeum | Limeum africanum |
| Eukaryota | Streptophyta | Magnoliopsida | Caryophyllales | Molluginaceae | Trigastrotheca | Trigastrotheca stricta |
| Eukaryota | Streptophyta | Magnoliopsida | Caryophyllales | Nepenthaceae | Nepenthes | Nepenthes ventricosa x Nepenthes alata |
| Eukaryota | Streptophyta | Magnoliopsida | Caryophyllales | Nyctaginaceae | Bougainvillea | Bougainvillea spectabilis |
| Eukaryota | Streptophyta | Magnoliopsida | Caryophyllales | Nyctaginaceae | Pisonia | Pisonia aculeata |
| Eukaryota | Streptophyta | Magnoliopsida | Caryophyllales | Plumbaginaceae | Limonium | Limonium aureum |
| Eukaryota | Streptophyta | Magnoliopsida | Caryophyllales | Polygonaceae | Atraphaxis | Atraphaxis bracteata |
| Eukaryota | Streptophyta | Magnoliopsida | Caryophyllales | Polygonaceae | Fagopyrum | Fagopyrum esculentum |
| Eukaryota | Streptophyta | Magnoliopsida | Caryophyllales | Polygonaceae | Persicaria | Persicaria chinensis |
| Eukaryota | Streptophyta | Magnoliopsida | Caryophyllales | Polygonaceae | Rheum | Rheum nobile |
| Eukaryota | Streptophyta | Magnoliopsida | Caryophyllales | Polygonaceae | Rheum | Rheum rhaponticum |
| Eukaryota | Streptophyta | Magnoliopsida | Caryophyllales | Polygonaceae | Rumex | Rumex sanguineus |
| Eukaryota | Streptophyta | Magnoliopsida | Caryophyllales | Tamaricaceae | Myricaria | Myricaria prostrata |
| Eukaryota | Streptophyta | Magnoliopsida | Caryophyllales | Tamaricaceae | Reaumuria | Reaumuria trigyna |
| Eukaryota | Streptophyta | Magnoliopsida | Celastrales | Celastraceae | Monimopetalum | Monimopetalum chinense |

| | | | | | | |
|---|---|---|---|---|---|---|
| Eukaryota | Streptophyta | Magnoliopsida | Chloranthales | Chloranthaceae | Sarcandra | Sarcandra glabra |
| Eukaryota | Streptophyta | Magnoliopsida | Commelinales | Commelinaceae | Tradescantia | Tradescantia pallida |
| Eukaryota | Streptophyta | Magnoliopsida | Commelinales | Pontederiaceae | Pontederia | Pontederia crassipes |
| Eukaryota | Streptophyta | Magnoliopsida | Cornales | Hydrangeaceae | Hydrangea | Hydrangea platyarguta |
| Eukaryota | Streptophyta | Magnoliopsida | Cucurbitales | Apodanthaceae | Pilostyles | Pilostyles hamiltonii |
| Eukaryota | Streptophyta | Magnoliopsida | Cucurbitales | Cucurbitaceae | Gynostemma | Gynostemma pentaphyllum |
| Eukaryota | Streptophyta | Magnoliopsida | Dipsacales | Caprifoliaceae | Valeriana | Valeriana dioica |
| Eukaryota | Streptophyta | Magnoliopsida | Ericales | Balsaminaceae | Impatiens | Impatiens glandulifera |
| Eukaryota | Streptophyta | Magnoliopsida | Ericales | Balsaminaceae | Impatiens | Impatiens stenosepala |
| Eukaryota | Streptophyta | Magnoliopsida | Ericales | Clethraceae | Clethra | Clethra delavayi |
| Eukaryota | Streptophyta | Magnoliopsida | Ericales | Ebenaceae | Diospyros | Diospyros celebica |
| Eukaryota | Streptophyta | Magnoliopsida | Ericales | Ebenaceae | Diospyros | Diospyros kaki |
| Eukaryota | Streptophyta | Magnoliopsida | Ericales | Lecythidaceae | Barringtonia | Barringtonia racemosa |
| Eukaryota | Streptophyta | Magnoliopsida | Ericales | Lecythidaceae | Eschweilera | Eschweilera micrantha |
| Eukaryota | Streptophyta | Magnoliopsida | Ericales | Primulaceae | Ardisia | Ardisia solanacea |
| Eukaryota | Streptophyta | Magnoliopsida | Ericales | Primulaceae | Cyclamen | Cyclamen purpurascens |
| Eukaryota | Streptophyta | Magnoliopsida | Ericales | Theaceae | Camellia | Camellia japonica |
| Eukaryota | Streptophyta | Magnoliopsida | Ericales | Theaceae | Camellia | Camellia sinensis |
| Eukaryota | Streptophyta | Magnoliopsida | Ericales | Theaceae | Gordonia | Gordonia brandegeei |
| Eukaryota | Streptophyta | Magnoliopsida | Fabales | Fabaceae | Acacia | Acacia jennerae |
| Eukaryota | Streptophyta | Magnoliopsida | Fabales | Fabaceae | Acacia | Acacia longispinea |
| Eukaryota | Streptophyta | Magnoliopsida | Fabales | Fabaceae | Acacia | Acacia puncticulata |
| Eukaryota | Streptophyta | Magnoliopsida | Fabales | Fabaceae | Acacia | Acacia stanleyi |
| Eukaryota | Streptophyta | Magnoliopsida | Fabales | Fabaceae | Aganope | Aganope dinghuensis |
| Eukaryota | Streptophyta | Magnoliopsida | Fabales | Fabaceae | Alhagi | Alhagi sparsifolia |
| Eukaryota | Streptophyta | Magnoliopsida | Fabales | Fabaceae | Ammopiptanthus | Ammopiptanthus nanus |
| Eukaryota | Streptophyta | Magnoliopsida | Fabales | Fabaceae | Angylocalyx | Angylocalyx braunii |
| Eukaryota | Streptophyta | Magnoliopsida | Fabales | Fabaceae | Apios | Apios americana |
| Eukaryota | Streptophyta | Magnoliopsida | Fabales | Fabaceae | Astragalus | Astragalus canadensis |
| Eukaryota | Streptophyta | Magnoliopsida | Fabales | Fabaceae | Astragalus | Astragalus crassicarpus |
| Eukaryota | Streptophyta | Magnoliopsida | Fabales | Fabaceae | Astragalus | Astragalus strictus |
| Eukaryota | Streptophyta | Magnoliopsida | Fabales | Fabaceae | Caragana | Caragana rosea |
| Eukaryota | Streptophyta | Magnoliopsida | Fabales | Fabaceae | Colvillea | Colvillea racemosa |
| Eukaryota | Streptophyta | Magnoliopsida | Fabales | Fabaceae | Dalbergia | Dalbergia oliveri |
| Eukaryota | Streptophyta | Magnoliopsida | Fabales | Fabaceae | Daniellia | Daniellia pilosa |
| Eukaryota | Streptophyta | Magnoliopsida | Fabales | Fabaceae | Glycyrrhiza | Glycyrrhiza pallidiflora |
| Eukaryota | Streptophyta | Magnoliopsida | Fabales | Fabaceae | Grona | Grona styracifolia |
| Eukaryota | Streptophyta | Magnoliopsida | Fabales | Fabaceae | Hedysarum | Hedysarum petrovii |
| Eukaryota | Streptophyta | Magnoliopsida | Fabales | Fabaceae | Lathyrus | Lathyrus aphaca |
| Eukaryota | Streptophyta | Magnoliopsida | Fabales | Fabaceae | Lathyrus | Lathyrus clymenum |
| Eukaryota | Streptophyta | Magnoliopsida | Fabales | Fabaceae | Lotus | Lotus broussonetii |

| | | | | | | |
|---|---|---|---|---|---|---|
| Eukaryota | Streptophyta | Magnoliopsida | Fabales | Fabaceae | Lotus | Lotus pedunculatus |
| Eukaryota | Streptophyta | Magnoliopsida | Fabales | Fabaceae | Medicago | Medicago disciformis |
| Eukaryota | Streptophyta | Magnoliopsida | Fabales | Fabaceae | Medicago | Medicago laciniata |
| Eukaryota | Streptophyta | Magnoliopsida | Fabales | Fabaceae | Medicago | Medicago marina |
| Eukaryota | Streptophyta | Magnoliopsida | Fabales | Fabaceae | Medicago | Medicago praecox |
| Eukaryota | Streptophyta | Magnoliopsida | Fabales | Fabaceae | Medicago | Medicago rigiduloides |
| Eukaryota | Streptophyta | Magnoliopsida | Fabales | Fabaceae | Medicago | Medicago sauvagei |
| Eukaryota | Streptophyta | Magnoliopsida | Fabales | Fabaceae | Medicago | Medicago strasseri |
| Eukaryota | Streptophyta | Magnoliopsida | Fabales | Fabaceae | Medicago | Medicago tetraprostrata |
| Eukaryota | Streptophyta | Magnoliopsida | Fabales | Fabaceae | Onobrychis | Onobrychis viciifolia |
| Eukaryota | Streptophyta | Magnoliopsida | Fabales | Fabaceae | Piliostigma | Piliostigma thonningii |
| Eukaryota | Streptophyta | Magnoliopsida | Fabales | Fabaceae | Pisum | Pisum sativum |
| Eukaryota | Streptophyta | Magnoliopsida | Fabales | Fabaceae | Poecilanthe | Poecilanthe parviflora |
| Eukaryota | Streptophyta | Magnoliopsida | Fabales | Fabaceae | Pterocarpus | Pterocarpus santalinus |
| Eukaryota | Streptophyta | Magnoliopsida | Fabales | Fabaceae | Securigera | Securigera varia |
| Eukaryota | Streptophyta | Magnoliopsida | Fabales | Fabaceae | Senna | Senna tora |
| Eukaryota | Streptophyta | Magnoliopsida | Fabales | Fabaceae | Styphnolobium | Styphnolobium japonicum |
| Eukaryota | Streptophyta | Magnoliopsida | Fabales | Fabaceae | Tibetia | Tibetia liangshanensis |
| Eukaryota | Streptophyta | Magnoliopsida | Fabales | Fabaceae | Trifolium | Trifolium alexandrinum |
| Eukaryota | Streptophyta | Magnoliopsida | Fabales | Fabaceae | Trifolium | Trifolium meduseum |
| Eukaryota | Streptophyta | Magnoliopsida | Fabales | Fabaceae | Vicia | Vicia americana |
| Eukaryota | Streptophyta | Magnoliopsida | Fabales | Fabaceae | Vicia | Vicia cracca |
| Eukaryota | Streptophyta | Magnoliopsida | Fabales | Fabaceae | Vicia | Vicia ramuliflora |
| Eukaryota | Streptophyta | Magnoliopsida | Fabales | Fabaceae | Vicia | Vicia sepium |
| Eukaryota | Streptophyta | Magnoliopsida | Fabales | Surianaceae | Cadellia | Cadellia pentastylis |
| Eukaryota | Streptophyta | Magnoliopsida | Fagales | Fagaceae | Castanopsis | Castanopsis tibetana |
| Eukaryota | Streptophyta | Magnoliopsida | Gentianales | Apocynaceae | Asclepias | Asclepias syriaca |
| Eukaryota | Streptophyta | Magnoliopsida | Gentianales | Apocynaceae | Catharanthus | Catharanthus roseus |
| Eukaryota | Streptophyta | Magnoliopsida | Gentianales | Apocynaceae | Chonemorpha | Chonemorpha megacalyx |
| Eukaryota | Streptophyta | Magnoliopsida | Gentianales | Apocynaceae | Hoya | Hoya hainanensis |
| Eukaryota | Streptophyta | Magnoliopsida | Gentianales | Apocynaceae | Lepiniopsis | Lepiniopsis trilocularis |
| Eukaryota | Streptophyta | Magnoliopsida | Gentianales | Apocynaceae | Strempeliopsis | Strempeliopsis strempelioides |
| Eukaryota | Streptophyta | Magnoliopsida | Gentianales | Gentianaceae | Gentiana | Gentiana apiata |
| Eukaryota | Streptophyta | Magnoliopsida | Gentianales | Gentianaceae | Gentiana | Gentiana lhassica |
| Eukaryota | Streptophyta | Magnoliopsida | Gentianales | Gentianaceae | Gentiana | Gentiana tongolensis |
| Eukaryota | Streptophyta | Magnoliopsida | Gentianales | Gentianaceae | Halenia | Halenia elliptica |
| Eukaryota | Streptophyta | Magnoliopsida | Gentianales | Rubiaceae | Coffea | Coffea canephora |
| Eukaryota | Streptophyta | Magnoliopsida | Gentianales | Rubiaceae | Coptosapelta | Coptosapelta flavescens |
| Eukaryota | Streptophyta | Magnoliopsida | Gentianales | Rubiaceae | Hedyotis | Hedyotis ovata |
| Eukaryota | Streptophyta | Magnoliopsida | Geraniales | Francoaceae | Viviania | Viviania marifolia |
| Eukaryota | Streptophyta | Magnoliopsida | Lamiales | Bignoniaceae | Amphilophium | Amphilophium dusenianum |

| | | | | | | |
|---|---|---|---|---|---|---|
| Eukaryota | Streptophyta | Magnoliopsida | Lamiales | Bignoniaceae | Oroxylum | Oroxylum indicum |
| Eukaryota | Streptophyta | Magnoliopsida | Lamiales | Bignoniaceae | Podranea | Podranea ricasoliana |
| Eukaryota | Streptophyta | Magnoliopsida | Lamiales | Bignoniaceae | Spathodea | Spathodea campanulata |
| Eukaryota | Streptophyta | Magnoliopsida | Lamiales | Bignoniaceae | Tecomaria | Tecomaria capensis |
| Eukaryota | Streptophyta | Magnoliopsida | Lamiales | Gesneriaceae | Oreocharis | Oreocharis mileensis |
| Eukaryota | Streptophyta | Magnoliopsida | Lamiales | Lamiaceae | Colquhounia | Colquhounia coccinea |
| Eukaryota | Streptophyta | Magnoliopsida | Lamiales | Lamiaceae | Salvia | Salvia hispanica |
| Eukaryota | Streptophyta | Magnoliopsida | Lamiales | Lamiaceae | Salvia | Salvia miltiorrhiza |
| Eukaryota | Streptophyta | Magnoliopsida | Lamiales | Lentibulariaceae | Utricularia | Utricularia amethystina |
| Eukaryota | Streptophyta | Magnoliopsida | Lamiales | Linderniaceae | Torenia | Torenia benthamiana |
| Eukaryota | Streptophyta | Magnoliopsida | Lamiales | Oleaceae | Chionanthus | Chionanthus axillaris |
| Eukaryota | Streptophyta | Magnoliopsida | Lamiales | Oleaceae | Chionanthus | Chionanthus ligustrinus |
| Eukaryota | Streptophyta | Magnoliopsida | Lamiales | Oleaceae | Chionanthus | Chionanthus parkinsonii |
| Eukaryota | Streptophyta | Magnoliopsida | Lamiales | Oleaceae | Nestegis | Nestegis sandwicensis |
| Eukaryota | Streptophyta | Magnoliopsida | Lamiales | Orobanchaceae | Aphyllon | Aphyllon epigalium |
| Eukaryota | Streptophyta | Magnoliopsida | Lamiales | Orobanchaceae | Aphyllon | Aphyllon fasciculatum |
| Eukaryota | Streptophyta | Magnoliopsida | Lamiales | Orobanchaceae | Aphyllon | Aphyllon uniflorum |
| Eukaryota | Streptophyta | Magnoliopsida | Lamiales | Orobanchaceae | Cistanche | Cistanche deserticola |
| Eukaryota | Streptophyta | Magnoliopsida | Lamiales | Orobanchaceae | Euphrasia | Euphrasia regelii |
| Eukaryota | Streptophyta | Magnoliopsida | Lamiales | Orobanchaceae | Phelipanche | Phelipanche ramosa |
| Eukaryota | Streptophyta | Magnoliopsida | Lamiales | Plantaginaceae | Veronica | Veronica nakaiana |
| Eukaryota | Streptophyta | Magnoliopsida | Lamiales | Plantaginaceae | Veronica | Veronica persica |
| Eukaryota | Streptophyta | Magnoliopsida | Lamiales | Scrophulariaceae | Myoporum | Myoporum bontioides |
| Eukaryota | Streptophyta | Magnoliopsida | Lamiales | Scrophulariaceae | Verbascum | Verbascum chinense |
| Eukaryota | Streptophyta | Magnoliopsida | Lamiales | Verbenaceae | Duranta | Duranta erecta |
| Eukaryota | Streptophyta | Magnoliopsida | Liliales | Alstroemeriaceae | Bomarea | Bomarea edulis |
| Eukaryota | Streptophyta | Magnoliopsida | Liliales | Liliaceae | Fritillaria | Fritillaria anhuiensis |
| Eukaryota | Streptophyta | Magnoliopsida | Liliales | Liliaceae | Lilium | Lilium tsingtauense |
| Eukaryota | Streptophyta | Magnoliopsida | Magnoliales | Myristicaceae | Myristica | Myristica yunnanensis |
| Eukaryota | Streptophyta | Magnoliopsida | Malpighiales | Achariaceae | Hydnocarpus | Hydnocarpus hainanensis |
| Eukaryota | Streptophyta | Magnoliopsida | Malpighiales | Erythroxylaceae | Erythroxylum | Erythroxylum novogranatense |
| Eukaryota | Streptophyta | Magnoliopsida | Malpighiales | Euphorbiaceae | Croton | Croton tiglium |
| Eukaryota | Streptophyta | Magnoliopsida | Malpighiales | Euphorbiaceae | Triadica | Triadica sebifera |
| Eukaryota | Streptophyta | Magnoliopsida | Malpighiales | Euphroniaceae | Euphronia | Euphronia guianensis |
| Eukaryota | Streptophyta | Magnoliopsida | Malpighiales | Malpighiaceae | Banisteriopsis | Banisteriopsis caapi |
| Eukaryota | Streptophyta | Magnoliopsida | Malpighiales | Passifloraceae | Passiflora | Passiflora affinis |
| Eukaryota | Streptophyta | Magnoliopsida | Malpighiales | Passifloraceae | Passiflora | Passiflora cerradensis |
| Eukaryota | Streptophyta | Magnoliopsida | Malpighiales | Passifloraceae | Passiflora | Passiflora vitifolia |
| Eukaryota | Streptophyta | Magnoliopsida | Malpighiales | Phyllanthaceae | Sauropus | Sauropus granulosus |
| Eukaryota | Streptophyta | Magnoliopsida | Malpighiales | Podostemaceae | Marathrum | Marathrum rubrum |
| Eukaryota | Streptophyta | Magnoliopsida | Malpighiales | Salicaceae | Casearia | Casearia decandra |

| | | | | | | |
|---|---|---|---|---|---|---|
| Eukaryota | Streptophyta | Magnoliopsida | Malpighiales | Salicaceae | Salix | Salix paraflabellaris |
| Eukaryota | Streptophyta | Magnoliopsida | Malpighiales | Salicaceae | Salix | Salix suchowensis |
| Eukaryota | Streptophyta | Magnoliopsida | Malpighiales | Salicaceae | Scyphostegia | Scyphostegia borneensis |
| Eukaryota | Streptophyta | Magnoliopsida | Malpighiales | Violaceae | Viola | Viola hirta |
| Eukaryota | Streptophyta | Magnoliopsida | Malvales | Malvaceae | Alcea | Alcea rosea |
| Eukaryota | Streptophyta | Magnoliopsida | Malvales | Malvaceae | Althaea | Althaea officinalis |
| Eukaryota | Streptophyta | Magnoliopsida | Malvales | Malvaceae | Bombax | Bombax ceiba |
| Eukaryota | Streptophyta | Magnoliopsida | Malvales | Malvaceae | Craigia | Craigia yunnanensis |
| Eukaryota | Streptophyta | Magnoliopsida | Malvales | Malvaceae | Cristaria | Cristaria insularis |
| Eukaryota | Streptophyta | Magnoliopsida | Malvales | Malvaceae | Gossypioides | Gossypioides kirkii |
| Eukaryota | Streptophyta | Magnoliopsida | Malvales | Malvaceae | Gossypium | Gossypium hirsutum |
| Eukaryota | Streptophyta | Magnoliopsida | Malvales | Malvaceae | Hibiscus | Hibiscus acetosella |
| Eukaryota | Streptophyta | Magnoliopsida | Malvales | Malvaceae | Hibiscus | Hibiscus cannabinus |
| Eukaryota | Streptophyta | Magnoliopsida | Malvales | Malvaceae | Hibiscus | Hibiscus syriacus |
| Eukaryota | Streptophyta | Magnoliopsida | Malvales | Malvaceae | Hibiscus | Hibiscus taiwanensis |
| Eukaryota | Streptophyta | Magnoliopsida | Malvales | Malvaceae | Malva | Malva verticillata |
| Eukaryota | Streptophyta | Magnoliopsida | Malvales | Malvaceae | Sphaeralcea | Sphaeralcea coccinea |
| Eukaryota | Streptophyta | Magnoliopsida | Malvales | Malvaceae | Theobroma | Theobroma cacao |
| Eukaryota | Streptophyta | Magnoliopsida | Malvales | Thymelaeaceae | Aquilaria | Aquilaria sinensis |
| Eukaryota | Streptophyta | Magnoliopsida | Malvales | Thymelaeaceae | Pimelea | Pimelea aquilonia |
| Eukaryota | Streptophyta | Magnoliopsida | Myrtales | Lythraceae | Sonneratia | Sonneratia apetala |
| Eukaryota | Streptophyta | Magnoliopsida | Myrtales | Melastomataceae | Bredia | Bredia hirsuta |
| Eukaryota | Streptophyta | Magnoliopsida | Myrtales | Melastomataceae | Bredia | Bredia yunnanensis |
| Eukaryota | Streptophyta | Magnoliopsida | Myrtales | Melastomataceae | Osbeckia | Osbeckia stellata |
| Eukaryota | Streptophyta | Magnoliopsida | Myrtales | Onagraceae | Oenothera | Oenothera biennis |
| Eukaryota | Streptophyta | Magnoliopsida | Myrtales | Onagraceae | Oenothera | Oenothera nuttallii |
| Eukaryota | Streptophyta | Magnoliopsida | Pandanales | Pandanaceae | Pandanus | Pandanus utilis |
| Eukaryota | Streptophyta | Magnoliopsida | Petrosaviales | Petrosaviaceae | Petrosavia | Petrosavia stellaris |
| Eukaryota | Streptophyta | Magnoliopsida | Piperales | Aristolochiaceae | Asarum | Asarum macranthum |
| Eukaryota | Streptophyta | Magnoliopsida | Piperales | Piperaceae | Piper | Piper laetispicum |
| Eukaryota | Streptophyta | Magnoliopsida | Poales | Eriocaulaceae | Eriocaulon | Eriocaulon buergerianum |
| Eukaryota | Streptophyta | Magnoliopsida | Poales | Poaceae | Aristida | Aristida glaziovii |
| Eukaryota | Streptophyta | Magnoliopsida | Poales | Poaceae | Hordeum | Hordeum vulgare |
| Eukaryota | Streptophyta | Magnoliopsida | Poales | Poaceae | Kuruna | Kuruna debilis |
| Eukaryota | Streptophyta | Magnoliopsida | Poales | Poaceae | Lolium | Lolium arundinaceum |
| Eukaryota | Streptophyta | Magnoliopsida | Poales | Poaceae | Phalaris | Phalaris arundinacea |
| Eukaryota | Streptophyta | Magnoliopsida | Poales | Poaceae | Tripsacum | Tripsacum dactyloides |
| Eukaryota | Streptophyta | Magnoliopsida | Poales | Poaceae | Triticum | Triticum timopheevii |
| Eukaryota | Streptophyta | Magnoliopsida | Poales | Poaceae | Zea | Zea diploperennis |
| Eukaryota | Streptophyta | Magnoliopsida | Proteales | Platanaceae | Platanus | Platanus occidentalis |
| Eukaryota | Streptophyta | Magnoliopsida | Ranunculales | Berberidaceae | Berberis | Berberis bealei |

| | | | | | | |
|---|---|---|---|---|---|---|
| Eukaryota | Streptophyta | Magnoliopsida | Ranunculales | Berberidaceae | Berberis | Berberis fortunei |
| Eukaryota | Streptophyta | Magnoliopsida | Ranunculales | Menispermaceae | Fibraurea | Fibraurea tinctoria |
| Eukaryota | Streptophyta | Magnoliopsida | Ranunculales | Menispermaceae | Legnephora | Legnephora moorei |
| Eukaryota | Streptophyta | Magnoliopsida | Ranunculales | Menispermaceae | Menispermum | Menispermum dauricum |
| Eukaryota | Streptophyta | Magnoliopsida | Rosales | Barbeyaceae | Barbeya | Barbeya oleoides |
| Eukaryota | Streptophyta | Magnoliopsida | Rosales | Cannabaceae | Gironniera | Gironniera subaequalis |
| Eukaryota | Streptophyta | Magnoliopsida | Rosales | Cannabaceae | Humulus | Humulus japonicus |
| Eukaryota | Streptophyta | Magnoliopsida | Rosales | Rhamnaceae | Rhamnus | Rhamnus cathartica |
| Eukaryota | Streptophyta | Magnoliopsida | Rosales | Rosaceae | Fragaria | Fragaria mandshurica |
| Eukaryota | Streptophyta | Magnoliopsida | Rosales | Rosaceae | Fragaria | Fragaria x ananassa |
| Eukaryota | Streptophyta | Magnoliopsida | Rosales | Rosaceae | Kageneckia | Kageneckia oblonga |
| Eukaryota | Streptophyta | Magnoliopsida | Rosales | Rosaceae | Prunus | Prunus avium |
| Eukaryota | Streptophyta | Magnoliopsida | Rosales | Rosaceae | Pyrus | Pyrus pyrifolia |
| Eukaryota | Streptophyta | Magnoliopsida | Rosales | Ulmaceae | Planera | Planera aquatica |
| Eukaryota | Streptophyta | Magnoliopsida | Rosales | Ulmaceae | Ulmus | Ulmus gaussenii |
| Eukaryota | Streptophyta | Magnoliopsida | Rosales | Ulmaceae | Ulmus | Ulmus uyematsui |
| Eukaryota | Streptophyta | Magnoliopsida | Rosales | Urticaceae | Urera | Urera sp. WangH-201001 |
| Eukaryota | Streptophyta | Magnoliopsida | Rosales | Urticaceae | Urtica | Urtica lobatifolia |
| Eukaryota | Streptophyta | Magnoliopsida | Santalales | Loranthaceae | Elytranthe | Elytranthe albida |
| Eukaryota | Streptophyta | Magnoliopsida | Santalales | Loranthaceae | Spragueanella | Spragueanella rhamnifolia |
| Eukaryota | Streptophyta | Magnoliopsida | Santalales | Olacaceae | Dulacia | Dulacia candida |
| Eukaryota | Streptophyta | Magnoliopsida | Santalales | Schoepfiaceae | Schoepfia | Schoepfia schreberi |
| Eukaryota | Streptophyta | Magnoliopsida | Santalales | Viscaceae | Viscum | Viscum album |
| Eukaryota | Streptophyta | Magnoliopsida | Sapindales | Anacardiaceae | Pistacia | Pistacia chinensis |
| Eukaryota | Streptophyta | Magnoliopsida | Sapindales | Rutaceae | Citrus | Citrus maxima |
| Eukaryota | Streptophyta | Magnoliopsida | Sapindales | Rutaceae | Citrus | Citrus sinensis |
| Eukaryota | Streptophyta | Magnoliopsida | Sapindales | Rutaceae | Flindersia | Flindersia xanthoxyla |
| Eukaryota | Streptophyta | Magnoliopsida | Sapindales | Sapindaceae | Acer | Acer platanoides |
| Eukaryota | Streptophyta | Magnoliopsida | Sapindales | Simaroubaceae | Eurycoma | Eurycoma longifolia |
| Eukaryota | Streptophyta | Magnoliopsida | Saxifragales | Crassulaceae | Crassula | Crassula perforata |
| Eukaryota | Streptophyta | Magnoliopsida | Saxifragales | Crassulaceae | Sinocrassula | Sinocrassula indica |
| Eukaryota | Streptophyta | Magnoliopsida | Saxifragales | Crassulaceae | Umbilicus | Umbilicus rupestris |
| Eukaryota | Streptophyta | Magnoliopsida | Saxifragales | Cynomoriaceae | Cynomorium | Cynomorium coccineum |
| Eukaryota | Streptophyta | Magnoliopsida | Saxifragales | Hamamelidaceae | Rhodoleia | Rhodoleia championii |
| Eukaryota | Streptophyta | Magnoliopsida | Solanales | Convolvulaceae | Cuscuta | Cuscuta australis |
| Eukaryota | Streptophyta | Magnoliopsida | Solanales | Convolvulaceae | Cuscuta | Cuscuta reflexa |
| Eukaryota | Streptophyta | Magnoliopsida | Solanales | Convolvulaceae | Ipomoea | Ipomoea maurandioides |
| Eukaryota | Streptophyta | Magnoliopsida | Solanales | Convolvulaceae | Ipomoea | Ipomoea nervosa |
| Eukaryota | Streptophyta | Magnoliopsida | Solanales | Solanaceae | Physochlaina | Physochlaina orientalis |
| Eukaryota | Streptophyta | Magnoliopsida | Solanales | Solanaceae | | Nicotiana tabacum/Hyoscyamus niger cybrid |

| | | | | | | |
|---|---|---|---|---|---|---|
| Eukaryota | Streptophyta | Magnoliopsida | Trochodendrales | Trochodendraceae | Trochodendron | Trochodendron aralioides |
| Eukaryota | Streptophyta | Magnoliopsida | Zingiberales | Musaceae | Musa | Musa acuminata |
| Eukaryota | Streptophyta | Magnoliopsida | Zingiberales | Musaceae | Musa | Musa schizocarpa |
| Eukaryota | Streptophyta | Magnoliopsida | Zygophyllales | Krameriaceae | Krameria | Krameria lanceolata |
| Eukaryota | Streptophyta | Marchantiopsida | Marchantiales | Marchantiaceae | Marchantia | Marchantia polymorpha |
| Eukaryota | Streptophyta | Pinopsida | | Pinaceae | Abies | Abies beshanzuensis |
| Eukaryota | Streptophyta | Pinopsida | | Pinaceae | Pinus | Pinus contorta |
| Eukaryota | Streptophyta | Polypodiopsida | Marattiales | Marattiaceae | Christensenia | Christensenia aesculifolia |
| Eukaryota | Streptophyta | Polypodiopsida | Polypodiales | Dennstaedtiaceae | Histiopteris | Histiopteris incisa |
| Eukaryota | Streptophyta | Polypodiopsida | Polypodiales | Saccolomataceae | Saccoloma | Saccoloma inaequale |
| Eukaryota | Streptophyta | Polypodiopsida | Salviniales | Salviniaceae | Salvinia | Salvinia cucullata |
| Eukaryota | Streptophyta | Polypodiopsida | | | | Dicksoniaceae environmental sample |
| Eukaryota | Streptophyta | Zygnemophyceae | Desmidiales | Desmidiaceae | Cosmarium | Cosmarium ochthodes |
| Eukaryota | | Choanoflagellata | Craspedida | Salpingoecidae | Salpingoeca | Salpingoeca urceolata |
| Eukaryota | | | | Apusomonadidae | | uncultured Apusomonadidae |
| Eukaryota | | | | | | uncultured soil eukaryote |

**Supplemental Figure S5:** *Megachile rotundata* cell class taxonomic composition

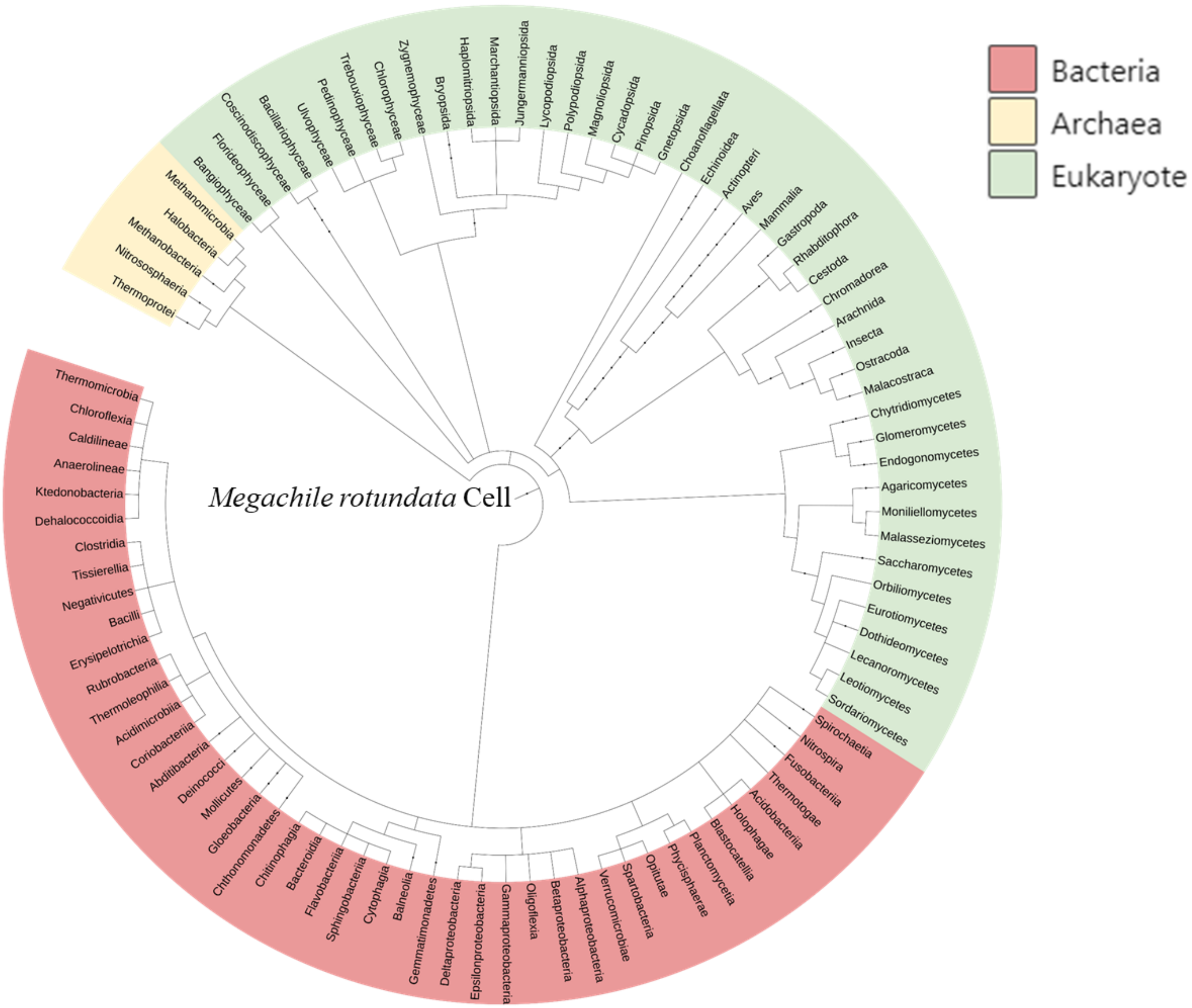


*Unable map the phylum Actinomycetia
